# Mapping Structural Distribution and Gating-Property Impacts of Disease-Associated Missense Mutations in Voltage-Gated Sodium Channels

**DOI:** 10.1101/2023.09.20.558623

**Authors:** Amin Akbari Ahangar, Eslam Elhanafy, Hayden Blanton, Jing Li

**Affiliations:** Department of Biomolecular Sciences, School of Pharmacy, University of Mississippi

## Abstract

Thousands of voltage-gated sodium (Nav) channel variants contribute to a variety of disorders, including epilepsy, autism, cardiac arrhythmia, and pain disorders. Yet variant effects of more mutations remain unclear. The conventional gain-of-function (GoF) or loss-of-function (LoF) classifications is frequently employed to interpret of variant effects on function and guide precision therapy for sodium channelopathies. Our study challenges this binary classification by analyzing 525 mutations associated with 34 diseases across 366 electrophysiology studies, revealing that diseases with similar phenotypic effects can stem from unique molecular mechanisms. Our results show a high biophysical agreement (86%) between homologous disease-associated variants in different Na_v_ genes, significantly surpassing the 60% phenotype (GoF_o_/LoF_o_) agreement among homologous mutants, suggesting the need for more nuanced disease categorization and treatment based on specific gating-property changes. Using UniProt data, we mapped over 2,400 disease-associated missense variants across nine human Nav channels and identified three clusters of mutation hotspots. Our findings indicate that mutations near the selectivity filter generally diminish the maximal current amplitude, while those in the fast inactivation region lean towards a depolarizing shift in half-inactivation voltage in steady-state activation, and mutations in the activation gate commonly enhance persistent current. In contrast to mutations in the PD, those within the VSD exhibit diverse impacts and subtle preferences on channel activity. This study shows great potential to enhance prediction accuracy for variant effects based on the structural context, laying the groundwork for targeted drug design in precision medicine.

## Introduction

Bioelectrical signals, responsible for phenomena such as heartbeats, muscle contractions, and rapid cognitive processing, hinge on the function of ion channels. Among these, voltage-gated sodium (Na_v_) channels are particularly prevalent, playing a crucial role in initiating action potentials and underpinning electrical excitability. Progress in gene research and functional assays has led to the identification of thousands of Na_v_ channel mutations associated with a range of excitability disorders affecting the heart, muscles, and brain^1–5^. For instance, the Na_v_1.5 channel, primarily associated with cardiac function, has been linked to an array of inherited arrhythmias due to a multitude of natural variants^6,7^. Similarly, various forms of periodic paralysis are caused by mutations in the Na_v_1.4 channel, which is primarily found in skeletal muscles ^5,8^. In the realm of neurological disorders, mutations in the Na_v_1.1, Na_v_1.2, Na_v_1.3, and Na_v_1.6 channels, predominantly located in the brain, have been tied to genetic epilepsy, autism, migraines, and other neurological conditions^5,9^. Lastly, dysfunction of the Na_v_1.7 channel, largely situated in peripheral neurons, is implicated in a wide range of pain disorders^10,11^.

Pathogenic mutations can disrupt the standard function of Na_v_ channels by either amplifying (Gain-of-function, or GoF) or diminishing (Loss-of-function, or LoF) Na^+^ current. These opposite effects on channel function can result in a spectrum of disorders ^5,12^. For example, GoF mutations in the Na_v_1.5 channel can trigger long QT syndrome type 3 (LQT3), while LoF mutations in the same channel have been linked to various conditions, such as Brugada syndrome (BRGDA1) and dilated cardiomyopathy (DCM)^6,7^. Similarly, GoF missense mutations in the Na_v_1.7 channel have been found to induce primary erythermalgia (PEM) and paroxysmal extreme pain disorder (PEXPD), while LoF mutations result in an insensitivity to pain ^10^. Likewise, GoF variants in the Na_v_1.2 channel have been associated with conditions like infantile epileptic encephalopathy and benign familial infantile seizures (BFIS3), whereas LoF variants can lead to autism (ASD) and/or intellectual disability ^5,13^. This genotype-phenotype relationship is similarly observed in other Na_v_ channels ^5^.

The GoF and LoF classifications in Na_v_ channels have frequently been employed to categorize disease-associated mutations for interpreting of genotype-phenotype relationships^5,12^, predicting the functional impact of novel variants^14^, and guiding precision therapy^15,16^ for sodium channelopathies. However, several critical questions cannot be addressed by the binary classification of GoF/LoF.

Firstly, the overall gain-of-function (GoF_o_) or loss-of-function (LoF_o_), which refers to the amplification or reduction of the total sodium current respectively, does not provide critical details for the nuanced impacts on channel function. Na_v_ channels undergo three primary steps within their functional cycle: activation, inactivation, and recovery from inactivation^17^. Each of these steps encompasses several gating properties, capable of modifying the Na^+^ current at a specific phase (Fig.1B). The alterations in gating properties associated with these steps not only change the total sodium current, but also modify the timing and shape of the action potential. Such changes in action potential, for instance in cardiac cells, could lead to differential conditions where the heart beats too fast, too slow, or irregularly ^6,7^. These diverse conditions can lead to distinct life-threatening arrhythmias, which cannot be accurately diagnosed or understood solely through the GoF_o_/LoF_o_ classification.

**Figure 1:**
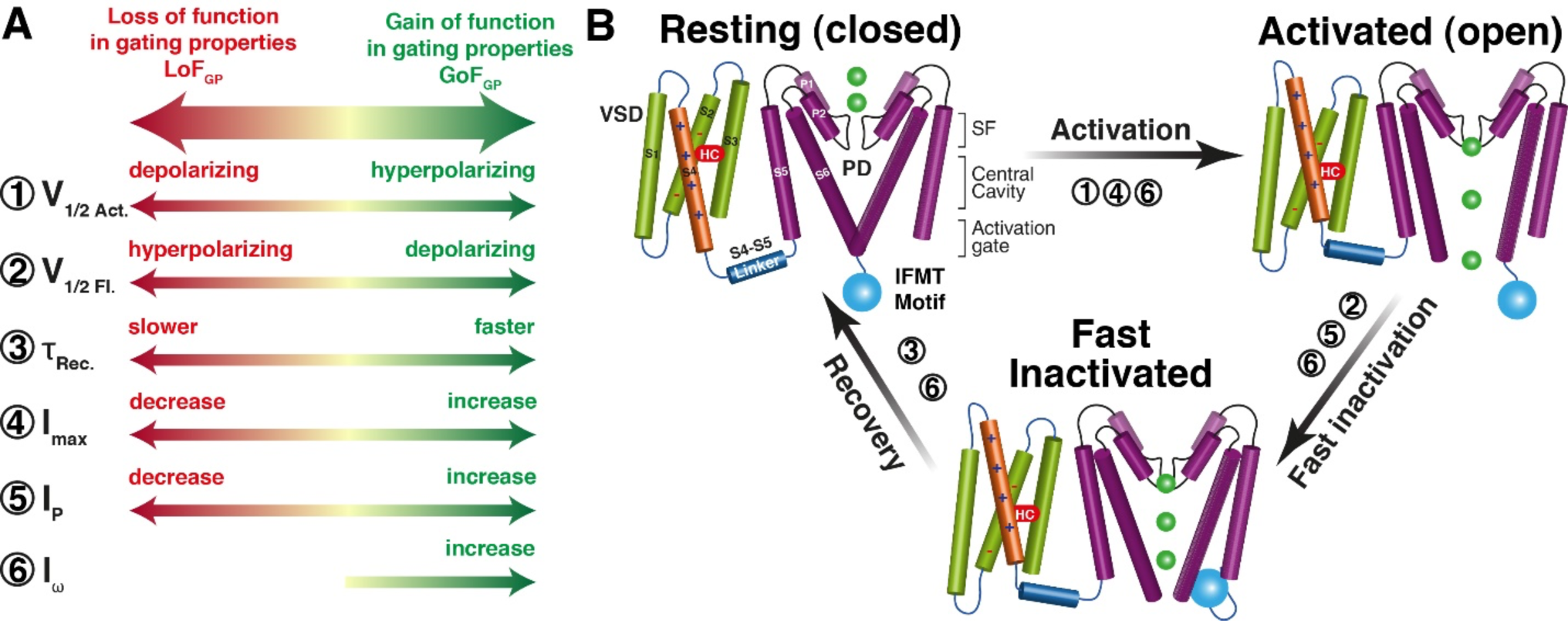
The gating properties and functional transitions of Na_v_ channels. (A) Gating properties are listed with their GoF_GP_ (green) or LoF_GP_ (red) effects. These properties comprised maximal current amplitude (I_max_), half-activation voltage in steady-state activation (V_1/2 Act_), half-inactivation voltage in steady-state fast inactivation (V_1/2 Inact_), recovery rate (τ_rec_), persistent current (I_P_), and gating pore current (or ω current, I_ω_). (B) The gating properties with their relevant transitions in the functional cycle of Na_v_ channels. The Na_v_ structure has four similar subunits (I to IV), and each subunit comprises six transmembrane helices (S1-S6). The first four TMs (S1 to S4) form a voltage-sensing domain (VSD), and the TMs S5 and S6 contribute to the pore domain (PD). Sensing the membrane depolarization, VSDs undergo resting-to-activated structural transition. Then the channel inactivates mediated by allosteric blocking of IFMT motif when depolarization is prolonged to a certain timescale. Thirdly, the repolarization of membrane potential allows recovery from the fast inactivation to the resting state. The gating properties are labeled with their relevant functional step.

Another challenging phenomenon is the occurrence of “overlapping syndromes”, where a single mutation can lead to phenotypes with different or even opposing overall effects ^6,18–20^. For example, more than 30 mutations in Na_v_1.5 have been linked to both GoF_o_-associated LQT3 and LoF_o_ - associated BRGDA1 ^18,21^. The mystery of how a single point mutation can lead to both GoF_o_ and LoF_o_ phenotypes ^22,23^ adds to the limitations of the conventional GoF_o_/LoF_o_ classification system. Understanding these complexities requires a more nuanced examination of Na_v_ channels and their mutations, going beyond the binary GoF_o_/LoF_o_ model.

Moreover, the same phenotype can arise from different gating property changes, mediated by diverse mechanisms^18,19,24,25^. For instance, over 200 LoF_o_ mutations are associated with BRGDA1, but their functional impacts are mediated through a range of mechanisms, such as depolarizing shift of activation, hyperpolarizing shift of inactivation, and/or slower recovery from inactivation^18^. This adds another level of complexity to the application of GoF_o_/LoF_o_ classification for predicting the variant effects and guiding precision therapy.

More importantly, the focus of therapeutic strategies should not be targeting GoF_o_/LoF_o_ effects, but rather be directed toward correcting the altered gating properties induced by mutations. Considering the broad array of mechanisms that lead to GoF_o_/LoF_o_ effects, the development of a universal therapy capable of addressing all similar overall outcomes seems unrealistic. By categorizing these mutations based on their impact on gating properties, it seems more promising to design a select group of drugs to treat a wide array of mutations that share similar mechanisms. In the current study, a large-scale analysis is undertaken, leveraging the most recent electrophysiological and genetic data to map the distribution of mutations and identify common patterns of variant effects on gating properties. Firstly, a large number of research articles have measured the variant effects on different gating properties over the past three decades. To synthesize this wealth of knowledge on variant effects, we conducted a meta-analysis of 366 independent studies on missense mutations based on human cell lines. A systematic literature search was performed to gather the gating properties of 525 mutations from previous electrophysiological measurements (Fig.2A). This meta-analysis enables us to compare homologous mutations across different channels and group mutations or phenotypes based on alterations in gating properties (Fig.3), providing deeper insights into the fundamental principles underlying the disturbed biophysical impacts induced by various mutations. Furthermore, approximately 2,400 annotated disease-associated missense mutations of the nine human Na_v_ channels from the UniProt database (Fig.2B) to equivalent positions are mapped into structural segments (Fig.4) and the multiple sequence alignment (MSA) (Fig.5). we can map mutation distributions, identify the most representative mutations with conserved functional significance (Table S1), and, crucially, explore the way to predict mutational effects based on their structural location and context.

**Figure 2:**
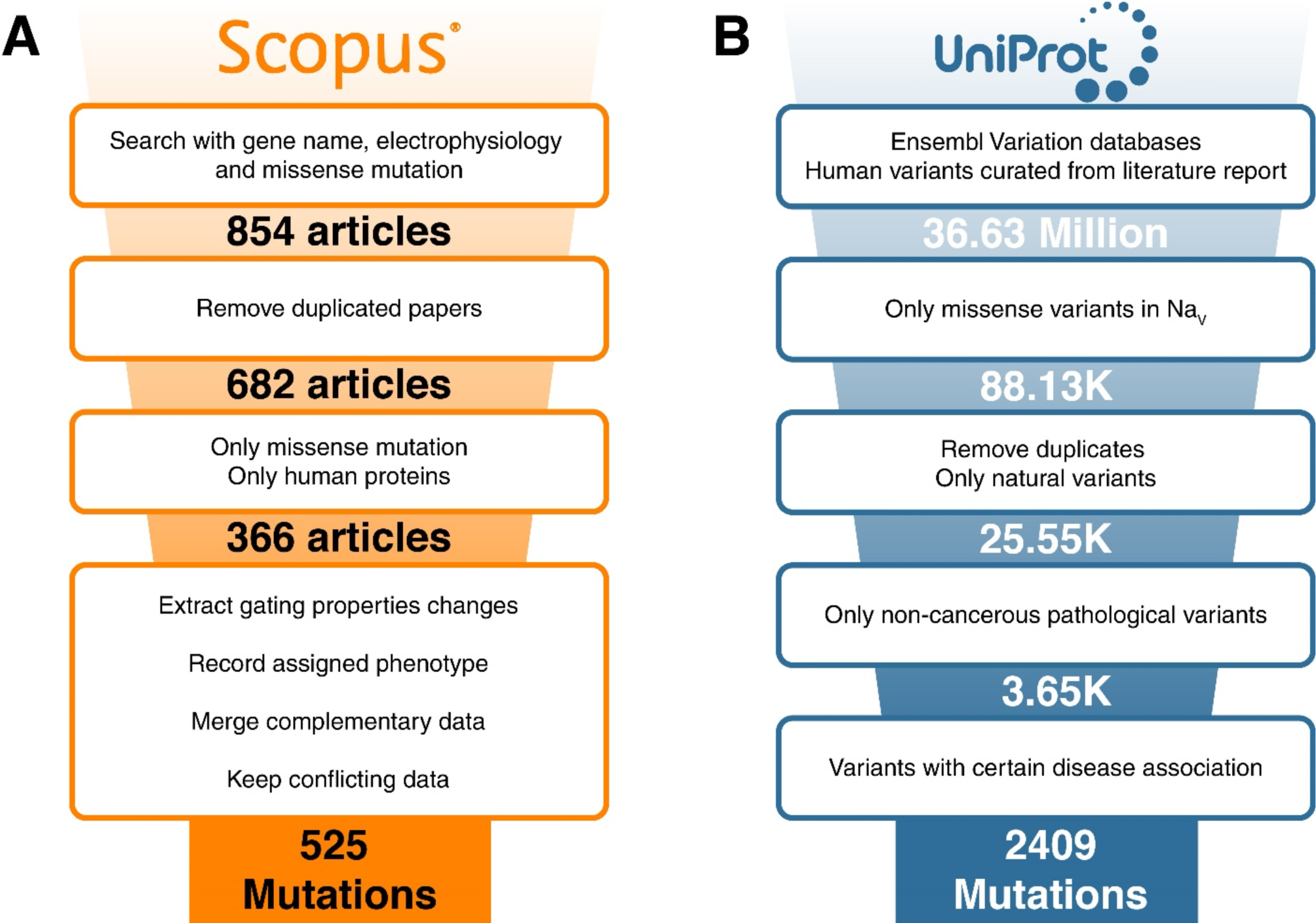
The workflow in the study for collecting electrophysiology and genetic data. (A) The workflow to search and extract electrophysiology research articles that study the mutational effects on gating properties. A total of 854 articles from Scopus were reviewed, and a rigorous selection process identified 366 articles relevant to this study. From these articles, 525 unique mutations with gating properties were identified and selected as the core data for further analysis and investigations. (B) Data extraction steps to retrieve disease and variant data from UniProt. Initially, more than 36 million mutations in UniProt were filtered, resulting in a refined dataset of 2.4K non-cancerous pathogenic missense mutations in Na_v_.

**Figure 3:**
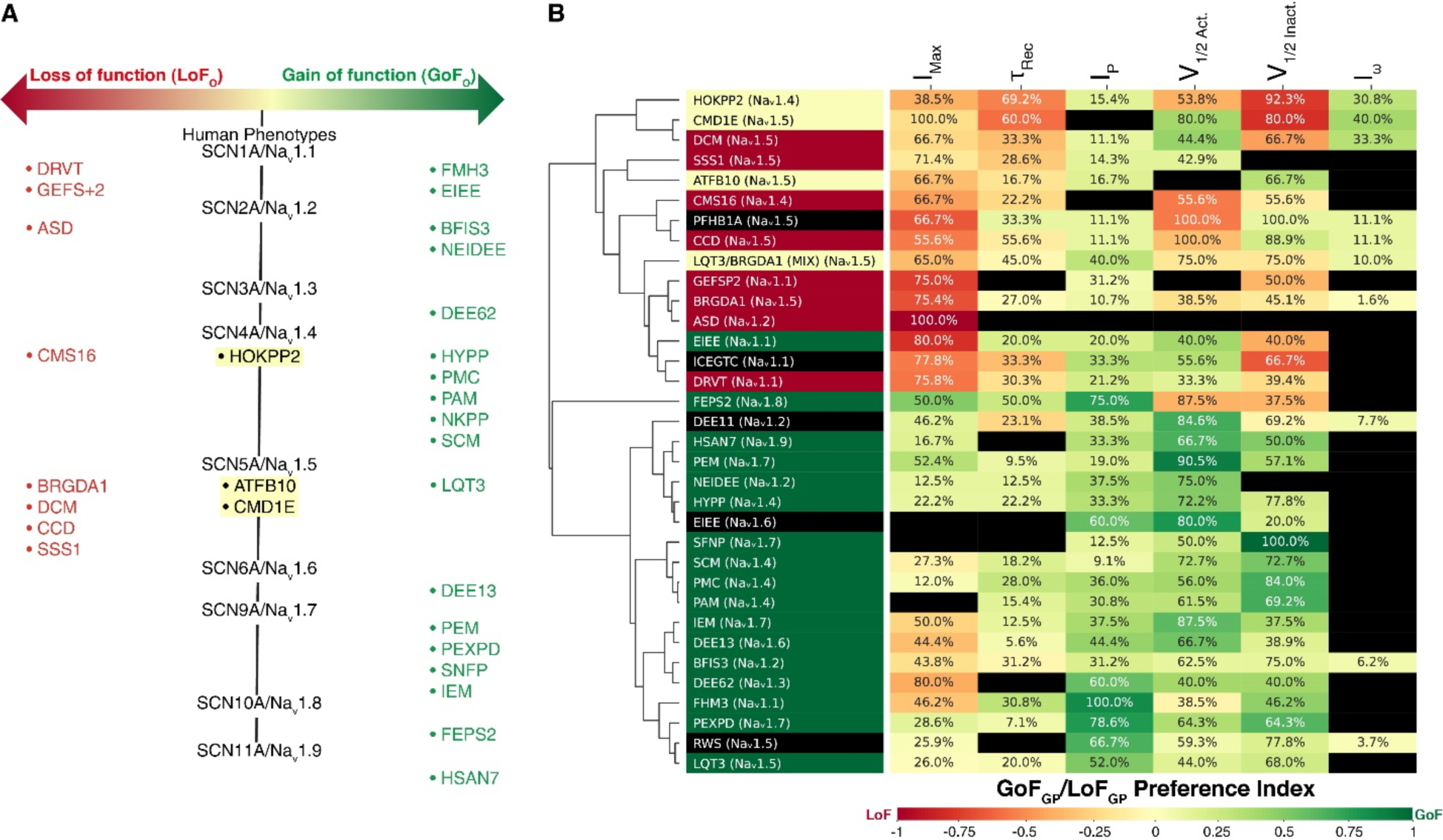
The classification of Na_v_ associated diseases based on overall mutational effect or impacts on gating properties. (A) 34 sodium channelopathies are grouped based on a binary GOF_o_/LOF_o_ classification according to previous literature (Table S2). Diseases are colored in green for GoF_o_ phenotypes, red for LoF_o_ phenotypes, and yellow for diseases with mixed overall effect (MIX_o_). (B) Gating-property impacts of 536 mutations are mapped into their associated 34 diseases. The diseases are also clustered based on the similarity of the gating-property impacts of their associated mutations. The GoF_GP_/LoF_GP_ preference index is depicted colorimetrically with dark green representing highly consistent GoF_GP_ effect, dark red for highly consistent LoF_GP_ effect, and black no such a gating-property data available. The percentage (%) of mutations affecting a certain gating property within a specific phenotype is shown in each grid.

**Figure 4.**
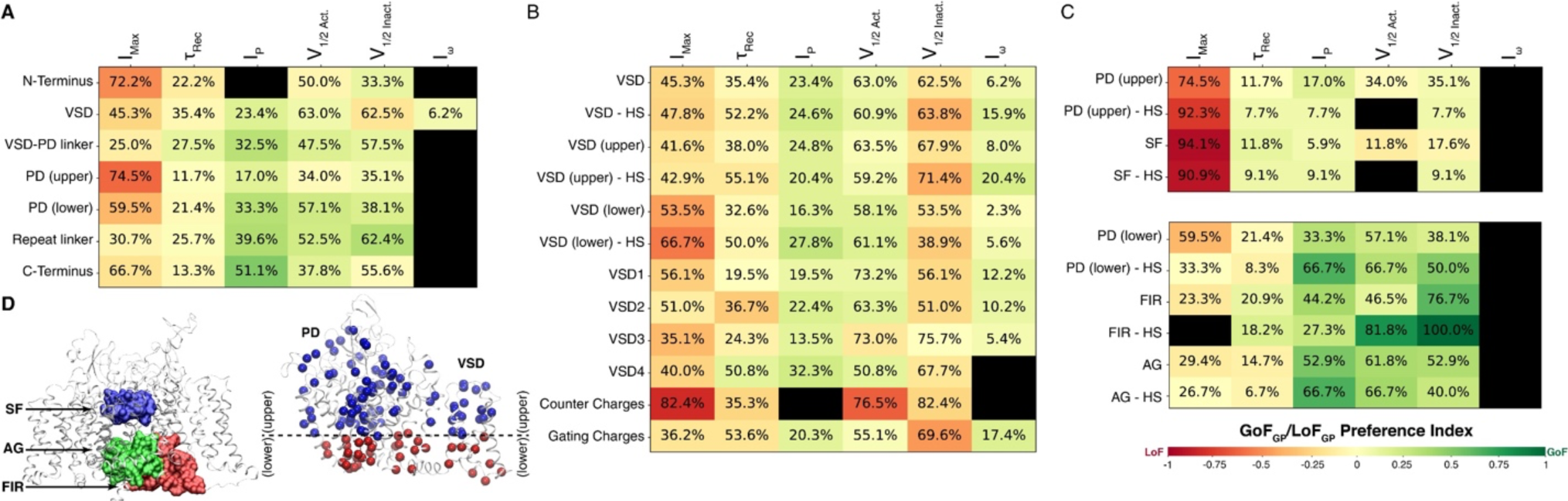
Preferences of gating-property impacts for disease-associated mutations in different structural segments. The variant effects on six gating properties of 525 mutations from 366 papers are mapped into seven major structural segments(A), different selections in VSDs (B), and PD (C). HS stands for the mutation hotspots of corresponding structural segments. The percentage (%) of mutations affecting a certain gating property within a specific structural segment are shown in each grid. GoF_GP_/LoF_GP_ preference index is depicted colorimetrically with dark green representing highly consistent GoF_GP_ effect, dark red for highly consistent LoF_GP_ effect, and black for no such gating-property data available. (D) Shows the selection for selectivity filter (SF), activation gate (AG), fast inactivation region (FIR), as well as the upper and lower part for PD and VSD. The disease-associated mutations are represented in blue (upper) and red (lower) spheres.

**Figure 5:**
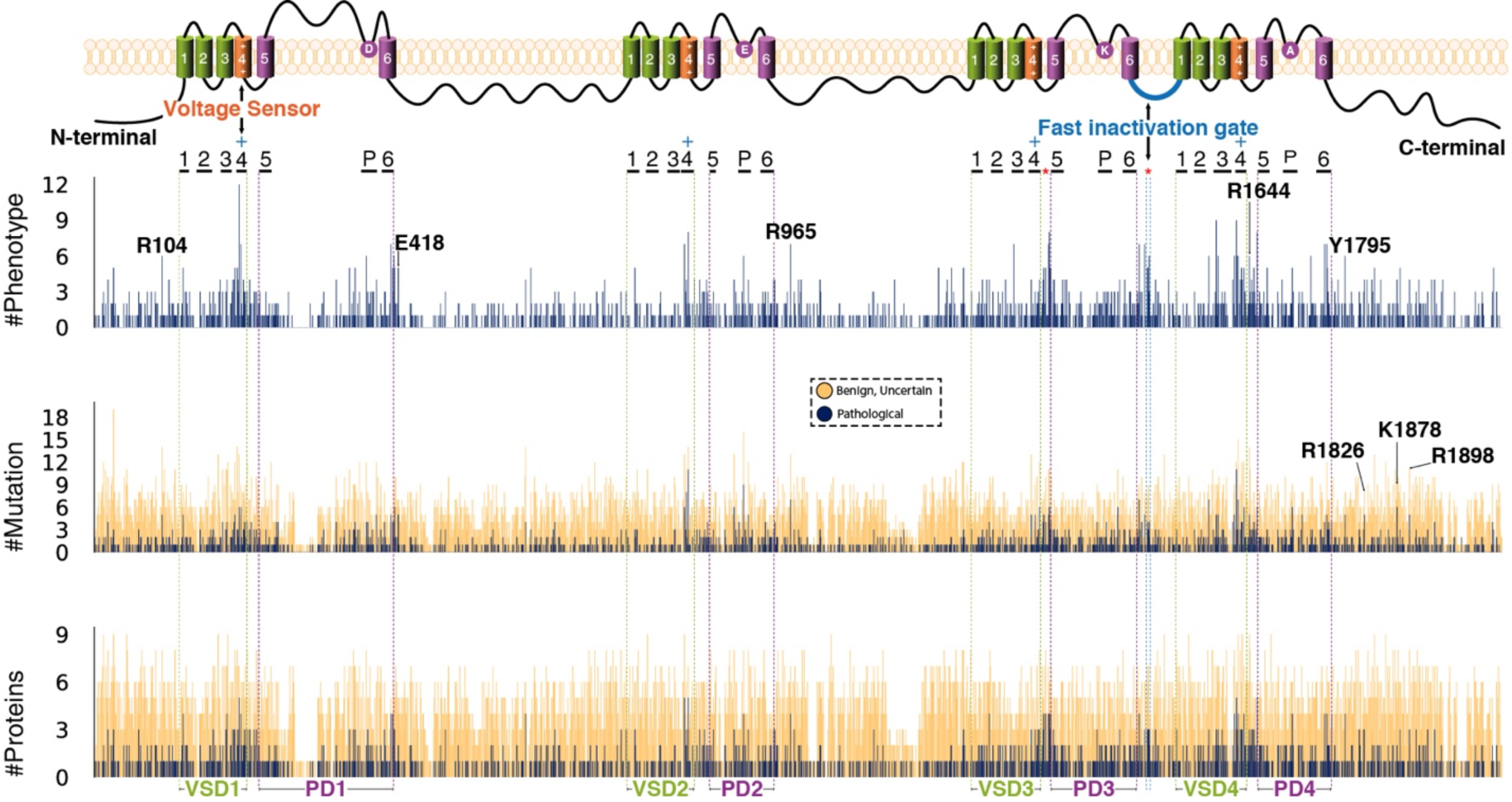
Mapping the mutation hotspots in Na_v_ channels. The annotated disease-associated mutations from UniProt are mapped to equivalent positions of MSA of the 9 human Na_v_ channels and the linear protein structure of the channel. The number of phenotypes (upper panel), the number of mutations (middle panel), and the number of proteins with mutation at the same position (lower panel) are used to determine the mutation hotspots. The data for pathogenic missense mutations are shown in blue bars and all information related to all missense (including pathogenic, benign, and uncertain) variants are shown in orange. Hotspots in the N terminal, C terminal, intracellular, and extracellular loops of the proteins are shown in this figure with residue IDs from Na_v_1.5.

## Materials and methods

## Data gathering and search strategy

In this study, we utilized two major databases to obtain pathogenic mutations in Nav channels and gather functional data related to these mutations.

Firstly, adhering to the PRISMA guidelines^26^, we systematically searched Scopus on March 29th, 2023, to identify English-language studies describing the functional characteristics of missense variants using the following mesh query: ’TITLE-ABS-KEY (SCNXA AND electrophysiology AND mutation) AND (LIMIT-TO (DOCTYPE, “ar”)),’ where X represents the corresponding sodium channel number and “ar” stands for a research article. Furthermore, we conducted a manual search of sodium channel mutation databases and reviewed relevant bibliographies obtained for our search. We specifically selected missense variants whose effects were characterized through whole-cell patch clamp electrophysiology using human Na_v_ channels. Mutations without any functional effects (no changes in biophysical properties) and double mutations were excluded. In cases where contradictory findings existed, duplicated data were considered as additional data points (Table S3). This process led to the selection of unique 525 mutations with their functional data (Fig.2A). Eighty mutations from this dataset were later added to the next dataset to address some missing phenotypes in UniProt (e.g., Autism Spectrum Disorder (ASD) in Na_v_1.2). Three researchers, namely A.A.A., E.E., and H.B., independently reviewed the data.

Secondly, we utilized UniProt’s “Index of Protein Altering Variants” (Release: 2022_03) in conjunction with UniProt’s “Index of human variants curated from literature reports” (Release: 2022_03)^27^ to extract pathogenic mutations within this protein family. We queried the combined database to identify non-cancerous missense mutations with clear OMIM disease assignments across all nine members of Na_v_ channels, including their respective naturally occurring isoforms (Fig.2B). Duplicate mutations resulting in the same phenotype within each family member were excluded. Manual annotations were performed to ensure consistency with OMIM, addressing certain phenotype assignments (e.g., LQTs in Na_v_1.5 to LQT3, PEPD to PEXPD in Na_v_1.7). This process led to the selection of 2409 mutations out of the 36.63 million presented in UniProt. Additionally, we employed a similar workflow to collect benign and uncertain mutations, resulting in a total of 10,348 mutations in this category.

## Data analysis

MSA was conducted using MAFFT^28^ to align all naturally occurring Na_v_ isoforms and was visualized in JalView^29^ 2.11.2.6. Subsequently, the pathogenic mutations obtained from UniProt were mapped onto this alignment to identify recurrent mutations and their respective phenotype in each residue throughout the alignment. The number of mutations, phenotypes, and proteins associated with these variants were tallied and graphed in relation to the MSA. The top 2% of mutations meeting any of the following criteria: more than 6 phenotypes, 5 mutations, or 4 proteins reported from a single residue position were selected as hotspots. Mapping these hotspots in protein sequences and visualizing them in 3D structure was carried out using VMD^30^ and Python 3.10^31^, utilizing Pandas^32^, Numpy^33^, BioPython^34^, and Matplotlib^35^ packages.

Mutations are mapped on different regions to characterize their structural distribution. The selectivity filter region (SF) is defined as including residues within 5Å of the four filter-forming residues (D372, E898, K1419, A1711 ^21^). Similarly, the activation gate region (AG) also includes residues within 5Å of gating residues (A413, L938, I1470, I1771^36^). Fast inactivation region (FIR) is the residues within 5Å of residues (1467 to 1500^36^). The midpoints are used to separate the “upper” and “lower” portions of both PD and VSD. The midpoint is determined by calculating the z-coordinate midpoint of the lowest atom in the SF and the highest atom in the AG, whereas the midpoint of VSDs is the z-coordinate of the C**α** atom of the aromatic residue (either Y or F) in the hydrophobic constriction site in each VSD. The residues in each structural selection were determined using VMD based on a Na_v_1.5 (PDB: 7DTC^21^) structure from the Orientations of Proteins in Membranes (OPM) database ^37^. All residues from other Na_v_ channels are also selected for each protein based on the multiple sequence alignment.

Gating properties altered by variants are documented in the literature, and they were categorized as either gain-of-function on a certain gating property (GoF_GP_) or loss-of-function (LoF_GP_) (Fig.1A). GoF_GP_ effects include an increase in maximal current amplitude (I_max_), an elevation in persistent current (I_P_), a decrease in the numerical values of recovery rate (τ_rec_) from fast inactivation, a hyperpolarizing shift in half-activation voltage in steady-state activation (V_1/2 Act_), a depolarizing shift in half-inactivation voltage (V_1/2 Inact_), and the presence of gating pore current (or ω current, I_ω_). Conversely, LoF_GP_ effects represented opposing effects on the same parameters (except I_ω_) (Fig.1A). Information regarding the phenotypes and the corresponding GoF_GP_ or LoF_GP_ assignment of each mutation, if provided in the literature, was recorded (Fig.2A). These mutations were then mapped onto distinct structural segments of the protein to examine the structural distribution of biophysical property changes (Fig.3). To assess the impact preference of a selected group of mutations (e.g., within a specific structural segment or associated with a certain disease) on a certain gating property, we calculate the GoF_GP_/LoF_GP_ Preference Index (GLPI). The GLPI for a certain gating property *i* is determined by two factors. The first factor is calculated based on the difference between the number of relevant mutations with GoF_GP_ effect (n_GoF_) and those with LoF_GP_ effect (n_LoF_), divided by the total number of mutations within the selected group that affect this gating property *i*. The second factor is the percentage of the mutations impacting gating property *i* within the selected group (*p*_!_), marked as X% in each grid of Figures 3, 4, and 8, was calculated by dividing the number of mutations showing property alterations by the total number of mutations reported for the selected group.

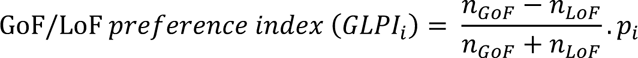

The assignment of phenotypes to specific mutations required meticulous attention and involved cross-referencing various sources such as review articles, research papers, and online databases, including UniProt and OMIM. However, this task presented considerable challenges due to the limitations of manual annotation and lack of widely used standardized disease nomenclature within the field. In this study, UniProt served as the primary source for obtaining phenotype names. In instances where UniProt did not furnish a specific phenotype name, we turned to the original paper that initially reported the mutation. This approach ensured consistency in nomenclature throughout our analysis. The phenotype association dataset for mutations was subsequently clustered into different groups using Seaborn’s clustermap, employing the cosine^38,39^ metric to compute the similarity:

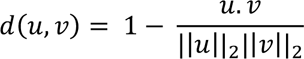

where || ∗ ||_2_ is the 2-norm of its argument *, and *u. v* is the dot product of u and v and average method to assess the relative “closeness” of phenotypes based on the resulting biophysical changes.

## Data availability

All data used in this study are available in the manuscript and the Supplementary material.

## Result

### Mapping the gating-property impacts of sodium channelopathies

We carried out an exhaustive literature search across Scopus, yielding 854 papers that reported electrophysiology measurements of gating properties for Na_v_ channel mutants. Duplicate research articles were eliminated, and the records lacking a patch-clamp experiment on a pathogenic missense mutation of human Na_v_ channels were excluded (Fig.2). Following this, we performed a thorough full-text analysis on 366 articles, documenting 525 variants and their influence on several crucial gating properties. These properties comprised maximal current amplitude (I_max_), half-activation voltage in steady-state activation (V_1/2 Act_), half-inactivation voltage in steady-state fast inactivation (V_1/2 Inact_), recovery rate (τ_rec_), persistent current (I_P_), and gating pore current (or ω current, I_ω_) (Fig.1). In this study, we recognize that alterations in each gating property in all electrophysiology measurements lead to amplification or reduction of the Na^+^ current at specific steps in the functional cycle. To effectively summarize the mutational effects on each gating property from 366 research articles, we have categorized the impacts on each gating property based on gain-of-function or loss-of-function, referred to as GoF_GP_ (Gain of Function on a Gating Property) and LoF_GP_ (Loss of Function on a Gating Property), respectively (Fig.1). This terminology helps to distinguish the effects on individual gating properties from the overall gain-of-function (GoF_o_) or loss-of-function (LoF_o_) effects on Na_v_ channels. This distinction is crucial as it provides a more nuanced understanding of how mutations affect Na_v_ channel function at specific stages, further aiding in understanding mutation impacts more precisely.

To categorize the sodium channelopathies and compare their gating-property impacts, a GoF_o_/LoF_o_ categorization was applied to 34 diseases across the nine Na_v_ channels based on overall effects in previous literature (Table S2). In order for a disease to be included in this analysis, it had to have at least five associated mutants that were reported in previous electrophysiological studies (Table S3). Based on previous literature (Table S2), eight diseases were associated with inferred LoF_o_ effect, eighteen were associated with inferred GoF_o_ effect, three were suggested to have mixed effects, and the effects of five diseases remain unclear (See Fig. 3A). Considering the fact that many mutants in Na_v_1.5 are associated with both LQT3 and BRGDA1, these mutants were exclusively grouped for a mixed phenotype (LQT3/BRGDA1). The mixed phenotype categorization highlights the complex relationships between specific mutations, channel functionality, and resulting diseases.

Sodium channelopathies, as identified in our analysis (Fig.3B), cluster into two primary branches based on their altered gating properties, predominantly aligning with the GoF_o_/LoF_o_ categorization. EIEE(Na_v_1.1) is the sole GoF_o_ phenotype grouped with most LoF_o_ diseases (Fig.3B), yet the overall effect of EIEE(Na_v_1.1) remains a subject of ongoing debate^5^.

However, within each of these branches, there are diseases with significant differences on their altered gating properties. This observation implies that diseases with similar overall effects (either GoF_o_ or LoF_o_) are triggered by distinctive molecular mechanisms. For instance, among GoF_o_ phenotypes, both Familial Episodic Pain Syndrome 2 (FEPS2, associated with Na_v_1.8) and Inherited Erythromelalgia (IEM, associated with Na_v_1.7) belong to this branch and both are pain disorders. However, their variant effects on several gating properties are almost opposite: a large portion of FEPS2 mutations increase I_max_, depolarizing shift V_1/2 Act_, and hyperpolarizing shift V_1/2 Inact_, whereas similar proportions of IEM mutations decrease I_max_, hyperpolarizing shift V_1/2 Act_, and depolarizing shift V_1/2 Inact_. This clear distinction suggests that these two pain disorders are driven by entirely different mechanisms, and as a result, their treatments for restoring normal channel function should also be unique. A similar divergence is observed within the GoF_o_ branch, DCM (associated with Na_v_1.5) mutations cause hyperpolarizing shift of V_1/2 Act_, hyperpolarizing shift V_1/2 Inact_, or slow τ_rec_. In contrast, most PFHB1A (associated with Na_v_1.5) mutations cause depolarizing shift of V_1/2 Act_, lead depolarizing shift of V_1/2 Inact_, or speed up τ_rec_. Given that a large portion of mutations associated with these two diseases affect these three gating properties, it is clear that these two diseases, despite having an LoF_o_ effect, must be driven by distinct mechanisms.

Our analysis shows that the same overall effect (either GoF_o_ or LoF_o_) can be caused by differential influences on different gating properties. This implies that certain gating properties might have a dominant role in shaping the overall effect, even when other gating properties are influenced in an opposite manner. For instance, in IEM (Nav1.7), 87.5% of mutations lead to a shift in V_1/2 Act_, with the majority trending towards a hyperpolarizing direction, producing a GoF_GP_ effect. At the same time, 50% of IEM mutations alter I_max_, with most reducing I_max_, resulting in a LoF_GP_ effect (Fig.3B). These two gating property changes are diametrically opposite, yet the net effect is categorized as GoF_o_. While in FEPS2 (Na_v_1.8), the mutations that alter V_1/2 Act_ and V_1/2 Inact_ lead to a LoF effect. However, a more dominant GoF effect is brought about by stronger impacts on the maximum ionic current (I_max_) and persistent current (I_p_) (Fig.3B). Similarly, in Dilated Cardiomyopathy 1E (CMD1E, Na_v_1.5), 80% of associated mutations prefer a hyperpolarizing shift of V_1/2 Inact_ and 60% of these mutations tend to slow the recovery time constant (τ_rec_). Although the mutational impact on V_1/2 act_ is GoF_GP,_ the alterations on V_1/2 Inact_ and τ_rec_ together bring about an LoF_o_ effect. In PFHB1A (Na_v_1.5), most associated mutations decrease I_max_ and cause a depolarizing shift of V_1/2 Act_, leading to the LoF_o_ effect, despite their GoF_GP_ effects on V_1/2 Inact_. In summary, the overall effect of a disease is not solely determined by the direction (either GoF_GP_ or LoF_GP_) of individual variant effects on gating properties. Instead, it is the cumulative effect of these alterations, with some gating properties potentially playing a more dominant role than others in determining the overall effect.

An additional pattern observed in our analysis is that almost all LoF_o_ diseases show mixed effects on different gating properties (Fig.3B). For instance, BRGDA1(Na_v_1.5) demonstrates not only a robust LoF_GP_ impact on I_max_, and moderate LoF_GP_ effects on V_1/2 Act_, V_1/2 Inact_, and τ_rec_, but also GoF_GP_ impacts on I_P_ and I_ω_. This is likely due to the fact that, as observed in electrophysiological measurements, impacts on I_P_ and I_ω_ always prefer to be GoF_GP_ for all phenotypes (Fig.3B). However, even when I_P_ and I_ω_ are excluded, many LoF_o_ diseases, such as DRVT(Na_v_1.1), ICEGTC (Na_v_1.1), DCM(Na_v_1.5), PHFB1A(Na_v_1.5), CCD(Na_v_1.5), show opposite impacts on V_1/2 Act_ and V_1/2 Inact_. Interestingly, all the diseases showing mixed GoF_GP_/LoF_GP_ effects, including HOKPP2(Na_v_1.4), ATFB10(Na_v_1.5), CMD1E(Na_v_1.5), and LQT3/BRGDA1(Na_v_1.5) are clustered within the LoF_o_ branch (Fig.3B). This reaffirms that mixed effects are a common feature for diseases within the LoF_o_ branch. Conversely, although there are several diseases in the GoF_o_ branch that present mixed effects, more diseases consistently show GoF_GP_ effects on all gating properties (Fig.3B).

In this study, we also compared the variant effects of homologous residues across different Na_v_ channels, focusing on their biophysical impacts on gating properties and their GoF_o_ or LoF_o_ categorizations. Our analysis revealed that the biophysical impacts of mutations in identical residues across different Na_v_ channels were more likely to align, with 37 out of 44 pairs of identical disease-associated mutations in different Na_v_ channels resulting in similar alterations in gating properties (Table S4), equating to 86% biophysical agreement. This observation aligns with prior research indicating that analogous positions in Na_v_ channels may elicit similar biophysical outcomes due to mutations^15^. In contrast, only 60% (74 out of 123) of these mutations showed a consistent GoF_o_/LoF_o_ agreement in their associated diseases (Table S5). There could be multiple reasons for the relatively lower agreement in phenotype classification compared to biophysical impact. Firstly, the predominant gating property influencing the overall effect could vary between different Na_v_ channels, leading to differential GoF_o_/LoF_o_ effects. For instance, the change in a specific gating property might be amplified due to the interaction between this Na_v_ channel and other proteins. Secondly, mutations associated with multiple diseases, encompassing both GoF_o_ or LoF_o_ phenotypes, might only be partially characterized in terms of their disease associations. Insufficient data could result in these variants being labeled as either GoF_o_ or LoF_o_ mutations. Consequently, predicting the impact of a new variant based on the known biophysical effects of equivalent mutations in other Na_v_ channels appears to be a more reliable approach than relying on the documented GoF_o_/LoF_o_ classification of the same mutation.

### Clustering mutations and their biophysical impacts in 3D structure

The biophysical/functional impact of a mutation should highly depend on its structural role, thus, mapping all disease-associated mutations in Nav channels and their impacts on gating properties in 3D structure would help us to understand and predict the mutational effects of the undocumented variants. The recurrence of mutations across independent samples in disease-associated cases is a robust indicator of functional significance ^40,41^. Na_v_ channels share high sequence, structure, and function similarity^42^. Considering mutations in analogous positions could cause similar biophysical effects in Na_v_ channels^14,15^, this evolutionarily conserved nature allows us to extrapolate the concept of recurring mutations from a single gene/protein to the Na_v_ channel family. Based on MSA across nine human Na_v_ channels, we mapped 2,409 annotated missense mutations from the UniProt database^43^ (Fig.2) to their corresponding positions, thus identifying a series of mutation hotspots (Fig.5). These pathogenic mutation hotspots were subsequently visualized within the 3D structure of human Na_v_1.5 (PDB ID:7DTC) to display their distribution. Thus, the residues discussed in the following sections are referred to using their residue ID in Na_v_1.5. Our structural mapping underscores that a significant number of mutation hotspots are predominantly situated within three distinct regions: 1) the voltage-sensing domain (VSD), 2) the upper section of the pore domain (PD (upper)) - near the selectivity filter and pore helices, and 3) the lower part of the pore domain (PD (lower)) - including the fast-inactivation segment and activation gate.

In this comprehensive study, we also mapped the effects of known Na_v_ channel mutations on six gating properties, across respective domains including VSDs, PD (upper), PD (lower), N/C terminals, and additional detailed structural segments (Fig.4). This thorough mapping procedure sheds light on the intricate relationships between the structural locations of mutations and their functional outcomes. While it is evident that mutations within each structural segment can influence most gating properties, no single segment appears to uniformly affect all gating properties in a consistent GoF_GP_ or LoF_GP_ direction. A key observation is that mutations across all structural segments tend to reduce the I_max_ (a LoF_GP_ effect), while also being inclined to increase the persistent current I_P_ and gating pore current I_ω_ (GoF_GP_ effects). This suggests that wild-type (WT) Nav channels have evolved to achieve maximum conductivity in the activated state while minimizing leakage in the inactivated and resting states. Beyond these general patterns, each structural segment showcases unique traits, displaying differential preferences for specific gating properties or exhibiting characteristic influences on channel behavior. These unique attributes may shape the manifestation of associated diseases. Thus, a detailed understanding of these individual segment features is imperative for gaining insight into disease mechanisms, which could be exploited for the prediction of undocumented mutational effects and the development of targeted therapeutic interventions.

### Diverse variant effects in VSDs are highly sensitive to structural context

A major cluster of mutations is located within the VSDs (Fig.3), critical structural components that sense membrane potential and trigger structural transitions of Na_v_ channels. Several features characterize the distribution of mutation hotspots within the VSDs. The first distinguishing feature of VSD mutations is their diverse impacts across a wide array of gating properties. Between one-third and two-thirds of these variants affect key properties such as I_max_, τ_rec_, V_1/2 Act_, and V_1/2 Inact_. As shown in Figure 3, there is no strong preference between GoF_GP_ and LoF_GP_ effects on these gating properties. In addition, only mutations within VSDs can induce gating pore currents, also termed omega-pore currents (Iω) ^44,45^. These currents are produced by protons or cations that pass the channel directly through the VSD ^46–49^. Consistently, VSD variants are implicated in most (32 out of 34) sodium channelopathies (Fig.8), each presenting with GoF_o_, LoF_o_, or mixed GoF_o_/LoF_o_ effect. More intriguingly, some hotspots have been identified as overlap-syndrome mutations that are linked to both GoF_o_ and LoF_o_ diseases. For instance, mutations like R222Q, R225W, and R1623Q in Na_v_1.5 are associated with both LQT3 (GoF_o_) and BRGDA1 (LoF_o_) ^50–52^. Similarly, R225W in Na_v_1.4, an equivalent mutation to R225W in Na_v_1.5, is linked to both Congenital Myasthenic Syndrome (CMS16, LoF_o_) and Sodium Channel Myotonia (SCM, GoF_o_) ^8,53,54^.

The second noteworthy feature is the prevalence of mutation hotspots within the S4 helix, specifically gating charges, i.e., R1635 (R5 in VSD_IV_) (Fig.6D). Na_v_ channel structures consistently illustrate that the structural transition of VSDs is mediated by the sliding of the S4 helix through the remaining portion of this domain (Fig.1B) ^17,55–57^. Arrayed along the S4 helix across the membrane, four to six gating charges (arginine or lysine) named as R1 to R6 from the extracellular to intracellular side, serve as the voltage sensors in VSDs. Our hotspot analysis indicates that mutations of gating charges tend to recur more frequently in diseases compared to non-gating charge mutations, thereby affirming their functional significance. Interestingly, these mutation hotspots are predominantly distributed in R1 to R3, including R808 (R1 in VSD_II_), R1303 (R1 in VSD_III_), R1623 (R1 in VSD_IV_), R222 (R2 in VSD_I_), R811 (R2 in VSD_II_), R1626 (R2 in VSD_IV_), R225 (R3 in VSD_I_), R814 (R3 in VSD_II_), and R1309 (R3 in VSD_III_) (Fig.6). In contrast, fewer hotspots are observed in the gating charges near the intracellular side, with only one hotspot in R5 (R1635 in VSD_IV_) and no hotspot in R4 in any VSD. Additionally, several conserved countercharge residues, like D1274 (VSD_III_) and D1595 (VSD_IV_), are also identified within mutation hotspots. As these countercharges form salt bridges with gating charges, they also contribute to voltage dependence and structural transition ^58^. However, hotspots in countercharges are considerably fewer than those in gating charges, and most countercharge hotspots are situated at the intracellular negatively charged region (INC) (Fig.6).

**Figure 6:**
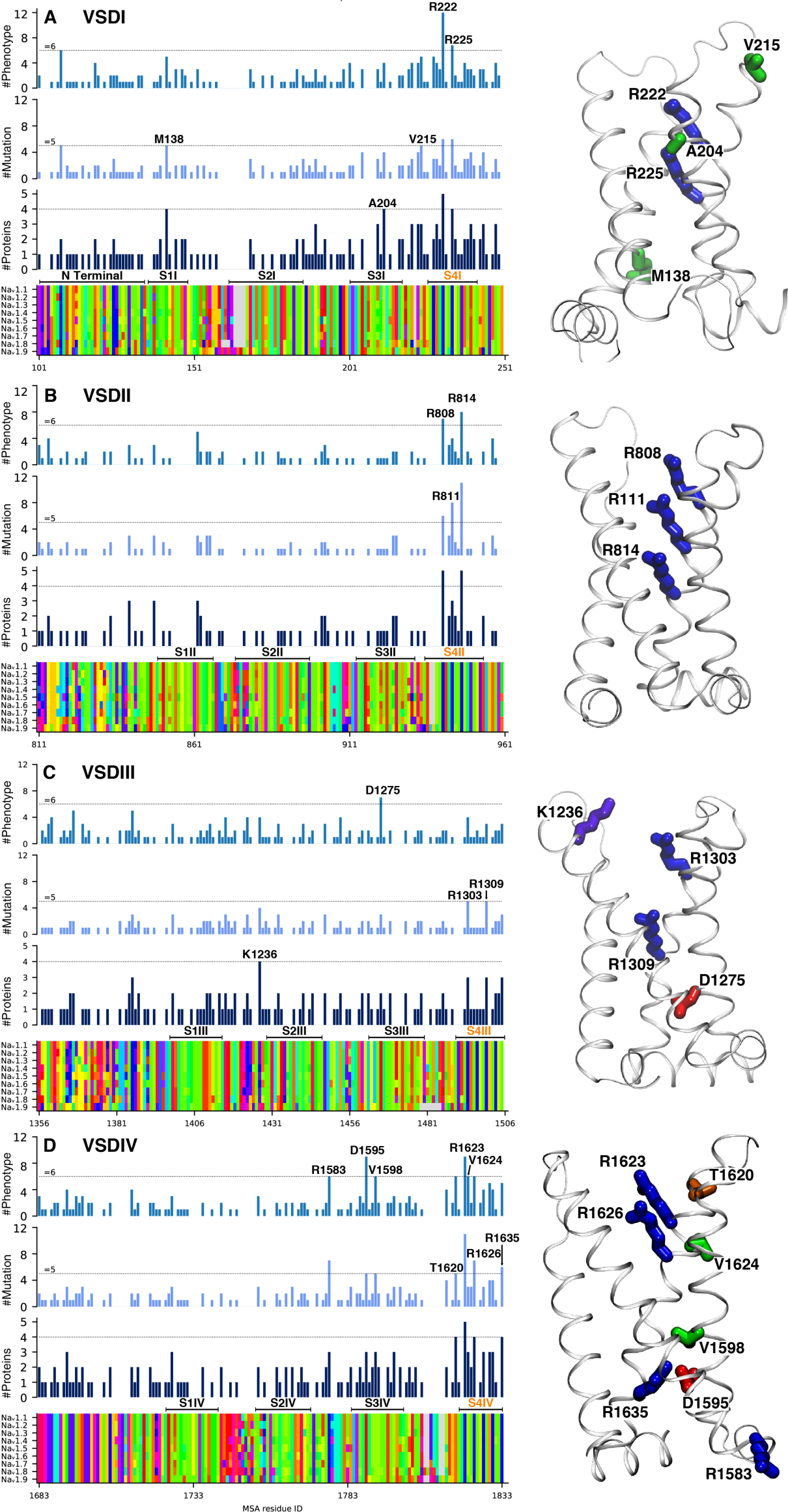
The mutation hotspots in VSDs. (Left) The hotspots showing in the MSA of Nav channels for 4 VSDs. All mutation hotspots in VSDs are labeled with the residue IDs in Na_v_1.5. (Right) Mapping the hotspots in the structure of VSDs. Residues are shown in licorice and colored according to Taylor color scheme.

Similar to all variants in VSDs, mutations of gating charges yield diverse effects on a variety of gating properties. Statistically, these gating-charge mutations show only a moderate preference toward LoF_GP_ on I_max_, τ_rec_, V_1/2 Act_, V_1/2 Inact_, and I_P_. Notably, gating-charge mutations are the primary variants that lead to omega-pore currents (I_ω_), which signifies a dominant GoF_GP_ effect at the molecular level. These mutations provide quintessential examples that the impacts of mutations are strongly influenced by the structural context. For instance, mutations of R2 in VSD_I_ and VSD_II_ can lead to diverse impacts due to the unique interactions within each VSD. Specifically, R2 mutations in VSD_I_ are likely to result in a hyperpolarizing shift in V_1/2 Act._ and V_1/2 InAct. 48,59_, or increasing the maximum sodium current^48^. Conversely, the same mutations in VSD_II_ can cause a depolarizing shift in V_1/2 Act._, a hyperpolarizing shift in V_1/2 InAct._, and a reduction in both maximum current amplitude and persistent current ^60–62^. It is important to note that studies on R2 mutations in VSD_I_ have reported mixed effects from these mutations^63–66^, indicating varying outcomes. However, mutations in VSD_II_ consistently exhibit a clear LoF_o_ effect. As such, predicting the effects of a gating-charge mutation remains a challenging task, given that the effects are highly sensitive to nearby residues in different structural states.

The third feature lies in the non-uniform distribution of mutation hotspots across the four VSDs. Specifically, VSD_IV_ hosts more mutation hotspots compared to the other three VSDs. These hotspots encompass the gating charges (R1623, R1626, R1635), countercharge (D1595), a positively charged residue in the S2-3 loop (R1583), as well as neutral residues (V1598, T1620, V1624) (Fig. 6). Prior studies have illustrated that the initial three VSDs collectively activate the channel, while VSD_IV_ serves as the critical determinant for both the onset of fast inactivation and recovery ^67–69^. The unique functional roles of the VSDs could potentially explain the disparity in the distribution of hotspots. VSD_IV_, in particular, plays a pivotal and distinctive role, mirroring its higher concentration of mutation hotspots. This functional variation amongst the VSDs is further reflected in their impacts on gating properties. For VSD_I_ and VSD_II_, there are more mutations influencing V_1/2 Act_ than V_1/2 Inact_. Conversely, more mutations in VSD_IV_ affect V_1/2 Inact_ than V_1/2 Act_. In addition, more mutations in VSD_IV_ impact the recovery rate (τ_rec_) than do those in VSD_I_ to VSD_III_ (Fig.4).

### Most mutations reduce ion conductivity in the upper pore domain

Being the pivotal structure of Na_v_ channels, the PD governs the permeation of Na^+^ ions via three key functional transitions: activation, inactivation, and recovery ^17^. Thus, it is reasonable that our disease-associated mutation map identifies an abundance of mutation hotspots in the PD. These hotspots are predominantly concentrated in two areas: the upper (PD (upper)) and lower (PD (lower)) parts of the pore domain.

Within the PD (upper), mutations are widespread in regions such as S5, S6, pore helices, pore loops, and extracellular loops (Fig.7). However, the majority of the mutation hotspots are found in structural segments proximate to the selectivity filter (SF), including pore helices (L1 and L2) and the pore loop (Fig.7). Apart from a few non-charged residues (P1438 (L2_III_), G1712 (P1_IV_), and C1728(L2_IV_)), most hotspots in this region are charged residues such as R367 (P1_I_), R383 (L2_I_), R878 (L1_II_), R893 (P1_II_), E901 (P1_II_), and D1741(L2_IV_). Pathogenic effects invariably ensue from mutations of these charged residues. For example, mutations such as R878C, R893C, and E901K in Na_v_1.5 all induce BRGDA1^50,70^, while R367C (P1_I_) exhibits mixed effects, being associated with both LQT3 and BRGDA1 ^50,51,71,72^.

**Figure 7:**
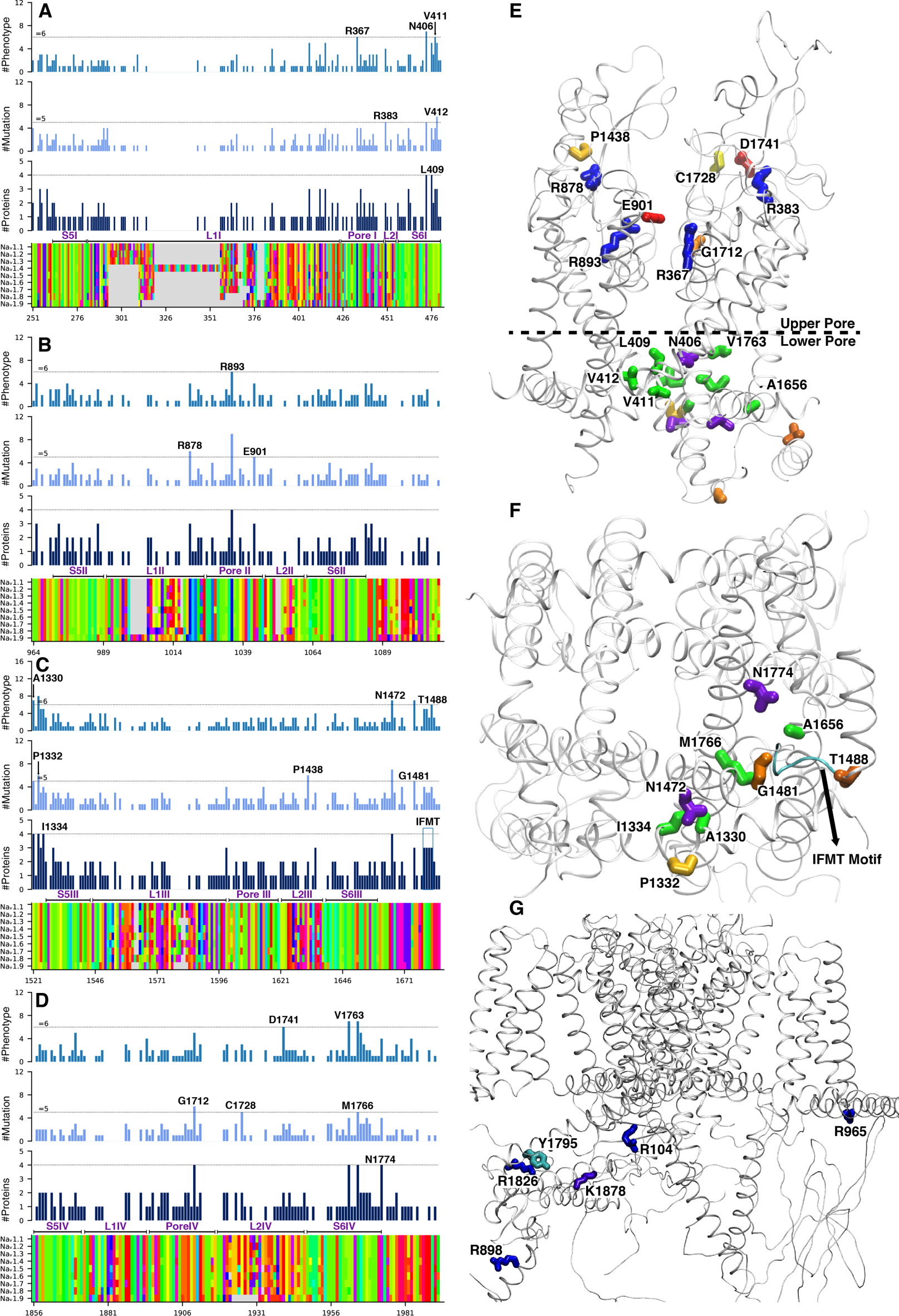
The mutation hotspots in the pore domain. (Left) The hotspots in PD showing in the MSA of Na_v_ channels. (Right) Mapping of the hotspots in the PD of the Na_v_1.5 structure from the sideview (top) and bottom view (middle). The mutation hotspots at the intracellular loops are shown in the right bottom image.

Contrary to VSD mutations, mutations in the PD (upper) region demonstrate greater consistency in their impacts on gating properties. 94.1% of mutations located near the SF affect I_max_ and most of them result in reduced I_max_ of Na_v_ channels, and only approximately 10% of mutations affect other gating properties such as τ_rec_, I_P_, V_1/2 Act_, and V_1/2 Inact_ (Fig.4). Our hypothesis is that these mutations disrupt critical interactions adjacent to the SF, such as salt bridges, potentially causing conformational changes to the SF and altering ion permeability. For instance, R893 directly forms salt bridges with two glutamate residues, E898 (SF_II_) and E901 (P2_II_). Notably, both R893C and E901K have been associated with BRGDA1^50^. Similarly, R878 forms a salt bridge with D1430 (P2_III_) from the neighboring repeat. It’s noteworthy that the probability of mutations impacting I_max_ in the entire PD (upper) decreases to 74.5%, whereas the mutations affecting V_1/2 Act_ or V_1/2 Inact_ increase to one-third (Fig.4). This suggests that these non-SF mutations affect the structural transitions rather than disrupting the SF conformation. Many of these residues are proximate to the VSD/PD interface, suggesting they likely influence VSD transitions through non-canonical coupling that observed in other voltage-gated ion channels ^73^.

### GoF_GP_ impacts of mutations in the lower pore domain

The PD (lower) contains the activation gate (AG) and encompasses the receptor site of the fast-inactivation IFMT motif situated at the edge of the AG (Fig.1) ^74^. When mutations occur near the AG or in the fast-inactivation region (FIR), they are primarily associated with GoF_o_ phenotypes such as LQT3 in Na_v_1.5 ^51,75^. The AG, mechanically coupled with the VSDs, responds to the membrane-potential depolarization by opening during activation and subsequently closing during repolarization to revert to the resting state ^17^. The hotspot mutations (N406 (S6I), V411 (S6I), V412 (S6I), V1763 (S6IV), M1766 (S6IV), and N1774 (S6IV)) are all situated near the intracellular end (Fig.7), a region critical to the opening and closing of the gate. The mutations of most AG residues are consistently linked with GoF_o_ phenotypes (Table S2), aligning with our analysis of their effects on gating properties. The majority of these AG mutation hotspots induce a higher persistent current (I_P_), hyperpolarize the V_1/2 Act_, or depolarize the V_1/2 Inact_ (Fig. 4). Accordingly, we propose that these GoF_o_ mutations may either facilitate the opening of the gate, stabilize the open state, or hinder its complete closure of the gate. Notably, it is observed that some AG mutations can also decrease I_max_, potentially explaining why a few residues in this region are associated with LoF_o_ phenotypes.

The fast-inactivation segment, located in the intracellular loop connecting the third and fourth domains (III-IV linker) (Fig.7) ^76^, is crucial for fast inactivation by binding to its own site at the edge of the AG (Fig.1) ^74^. The IFMT motif, a triple-hydrophobic motif combined with a polar residue in the III-IV linker (Fig.7), is the essential motif for binding ^77^. According to our analysis of mutation hotspots, mutations associated with LQT3 are primarily located in the IFMT motif and its receptor site, with all four amino acids in the IFMT identified as mutation hotspots (Fig.7). While mutations linked to BRGDA1 or overlap syndromes (both LQT3 and BRGDA1) are also seen among the hotspot mutations, they occur less frequently. For instance, F1486L, M1487L, and T1488R mutations are all associated with LQT3, while I1485V is linked to BRGDA1 ^51^. The III-IV linker, an elongated loop containing approximately 60 amino acids, also contains mutation hotspots like N1472S and G1481E (Fig.7). Near these residues, additional mutation hotspots can be found in other structural segments, such as the S4-S5III linker, the intracellular ends of S5III and S6III (Fig.7), S4-S5IV linker, and the C-terminal domain (CTD). For example, A1330 from the S4-S5III linker and N1472 from the III-IV linker (Fig.7), both mutation hotspots, are in contact and associated with LQT3 ^51^. Nearby A1330, residues P1332 and I1334 are also mutation hotspots. I1334V is associated with LQT3 ^51^, while P1332L is linked to BRGDA1 ^23^. These residues contribute to fast inactivation by binding the III-IV linker to the PD, thereby allosterically blocking the activation gate. According to our gating property map, 100% of mutation hotspots in the fast inactivation region (FIR) depolarize V_1/2 Inact_, thus leading to a GoF_GP_ effect on channel activity. Fewer FIR residues impact the maximal current (I_max_) compared to AG mutations. Our analysis of variant effects on gating properties confirms, rather than directly blocking the activation gate, the fast-inactivation segment blocks the gate allosterically. This is consistent with the allosteric blocking mechanism for fast inactivation in Na_v_ channels ^78,79^.

### Gating-property impacts of mutations in other structural segments

Although structural segments such as N/C terminals, VSD-PD linkers, and repeat linkers are significantly less conserved than VSDs and PDs, they host a substantial number of disease-associated mutations and exhibit distinct tendencies in their impacts on gating properties. Approximately 60% of mutations located in the VSD-PD linkers and repeat linkers favor a depolarizing shift in V_1/2 Inact_, a characteristic LoF_GP_ effect. Over one-third of mutations in these linkers also show a strong propensity to increase I_P_, another LoF_GP_ effect (Fig.4). These effects mirror those observed in the lower PD, which could be attributed to the fact that these linkers also contribute to fast inactivation. Notably, over 70% of mutations in the N-terminus tend to reduce I_max_ (Fig.4), likely due to the influential role of residues in this region on Na_v_ channel expression and folding. On the other hand, mutations in the C-terminus have the highest likelihood to increase I_P_ (Fig.4). Although the structural role of the C-terminus remains largely undeciphered, the observed impacts on gating properties underscore its crucial role in maintaining the channel impermeable in its inactivated and resting states.

### The relationship between phenotypes and structural segments

Mapping mutations corresponding to each phenotype onto the structural segments of the sodium channels shows no evident pattern for the overall effect of each phenotype based on their distributions across structural segments (Fig.8). There is even no significant preference for GoF_o_ (or LoF_o_) phenotypes for their structural distributions. Only in the case of CMD1E (Na_v_1.5) are all mutations located in one structural segment (VSDs). Additionally, the mutations associated with ASD, NKPP, and HOKPP2 are located in only two structural segments. For most other phenotypes, mutations are found in at least three distinct structural segments. This observation underlines the fact that mutations across different regions of the sodium channels are capable of similarly altering the channel function, leading to comparable phenotypic effects. It highlights the complexity of the relationships between genetic mutations, the structural domains of ion channels, and the phenotypic outcomes of these mutations, suggesting that the overall effect on channel function is determined by a combination of alterations in multiple channel properties and regions.

**Figure 8:**
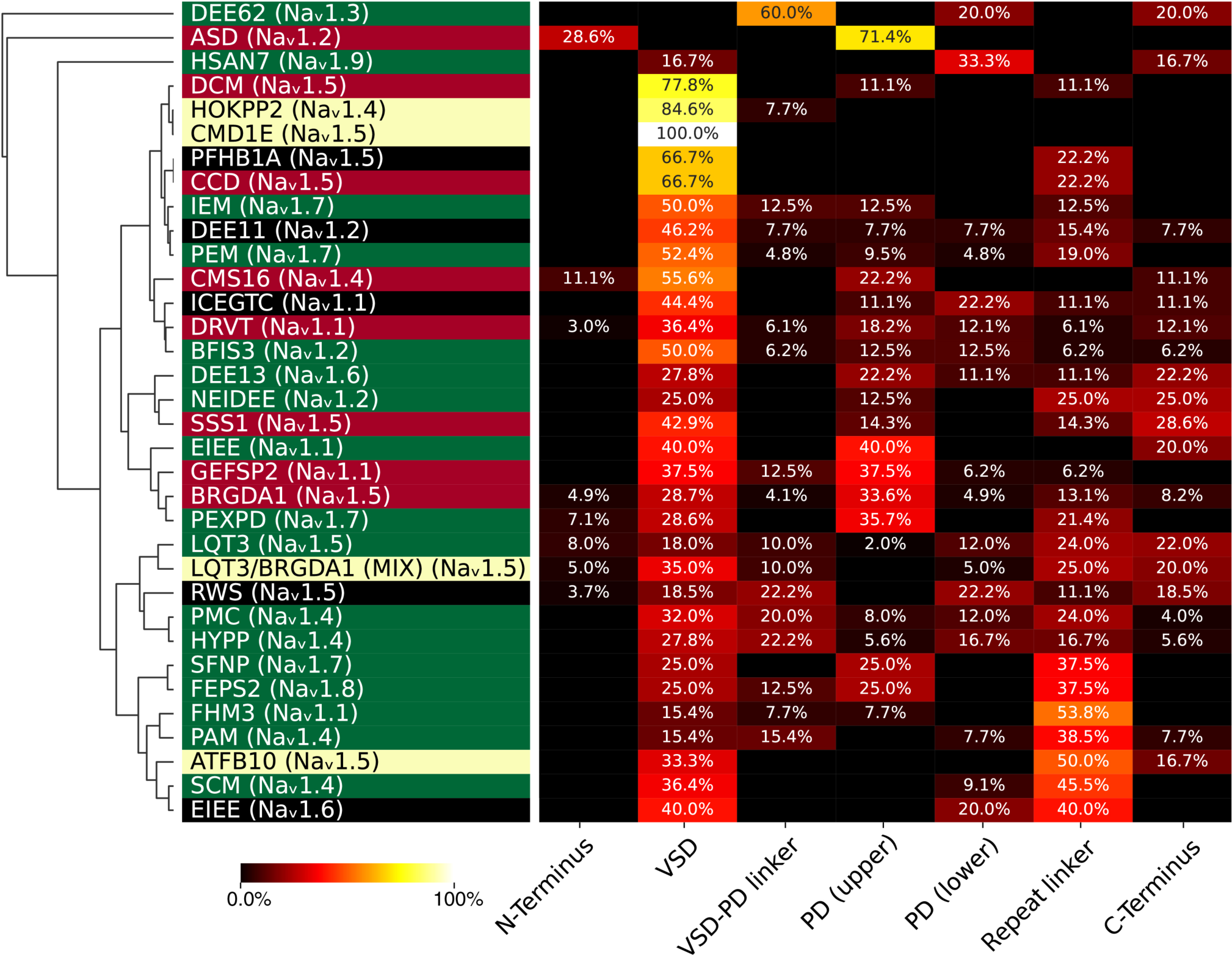
Mapping the phenotypes based on the gating-property impacts of associated mutations. Phenotype clustering of missense mutation in Na_v_ based on the similarity of gating-property impacts. Black cells represent no data for the corresponding disease segment. The heatmap is colored based on the percentage (%) of mutations affecting a certain gating property within a specific phenotype. Diseases are colored in green (GoF_o_), red (LoF_o_), yellow (MiX_o_), and black for undetermined phenotypes.

On the other hand, when a phenotype displays a dominant distribution in specific structural segments, it often aligns with the structural preference for gating properties. For instance, mutations situated in VSD-PD linkers and D linkers preferentially demonstrate a GoF_o_ effect. Correspondingly, conditions such as DEE62 (Na_v_1.3), where 60% of mutations are located in the VSD-PD linker, and several other phenotypes (including FHM3(Na_v_1.1) and SCM(Na_v_1.4)) with close to 50% mutations in Repeat linkers, are mainly categorized as GoF_o_ overall effect phenotypes (Fig.8). In contrast, a large portion of mutations in the upper pore domain (PD (upper)) reduce the maximum current (I_max_), causing a loss of function (LoF_GP_) effect. Several phenotypes with over 30% mutations in PD (upper), such as ASD (Na_v_1.2), BRGDA1 (Na_v_1.5), and GEFSP2 (Na_v_1.1), tend to be classified as LoF_o_ phenotypes. Nonetheless, there are exceptions showing a GoF_o_ effect, including PEXPD (Na_v_1.7) and EIEE (Na_v_1.1). There is only one phenotype, HSAN7 (Na_v_1.9), with over 30% mutations in the lower pore domain (PD (lower)). HSAN7 (Na_v_1.9), along with several other diseases with over 20% mutations in PD (lower) (such as DEE62(Na_v_1.3) and EIEE (Na_v_1.6)), tend to show GoF_o_ overall effect phenotypes. Many phenotypes exhibit 30% mutations in VSDs, but there is no clear preference for GoF_o_ or LoF_o_ effect. This is congruent with the observation that mutations in the VSD have diverse impacts on all gating properties and display modest preferences on a range of gating properties.

### Pathogenic mutations are rich in Arginine

In terms of amino-acid mutation frequencies, a significant feature is that the pathogenic mutations of arginines (Arg) are enriched in in voltage-gated sodium channels. The probability of a disease mutation at different amino acids was calculated and shown in Fig. 9A. For comparison, the probability of benign or uncertain mutations and expected frequencies were also calculated (Fig.9B). Accordingly, a mutation at an Arg residue has the highest probability of causing a disease. As a positively charged amino acid, Arg is often involved in many essential biochemical processes. Arg mutation is well-known for its high pathogenicity mainly due to the fact that Arg mutates to residues with very different chemical properties, such as glutamine (Gln), glycine (Gly), cysteine (Cys), histidine (His), and tryptophan (Trp)^80^. Different from Arg, another positively charged residue, lysine (Lys) shows the lowest relative pathogenicity in its mutations. This difference indicates that the highest probability of disease mutations at Arg may be related to important and unique structural roles in voltage-gated ion channels.

Almost all Arg mutation hotspots are located in three critical regions. These Arg residues either work as gating charges in VSDs, locate in the upper part of PD behind the SF, or distributed in intracellular loops. As the essential voltage sensors in VSDs, it is reasonable that the mutations of gating charges change activation or inactivation voltage dependence and then induce dysfunctions of ion channels. ∼70% of the gating-charge mutations exhibit strong preference to hyperpolarize V_1/2 Inact_. The Args behind the SF, such as R367, R383, R878, R893, form salt bridges with negatively charged residues (Fig.7A). Breaking critical salt bridges, these Arg mutations can induce the conformational changes of the SF and then reduce the ion permeability. Meanwhile, pathogenic Arg mutations are also popular in different intracellular loops, such as R104 in the N-terminal, R965 in repeat (II-III) linker, R1583 in S2-3 linker of VSD4, R1644 in VSD-PD linker of repeat 4, and R1826 as well as R1898 in the C-terminal. Due to their locations, these Arg may play key roles in fast inactivation, persistent current, membrane protein orientation^81^, protein-lipid interactions^82^, and protein-protein interactions^83^. All of these pathogenic Arg mutations indicate that Na_v_ channels evolved diverse voltage-dependent mechanisms, not only for activation/inactivation, but also for regulation by lipids or other proteins.

## Discussion

The ability to predict variant effects on gating properties based on their structural location is highly dependent on the precise location of the mutation. Mutations in the PD (upper), particularly those near the SF, are highly likely to significantly reduce the I_max_ (Fig.4). Conversely, within the PD (lower), mutations in the FIR are likely to shift V_1/2 Inact_ in the depolarizing direction, while mutations in the AG often enhance I_p_ (Fig.4). For mutations within the VSDs, predicting their impacts on gating properties proves to be a challenging task. Our analysis illustrates that mutations in VSDs can have a broad range of effects on all gating properties with modest preferences towards V_1/2 Act_, V_1/2 Inact_, τ_rec_, and I_max_. As the voltage-sensing components of Na_v_ channels, VSDs initiate all three major functional transitions: activation, inactivation, and recovery ^17,68^. Mutations within VSDs can either enhance or reduce channel activity by altering the structural transition between the resting (“down”) and activated/inactivated (“up”) states. When a mutation introduces or disrupts specific interactions, it could potentially influence the conformational equilibrium between the “up” and “down” states, as well as the rates of their transitions. In some instances, mutations can result in leaky channels directly via the VSD, leading to the generation of omega pore currents (Iω). These combined factors account for the extensive range of functional effects associated with VSD mutations, providing a structural foundation for their diverse impacts. The impacts of VSD mutations are highly sensitive to the structural context of the mutated residue. Notably, this structural context is dependent on the state, selective to the subtype, and varies across VSDs of the same channel. Therefore, theoretically, if the interactions between a specific residue and its neighboring amino acids (or other molecules) across multiple functional states (including intermediates between the “down” and “up” states) are known, it could be possible to predict its impacts on gating properties based on structures in different functional states.

Our analysis shows that the mapping of phenotypes to their mutant-located structural segments does not reveal a significant correlation between structural distribution and phenotypes. The weak relationship between phenotype and structural distribution could be attributed to several factors. First, residues in different structural segments may have similar impacts on gating properties. Consequently, mutants associated with the same phenotype are not necessarily concentrated in a specific structural segment. Second, the number of identified mutations for many phenotypes is still insufficient, and the accuracy of phenotype association requires improvement. With the discovery of more disease-associated mutations, a clearer pattern may emerge linking the structural distribution of mutations, their effects on gating properties, and associated phenotypes. Thus, the ongoing discovery and characterization of disease-associated mutations will likely continue to refine our understanding of the relationship between mutations, structural segments, gating properties, and diseases.

The study’s mapping of gating-property impacts on 34 sodium channelopathies reveals that, even within the same overall effect (GoF_o_ or LoF_o_) classification, different phenotypes demonstrate unique patterns of gating property alterations. This suggests that the same overall effect (either GoF_o_ or LoF_o_) can result from different molecular mechanisms. The similarities in overall effects across certain phenotypes may be coincidental and arise from dominant impacts of different gating properties or accumulative effects of distinctive alterations. This finding highlights the challenges of the binary prediction model of gain or loss of function, both for interpreting pathophysiology and designing personalized treatments. Conversely, phenotypes belonging to the same sub-branch (as shown in Fig.4), such as IEM (Na_v_1.7) vs DEE13 (Na_v_1.6), and PEXPD (Na_v_1.7) vs PMC (Na_v_1.4), show high similarities in gating properties. This implies that categorizing diseases based on altered gating properties may provide more accurate clustering of diseases driven by similar pathophysiological mechanisms. This nuanced understanding of gating property impacts has the potential to improve prediction accuracy and the design of more effective treatments. There may also be opportunities for repurposing drugs that treat a particular phenotype for the treatment of other diseases with a similar effect on gating properties.

## Acknowledgments

Discussions with Claudio Grosman, Joao Luis Carvalho De Souza, and Tamer M. Gamal El-Din are gratefully acknowledged. Computer resources came from a Maximize ACCESS allocation through project BIO210015, an allocation (MCB200085P) on Antons at the Pittsburgh Supercomputing Center provided by the National Center for Multiscale Modeling of Biological Systems through National Institutes of Health grant P41GM103712-1 and from a loan from D. E. Shaw Research, and a Frontera Pathways allocation (MCB21012) at the Texas Advanced Computing Center (TACC).

## Author contributions

Conceptualization: AA, JL

Methodology: AA, JL

Investigation: AA, EE, HB

Visualization: AA

Supervision: JL

Writing—original draft: JL

Writing—review & editing: AA, EE, HB, JL

## Data and materials availability

All data used in this study are available in the manuscript and the Supplementary material.

## Author contributions

All authors contributed to conception and interpretation of data.

A.A., E.E., and H.B. contributed to the meta-analysis of 366 electrophysiology research article.

A.A. carried out bioinformatics analysis of 2,400 disease-associated variants.

A.A. carried out data visualization.

J.L. carried out design and writing the manuscript.

