## supplemental tables and figures for "Mapping Structural Distribution and Gating-Property Impacts of Disease-Associated Missense Mutations in Voltage-Gated Sodium Channels"

This file includes:

Tables S1 to S5

Figure S1

References cited in Tables S1 to S5

Table S1: Diseases with clear GoF/LoF assignment in Voltage gated sodium channels that have more than 5 mutations

| # | Protein | Phenotype | OMIM | GOF/LOF | Reference |
|---|---|---|---|---|---|
| 1 | Nav1.1 | Familial Hemiplegic Migraine Type 3 | FHM3 | GOF | (Mantegazza et al., 2021) |
| 2 | Nav1.1 | Early Infantile Epileptic Encephalopathy | EIEE | GOF | (Mantegazza et al., 2021) |
| 3 | Nav1.1 | Generalized Epilepsy With Febrile Seizures Plus Type 2 | GEFS+2 | LOF | (Mantegazza et al., 2021) (Ragsdale, 2008) |
| 4 | Nav1.1 | Dravet Syndrome | DRVT | LOF | (Mantegazza et al., 2021) (Ragsdale, 2008) |
| 5 | Nav1.2 | Neonatal-early Infantile Developmental And Epileptic Encephalopathies | NEIDEE | GOF | (Mantegazza et al., 2021) (Sanders et al., 2018) |
| 6 | Nav1.2 | Benign Familial Infantile, 3 | BFIS3 | GOF | (Mantegazza et al., 2021) |
| 7 | Nav1.2 | Autism Spectrum Disorder | ASD | LOF | (Sanders et al., 2018) (Mantegazza et al., 2021) |
| 8 | Nav1.3 | Developmental And Epileptic Encephalopathy 62 | DEE62 | GOF | (Sanders et al., 2018) |
| 9 | Nav1.4 | Hyperkalemic Periodic Paralysis | HYPP | GOF | (Sanders et al., 2018) (Nicole & Fontaine, 2015) |
| 10 | Nav1.4 | Paramyotonia Congenita | PMC | GOF | (Nicole & Fontaine, 2015) (Mantegazza et al., 2021) |
| 11 | Nav1.4 | Sodium Channel Myotonias | SCM | LOF | (Nicole & Fontaine, 2015) (Nicole & Lory, 2021) |
| 12 | Nav1.4 | Congenital Myasthenic Syndrome-16 | CMS16 | LOF | (Nicole & Fontaine, 2015) (Nicole & Lory, 2021) (Mantegazza et al., 2021) |
| 13 | Nav1.4 | Hypokalemic Periodic Paralysis, Type 2 | HOKPP2 | MIX | (Mantegazza et al., 2021) (Nicole & Fontaine, 2015) (Nicole & Lory, 2021) |
| 14 | Nav1.4 | Potassium-aggravated Myotonia | PAM | GOF | (Mantegazza et al., 2021) (Nicole & Fontaine, 2015) (Nicole & Lory, 2021) |
| 15 | Nav1.4 | Normokalemic Periodic Paralysis | NKPP | GOF | (Mantegazza et al., 2021) |
| 16 | Nav1.5 | LQT syndrome type 3 | LQT3 | GOF | (W. Li et al., 2018) (Wilde & Amin, 2018) (M. Liu et al., 2016) (Remme et al., 2008) |
| 17 | Nav1.5 | Dilated Cardiomyopathy | DCM | LOF | (M. Liu et al., 2016) |
| 18 | Nav1.5 | Brugada Syndrome 1 | BRGDA1 | LOF | (Wilde & Amin, 2018) (M. Liu et al., 2016) (Remme et al., 2008) |
| 19 | Nav1.5 | Cardiac Conduction Disease | CCD | LOF | (W. Li et al., 2018) (Wilde & Amin, 2018) (M. Liu et al., 2016) (Remme et al., 2008) |

| # | Protein | Phenotype | OMIM | GOF/LOF | Reference |
|---|---|---|---|---|---|
| 20 | Nav1.5 | Sick Sinus Syndrome 1 | SSS1 | LOF | (W. Li et al., 2018)<br>(Wilde & Amin, 2018)<br>(M. Liu et al., 2016)<br>(Remme et al., 2008) |
| 21 | Nav1.5 | Atrial Fibrillation, Familial, 10 | ATFB10 | MIX | (W. Huang et al., 2017) |
| 22 | Nav1.5 | Cardiomyopathy, Dilated, 1E | CMD1E | MIX | (W. Huang et al., 2017) |
| 23 | Nav1.6 | Developmental And Epileptic Encephalopathy 13 | DEE13 | GOF | (Mantegazza et al., 2021)<br>(Solé et al., 2020) |
| 24 | Nav1.7 | Primary Erythromelalgia Mutations | PEM | GOF | (Drenth & Waxman, 2007)<br>(Mantegazza et al., 2021) |
| 25 | Nav1.7 | Small Nerve Fiber Neuropathy | SFNP | GOF | (Lee et al., 2020)<br>(Tanaka et al., 2017) |
| 26 | Nav1.7 | Paroxysmal Extreme Pain Disorder | PEXPD | GOF | (Mantegazza et al., 2021)<br>(Drenth & Waxman, 2007) |
| 27 | Nav1.7 | Inherited Erythromelalgia | IEM | GOF | (Estacion et al., 2008) |
| 28 | Nav1.8 | Episodic Pain Syndrome, Familial, 2 | FEPS2 | GOF | (Shen et al., 2022)<br>(Faber, Lauria, Merkies, Cheng, Han, Ahn, Persson, Hoeijmakers, Gerrits, Pierro, Dib-Hajj, et al., 2012) |
| 29 | Nav1.9 | Neuropathy, Hereditary Sensory And Autonomic, Type VII | HSAN7 | GOF | (X. Y. Zhang et al., 2013)<br>(C. Han et al., 2017)<br>(Leipold et al., 2013) |

Table S2: Hotspot Overlay in Voltage Gated Sodium Channels:

| MSA | Na$_v$1.1 | | Na$_v$1.2 | | Na$_v$1.3 | | Na$_v$1.4 | | Na$_v$1.5 | | Na$_v$1.6 | | Na$_v$1.7 | | Na$_v$1.8 | | Na$_v$1.9 | |
|---|---|---|---|---|---|---|---|---|---|---|---|---|---|---|---|---|---|---|
| | MUT* | DIS** | MUT | DIS | MUT | DIS | MUT | DIS | MUT | DIS | MUT | DIS | MUT | DIS | MUT | DIS | MUT | DIS |
| | R101W† | EIEE | | | | | R104H | CMS16 | R104G | LQT3 | | | R99H | CIP | | | | |
| | | DRVT | | | | | | | | RWS | | | | | | | | |
| | R101Q | DRVT | | | | | | | R104Q | BRGDA1 | | | | | | | | |
| 108 | | ICEGTC | R102† | | R101 | | | | R104W | BRGDA1 | 105R | | | | R103 | | R102 | |
| | M135T | DRVT | M136I | DEE11 | | | | | M138T | DCM | M139I | EIEE | | | | | | |
| 142 | | | | | M135 | | M138 | | M138I | ATFB10 | | | M133 | | M137 | | M136 | |
| 212 | A201E | EIEE | A202 | | A201 | | A204V | HYPP | A204V | BRGDA1 | A205 | | A199 | | A200V | BRGDA1 | A206 | |
| | V212A | DRVT | V213A | BFIS3 | | | | | | | V216D | DEE13 | | | | | | |
| 224 | | | V213D | DEE11 | V212 | | I215 | | V215 | | | | V210 | | I211 | | L218 | |
| | | | | BFIS3 | | | R222W | HOKPP2 | | CMD1E | R223G | DEE13 | | | | | | FEPS3 |
| | | | R220G | DEE11 | | | | | | RWS | | | | | | | R225C | HSAN7 |
| | | | | | | | | | | DCM | | | | | | | | |
| | | | | | | | | | | LQT3 | | | | | | | | |
| 231 | R219 | | | | R219 | | | | R222Q | BRGDA1 | | | R217 | | R218 | | | |
| 234 | R222 | | R223Q | BFIS3 | R222 | | R225Q | HYPP | R225Q | LQT3 | R226G | CIAT | R220 | | R221 | | R228 | |

| MSA | $Na_v1.1$ | | $Na_v1.2$ | | $Na_v1.3$ | | $Na_v1.4$ | | $Na_v1.5$ | | $Na_v1.6$ | | $Na_v1.7$ | | $Na_v1.8$ | | $Na_v1.9$ | |
|---|---|---|---|---|---|---|---|---|---|---|---|---|---|---|---|---|---|---|
| | MUT* | DIS** | MUT | DIS | MUT | DIS | MUT | DIS | MUT | DIS | MUT | DIS | MUT | DIS | MUT | DIS | MUT | DIS |
| | | | | | | | R225W | MYOSCN4A | R225W | PFHB1A | | | | | | | | |
| | | | | | | | | SCM | | LQT3 | | | | | | | | |
| | | | | | | | | | | BRGDA1 | | | | | | | | |
| | | | | | | | | | | CCD | | | | | | | | |
| | R377Q | DRVT | R379H | ASD | | | | | R367L | BRGDA1 | | | | | | | | |
| | | GEFSP2 | | | | | | | R367H | BRGDA1 | | | | | | | | |
| | | GEFSP1 | | | | | | | R367C | LQT3 | | | | | | | | |
| | | | | | | | | | | BRGDA1 | | | | | | | | |
| 434 | | | | | R378 | | R401 | | | RWS | R365 | | R356 | | R351 | | R354 | |
| | R393C | EIEE | | | | | | | | | | | R372C | DRVT | | | | |
| | | DRVT | | | | | | | | | | | | | | | | |
| | R393H | EIEE | | | | | | | | | | | | | | | | |
| | | ICEGTC | | | | | | | | | | | | | | | | |
| | | DRVT | | | | | | | | | | | | | | | | |
| | R393P | EIEE | | | | | | | | | | | | | | | | |
| 450 | R393S | DRVT | R395 | | R394 | | R417 | | R383 | | R381 | | | | R367 | | R370 | |
| | | | N418S | BFIS3 | | | N440K | HYPP | N406K | LQT1 | | | N395K | PERYTHM | | | | |
| 473 | N416 | | | | N417 | | | PMC | | LQT3 | N404 | | | | N390 | | N393 | |

| MSA | $Na_v1.1$ | | $Na_v1.2$ | | $Na_v1.3$ | | $Na_v1.4$ | | $Na_v1.5$ | | $Na_v1.6$ | | $Na_v1.7$ | | $Na_v1.8$ | | $Na_v1.9$ | |
|---|---|---|---|---|---|---|---|---|---|---|---|---|---|---|---|---|---|---|
| | MUT* | DIS** | MUT | DIS | MUT | DIS | MUT | DIS | MUT | DIS | MUT | DIS | MUT | DIS | MUT | DIS | MUT | DIS |
| | | | | | | | | | | RWS | | | | | | | | |
| | | | | | | | | | N406S | BRGDA1 | | | | | | | | |
| | | | | | | | L443P | HYPP | | LQT3 | L407F | DEE13 | | | | | L396P | HSAN7 |
| 476 | L419 | | L421 | | L420 | | | | L409V | RWS | | | L398 | | L393 | | | |
| | V421M | DRVT | V423L | NEIDEE | | | | MYOSCN4A | | LQTS | | | | | | | | |
| | | | | | | | | HYPP | | LQT3 | | | | | | | | |
| | | | | | | | V445M | PAM | V411M | RWS | | | | | | | | |
| 478 | | | | | V422 | | V445L | HYPP | | | V409 | | V400 | | V395 | | V398 | |
| | V422M | DRVT | V424L | DD | | | | | | | V410L | DEE13 | | | | | | |
| | V422A | DRVT | V424A | BFIS3 | | | | | | | | | | | | | | |
| 479 | V422L | DEE6B | | | V423 | | V446 | | V412 | | | | V401 | | V396 | | V399 | |
| | | | E430A | BFIS3 | | | | HYPP | | | | | | | | | | |
| | | | | BFIS3 | | | E452D | HOKPP2 | | | | | | | | | | |
| | | | E430G | DEE11 | | | | MYOSCN4A | | | | | | | | | | |
| 485 | E428 | | E430Q | BFIS3 | E429 | | E452K | HYPP | E418 | | E416 | | E407 | | E402 | | E405 | |
| | | DRVT | | | | | R669H | HOKPP2 | | BRGDA1 | R844Q | CIAT | | | R756W | FEPS2 | | |
| 941 | R859C | GEFSP2 | R850 | | R851 | | R669W | NKPP | R808P | RWS | | | R835 | | | | R670 | |

| MSA | $Na_v1.1$ | | $Na_v1.2$ | | $Na_v1.3$ | | $Na_v1.4$ | | $Na_v1.5$ | | $Na_v1.6$ | | $Na_v1.7$ | | $Na_v1.8$ | | $Na_v1.9$ | |
|---|---|---|---|---|---|---|---|---|---|---|---|---|---|---|---|---|---|---|
| | MUT* | DIS** | MUT | DIS | MUT | DIS | MUT | DIS | MUT | DIS | MUT | DIS | MUT | DIS | MUT | DIS | MUT | DIS |
| | R859H | GEFSP2 | | | | | R669Q | NKPP | | | | | | | | | | |
| | | | | | | | R669G | NKPP | | | | | | | | | | |
| | R862G | DRVT | | BFIS3 | | | R672L | HOKPP2 | | | | | | | | | | |
| | R862Q | DRVT | | DEE11 | | | R672H | HOKPP2 | | | | | | | | | | |
| | | | R853Q | ICDEE | | | R672G | HOKPP2 | | | | | | | | | | |
| | | | | | | | R672S | HOKPP2 | | | | | | | | | | |
| 944 | | | | | R854 | | R672C | HOKPP2 | R811 | | R847 | | R838 | | R759 | | R673 | |
| | R865G | DRVT | R856P | BFIS3 | | | R675G | NKPP | | CMD1E | R850Q | DEE13 | | | | | | |
| | | | R856Q | BFIS3 | | | R675Q | NKPP | R814W | DCM | R850E | DEE13 | | | | | | |
| 947 | | | R856L | DEE11 | R857 | | R675W | NKPP | R814Q | BRGDA1 | | | R841 | | R762 | | R676 | |
| | R931P | DRVT | | | | | | | R878C | BRGDA1 | | | R907Q | CIP | | | | |
| | | DRVT | | | | | | | R878H | BRGDA1 | | | | | | | | |
| | R931H | EIEE | | | | | | | | | | | | | | | | |
| 1020 | R931C | DRVT | R922 | | R923 | | R741 | | | | R916 | | | | R829 | | R748 | |
| | | EIEE | | BFIS3 | | | | | R893L | BRGDA1 | | | | | | | R763H | HSAN7 |
| 1035 | R946C | DRVT | R937C | ASD | R938 | | R756 | | R893H | BRGDA1 | R931 | | R922 | | R844 | | | |

| MSA | $Na_v1.1$ | | $Na_v1.2$ | | $Na_v1.3$ | | $Na_v1.4$ | | $Na_v1.5$ | | $Na_v1.6$ | | $Na_v1.7$ | | $Na_v1.8$ | | $Na_v1.9$ | |
|---|---|---|---|---|---|---|---|---|---|---|---|---|---|---|---|---|---|---|
| | MUT* | DIS** | MUT | DIS | MUT | DIS | MUT | DIS | MUT | DIS | MUT | DIS | MUT | DIS | MUT | DIS | MUT | DIS |
| | | GEFSP2 | | ID | | | | | R893C | BRGDA1 | | | | | | | | |
| | R946P | DRVT | R937H | ASD | | | | | | | | | | | | | | |
| | R946S | DRVT | | | | | | | | | | | | | | | | |
| | | DRVT | | | | | | | | | | | | | | | | |
| | R946H | GEFSP2 | | | | | | | | | | | | | | | | |
| | E954K | DRVT | | | | | E764K | HYPP | E901K | BRGDA1 | | | | | | | | |
| | E954V | EIEE | | | | | | | | | | | | | | | | |
| 1043 | E954D | EIEE | E945 | | E946 | | | | | | E939 | | E930 | | E852 | | E771 | |
| | | | | | | | | HYPP | R965L | LQT3 | R1003H | EIEE | | | | | R838Q | HSAN7 |
| | | | | | | | R828H | HOKPP2 | R965L | RWS | | | | | | | R838L | HSAN7 |
| | | | | | | | | | R965C | BRGDA1 | | | | | | | | |
| 1110 | R1018 | | R1009 | | R1010 | | | | R965H | BRGDA1 | | | R994 | | R916 | | | |
| 1427 | K1249N | GEFSP2 | K1239 | | K1237 | | R1062P | HYPP | K1236N | BRGDA1 | R1229H | EIEE | K1223 | | K1183 | | Q1087 | |
| | D1288N | DRVT | | | | | | | | PFHB1A | | | | | | | | |
| | | | | | | | | | | CMD1E | | | | | | | | |
| | | | | | | | | | | ATFB10 | | | | | | | | |
| 1466 | | | D1278 | | D1276 | | D1101 | | D1275N | LQT3 | D1268 | | D1262 | | D1222 | | D1126 | |

| MSA | Na$_v$1.1 | | Na$_v$1.2 | | Na$_v$1.3 | | Na$_v$1.4 | | Na$_v$1.5 | | Na$_v$1.6 | | Na$_v$1.7 | | Na$_v$1.8 | | Na$_v$1.9 | |
|---|---|---|---|---|---|---|---|---|---|---|---|---|---|---|---|---|---|---|
| | MUT* | DIS** | MUT | DIS | MUT | DIS | MUT | DIS | MUT | DIS | MUT | DIS | MUT | DIS | MUT | DIS | MUT | DIS |
| | | | | | | | | | | BRGDA1 | | | | | | | | |
| | | | | | | | | | | ATRST1 | | | | | | | | |
| | | | | | | | | | | CCD | | | | | | | | |
| | | | | | | | | | D1275Y | ATFB10 | | | | | | | | |
| | R1316G | DRVT | | | | | | NKPP | | | | | | | | | R1147W | HSAN7 |
| | R1316S | DRVT | | | | | R1129Q | HOKPP2 | | | | | | | | | | |
| 1494 | | | R1306 | | R1304 | | R1129P | HOKPP2 | R1303 | | R1296 | | R1290 | | R1250 | | | |
| | | | R1312T | DEE11 | | | R1135S | HOKPP2 | | | | | | | R1256Q | FEPS2 | | |
| | | | | | | | R1135H | HOKPP2 | | | | | | | | | | |
| 1500 | R1322 | | | | R1310 | | R1135C | HOKPP2 | R1309 | | R1302 | | R1296 | | | | R1153 | |
| | A1343S | EIEE | | | | | | MYOSCN4A | | LQT3 | A1323S | DEE13 | | | | | | |
| | | | | | | | | HYPP | A1330T | RWS | | | | | | | | |
| | | | | | | | A1156T | PMC | | LQT3 | | | | | | | | |
| 1521 | | | A1333 | | A1331 | | | | A1330P | RWS | | | A1317 | | A1277 | | R1174 | |
| | | EIEE | | | P1333L | DEE62 | P1158S | HOKPP2 | | LQT3 | | | | | | | | |
| | P1345S | DEE6B | | | | | P1158L | HYPP | P1332L | BRGDA1 | | | | | | | | |
| 1523 | P1345L | DRVT | P1335 | | | | | | | | P1325 | | P1319 | | P1279 | | P1176 | |

| MSA | $Na_v1.1$ | | $Na_v1.2$ | | $Na_v1.3$ | | $Na_v1.4$ | | $Na_v1.5$ | | $Na_v1.6$ | | $Na_v1.7$ | | $Na_v1.8$ | | $Na_v1.9$ | |
|---|---|---|---|---|---|---|---|---|---|---|---|---|---|---|---|---|---|---|
| | MUT* | DIS** | MUT | DIS | MUT | DIS | MUT | DIS | MUT | DIS | MUT | DIS | MUT | DIS | MUT | DIS | MUT | DIS |
| | I1347N | DRVT | | | | | I1160V | MYOSCN4A | I1334V | LQT3 | I1327V | DEE13 | | | | | | |
| 1525 | | | I1337 | | I1335 | | | PAM | | RWS | | | I1321 | | I1281 | | I1178 | |
| | P1451L | DRVT | | | | | | | P1438S | BRGDA1 | | | | | | | | |
| | P1451T | DRVT | | | | | | | P1438L | BRGDA1 | | | | | | | | |
| | P1451A | EIEE | | | | | | | | | | | | | | | | |
| 1632 | P1451S | DRVT | P1441 | | P1436 | | P1263 | | | | P1432 | | P1425 | | P1386 | | P1276 | |
| | N1485D | EIEE | | | | | N1297K | MYOSCN4A | N1472S | LQT3 | N1466S | EIEE | | | | | | |
| | N1485Y | DRVT | | | | | N1297K | CAM | N1472S | RWS | N1466K | DEE13 | | | | | | |
| 1666 | | | N1475 | | N1470 | | | | | | N1466T | DEE13 | N1459 | | N1420 | | N1310 | |
| | | | | | | | G1306V | MYOSCN4A | G1481E | LQT3 | G1475R | EIEE | | | | | | |
| | | | | | | | | HYPP | G1481E | RWS | G1475R | DEE13 | | | | | | |
| | | | | | | | | PMC | | | | | | | | | | |
| | | | | | | | | PAM | | | | | | | | | | |
| | | | | | | | | SCM | | | | | | | | | | |
| | | | | | | | G1306E | MYOSCN4A | | | | | | | | | | |
| | | | | | | | | PMC | | | | | | | | | | |
| | | | | | | | | PAM | | | | | | | | | | |
| 1675 | G1494 | | G1484 | | G1479 | | | SCM | | | | | G1468 | | G1429 | | G1319 | |

| MSA | $Na_v1.1$ | | $Na_v1.2$ | | $Na_v1.3$ | | $Na_v1.4$ | | $Na_v1.5$ | | $Na_v1.6$ | | $Na_v1.7$ | | $Na_v1.8$ | | $Na_v1.9$ | |
|---|---|---|---|---|---|---|---|---|---|---|---|---|---|---|---|---|---|---|
| | MUT* | DIS** | MUT | DIS | MUT | DIS | MUT | DIS | MUT | DIS | MUT | DIS | MUT | DIS | MUT | DIS | MUT | DIS |
| | | | | | | | G1306A | PMC | | | | | | | | | | |
| | | | | | | | | PAM | | | | | | | | | | |
| | | | | | | | | SCM | | | | | | | | | | |
| | | | | | | | T1313M | HYPP | T1488R | LQT3 | | | T1475I | PEXPD | | | | |
| | | | | | | | | PMC | | RWS | | | | PEPD | | | | |
| 1682 | T1501 | | T1491 | | T1486 | | | | T1488K | LQT3 | T1482 | | | | T1436 | | T1326 | |
| | R1596H | EIEE | | | | | R1408C | HYPP | R1583C | BRGDA1 | | | | | | | | |
| | | GEFSP2 | | | | | R1408S | HYPP | R1583H | BRGDA1 | | | | | | | | |
| | R1596C | DRVT | | | | | R1408S | HOKPP2 | | | | | | | | | | |
| | | GEFSP2 | | | | | | | | | | | | | | | | |
| 1777 | R1596L | DRVT | R1586 | | R1581 | | | | | | R1577 | | R1570 | | R1531 | | R1421 | |
| | D1608Y | EIEE | | | | | D1420N | HYPP | D1595N | PFHB1A | | | | | | | | |
| | | DRVT | | | | | | HOKPP2 | | ATFB10 | | | | | | | | |
| | D1608G | DRVT | | | | | | | | CCD | | | | | | | | |
| | | | | | | | | | D1595H | CMD1E | | | | | | | | |
| | | | | | | | | | | BRGDA1 | | | | | | | | |
| 1789 | | | D1598 | | D1593 | | | | | DCM | D1589 | | D1582 | | D1543 | | D1433 | |
| 1792 | V1611F | DRVT | V1601M | BFIS3 | V1596 | | V1423 | | V1598 | | V1592I | EIEE | V1585 | | V1546 | | V1436 | |

| MSA | $Na_v1.1$ | | $Na_v1.2$ | | $Na_v1.3$ | | $Na_v1.4$ | | $Na_v1.5$ | | $Na_v1.6$ | | $Na_v1.7$ | | $Na_v1.8$ | | $Na_v1.9$ | |
|---|---|---|---|---|---|---|---|---|---|---|---|---|---|---|---|---|---|---|
| | MUT* | DIS** | MUT | DIS | MUT | DIS | MUT | DIS | MUT | DIS | MUT | DIS | MUT | DIS | MUT | DIS | MUT | DIS |
| | | GEFSP2 | V1601L | BFIS3 | | | | | | | V1592L | DEE13 | | | | | | |
| | | ICEGTC | | | | | | | | | | | | | | | | |
| | T1633 | | T1623N | DEE11 | | | T1445M | HYPP | T1620M | BRGDA1 | T1614A | EIEE | | | | | | |
| | | | | | | | | | | PFHB1A | | | | | | | | |
| | | | | | | | | | | LQT3 | | | | | | | | |
| | | | | | | | | | | BRGDA1 | | | | | | | | |
| 1818 | | | | | T1618 | | | | T1620K | CCD | | | T1607 | | T1570 | | T1460 | |
| | R1636Q | EIEE | R1626Q | BFIS3 | | | R1448P | PMC | | LQT3 | R1617L | EIEE | | | | | | |
| | | | | | | | R1448L | PMC | | BRGDA1 | R1617Q | DEE13 | | | | | | |
| | | | | | | | R1448C | PMC | | LQT3/6, DIGENIC | R1617Q | DEE13 | | | | | | |
| | | | | | | | R1448S | PMC | R1623Q | RWS | | | | | | | | |
| | | | | | | | R1448H | PMC | | LQT3 | | | | | | | | |
| 1821 | | | | | R1621 | | | | R1623L | RWS | | | R1610 | | R1573 | | R1463 | |
| | V1637E | DRVT | V1627M | BFIS3 | | | | | | PFHB1A | | | | | | | | |
| | | | | | | | | | | BRGDA1 | | | | | | | | |
| 1822 | | | | | V1622 | | V1449 | | V1624I | CMD1E | V1618 | | V1611 | | V1574 | | I1464 | |

| MSA | $Na_v1.1$ | | $Na_v1.2$ | | $Na_v1.3$ | | $Na_v1.4$ | | $Na_v1.5$ | | $Na_v1.6$ | | $Na_v1.7$ | | $Na_v1.8$ | | $Na_v1.9$ | |
|---|---|---|---|---|---|---|---|---|---|---|---|---|---|---|---|---|---|---|
| | MUT* | DIS** | MUT | DIS | MUT | DIS | MUT | DIS | MUT | DIS | MUT | DIS | MUT | DIS | MUT | DIS | MUT | DIS |
| | | | | | | | | | | LQT3 | | | | | | | | |
| | R1639G | DRVT | R1629H | BFIS3 | | | R1451H | HYPP | R1626L | LQT3 | R1620L | ID | | | | | | |
| | | | | BFIS3 | | | | HOKPP2 | R1626H | LQT3 | | | | | | | | |
| | | | R1629L | DEE11 | | | R1451L | SCM | | LQT3 | | | | | | | | |
| 1824 | | | | | R1624 | | | | R1626P | RWS | | | R1613 | | R1576 | | R1466 | |
| | | DRVT | R1638C | BFIS3 | | | R1460W | HYPP | | | R1629C | EIEE | | | | | | |
| | R1648H | GEFSP2 | | | | | | | | | | | | | | | | |
| | R1648C | DRVT | | | | | | | | | | | | | | | | |
| 1833 | R1648L | GEFSP2 | | | R1633 | | | | R1635 | | | | R1622 | | R1585 | | R1475 | |
| | | GEFSP2 | | | R1642C | DEE62 | | | R1644L | LQT3 | R1638C | NDD | | | | | | |
| | R1657C | GEFSP1 | | | | | | | | LQT3 | | | | | | | | |
| | | | | | | | | | R1644H | RWS | | | | | | | | |
| | | | | | | | | | | LQT3 | | | | | | | | |
| | | | | | | | | | | BRGDA1 | | | | | | | | |
| 1842 | | | R1647 | | | | R1469 | | R1644C | RWS | | | R1631 | | R1594 | | R1484 | |
| | A1669E | DRVT | A1659V | BFIS3 | | | | MYOSCN4A | | | A1650T | DEE13 | | PEXPD | | | | |
| | | | | | | | A1481D | HYPP | | | | | | PEPD | | | | |
| 1854 | | | | | A1654 | | | | A1656 | | | | A1643E | PERYTHM | A1606 | | S1496 | |

| MSA | $Na_v1.1$ MUT* | $Na_v1.1$ DIS** | $Na_v1.2$ MUT | $Na_v1.2$ DIS | $Na_v1.3$ MUT | $Na_v1.3$ DIS | $Na_v1.4$ MUT | $Na_v1.4$ DIS | $Na_v1.5$ MUT | $Na_v1.5$ DIS | $Na_v1.6$ MUT | $Na_v1.6$ DIS | $Na_v1.7$ MUT | $Na_v1.7$ DIS | $Na_v1.8$ MUT | $Na_v1.8$ DIS | $Na_v1.9$ MUT | $Na_v1.9$ DIS |
|---|---|---|---|---|---|---|---|---|---|---|---|---|---|---|---|---|---|---|
| | | | | | | | | | | | | | A1643T | PERYTHM | | | | |
| | G1725D | EIEE | | | | | G1537S | HYPP | G1712S | BRGDA1 | | | | | G1662S | BRGDA1 | | |
| | G1725S | DRVT | | | | | | | | | | | | | | | | |
| 1910 | G1725C | DRVT | G1715 | | G1710 | | | | | | G1706 | | G1699 | | | | G1552 | |
| | C1741Y | DRVT | | | | | | | C1728R | BRGDA1 | | | | | | | | |
| | C1741S | EIEE | | | | | | | C1728W | BRGDA1 | | | | | | | | |
| 1926 | C1741R | DRVT | C1731 | | C1726 | | C1553 | | | | C1721 | | C1715 | | C1678 | | -- | |
| | D1755Y | EIEE | | | | | | | | PFHB1A | | | | | | | | |
| | D1755G | DRVT | | | | | | | | LQT3 | | | | | | | | |
| | | | | | | | | | | BRGDA1 | | | | | | | | |
| 1940 | | | D1745 | | D1740 | | D1567 | | D1741N | CMD1E | D1735 | | D1729 | | D1691 | | N1573 | |
| | | | | | | | V1589M | PMC | V1763M | LQT3 | V1757I | EIEE | | | M1713V | BRGDA1 | | |
| | | | | | | | | PAM | | RWS | | | | | | FEPS2 | | |
| 1962 | V1777 | | V1767 | | V1762 | | | | | SIDS | | | V1751 | | | | V1595 | |
| | M1780T | EIEE | | | | | M1592I | HYPP | M1766L | LQT3 | M1760I | EIEE | | | | | | |
| | | DRVT | | | | | M1592V | NKPP | M1766L | RWS | M1760I | DEE13 | | | | | | |
| 1965 | | | M1770 | | M1765 | | | HYPP | | | | | M1754 | | M1716 | | M1598 | |

| MSA | Na$_v$1.1 | | Na$_v$1.2 | | Na$_v$1.3 | | Na$_v$1.4 | | Na$_v$1.5 | | Na$_v$1.6 | | Na$_v$1.7 | | Na$_v$1.8 | | Na$_v$1.9 | |
|---|---|---|---|---|---|---|---|---|---|---|---|---|---|---|---|---|---|---|
| | MUT* | DIS** | MUT | DIS | MUT | DIS | MUT | DIS | MUT | DIS | MUT | DIS | MUT | DIS | MUT | DIS | MUT | DIS |
| 1973 | N1788K | DRVT | N1778I | BFIS3 | N1773 | | N1600 | | N1774D | RWS | N1768D | DEE13 | N1762 | | N1724 | | N1606 | |
| | Y1809H | GEFSP2 | | | | | | | Y1795C | LQT3 | Y1789C | EIEE | | | | | Y1627H | HSAN7 |
| | | | | | | | | | | RWS | | | | | | | | |
| 1994 | | | Y1799 | | Y1794 | | Y1621 | | Y1795H | BRGDA1 | | | Y1783 | | Y1745 | | | |
| | | | | | | | R1652K | HYPP | R1826H | LQT3 | | | | | | | R1658H | HSAN7 |
| 2025 | N1840 | | L1830 | | L1825 | | | | R1826C | ATFB10 | R1820 | | L1814 | | R1776 | | R1658P | HSAN7 |
| | | | | EA9 | | | | | | | R1872W | DEE13 | | | | | | |
| | | | | BFIS3 | | | | | | | R1872L | DEE13 | | | | | | |
| | | | R1882G | NEIDEE | | | | | | | R1872Q | DEE13 | | | | | | |
| | | | | DEE11 | | | | | | | | | | | | | | |
| | | | R1882Q | NEIDEE | | | | | | | | | | | | | | |
| 2077 | R1892 | | R1882L | DEE11 | R1877 | | K1704 | | K1878 | | | | R1866 | | K1828 | | K1710 | |
| | | | R1902H | BFIS3 | | | | | R1898C | BRGDA1 | R1892H | EIEE | | | | | | |
| 2097 | R1912 | | R1902C | BFIS3 | R1897 | | R1724 | | R1898H | LQT3 | | | R1886 | | W1848 | | R1730 | |

*MUT: Mutation

**DIS: Diseases/Phenotype

†The mutations are denoted as [The original Amino Acid]MSA/Residue ID[Mutated Amino Acid] so R222Q mean an arginine in the position 222 mutated to a Glutamine, if there is no “Mutated Amino Acid” it means that there is no mutation in that position form that protein.

DEE6B: Developmental And Epileptic Encephalopathy 6B, DEE11: Early Infantile Epileptic Encephalopathy 11, FEB3A: Febrile Seizures, Familial, 3A, EPD: Pyridoxine-dependent epilepsy, FFEVF4: Epilepsy, Familial Focal, With Variable Foci 4, FEB3B: Febrile seizures, familial, 3B, SIDS: Sudden Infant Death Syndrome, VF1: Paroxysmal Familial Ventricular Fibrillation 1, RWS: Romano-Ward syndrome, EA9: Episodic Ataxia 9, CIP: Indifference To Pain, Congenital, Autosomal Recessive

Table S3: Mutations and their respective electrorheological characteristics cultured from literature

| | Protein | Mutation | Isoform | Mutation (MSA) | Phenotype | GoF/LoF | Gating Properties | Reference |
|---|---|---|---|---|---|---|---|---|
| 1 | Na$_v$1.1 | E78D | Na$_v$1.1:E78<br>Na$_v$1.1-2:E78<br>Na$_v$1.1-3:E78 | E85D | DRVT | | Imax:none<br>τrec:↓<br>Ip:none<br>V1/2act:none<br>V1/2inact:none<br>Iω:none<br>ExTraFol:none | (Kluckova et al., 2020) |
| 2 | Na$_v$1.1 | M145T | Na$_v$1.1:M145<br>Na$_v$1.1-2:M145<br>Na$_v$1.1-3:M145 | M152T | FEB3A*<br>GEFSP2*<br>FS | LOF | Imax:↓<br>τrec:none<br>Ip:↓<br>V1/2act:→<br>V1/2inact:none<br>Iω:none<br>ExTraFol:none | (Bender et al., 2013)<br>(Mantegazza et al., 2005) |
| 3 | Na$_v$1.1 | G177A | Na$_v$1.1:G177<br>Na$_v$1.1-2:G177<br>Na$_v$1.1-3:G177 | G188A | DRVT | LOF | Imax:↓<br>τrec:none<br>Ip:none<br>V1/2act:none<br>V1/2inact:none<br>Iω:none<br>ExTraFol:↓ | (Nissenkorn et al., 2019) |
| 4 | Na$_v$1.1 | T226M | Na$_v$1.1:T226<br>Na$_v$1.1-2:T226<br>Na$_v$1.1-3:T226 | T238M | DRVT*<br>DEE6B*<br>EIEE | Mixed | Imax:↓<br>τrec:none<br>Ip:none<br>V1/2act:←<br>V1/2inact:←<br>Iω:none<br>ExTraFol:none | (Berecki et al., 2019) |
| 5 | Na$_v$1.1 | I236V | Na$_v$1.1:I236<br>Na$_v$1.1-2:I236<br>Na$_v$1.1-3:I236 | I248V | NDEEMA | GOF | Imax:none<br>τrec:↑<br>Ip:none<br>V1/2act:none<br>V1/2inact:none<br>Iω:none<br>ExTraFol:none | (Brunklaus et al., 2022) |
| 6 | Na$_v$1.1 | D249E | Na$_v$1.1:D249<br>Na$_v$1.1-2:D249<br>Na$_v$1.1-3:D249 | D261E | DRVT | | Imax:none<br>τrec:↓<br>Ip:none<br>V1/2act:←<br>V1/2inact:← | (Kluckova et al., 2020) |

| | Protein | Mutation | Isoform | Mutation (MSA) | Phenotype | GoF/LoF | Gating Properties | Reference |
|---|---|---|---|---|---|---|---|---|
| | | | | | | | Iω:none<br>ExTraFol:none | |
| 7 | Naᵥ1.1 | S259R | Naᵥ1.1:S259<br>Naᵥ1.1-2:S259<br>Naᵥ1.1-3:S259 | S271R | DRVT | LOF | Imax:↓<br>τrec:none<br>Ip:none<br>V1/2act:none<br>V1/2inact:none<br>Iω:none<br>ExTraFol:none | (Nissenkorn et al., 2019) |
| 8 | Naᵥ1.1 | L263V | Naᵥ1.1:L263<br>Naᵥ1.1-2:L263<br>Naᵥ1.1-3:L263 | L275V | FHM3 | GOF | Imax:none<br>τrec:none<br>Ip:↑<br>V1/2act:←<br>V1/2inact:→<br>Iω:none<br>ExTraFol:none | (Kahlig et al., 2008) |
| 9 | Naᵥ1.1 | N301S | Naᵥ1.1:N301<br>Naᵥ1.1-2:N301<br>Naᵥ1.1-3:N301 | N314S | DRVT | LOF | Imax:↓<br>τrec:none<br>Ip:none<br>V1/2act:none<br>V1/2inact:none<br>Iω:none<br>ExTraFol:none | (Y.-J. J. Chen et al., 2015) |
| 10 | Naᵥ1.1 | Y426N | Naᵥ1.1:Y426<br>Naᵥ1.1-2:Y426<br>Naᵥ1.1-3:Y426 | Y483N | DRVT | | Imax:↓<br>τrec:↓<br>Ip:none<br>V1/2act:none<br>V1/2inact:←<br>Iω:none<br>ExTraFol:none | (Ohmori et al., 2006) |
| 11 | Naᵥ1.1 | E788K | Naᵥ1.1:E788<br>Naᵥ1.1-2:E777<br>Naᵥ1.1-3:E760 | E868K | EPI | LOF | Imax:↓<br>τrec:↓<br>Ip:none<br>V1/2act:→<br>V1/2inact:←<br>Iω:none<br>ExTraFol:none | (Kluckova et al., 2020) |
| 12 | Naᵥ1.1 | T808S | Naᵥ1.1:T808<br>Naᵥ1.1-2:T797<br>Naᵥ1.1-3:T780 | T888S | ICEGTC*<br>DRVT | | Imax:↑<br>τrec:↓<br>Ip:none<br>V1/2act:none<br>V1/2inact:none | (Rhodes et al., 2005) |

| | Protein | Mutation | Isoform | Mutation (MSA) | Phenotype | GoF/LoF | Gating Properties | Reference |
|---|---|---|---|---|---|---|---|---|
| | | | | | | | Iω:none<br>ExTraFol:none | |
| 13 | $Na_v1.1$ | R859C | $Na_v1.1$:R859<br>$Na_v1.1$-2:R848<br>$Na_v1.1$-3:R831 | R941C | DRVT*<br>GEFSP2*<br>FHM3 | LOF | Imax:↓<br>τrec:none<br>Ip:none<br>V1/2act:none<br>V1/2inact:none<br>Iω:none<br>ExTraFol:none | (Bechi et al., 2015) |
| 14 | $Na_v1.1$ | R859H | $Na_v1.1$:R859<br>$Na_v1.1$-2:R848<br>$Na_v1.1$-3:R831 | R941H | GEFSP2 | LOF | Imax:none<br>τrec:↓<br>Ip:↑<br>V1/2act:←<br>V1/2inact:←<br>Iω:none<br>ExTraFol:none | (Brunklaus et al., 2020) (Volkers et al., 2011) |
| 15 | $Na_v1.1$ | R865G | $Na_v1.1$:R865<br>$Na_v1.1$-2:R854<br>$Na_v1.1$-3:R837 | R947G | DRVT | GOF | Imax:none<br>τrec:↓<br>Ip:↑<br>V1/2act:←<br>V1/2inact:none<br>Iω:none<br>ExTraFol:none | (Brunklaus et al., 2020) (Volkers et al., 2011) |
| 16 | $Na_v1.1$ | M909K | $Na_v1.1$:M909<br>$Na_v1.1$-2:M898<br>$Na_v1.1$-3:M881 | M991K | GEFSP2 | LOF | Imax:↓<br>τrec:↓<br>Ip:none<br>V1/2act:→<br>V1/2inact:←<br>Iω:none<br>ExTraFol:none | (Kluckova et al., 2020) |
| 17 | $Na_v1.1$ | R946C | $Na_v1.1$:R946<br>$Na_v1.1$-2:R935<br>$Na_v1.1$-3:R918 | R1035C | EIEE*<br>GEFSP2*<br>DRVT | LOF | Imax:↓<br>τrec:none<br>Ip:none<br>V1/2act:none<br>V1/2inact:none<br>Iω:none<br>ExTraFol:none | (Brunklaus et al., 2020) |
| 18 | $Na_v1.1$ | R946H | $Na_v1.1$:R946<br>$Na_v1.1$-2:R935<br>$Na_v1.1$-3:R918 | R1035H | GEFSP2*<br>DRVT | LOF | Imax:↓<br>τrec:none<br>Ip:none<br>V1/2act:none<br>V1/2inact:none | (Brunklaus et al., 2020) |

| | Protein | Mutation | Isoform | Mutation (MSA) | Phenotype | GoF/LoF | Gating Properties | Reference |
|---|---|---|---|---|---|---|---|---|
| | | | | | | | Iω:none<br>ExTraFol:none | |
| 19 | $Na_v$1.1 | G979R | $Na_v$1.1:G979<br>$Na_v$1.1-2:G968<br>$Na_v$1.1-3:G951 | G1070R | DRVT*<br>ICEGTC | LOF | Imax:↓<br>τrec:none<br>Ip:none<br>V1/2act:none<br>V1/2inact:none<br>Iω:none<br>ExTraFol:none | (Rhodes et al., 2005) |
| 20 | $Na_v$1.1 | V983A | $Na_v$1.1:V983<br>$Na_v$1.1-2:V972<br>$Na_v$1.1-3:V955 | V1074A | ICEGTC*<br>DRVT | | Imax:↓<br>τrec:↓<br>Ip:none<br>V1/2act:→<br>V1/2inact:←<br>Iω:none<br>ExTraFol:none | (Rhodes et al., 2005) |
| 21 | $Na_v$1.1 | N1011I | $Na_v$1.1:N1011<br>$Na_v$1.1-2:N1000<br>$Na_v$1.1-3:N983 | N1103I | ICEGTC*<br>DRVT | | Imax:↓<br>τrec:↓<br>Ip:none<br>V1/2act:none<br>V1/2inact:←<br>Iω:none<br>ExTraFol:none | (Rhodes et al., 2005) |
| 22 | $Na_v$1.1 | T1174S | $Na_v$1.1:T1174<br>$Na_v$1.1-2:T1163<br>$Na_v$1.1-3:T1146 | T1352S | FHM3 | GOF | Imax:↓<br>τrec:none<br>Ip:↑<br>V1/2act:→<br>V1/2inact:none<br>Iω:none<br>ExTraFol:none | (Cestèle, Labate, et al., 2013) |
| 23 | $Na_v$1.1 | W1204R | $Na_v$1.1:W1204<br>$Na_v$1.1-2:W1193<br>$Na_v$1.1-3:W1176 | W1382R | GEFSP2 | LOF | Imax:none<br>τrec:none<br>Ip:none<br>V1/2act:←<br>V1/2inact:←<br>Iω:none<br>ExTraFol:none | (Spampanato et al., 2003) |
| 24 | $Na_v$1.1 | M1267I | $Na_v$1.1:M1267<br>$Na_v$1.1-2:M1256<br>$Na_v$1.1-3:M1239 | M1445I | DRVT | LOF | Imax:none<br>τrec:↓<br>Ip:none<br>V1/2act:←<br>V1/2inact:none | (Nissenkorn et al., 2019) |

|  | Protein | Mutation | Isoform | Mutation (MSA) | Phenotype | GoF/LoF | Gating Properties | Reference |
|---|---|---|---|---|---|---|---|---|
|  |  |  |  |  |  |  | Iω:none<br>ExTraFol:none |  |
| 25 | $Na_v1.1$ | W1271R | $Na_v1.1$:W1271<br>$Na_v1.1$-2:W1260<br>$Na_v1.1$-3:W1243 | W1449R | DRVT | LOF | Imax:↓<br>τrec:none<br>Ip:none<br>V1/2act:→<br>V1/2inact:←<br>Iω:none<br>ExTraFol:none | (Fang et al., 2022) |
| 26 | $Na_v1.1$ | S1328P | $Na_v1.1$:S1328<br>$Na_v1.1$-2:S1317<br>$Na_v1.1$-3:S1300 | S1506P | ICEGTC*<br>DRVT | | Imax:↓<br>τrec:none<br>Ip:none<br>V1/2act:←<br>V1/2inact:←<br>Iω:none<br>ExTraFol:none | (Y. Sun et al., 2016) |
| 27 | $Na_v1.1$ | V1353L | $Na_v1.1$:V1353<br>$Na_v1.1$-2:V1342<br>$Na_v1.1$-3:V1325 | V1531L | GEFSP2*<br>EIEE | Mixed | Imax:↓<br>τrec:↑<br>Ip:none<br>V1/2act:none<br>V1/2inact:→<br>Iω:none<br>ExTraFol:none | (Berecki et al., 2019) |
| 28 | $Na_v1.1$ | F1415I | $Na_v1.1$:F1415<br>$Na_v1.1$-2:F1404<br>$Na_v1.1$-3:F1387 | F1596I | DRVT | | Imax:↑<br>τrec:none<br>Ip:none<br>V1/2act:none<br>V1/2inact:→<br>Iω:none<br>ExTraFol:none | (Jiao et al., 2013) |
| 29 | $Na_v1.1$ | N1417H | $Na_v1.1$:N1417<br>$Na_v1.1$-2:N1406<br>$Na_v1.1$-3:N1389 | N1598H | FS | | Imax:none<br>τrec:↓<br>Ip:↑<br>V1/2act:none<br>V1/2inact:←<br>Iω:none<br>ExTraFol:none | (Suleimanova et al., 2021) |
| 30 | $Na_v1.1$ | Q1489H | $Na_v1.1$:Q1489<br>$Na_v1.1$-2:Q1478<br>$Na_v1.1$-3:Q1461 | Q1670H | FHM3 | GOF | Imax:↑<br>τrec:none<br>Ip:↑<br>V1/2act:none<br>V1/2inact:none | (Barbieri et al., 2019) |

| | Protein | Mutation | Isoform | Mutation (MSA) | Phenotype | GoF/LoF | Gating Properties | Reference |
|---|---|---|---|---|---|---|---|---|
| | | | | | | | Iω:none<br>ExTraFol:none | |
| 31 | $Na_v1.1$ | Q1489K | $Na_v1.1$:Q1489<br>$Na_v1.1$-2:Q1478<br>$Na_v1.1$-3:Q1461 | Q1670K | FHM3 | | Imax:none<br>τrec:none<br>Ip:↑<br>V1/2act:←<br>V1/2inact:none<br>Iω:none<br>ExTraFol:none | (Kahlig et al., 2008)<br>(Cestèle et al., 2008) |
| 32 | $Na_v1.1$ | I1498M | $Na_v1.1$:I1498<br>$Na_v1.1$-2:I1487<br>$Na_v1.1$-3:I1470 | I1679M | FHM3 | LOF | Imax:↓<br>τrec:none<br>Ip:↓<br>V1/2act:none<br>V1/2inact:none<br>Iω:none<br>ExTraFol:none | (Barbieri et al., 2019) |
| 33 | $Na_v1.1$ | I1498M | $Na_v1.1$:I1498<br>$Na_v1.1$-2:I1487<br>$Na_v1.1$-3:I1470 | I1679M | FHM3 | | Imax:↓<br>τrec:↑<br>Ip:↑<br>V1/2act:none<br>V1/2inact:→<br>Iω:none<br>ExTraFol:none | (Brunklaus et al., 2022) |
| 34 | $Na_v1.1$ | I1498T | $Na_v1.1$:I1498<br>$Na_v1.1$-2:I1487<br>$Na_v1.1$-3:I1470 | I1679T | MD/EIDEE | | Imax:none<br>τrec:↑<br>Ip:none<br>V1/2act:none<br>V1/2inact:→<br>Iω:none<br>ExTraFol:none | (Brunklaus et al., 2022) |
| 35 | $Na_v1.1$ | F1499L | $Na_v1.1$:F1499<br>$Na_v1.1$-2:F1488<br>$Na_v1.1$-3:F1471 | F1680L | FHM3 | GOF | Imax:none<br>τrec:none<br>Ip:↑<br>V1/2act:none<br>V1/2inact:→<br>Iω:none<br>ExTraFol:none | (Barbieri et al., 2019) |
| 36 | $Na_v1.1$ | M1500V | $Na_v1.1$:M1500<br>$Na_v1.1$-2:M1489<br>$Na_v1.1$-3:M1472 | M1681V | FHM3 | Mixed | Imax:↓<br>τrec:none<br>Ip:↑<br>V1/2act:none<br>V1/2inact:none | (Barbieri et al., 2019) |

| | Protein | Mutation | Isoform | Mutation (MSA) | Phenotype | GoF/LoF | Gating Properties | Reference |
|---|---|---|---|---|---|---|---|---|
| | | | | | | | Iω:none<br>ExTraFol:none | |
| 37 | $Na_v1.1$ | R1575C | $Na_v1.1$:R1575<br>$Na_v1.1$-2:R1564<br>$Na_v1.1$-3:R1547 | R1756C | Rasmussen encephalitis | | Imax:none<br>τrec:none<br>Ip:↑<br>V1/2act:none<br>V1/2inact:→<br>Iω:none<br>ExTraFol:none | (Ohmori et al., 2008) |
| 38 | $Na_v1.1$ | E1587K | $Na_v1.1$:E1587<br>$Na_v1.1$-2:E1576<br>$Na_v1.1$-3:E1559 | E1768K | EPI | LOF | Imax:↓<br>τrec:none<br>Ip:none<br>V1/2act:none<br>V1/2inact:none<br>Iω:none<br>ExTraFol:none | (Kluckova et al., 2020) |
| 39 | $Na_v1.1$ | R1596C | $Na_v1.1$:R1596<br>$Na_v1.1$-2:R1585<br>$Na_v1.1$-3:R1568 | R1777C | DRVT<br>GEFSP2<br>FS | LOF | Imax:↓<br>τrec:none<br>Ip:none<br>V1/2act:none<br>V1/2inact:none<br>Iω:none<br>ExTraFol:none | (Kluckova et al., 2020) |
| 40 | $Na_v1.1$ | V1611F | $Na_v1.1$:V1611<br>$Na_v1.1$-2:V1600<br>$Na_v1.1$-3:V1583 | V1792F | ICEGTC*<br>GEFSP2<br>DRVT | | Imax:none<br>τrec:none<br>Ip:↑<br>V1/2act:←<br>V1/2inact:←<br>Iω:none<br>ExTraFol:none | (Rhodes et al., 2005) |
| 41 | $Na_v1.1$ | E1623A | $Na_v1.1$:E1623<br>$Na_v1.1$-2:E1612<br>$Na_v1.1$-3:E1595 | E1804A | EPI | LOF | Imax:↓<br>τrec:↓<br>Ip:none<br>V1/2act:none<br>V1/2inact:←<br>Iω:none<br>ExTraFol:none | (Su et al., 2022) |
| 42 | $Na_v1.1$ | E1623D | $Na_v1.1$:E1623<br>$Na_v1.1$-2:E1612<br>$Na_v1.1$-3:E1595 | E1804D | EPI | LOF | Imax:none<br>τrec:none<br>Ip:↓<br>V1/2act:←<br>V1/2inact:none | (Su et al., 2022) |

| | Protein | Mutation | Isoform | Mutation (MSA) | Phenotype | GoF/LoF | Gating Properties | Reference |
|---|---|---|---|---|---|---|---|---|
| | | | | | | | Iω:none<br>ExTraFol:none | |
| 43 | $Na_v1.1$ | E1623K | $Na_v1.1$:E1623<br>$Na_v1.1$-2:E1612<br>$Na_v1.1$-3:E1595 | E1804K | EPI | LOF | Imax:↓<br>τrec:none<br>Ip:↓<br>V1/2act:none<br>V1/2inact:←<br>Iω:none<br>ExTraFol:none | (Su et al., 2022) |
| 44 | $Na_v1.1$ | E1623Q | $Na_v1.1$:E1623<br>$Na_v1.1$-2:E1612<br>$Na_v1.1$-3:E1595 | E1804Q | EPI | GOF | Imax:↑<br>τrec:none<br>Ip:none<br>V1/2act:←<br>V1/2inact:←<br>Iω:none<br>ExTraFol:none | (Su et al., 2022) |
| 45 | $Na_v1.1$ | E1623V | $Na_v1.1$:E1623<br>$Na_v1.1$-2:E1612<br>$Na_v1.1$-3:E1595 | E1804V | EPI | LOF | Imax:none<br>τrec:none<br>Ip:↓<br>V1/2act:none<br>V1/2inact:←<br>Iω:none<br>ExTraFol:none | (Su et al., 2022) |
| 46 | $Na_v1.1$ | L1624P | $Na_v1.1$:L1624<br>$Na_v1.1$-2:L1613<br>$Na_v1.1$-3:L1596 | L1805P | FHM3 | GOF | Imax:none<br>τrec:↑<br>Ip:↑<br>V1/2act:none<br>V1/2inact:→<br>Iω:none<br>ExTraFol:none | (Fan et al., 2016) |
| 47 | $Na_v1.1$ | P1632S | $Na_v1.1$:P1632<br>$Na_v1.1$-2:P1621<br>$Na_v1.1$-3:P1604 | P1817S | ICEGTC*<br>DRVT | LOF | Imax:none<br>τrec:none<br>Ip:↑<br>V1/2act:←<br>V1/2inact:←<br>Iω:none<br>ExTraFol:none | (Rhodes et al., 2005) |
| 48 | $Na_v1.1$ | R1636Q | $Na_v1.1$:R1636<br>$Na_v1.1$-2:R1625<br>$Na_v1.1$-3:R1608 | R1821Q | EIEE*<br>MD<br>EIDEE | | Imax:none<br>τrec:↑<br>Ip:none<br>V1/2act:none<br>V1/2inact:none | (Brunklaus et al., 2022) |

| | Protein | Mutation | Isoform | Mutation (MSA) | Phenotype | GoF/LoF | Gating Properties | Reference |
|---|---|---|---|---|---|---|---|---|
| | | | | | | | Iω:none<br>ExTraFol:none | |
| 49 | $Na_v$1.1 | R1648H | $Na_v$1.1:R1648<br>$Na_v$1.1-2:R1637<br>$Na_v$1.1-3:R1620 | R1833H | DRVT*<br>GEFSP2 | LOF | Imax:↓<br>τrec:↑<br>Ip:↑<br>V1/2act:none<br>V1/2inact:none<br>Iω:none<br>ExTraFol:none | (Hedrich et al., 2014) |
| 50 | $Na_v$1.1 | L1649Q | $Na_v$1.1:L1649<br>$Na_v$1.1-2:L1638<br>$Na_v$1.1-3:L1621 | L1834Q | FHM3 | GOF | Imax:none<br>τrec:↑<br>Ip:↑<br>V1/2act:→<br>V1/2inact:→<br>Iω:none<br>ExTraFol:none | (Cestèle, Schiavon, et al., 2013) |
| 51 | $Na_v$1.1 | I1656M | $Na_v$1.1:I1656<br>$Na_v$1.1-2:I1645<br>$Na_v$1.1-3:I1628 | I1841M | GEFSP2 | LOF | Imax:↓<br>τrec:↓<br>Ip:none<br>V1/2act:→<br>V1/2inact:←<br>Iω:none<br>ExTraFol:none | (Lossin et al., 2003) |
| 52 | $Na_v$1.1 | R1657C | $Na_v$1.1:R1657<br>$Na_v$1.1-2:R1646<br>$Na_v$1.1-3:R1629 | R1842C | GEFSP2 | LOF | Imax:↓<br>τrec:↑<br>Ip:none<br>V1/2act:→<br>V1/2inact:none<br>Iω:none<br>ExTraFol:none | (Lossin et al., 2003) |
| 53 | $Na_v$1.1 | F1661L | $Na_v$1.1:F1661<br>$Na_v$1.1-2:F1650<br>$Na_v$1.1-3:F1633 | F1846L | FHM3 | GOF | Imax:none<br>τrec:none<br>Ip:↑<br>V1/2act:none<br>V1/2inact:none<br>Iω:none<br>ExTraFol:none | (Barbieri et al., 2019) |
| 54 | $Na_v$1.1 | F1661S | $Na_v$1.1:F1661<br>$Na_v$1.1-2:F1650<br>$Na_v$1.1-3:F1633 | F1846S | DRVT | | Imax:↓<br>τrec:↑<br>Ip:↑<br>V1/2act:none<br>V1/2inact:→ | (Rhodes et al., 2004)<br>(Thompson et al., 2012) |

| | Protein | Mutation | Isoform | Mutation (MSA) | Phenotype | GoF/LoF | Gating Properties | Reference |
|---|---|---|---|---|---|---|---|---|
| | | | | | | | Iω:none<br>ExTraFol:↓ | |
| 55 | $Na_v$1.1 | M1664K | $Na_v$1.1:M1664<br>$Na_v$1.1-2:M1653<br>$Na_v$1.1-3:M1636 | M1849K | GEFSP2<br>DRVT | | Imax:none<br>τrec:none<br>Ip:none<br>V1/2act:none<br>V1/2inact:none<br>Iω:none<br>ExTraFol:↓ | (Bechi et al., 2015) |
| 56 | $Na_v$1.1 | L1670W | $Na_v$1.1:L1670<br>$Na_v$1.1-2:L1659<br>$Na_v$1.1-3:L1642 | L1855W | FHM3 | GOF | Imax:↓<br>τrec:none<br>Ip:↑<br>V1/2act:→<br>V1/2inact:→<br>Iω:none<br>ExTraFol:↓ | (Dhifallah et al., 2018)<br>(Bertelli et al., 2019) |
| 57 | $Na_v$1.1 | G1674R | $Na_v$1.1:G1674<br>$Na_v$1.1-2:G1663<br>$Na_v$1.1-3:G1646 | G1859R | DRVT | LOF | Imax:↓<br>τrec:none<br>Ip:none<br>V1/2act:none<br>V1/2inact:none<br>Iω:none<br>ExTraFol:none | (Rhodes et al., 2004) |
| 58 | $Na_v$1.1 | A1685D | $Na_v$1.1:A1685<br>$Na_v$1.1-2:A1674<br>$Na_v$1.1-3:A1657 | A1870D | DRVT | | Imax:↓<br>τrec:none<br>Ip:none<br>V1/2act:none<br>V1/2inact:none<br>Iω:none<br>ExTraFol:none | (Sugiura et al., 2012) |
| 59 | $Na_v$1.1 | A1685V | $Na_v$1.1:A1685<br>$Na_v$1.1-2:A1674<br>$Na_v$1.1-3:A1657 | A1870V | EIEE*<br>GEFSP2 | | Imax:↓<br>τrec:none<br>Ip:none<br>V1/2act:←<br>V1/2inact:←<br>Iω:none<br>ExTraFol:none | (Sugiura et al., 2012) |
| 60 | $Na_v$1.1 | T1709I | $Na_v$1.1:T1709<br>$Na_v$1.1-2:T1698<br>$Na_v$1.1-3:T1681 | T1894I | ICEGTC*<br>DRVT<br>GEFSP2 | | Imax:↓<br>τrec:none<br>Ip:none<br>V1/2act:none<br>V1/2inact:none | (Rhodes et al., 2005) |

| | Protein | Mutation | Isoform | Mutation (MSA) | Phenotype | GoF/LoF | Gating Properties | Reference |
|---|---|---|---|---|---|---|---|---|
| | | | | | | | Iω:none<br>ExTraFol:none | |
| 61 | Na$_v$1.1 | R1916G | Na$_v$1.1:R1927<br>Na$_v$1.1-2:R1916<br>Na$_v$1.1-3:R1899 | R1927G | GEFSP2 | LOF | Imax:none<br>τrec:none<br>Ip:↓<br>V1/2act:none<br>V1/2inact:←<br>Iω:none<br>ExTraFol:↓ | (Bechi et al., 2015) |
| 62 | Na$_v$1.1 | F1774S | Na$_v$1.1:F1774<br>Na$_v$1.1-2:F1763<br>Na$_v$1.1-3:F1746 | F1959S | FHM3 | | Imax:none<br>τrec:↑<br>Ip:↑<br>V1/2act:none<br>V1/2inact:none<br>Iω:none<br>ExTraFol:none | (Bertelli et al., 2019) |
| 63 | Na$_v$1.1 | A1783V | Na$_v$1.1:A1783<br>Na$_v$1.1-2:A1772<br>Na$_v$1.1-3:A1755 | A1968V | DRVT | LOF | Imax:none<br>τrec:none<br>Ip:none<br>V1/2act:→<br>V1/2inact:none<br>Iω:none<br>ExTraFol:none | (Layer et al., 2021) |
| 64 | Na$_v$1.1 | F1808L | Na$_v$1.1:F1808<br>Na$_v$1.1-2:F1797<br>Na$_v$1.1-3:F1780 | F1993L | DRVT*<br>ICEGTC | | Imax:↓<br>τrec:none<br>Ip:↑<br>V1/2act:←<br>V1/2inact:←<br>Iω:none<br>ExTraFol:none | (Rhodes et al., 2005) |
| 65 | Na$_v$1.1 | T1909I | Na$_v$1.1:T1909<br>Na$_v$1.1-2:T1898<br>Na$_v$1.1-3:T1881 | T2094I | EIEE*<br>DRVT | | Imax:↓<br>τrec:none<br>Ip:↑<br>V1/2act:none<br>V1/2inact:none<br>Iω:none<br>ExTraFol:none | (Ohmori et al., 2006) |
| 66 | Na$_v$1.1 | Q1923R | Na$_v$1.1:Q1923<br>Na$_v$1.1-2:Q1912<br>Na$_v$1.1-3:Q1895 | Q2108R | DRVT | LOF | Imax:↓<br>τrec:none<br>Ip:none<br>V1/2act:none<br>V1/2inact:none | (Nissenkorn et al., 2019) |

| | Protein | Mutation | Isoform | Mutation (MSA) | Phenotype | GoF/LoF | Gating Properties | Reference |
|---|---|---|---|---|---|---|---|---|
| | | | | | | | Iω:none<br>ExTraFol:none | |
| 67 | $Na_v$1.1 | Q1923R | $Na_v$1.1:Q1923<br>$Na_v$1.1-2:Q1912<br>$Na_v$1.1-3:Q1895 | Q2108R | FS | | Imax:↑<br>τrec:none<br>Ip:none<br>V1/2act:none<br>V1/2inact:→<br>Iω:none<br>ExTraFol:none | (Jiao et al., 2013) |
| 68 | $Na_v$1.1 | T1934I | $Na_v$1.1:T1934<br>$Na_v$1.1-2:T1923<br>$Na_v$1.1-3:T1906 | T2119I | EE | LOF | Imax:↓<br>τrec:↓<br>Ip:none<br>V1/2act:none<br>V1/2inact:none<br>Iω:none<br>ExTraFol:none | (Kluckova et al., 2020) |
| 69 | $Na_v$1.2 | D12N | $Na_v$1.2:D12<br>$Na_v$1.2-2:D12 | D15N | ASD | LOF | Imax:↓<br>τrec:none<br>Ip:none<br>V1/2act:none<br>V1/2inact:none<br>Iω:none<br>ExTraFol:none | (Ben-Shalom et al., 2017) |
| 70 | $Na_v$1.2 | D82G | $Na_v$1.2:D82<br>$Na_v$1.2-2:D82 | D88G | ASD | LOF | Imax:↓<br>τrec:none<br>Ip:none<br>V1/2act:none<br>V1/2inact:none<br>Iω:none<br>ExTraFol:none | (Ben-Shalom et al., 2017) |
| 71 | $Na_v$1.2 | R188W | $Na_v$1.2:R188<br>$Na_v$1.2-2:R188 | R198W | BFIS3*<br>FS | GOF | Imax:none<br>τrec:none<br>Ip:none<br>V1/2act:none<br>V1/2inact:←<br>Iω:none<br>ExTraFol:none | (Sugawara et al., 2001) |
| 72 | $Na_v$1.2 | R223Q | $Na_v$1.2:R223<br>$Na_v$1.2-2:R223 | R234Q | BFIS3 | GOF | Imax:none<br>τrec:none<br>Ip:none<br>V1/2act:→<br>V1/2inact:→ | (Scalmani et al., 2006) |

| | Protein | Mutation | Isoform | Mutation (MSA) | Phenotype | GoF/LoF | Gating Properties | Reference |
|---|---|---|---|---|---|---|---|---|
| | | | | | | | Iω:none<br>ExTraFol:none | |
| 73 | $Na_v1.2$ | T236S | $Na_v1.2$:T236<br>$Na_v1.2$-2:T236 | T247S | DEE11*<br>EOEE | Mixed | Imax:none<br>τrec:none<br>Ip:none<br>V1/2act:←<br>V1/2inact:none<br>Iω:none<br>ExTraFol:none | (Thompson et al., 2020) |
| 74 | $Na_v1.2$ | M252V | $Na_v1.2$:M252<br>$Na_v1.2$-2:M252 | M263V | BFIS3 | GOF | Imax:none<br>τrec:none<br>Ip:↑<br>V1/2act:none<br>V1/2inact:none<br>Iω:none<br>ExTraFol:none | (Liao, Deprez, et al., 2010) |
| 75 | $Na_v1.2$ | A263V | $Na_v1.2$:A263<br>$Na_v1.2$-2:A263 | A274V | EA9*<br>BFIS3*<br>DEE11*<br>NEIDEE*<br>ICDEE | GOF | Imax:none<br>τrec:none<br>Ip:↑<br>V1/2act:→<br>V1/2inact:→<br>Iω:none<br>ExTraFol:none | (Liao, Anttonen, et al., 2010) (Schwarz et al., 2015) |
| 76 | $Na_v1.2$ | R379H | $Na_v1.2$:R379<br>$Na_v1.2$-2:R379 | R434H | ASD | LOF | Imax:↓<br>τrec:none<br>Ip:none<br>V1/2act:none<br>V1/2inact:none<br>Iω:none<br>ExTraFol:none | (Ben-Shalom et al., 2017) |
| 77 | $Na_v1.2$ | V423L | $Na_v1.2$:V423<br>$Na_v1.2$-2:V423 | V478L | NEIDEE*<br>OS | GOF | Imax:none<br>τrec:none<br>Ip:↑<br>V1/2act:none<br>V1/2inact:none<br>Iω:none<br>ExTraFol:none | (Wolff et al., 2017) |
| 78 | $Na_v1.2$ | A575V | $Na_v1.2$:A575<br>$Na_v1.2$-2:A575 | A640V | NEIDEE | | Imax:none<br>τrec:none<br>Ip:none<br>V1/2act:none<br>V1/2inact:← | (Ogiwara et al., 2009) |

| | Protein | Mutation | Isoform | Mutation (MSA) | Phenotype | GoF/LoF | Gating Properties | Reference |
|---|---|---|---|---|---|---|---|---|
| | | | | | | | Iω:none<br>ExTraFol:none | |
| 79 | $Na_v$1.2 | T773I | $Na_v$1.2:T773<br>$Na_v$1.2-2:T773 | T862I | DEE11*<br>DEE | GoF | Imax:none<br>τrec:none<br>Ip:↑<br>V1/2act:←<br>V1/2inact:none<br>Iω:none<br>ExTraFol:none | (Lauxmann et al., 2018) |
| 80 | $Na_v$1.2 | R853Q | $Na_v$1.2:R853<br>$Na_v$1.2-2:R853 | R944Q | BFIS3*<br>DEE11*<br>ICDEE*<br>West syndrome | LOF | Imax:↓<br>τrec:↓<br>Ip:↓<br>V1/2act:none<br>V1/2inact:←<br>Iω:↑<br>ExTraFol:none | (Mason et al., 2019)<br>(Ganguly et al., 2021) |
| 81 | $Na_v$1.2 | R853Q | $Na_v$1.2:R853<br>$Na_v$1.2-2:R853 | R944Q | BFIS3*<br>DEE11*<br>ICDEE*<br>NEIDEE | GOF | Imax:↑<br>τrec:none<br>Ip:none<br>V1/2act:←<br>V1/2inact:←<br>Iω:none<br>ExTraFol:none | (Berecki et al., 2018) |
| 82 | $Na_v$1.2 | G879R | $Na_v$1.2:G879<br>$Na_v$1.2-2:G879 | G970R | EIMFS | LOF | Imax:↓<br>τrec:none<br>Ip:none<br>V1/2act:←<br>V1/2inact:→<br>Iω:none<br>ExTraFol:↓ | (X. R. Yang et al., 2022) |
| 83 | $Na_v$1.2 | G899S | $Na_v$1.2:G899<br>$Na_v$1.2-2:G899 | G990S | ICDEE*<br>Other | LOF | Imax:none<br>τrec:none<br>Ip:none<br>V1/2act:→<br>V1/2inact:none<br>Iω:none<br>ExTraFol:none | (Wolff et al., 2017) |
| 84 | $Na_v$1.2 | R937C | $Na_v$1.2:R937<br>$Na_v$1.2-2:R937 | R1035C | BFIS3*<br>ID*<br>ASD | LOF | Imax:↓<br>τrec:none<br>Ip:none<br>V1/2act:none<br>V1/2inact:none | (Ben-Shalom et al., 2017) |

| | Protein | Mutation | Isoform | Mutation (MSA) | Phenotype | GoF/LoF | Gating Properties | Reference |
|---|---|---|---|---|---|---|---|---|
| | | | | | | | Iω:none<br>ExTraFol:none | |
| 85 | $Na_v$1.2 | R937H | $Na_v$1.2:R937<br>$Na_v$1.2-2:R937 | R1035H | ASD | LOF | Imax:↓<br>τrec:none<br>Ip:none<br>V1/2act:none<br>V1/2inact:none<br>Iω:none<br>ExTraFol:none | (Ben-Shalom et al., 2017) |
| 86 | $Na_v$1.2 | E999K | $Na_v$1.2:E999<br>$Na_v$1.2-2:E999 | E1100K | BFIS3*<br>DEE11*<br>EOEE | GOF | Imax:none<br>τrec:none<br>Ip:none<br>V1/2act:←<br>V1/2inact:none<br>Iω:none<br>ExTraFol:none | (Thompson et al., 2020)<br>(Miao et al., 2020) |
| 87 | $Na_v$1.2 | E1211K | $Na_v$1.2:E1211<br>$Na_v$1.2-2:E1211 | E1399K | BFIS3*<br>DEE11*<br>NEIDEE | GOF | Imax:none<br>τrec:none<br>Ip:none<br>V1/2act:←<br>V1/2inact:←<br>Iω:none<br>ExTraFol:none | (Ogiwara et al., 2009) |
| 88 | $Na_v$1.2 | R1312T | $Na_v$1.2:R1312<br>$Na_v$1.2-2:R1312 | R1500T | DEE11*<br>DRVT | | Imax:none<br>τrec:↓<br>Ip:none<br>V1/2act:←<br>V1/2inact:←<br>Iω:none<br>ExTraFol:none | (Lossin, Shi, et al., 2012) |
| 89 | $Na_v$1.2 | R1319Q | $Na_v$1.2:R1319<br>$Na_v$1.2-2:R1319 | R1507Q | BFIS3 | | Imax:↓<br>τrec:↓<br>Ip:none<br>V1/2act:→<br>V1/2inact:→<br>Iω:none<br>ExTraFol:↓ | (Misra et al., 2008)<br>(Scalmani et al., 2006) |
| 90 | $Na_v$1.2 | L1330F | $Na_v$1.2:L1330<br>$Na_v$1.2-2:L1330 | L1518F | BFIS3 | GOF | Imax:none<br>τrec:none<br>Ip:none<br>V1/2act:none<br>V1/2inact:→ | (Scalmani et al., 2006) |

|  | Protein | Mutation | Isoform | Mutation (MSA) | Phenotype | GoF/LoF | Gating Properties | Reference |
|---|---|---|---|---|---|---|---|---|
|  |  |  |  |  |  |  | Iω:none<br>ExTraFol:none |  |
| 91 | Na$_v$1.2 | S1336Y | Na$_v$1.2:S1336<br>Na$_v$1.2-2:S1336 | S1524Y | DEE11*<br>EOEE | MIXed | Imax:↓<br>τrec:none<br>Ip:none<br>V1/2act:←<br>V1/2inact:→<br>Iω:none<br>ExTraFol:none | (Thompson et al., 2020) |
| 92 | Na$_v$1.2 | N1339D | Na$_v$1.2:N1339<br>Na$_v$1.2-2:N1339 | N1527D | DEE | | Imax:none<br>τrec:↑<br>Ip:none<br>V1/2act:←<br>V1/2inact:none<br>Iω:none<br>ExTraFol:none | (Miao et al., 2020) |
| 93 | Na$_v$1.2 | L1342P | Na$_v$1.2:L1342<br>Na$_v$1.2-2:L1342 | L1530P | DEE11*<br>EOEE | GOF | Imax:↑<br>τrec:↓<br>Ip:↑<br>V1/2act:←<br>V1/2inact:←<br>Iω:none<br>ExTraFol:none | (Begemann et al., 2019)<br>(Que et al., 2021) |
| 94 | Na$_v$1.2 | C1386R | Na$_v$1.2:C1386<br>Na$_v$1.2-2:C1386 | C1576R | ASD | LOF | Imax:↓<br>τrec:none<br>Ip:none<br>V1/2act:none<br>V1/2inact:none<br>Iω:none<br>ExTraFol:none | (Ben-Shalom et al., 2017) |
| 95 | Na$_v$1.2 | T1420M | Na$_v$1.2:T1420<br>Na$_v$1.2-2:T1420 | T1611M | ASD | LOF | Imax:↓<br>τrec:none<br>Ip:none<br>V1/2act:none<br>V1/2inact:none<br>Iω:none<br>ExTraFol:none | (Ben-Shalom et al., 2017) |
| 96 | Na$_v$1.2 | I1473M | Na$_v$1.2:I1473<br>Na$_v$1.2-2:I1473 | I1664M | DEE11*<br>NEIDEE | GOF | Imax:none<br>τrec:none<br>Ip:none<br>V1/2act:←<br>V1/2inact:none | (Ogiwara et al., 2009) |

| | Protein | Mutation | Isoform | Mutation (MSA) | Phenotype | GoF/LoF | Gating Properties | Reference |
|---|---|---|---|---|---|---|---|---|
| | | | | | | | Iω:none<br>ExTraFol:none | |
| 97 | $Na_v1.2$ | L1563V | $Na_v1.2$:L1563<br>$Na_v1.2$-2:L1563 | L1754V | BFIS3 | GOF | Imax:↓<br>τrec:↑<br>Ip:none<br>V1/2act:←<br>V1/2inact:→<br>Iω:none<br>ExTraFol:↓ | (Misra et al., 2008)<br>(Scalmani et al., 2006) |
| 98 | $Na_v1.2$ | L1563V | $Na_v1.2$:L1563<br>$Na_v1.2$-2:L1563 | L1754V | BFIS3 | GOF | Imax:↑<br>τrec:↑<br>Ip:↑<br>V1/2act:→<br>V1/2inact:→<br>Iω:none<br>ExTraFol:none | (Begemann et al., 2019)<br>(Berecki et al., 2018) |
| 99 | $Na_v1.2$ | I1571T | $Na_v1.2$:I1571<br>$Na_v1.2$-2:I1571 | I1762T | DEE | | Imax:none<br>τrec:↑<br>Ip:none<br>V1/2act:←<br>V1/2inact:none<br>Iω:none<br>ExTraFol:none | (Miao et al., 2020) |
| 100 | $Na_v1.2$ | Y1589C | $Na_v1.2$:Y1589<br>$Na_v1.2$-2:Y1589 | Y1780C | BFIS3 | GOF | Imax:none<br>τrec:none<br>Ip:↑<br>V1/2act:none<br>V1/2inact:→<br>Iω:none<br>ExTraFol:none | (Lauxmann et al., 2013) |
| 101 | $Na_v1.2$ | F1597L | $Na_v1.2$:F1597<br>$Na_v1.2$-2:F1597 | F1788L | NEIDEE*<br>EIMFS | GOF | Imax:none<br>τrec:↑<br>Ip:none<br>V1/2act:←<br>V1/2inact:none<br>Iω:none<br>ExTraFol:none | (Wolff et al., 2017) |
| 102 | $Na_v1.2$ | P1622S | $Na_v1.2$:P1622<br>$Na_v1.2$-2:P1622 | P1817S | ICDEE*<br>MAE | LOF | Imax:none<br>τrec:↓<br>Ip:none<br>V1/2act:none<br>V1/2inact:← | (Wolff et al., 2017) |

| | Protein | Mutation | Isoform | Mutation (MSA) | Phenotype | GoF/LoF | Gating Properties | Reference |
|---|---|---|---|---|---|---|---|---|
| | | | | | | | Iω:none<br>ExTraFol:none | |
| 103 | Na$_v$1.2 | T1623N | Na$_v$1.2:T1623<br>Na$_v$1.2-2:T1623 | T1818N | DEE11*<br>EOEE | Mixed | Imax:↑<br>τrec:none<br>Ip:none<br>V1/2act:none<br>V1/2inact:→<br>Iω:none<br>ExTraFol:none | (Thompson et al., 2020) |
| 104 | Na$_v$1.2 | P1658S | Na$_v$1.2:P1658<br>Na$_v$1.2-2:P1658 | P1853S | | | Imax:↓<br>τrec:none<br>Ip:none<br>V1/2act:none<br>V1/2inact:none<br>Iω:none<br>ExTraFol:none | (Miao et al., 2020) |
| 105 | Na$_v$1.2 | A1773T | Na$_v$1.2:A1773<br>Na$_v$1.2-2:A1773 | A1968T | BFIS3*<br>DEE | | Imax:↓<br>τrec:↓<br>Ip:none<br>V1/2act:←<br>V1/2inact:←<br>Iω:none<br>ExTraFol:none | (Miao et al., 2020) |
| 106 | Na$_v$1.2 | E1803G | Na$_v$1.2:E1803<br>Na$_v$1.2-2:E1803 | E1998G | EOEE | GOF | Imax:↑<br>τrec:↓<br>Ip:↑<br>V1/2act:none<br>V1/2inact:→<br>Iω:none<br>ExTraFol:none | (Begemann et al., 2019) |
| 107 | Na$_v$1.2 | M1879T | Na$_v$1.2:M1879<br>Na$_v$1.2-2:M1879 | M2074T | EOEE | | Imax:none<br>τrec:none<br>Ip:none<br>V1/2act:none<br>V1/2inact:→<br>Iω:none<br>ExTraFol:none | (Adney et al., 2020) |
| 108 | Na$_v$1.2 | R1882G | Na$_v$1.2:R1882<br>Na$_v$1.2-2:R1882 | R2077G | EA9*<br>BFIS3*<br>NEIDEE*<br>EPI | | Imax:none<br>τrec:none<br>Ip:none<br>V1/2act:←<br>V1/2inact:none | (Schwarz et al., 2015) |

| | Protein | Mutation | Isoform | Mutation (MSA) | Phenotype | GoF/LoF | Gating Properties | Reference |
|---|---|---|---|---|---|---|---|---|
| | | | | | | | Iω:none<br>ExTraFol:none | |
| 109 | $Na_v1.2$ | R1882Q | $Na_v1.2$:R1882<br>$Na_v1.2$-2:R1882 | R2077Q | DEE11*<br>NEIDEE | GOF | Imax:↑<br>τrec:none<br>Ip:↑<br>V1/2act:←<br>V1/2inact:→<br>Iω:none<br>ExTraFol:none | (Schwarz et al., 2015)<br>(Mason et al., 2019)<br>(Berecki et al., 2018)<br>(Wolff et al., 2017)<br>(Thompson et al., 2020) |
| 110 | $Na_v1.3$ | L247P | $Na_v1.3$:L247<br>$Na_v1.3$-2:L247<br>$Na_v1.3$-3:L247<br>$Na_v1.3$-4:L247 | L259P | FFEVF4*<br>FE | LOF | Imax:↓<br>τrec:none<br>Ip:none<br>V1/2act:none<br>V1/2inact:none<br>Iω:none<br>ExTraFol:↓ | (Lamar et al., 2017) |
| 111 | $Na_v1.3$ | N302S | $Na_v1.3$:N302<br>$Na_v1.3$-2:N302<br>$Na_v1.3$-3:N302<br>$Na_v1.3$-4:N302 | N314S | FE | LOF | Imax:none<br>τrec:none<br>Ip:none<br>V1/2act:→<br>V1/2inact:→<br>Iω:none<br>ExTraFol:none | (Y.-J. J. Chen et al., 2015) |
| 112 | $Na_v1.3$ | R357Q | $Na_v1.3$:R357<br>$Na_v1.3$-2:R357<br>$Na_v1.3$-3:R357<br>$Na_v1.3$-4:R357 | R413Q | FFEVF4*<br>FE | GOF | Imax:↓<br>τrec:none<br>Ip:none<br>V1/2act:←<br>V1/2inact:←<br>Iω:none<br>ExTraFol:none | (Vanoye et al., 2014)<br>(Ademuwagun et al., 2021)<br>(Vanoye et al., 2014) |
| 113 | $Na_v1.3$ | L855P | $Na_v1.3$:L855<br>$Na_v1.3$-2:L806<br>$Na_v1.3$-3:L855<br>$Na_v1.3$-4:L806 | L945P | | | Imax:none<br>τrec:none<br>Ip:↑<br>V1/2act:←<br>V1/2inact:none<br>Iω:none<br>ExTraFol:none | (Zaman et al., 2020) |
| 114 | $Na_v1.3$ | I875T | $Na_v1.3$:I875<br>$Na_v1.3$-2:I826 | I965T | DEE62 | GOF | Imax:↑<br>τrec:none<br>Ip:↑<br>V1/2act:← | (Zaman et al., 2020)<br>(Zaman et al., 2018) |

| | Protein | Mutation | Isoform | Mutation (MSA) | Phenotype | GoF/LoF | Gating Properties | Reference |
|---|---|---|---|---|---|---|---|---|
| | | | Na$_v$1.3-3:I875<br>Na$_v$1.3-4:I826 | | | | V1/2inact:→<br>Iω:none<br>ExTraFol:none | |
| 115 | Na$_v$1.3 | E1111K | Na$_v$1.3:E1160<br>Na$_v$1.3-2:E1111<br>Na$_v$1.3-3:E1160<br>Na$_v$1.3-4:E1111 | E1160K | FE | GOF | Imax:none<br>τrec:none<br>Ip:↑<br>V1/2act:none<br>V1/2inact:none<br>Iω:none<br>ExTraFol:none | (Vanoye et al., 2014)<br>(Ademuwagun et al., 2021) |
| 116 | Na$_v$1.3 | P1333L | Na$_v$1.3:P1333<br>Na$_v$1.3-2:P1284<br>Na$_v$1.3-3:P1333<br>Na$_v$1.3-4:P1284 | P1523L | DEE62 | GOF | Imax:none<br>τrec:none<br>Ip:↑<br>V1/2act:←<br>V1/2inact:none<br>Iω:none<br>ExTraFol:none | (Zaman et al., 2020)<br>(Zaman et al., 2018) |
| 117 | Na$_v$1.3 | I1468R | Na$_v$1.3:I1468<br>Na$_v$1.3-2:I1419<br>Na$_v$1.3-3:I1468<br>Na$_v$1.3-4:I1419 | I1664R | | | Imax:↓<br>τrec:↓<br>Ip:↑<br>V1/2act:none<br>V1/2inact:→<br>Iω:none<br>ExTraFol:none | (Zaman et al., 2020) |
| 118 | Na$_v$1.3 | T1486I | Na$_v$1.3:T1486<br>Na$_v$1.3-2:T1437<br>Na$_v$1.3-3:T1486<br>Na$_v$1.3-4:T1437 | T1682I | PMG/ID | GOF | Imax:none<br>τrec:none<br>Ip:↑<br>V1/2act:none<br>V1/2inact:→<br>Iω:none<br>ExTraFol:none | (Zaman et al., 2020) |
| 119 | Na$_v$1.3 | R1621G | Na$_v$1.3:R1621<br>Na$_v$1.3-2:R1572<br>Na$_v$1.3-3:R1621<br>Na$_v$1.3-4:R1572 | R1821G | | | Imax:none<br>τrec:↓<br>Ip:↑<br>V1/2act:←<br>V1/2inact:none<br>Iω:none<br>ExTraFol:none | (Zaman et al., 2020) |
| 120 | Na$_v$1.3 | R1621Q | Na$_v$1.3:R1621<br>Na$_v$1.3-2:R1572<br>Na$_v$1.3-3:R1621<br>Na$_v$1.3-4:R1572 | R1821Q | PMG | GOF | Imax:none<br>τrec:none<br>Ip:↑<br>V1/2act:←<br>V1/2inact:none | (Zaman et al., 2020) |

| | Protein | Mutation | Isoform | Mutation (MSA) | Phenotype | GoF/LoF | Gating Properties | Reference |
|---|---|---|---|---|---|---|---|---|
| | | | | | | | Iω:none<br>ExTraFol:none | |
| 121 | Nav1.3 | R1642C | Nav1.3:R1642<br>Nav1.3-2:R1593<br>Nav1.3-3:R1642<br>Nav1.3-4:R1593 | R1842C | DEE62 | | Imax:↓<br>τrec:none<br>Ip:none<br>V1/2act:none<br>V1/2inact:none<br>Iω:none<br>ExTraFol:none | (Zaman et al., 2018) |
| 122 | Nav1.3 | F1646C | Nav1.3:F1646<br>Nav1.3-2:F1597<br>Nav1.3-3:F1646<br>Nav1.3-4:F1597 | F1846C | Familial polymicrogyria | | Imax:none<br>τrec:none<br>Ip:↑<br>V1/2act:none<br>V1/2inact:→<br>Iω:none<br>ExTraFol:none | (Zaman et al., 2020) |
| 123 | Nav1.3 | Y1669C | Nav1.3:Y1669<br>Nav1.3-2:Y1620<br>Nav1.3-3:Y1669<br>Nav1.3-4:Y1620 | Y1869C | ID | LOF | Imax:↓<br>τrec:none<br>Ip:none<br>V1/2act:none<br>V1/2inact:none<br>Iω:none<br>ExTraFol:none | (Zaman et al., 2020) |
| 124 | Nav1.3 | M1765I | Nav1.3:M1765<br>Nav1.3-2:M1716<br>Nav1.3-3:M1765<br>Nav1.3-4:M1716 | M1965I | EPI | GOF | Imax:↑<br>τrec:none<br>Ip:↑<br>V1/2act:←<br>V1/2inact:none<br>Iω:none<br>ExTraFol:none | (Zaman et al., 2020) |
| 125 | Nav1.3 | V1769A | Nav1.3:V1769<br>Nav1.3-2:V1720<br>Nav1.3-3:V1769<br>Nav1.3-4:V1720 | V1969A | DEE62 | GOF | Imax:↓<br>τrec:none<br>Ip:↑<br>V1/2act:none<br>V1/2inact:→<br>Iω:none<br>ExTraFol:none | (Zaman et al., 2018) |
| 126 | Nav1.3 | K1799Q | Nav1.3:K1799<br>Nav1.3-2:K1750<br>Nav1.3-3:K1799<br>Nav1.3-4:K1750 | K1999Q | DEE62 | | Imax:↓<br>τrec:none<br>Ip:none<br>V1/2act:none<br>V1/2inact:none | (Zaman et al., 2018) |

| | Protein | Mutation | Isoform | Mutation (MSA) | Phenotype | GoF/LoF | Gating Properties | Reference |
|---|---|---|---|---|---|---|---|---|
| | | | | | | | Iω:none<br>ExTraFol:none | |
| 127 | Na$_v$1.4 | R663H | - | 0 | HOKPP2 | | Imax:↓<br>τrec:none<br>Ip:↑<br>V1/2act:none<br>V1/2inact:none<br>Iω:↑<br>ExTraFol:none | (Struyk & Cannon, 2007) |
| 128 | Na$_v$1.4 | P72L | Na$_v$1.4:P72 | P76L | DM2 | GOF | Imax:none<br>τrec:↓<br>Ip:none<br>V1/2act:←<br>V1/2inact:none<br>Iω:none<br>ExTraFol:none | (Bugiardini et al., 2015) |
| 129 | Na$_v$1.4 | R104H | Na$_v$1.4:R104 | R108H | CMS16 | LOF | Imax:↓<br>τrec:none<br>Ip:none<br>V1/2act:→<br>V1/2inact:none<br>Iω:none<br>ExTraFol:none | (Zaharieva et al., 2016) |
| 130 | Na$_v$1.4 | I141V | Na$_v$1.4:I141 | I145V | PAM*<br>PMC*<br>SCM | GOF | Imax:↑<br>τrec:none<br>Ip:↑<br>V1/2act:←<br>V1/2inact:none<br>Iω:none<br>ExTraFol:none | (Petitprez, Tiab, et al., 2008) |
| 131 | Na$_v$1.4 | M203K | Na$_v$1.4:M203 | M211K | CMS16 | LOF | Imax:↓<br>τrec:none<br>Ip:none<br>V1/2act:→<br>V1/2inact:→<br>Iω:none<br>ExTraFol:none | (Zaharieva et al., 2016) |
| 132 | Na$_v$1.4 | A204E | Na$_v$1.4:A204 | A212E | HYPP | | Imax:none<br>τrec:none<br>Ip:none<br>V1/2act:←<br>V1/2inact:← | (Kokunai et al., 2018) |

| | Protein | Mutation | Isoform | Mutation (MSA) | Phenotype | GoF/LoF | Gating Properties | Reference |
|---|---|---|---|---|---|---|---|---|
| | | | | | | | Iω:none<br>ExTraFol:none | |
| 133 | $Na_v$1.4 | I215T | $Na_v$1.4:I215 | I224T | SCM | LOF | Imax:none<br>τrec:none<br>Ip:none<br>V1/2act:←<br>V1/2inact:none<br>Iω:none<br>ExTraFol:none | (Pagliarani et al., 2020) |
| 134 | $Na_v$1.4 | R222W | $Na_v$1.4:R222 | R231W | HOKPP2 | | Imax:↓<br>τrec:↓<br>Ip:none<br>V1/2act:→<br>V1/2inact:←<br>Iω:↑<br>ExTraFol:none | (Bayless-Edwards et al., 2018) |
| 135 | $Na_v$1.4 | R225W | $Na_v$1.4:R225 | R234W | PAM*<br>SCM | LOF | Imax:↓<br>τrec:none<br>Ip:none<br>V1/2act:→<br>V1/2inact:none<br>Iω:none<br>ExTraFol:none | (Zaharieva et al., 2016) |
| 136 | $Na_v$1.4 | G241V | $Na_v$1.4:G241 | G250V | MYOTONIA | LOF | Imax:none<br>τrec:none<br>Ip:none<br>V1/2act:←<br>V1/2inact:←<br>Iω:none<br>ExTraFol:none | (Pagliarani et al., 2020) |
| 137 | $Na_v$1.4 | L266V | $Na_v$1.4:L266 | L275V | PMC*<br>CAM | GOF | Imax:none<br>τrec:none<br>Ip:none<br>V1/2act:none<br>V1/2inact:→<br>Iω:none<br>ExTraFol:none | (F.-F. F. Wu et al., 2001) |
| 138 | $Na_v$1.4 | Q270K | $Na_v$1.4:Q270 | Q279K | PMC*<br>PAM | GOF | Imax:none<br>τrec:↑<br>Ip:none<br>V1/2act:→<br>V1/2inact:→ | (Carle et al., 2009) |

| | Protein | Mutation | Isoform | Mutation (MSA) | Phenotype | GoF/LoF | Gating Properties | Reference |
|---|---|---|---|---|---|---|---|---|
| | | | | | | | Iω:none ExTraFol:none | |
| 139 | $Na_v1.4$ | P382T | $Na_v1.4$:P382 | P415T | CMS16 | LOF | Imax:↓ τrec:none Ip:none V1/2act:→ V1/2inact:none Iω:none ExTraFol:none | (Zaharieva et al., 2016) |
| 140 | $Na_v1.4$ | N440K | $Na_v1.4$:N440 | N473K | HYPP* PMC | GOF | Imax:none τrec:none Ip:↑ V1/2act:none V1/2inact:→ Iω:none ExTraFol:none | (Lossin, Nam, et al., 2012) |
| 141 | $Na_v1.4$ | V445M | $Na_v1.4$:V445 | V478M | HYPP* PAM | GOF | Imax:none τrec:none Ip:↑ V1/2act:← V1/2inact:none Iω:none ExTraFol:none | (D. W. Wang et al., 1999) |
| 142 | $Na_v1.4$ | R669G | $Na_v1.4$:R669 | R941G | NKPP* NormoPP | GOF | Imax:↑ τrec:none Ip:none V1/2act:none V1/2inact:none Iω:↑ ExTraFol:none | (Sokolov et al., 2008)V |
| 143 | $Na_v1.4$ | R669H | $Na_v1.4$:R669 | R941H | HOKPP2 | LOF | Imax:none τrec:↓ Ip:none V1/2act:none V1/2inact:← Iω:none ExTraFol:none | (Kuzmenkin et al., 2002) |
| 144 | $Na_v1.4$ | R669Q | $Na_v1.4$:R669 | R941Q | NKPP* NormoPP | GOF | Imax:↑ τrec:none Ip:none V1/2act:none V1/2inact:none | (Sokolov et al., 2008) |

| | Protein | Mutation | Isoform | Mutation (MSA) | Phenotype | GoF/LoF | Gating Properties | Reference |
|---|---|---|---|---|---|---|---|---|
| | | | | | | | Iω:↑<br>ExTraFol:none | |
| 145 | $Na_v1.4$ | R669W | $Na_v1.4$:R669 | R941W | NKPP*<br>NormoPP | GOF | Imax:↑<br>τrec:none<br>Ip:none<br>V1/2act:none<br>V1/2inact:none<br>Iω:↑<br>ExTraFol:↓ | (Sokolov et al., 2008) |
| 146 | $Na_v1.4$ | R672G | $Na_v1.4$:R672 | R944G | HOKPP2 | LOF | Imax:none<br>τrec:↓<br>Ip:none<br>V1/2act:→<br>V1/2inact:←<br>Iω:none<br>ExTraFol:↓ | (Kuzmenkin et al., 2002)<br>(Jurkat-Rott et al., 2000) |
| 147 | $Na_v1.4$ | R672H | $Na_v1.4$:R672 | R944H | HOKPP2 | LOF | Imax:none<br>τrec:↓<br>Ip:none<br>V1/2act:→<br>V1/2inact:←<br>Iω:none<br>ExTraFol:↓ | (Jurkat-Rott et al., 2000)<br>(Kuzmenkin et al., 2002) |
| 148 | $Na_v1.4$ | R672S | $Na_v1.4$:R672 | R944S | HOKPP2 | LOF | Imax:none<br>τrec:↓<br>Ip:none<br>V1/2act:→<br>V1/2inact:←<br>Iω:none<br>ExTraFol:none | (Kuzmenkin et al., 2002) |
| 149 | $Na_v1.4$ | L689I | $Na_v1.4$:L689 | L961I | PMC*<br>HYPP | GOF | Imax:none<br>τrec:none<br>Ip:↑<br>V1/2act:←<br>V1/2inact:none<br>Iω:none<br>ExTraFol:none | (Bendahhou et al., 2002) |
| 150 | $Na_v1.4$ | I692M | $Na_v1.4$:I692 | I964M | HYPP | | Imax:↑<br>τrec:↓<br>Ip:none<br>V1/2act:none<br>V1/2inact:← | (Fan et al., 2017) |

| | Protein | Mutation | Isoform | Mutation (MSA) | Phenotype | GoF/LoF | Gating Properties | Reference |
|---|---|---|---|---|---|---|---|---|
| | | | | | | | Iω:none<br>ExTraFol:none | |
| 151 | $Na_v1.4$ | I693T | $Na_v1.4$:I693 | I965T | HYPP*<br>PMC | GOF | Imax:none<br>τrec:none<br>Ip:none<br>V1/2act:←<br>V1/2inact:none<br>Iω:none<br>ExTraFol:none | (Bendahhou et al., 2002) |
| 152 | $Na_v1.4$ | T704M | $Na_v1.4$:T704 | T976M | SOTOS1*<br>PMC*<br>HYPP | GOF | Imax:none<br>τrec:none<br>Ip:↑<br>V1/2act:←<br>V1/2inact:→<br>Iω:none<br>ExTraFol:none | (Cummins & Sigworth, 1996)<br>(Hayward et al., 1997)<br>(Cannon, 2018)<br>(Cannon & Strittmatter, 1993) |
| 153 | $Na_v1.4$ | L796V | $Na_v1.4$:L796 | L1077V | Myotonic Myopathy | GOF | Imax:↓<br>τrec:none<br>Ip:none<br>V1/2act:←<br>V1/2inact:none<br>Iω:none<br>ExTraFol:none | (Elia et al., 2020) |
| 154 | $Na_v1.4$ | A799S | $Na_v1.4$:A799 | A1080S | SCM | GOF | Imax:none<br>τrec:none<br>Ip:none<br>V1/2act:←<br>V1/2inact:→<br>Iω:none<br>ExTraFol:none | (Lion-François et al., 2010) |
| 155 | $Na_v1.4$ | S804F | $Na_v1.4$:S804 | S1085F | PMC*<br>PAM | GOF | Imax:none<br>τrec:none<br>Ip:↑<br>V1/2act:none<br>V1/2inact:→<br>Iω:none<br>ExTraFol:none | (Green et al., 1998) |
| 156 | $Na_v1.4$ | D1069N | $Na_v1.4$:D1069 | D1434N | HOKPP2<br>CMS16 | LOF | Imax:↓<br>τrec:none<br>Ip:none<br>V1/2act:→ | (Zaharieva et al., 2016) |

| | Protein | Mutation | Isoform | Mutation (MSA) | Phenotype | GoF/LoF | Gating Properties | Reference |
|---|---|---|---|---|---|---|---|---|
| | | | | | | | V1/2inact:→<br>Iω:none<br>ExTraFol:none | |
| 157 | $Na_v1.4$ | R1132Q | $Na_v1.4$:R1132 | R1497Q | HYPP*<br>HOKPP2 | LOF | Imax:none<br>τrec:none<br>Ip:none<br>V1/2act:→<br>V1/2inact:←<br>Iω:none<br>ExTraFol:none | (Carle et al., 2006) |
| 158 | $Na_v1.4$ | R1135C | $Na_v1.4$:R1135 | R1500C | HOKPP2 | LOF | Imax:none<br>τrec:↓<br>Ip:none<br>V1/2act:←<br>V1/2inact:←<br>Iω:↑<br>ExTraFol:none | (Groome et al., 2014) |
| 159 | $Na_v1.4$ | R1135H | $Na_v1.4$:R1135 | R1500H | HOKPP2 | LOF | Imax:none<br>τrec:↓<br>Ip:none<br>V1/2act:none<br>V1/2inact:←<br>Iω:↑<br>ExTraFol:none | (Groome et al., 2014) |
| 160 | $Na_v1.4$ | A1152D | $Na_v1.4$:A1152 | A1517D | PMC | GOF | Imax:none<br>τrec:none<br>Ip:none<br>V1/2act:none<br>V1/2inact:→<br>Iω:none<br>ExTraFol:none | (Bouhours et al., 2005) |
| 161 | $Na_v1.4$ | A1156T | $Na_v1.4$:A1156 | A1521T | PAM*<br>HYPP*<br>PMC | GOF | Imax:none<br>τrec:↑<br>Ip:none<br>V1/2act:none<br>V1/2inact:→<br>Iω:none<br>ExTraFol:none | (Palmio et al., 2017) |
| 162 | $Na_v1.4$ | P1158L | $Na_v1.4$:P1158 | P1523L | HYPP*<br>SCM | GOF | Imax:none<br>τrec:none<br>Ip:none<br>V1/2act:none<br>V1/2inact:→ | (Desaphy et al., 2016) |

| | Protein | Mutation | Isoform | Mutation (MSA) | Phenotype | GoF/LoF | Gating Properties | Reference |
|---|---|---|---|---|---|---|---|---|
| | | | | | | | Iω:none<br>ExTraFol:none | |
| 163 | $Na_v1.4$ | P1158S | $Na_v1.4$:P1158 | P1523S | HOKPP2 | LOF | Imax:none<br>τrec:none<br>Ip:↑<br>V1/2act:none<br>V1/2inact:←<br>Iω:none<br>ExTraFol:none | (Sugiura et al., 2003) |
| 164 | $Na_v1.4$ | I1160V | $Na_v1.4$:I1160 | I1525V | PAM | GOF | Imax:none<br>τrec:none<br>Ip:none<br>V1/2act:←<br>V1/2inact:none<br>Iω:none<br>ExTraFol:none | (Sugiura et al., 2003) |
| 165 | $Na_v1.4$ | C1209F | $Na_v1.4$:C1209 | C1576F | CMS16 | LOF | Imax:↓<br>τrec:none<br>Ip:none<br>V1/2act:→<br>V1/2inact:none<br>Iω:none<br>ExTraFol:none | (Zaharieva et al., 2016) |
| 166 | $Na_v1.4$ | G1292D | $Na_v1.4$:G1292 | G1661D | NDM | GOF | Imax:none<br>τrec:none<br>Ip:none<br>V1/2act:←<br>V1/2inact:→<br>Iω:none<br>ExTraFol:none | (Kokunai et al., 2012) |
| 167 | $Na_v1.4$ | V1293I | $Na_v1.4$:V1293 | V1662I | PMC | GOF | Imax:none<br>τrec:none<br>Ip:↑<br>V1/2act:←<br>V1/2inact:→<br>Iω:none<br>ExTraFol:none | (Bendahhou et al., 2002)<br>(Green et al., 1998)<br>(Farinato et al., 2019) |
| 168 | $Na_v1.4$ | N1297S | $Na_v1.4$:N1297 | N1666S | SCM | GOF | Imax:none<br>τrec:none<br>Ip:none<br>V1/2act:none<br>V1/2inact:→ | (Maggi et al., 2017) |

| | Protein | Mutation | Isoform | Mutation (MSA) | Phenotype | GoF/LoF | Gating Properties | Reference |
|---|---|---|---|---|---|---|---|---|
| | | | | | | | Iω:none<br>ExTraFol:none | |
| 169 | $Na_v1.4$ | F1298C | $Na_v1.4$:F1298 | F1667C | SCM | GOF | Imax:none<br>τrec:none<br>Ip:none<br>V1/2act:→<br>V1/2inact:→<br>Iω:none<br>ExTraFol:none | (Farinato et al., 2019) |
| 170 | $Na_v1.4$ | G1306A | $Na_v1.4$:G1306 | G1675A | PMC*<br>SCM<br>PAM | GOF | Imax:none<br>τrec:none<br>Ip:none<br>V1/2act:←<br>V1/2inact:→<br>Iω:none<br>ExTraFol:none | (Bendahhou et al., 2002)<br>(Hayward et al., 1996) |
| 171 | $Na_v1.4$ | G1306E | $Na_v1.4$:G1306 | G1675E | PMC<br>PAM<br>SCM | GOF | Imax:none<br>τrec:none<br>Ip:none<br>V1/2act:←<br>V1/2inact:→<br>Iω:none<br>ExTraFol:none | (Bendahhou et al., 2002)<br>(Hayward et al., 1996) |
| 172 | $Na_v1.4$ | G1306V | $Na_v1.4$:G1306 | G1675V | HYPP*<br>PMC*<br>SCM<br>PAM | GOF | Imax:none<br>τrec:none<br>Ip:none<br>V1/2act:←<br>V1/2inact:→<br>Iω:none<br>ExTraFol:none | (Bendahhou et al., 2002)<br>(Hayward et al., 1996) |
| 173 | $Na_v1.4$ | I1310N | $Na_v1.4$:I1310 | I1679N | PAM*<br>HYPP*<br>CAM | | Imax:none<br>τrec:none<br>Ip:none<br>V1/2act:→<br>V1/2inact:→<br>Iω:none<br>ExTraFol:none | (Farinato et al., 2019) |
| 174 | $Na_v1.4$ | T1313A | $Na_v1.4$:T1313 | T1682A | PC | | Imax:none<br>τrec:↑<br>Ip:none<br>V1/2act:none<br>V1/2inact:→ | (Bouhours et al., 2004) |

| | Protein | Mutation | Isoform | Mutation (MSA) | Phenotype | GoF/LoF | Gating Properties | Reference |
|---|---|---|---|---|---|---|---|---|
| | | | | | | | Iω:none<br>ExTraFol:none | |
| 175 | $Na_v1.4$ | T1313M | $Na_v1.4$:T1313 | T1682M | HYPP*<br>PMC | GOF | Imax:none<br>τrec:↑<br>Ip:↑<br>V1/2act:←<br>V1/2inact:→<br>Iω:none<br>ExTraFol:none | (Hayward et al., 1996) (Bouhours et al., 2004) |
| 176 | $Na_v1.4$ | M1360V | $Na_v1.4$:M1360 | M1729V | HYPP | GOF | Imax:↑<br>τrec:none<br>Ip:none<br>V1/2act:←<br>V1/2inact:←<br>Iω:none<br>ExTraFol:none | (Wagner et al., 1997) |
| 177 | $Na_v1.4$ | I1363T | $Na_v1.4$:I1363 | I1732T | PMC | GOF | Imax:↑<br>τrec:none<br>Ip:none<br>V1/2act:none<br>V1/2inact:→<br>Iω:none<br>ExTraFol:none | (S. Tang et al., 2019) |
| 178 | $Na_v1.4$ | N1366S | $Na_v1.4$:N1366 | N1735S | PC | | Imax:none<br>τrec:↑<br>Ip:none<br>V1/2act:←<br>V1/2inact:→<br>Iω:none<br>ExTraFol:none | (Ke et al., 2017) |
| 179 | $Na_v1.4$ | L1433R | $Na_v1.4$:L1433 | L1802R | HYPP*<br>PMC | GOF | Imax:none<br>τrec:none<br>Ip:none<br>V1/2act:←<br>V1/2inact:→<br>Iω:none<br>ExTraFol:none | (N. Yang et al., 1994) |
| 180 | $Na_v1.4$ | V1442E | $Na_v1.4$:V1442 | V1815E | CMS16 | LOF | Imax:none<br>τrec:none<br>Ip:none<br>V1/2act:none<br>V1/2inact:← | (Tsujino et al., 2003) |

| | Protein | Mutation | Isoform | Mutation (MSA) | Phenotype | GoF/LoF | Gating Properties | Reference |
|---|---|---|---|---|---|---|---|---|
| | | | | | | | Iω:none<br>ExTraFol:none | |
| 181 | $Na_v1.4$ | R1448C | $Na_v1.4$:R1448 | R1821C | PMC | GOF | Imax:none<br>τrec:none<br>Ip:↑<br>V1/2act:←<br>V1/2inact:→<br>Iω:none<br>ExTraFol:none | (N. Yang et al., 1994) |
| 182 | $Na_v1.4$ | R1448H | $Na_v1.4$:R1448 | R1821H | PMC | GOF | Imax:none<br>τrec:none<br>Ip:none<br>V1/2act:←<br>V1/2inact:→<br>Iω:none<br>ExTraFol:none | (Chahine et al., 1994) |
| 183 | $Na_v1.4$ | R1448P | $Na_v1.4$:R1448 | R1821P | PMC | GOF | Imax:none<br>τrec:↑<br>Ip:none<br>V1/2act:none<br>V1/2inact:←<br>Iω:none<br>ExTraFol:none | (Featherstone et al., 1998) |
| 184 | $Na_v1.4$ | R1448S | $Na_v1.4$:R1448 | R1821S | PMC | GOF | Imax:none<br>τrec:↑<br>Ip:none<br>V1/2act:none<br>V1/2inact:←<br>Iω:none<br>ExTraFol:none | (Bendahhou, Cummins, Kwiecinski, et al., 1999) |
| 185 | $Na_v1.4$ | R1451C | $Na_v1.4$:R1451 | R1824C | HOKPP2/ MYOTONIA | LOF | Imax:↓<br>τrec:↓<br>Ip:none<br>V1/2act:none<br>V1/2inact:←<br>Iω:none<br>ExTraFol:none | (Poulin et al., 2018) |
| 186 | $Na_v1.4$ | R1451L | $Na_v1.4$:R1451 | R1824L | SCM*<br>HOKPP2<br>MYOTONIA | LOF | Imax:↓<br>τrec:↑<br>Ip:none<br>V1/2act:none<br>V1/2inact:← | (Poulin et al., 2018)<br>(Luo et al., 2018) |

| | Protein | Mutation | Isoform | Mutation (MSA) | Phenotype | GoF/LoF | Gating Properties | Reference |
|---|---|---|---|---|---|---|---|---|
| | | | | | | | Iω:none<br>ExTraFol:none | |
| 187 | $Na_v1.4$ | R1454W | $Na_v1.4$:R1454 | R1827W | CMS16 | | Imax:none<br>τrec:↓<br>Ip:none<br>V1/2act:none<br>V1/2inact:←<br>Iω:none<br>ExTraFol:none | (Habbout et al., 2016) |
| 188 | $Na_v1.4$ | I1455T | $Na_v1.4$:I1455 | I1828T | PMC | LOF | Imax:↓<br>τrec:↑<br>Ip:none<br>V1/2act:→<br>V1/2inact:none<br>Iω:none<br>ExTraFol:↓ | (Bednarz et al., 2017) |
| 189 | $Na_v1.4$ | R1457H | $Na_v1.4$:R1457 | R1830H | CMS16 | LOF | Imax:none<br>τrec:↓<br>Ip:none<br>V1/2act:none<br>V1/2inact:←<br>Iω:none<br>ExTraFol:none | (Arnold et al., 2015) |
| 190 | $Na_v1.4$ | R1460Q | $Na_v1.4$:R1460 | R1833Q | congenital hypotonia | | Imax:↓<br>τrec:↑<br>Ip:none<br>V1/2act:←<br>V1/2inact:none<br>Iω:none<br>ExTraFol:none | (Elia et al., 2019) |
| 191 | $Na_v1.4$ | R1460W | $Na_v1.4$:R1460 | R1833W | HYPP* congenital hypotonia | | Imax:↓<br>τrec:↑<br>Ip:none<br>V1/2act:←<br>V1/2inact:none<br>Iω:none<br>ExTraFol:none | (Elia et al., 2019) |
| 192 | $Na_v1.4$ | R1463H | $Na_v1.4$:R1463 | R1836H | SCM | GOF | Imax:none<br>τrec:↑<br>Ip:none<br>V1/2act:none<br>V1/2inact:→ | (Brenes et al., 2021) |

| | Protein | Mutation | Isoform | Mutation (MSA) | Phenotype | GoF/LoF | Gating Properties | Reference |
|---|---|---|---|---|---|---|---|---|
| | | | | | | | Iω:none<br>ExTraFol:none | |
| 193 | $Na_v$1.4 | F1473S | $Na_v$1.4:F1473 | F1846S | PMC*<br>PAM | GOF | Imax:none<br>τrec:↑<br>Ip:↑<br>V1/2act:none<br>V1/2inact:→<br>Iω:none<br>ExTraFol:none | (Fleischhauer et al., 1998)<br>(Groome et al., 2008) |
| 194 | $Na_v$1.4 | I1495F | $Na_v$1.4:I1495 | I1868F | HYPP | GOF | Imax:↑<br>τrec:none<br>Ip:↑<br>V1/2act:←<br>V1/2inact:←<br>Iω:none<br>ExTraFol:none | (Bendahhou et al., 2002)<br>(Bendahhou, Cummins, Tawil, et al., 1999) |
| 195 | $Na_v$1.4 | V1589M | $Na_v$1.4:V1589 | V1962M | PMC*<br>PAM | GOF | Imax:none<br>τrec:none<br>Ip:none<br>V1/2act:none<br>V1/2inact:→<br>Iω:none<br>ExTraFol:none | (Bendahhou, Cummins, Tawil, et al., 1999) |
| 196 | $Na_v$1.4 | M1592V | $Na_v$1.4:M1592 | M1965V | NKPP*<br>HYPP | GOF | Imax:none<br>τrec:none<br>Ip:↑<br>V1/2act:←<br>V1/2inact:none<br>Iω:none<br>ExTraFol:none | (Hayward et al., 1997)<br>(Rojas et al., 1999) |
| 197 | $Na_v$1.4 | Q1633E | $Na_v$1.4:Q1633 | Q2006E | PAM | GOF | Imax:none<br>τrec:none<br>Ip:none<br>V1/2act:none<br>V1/2inact:→<br>Iω:none<br>ExTraFol:none | (Kubota et al., 2009) |
| 198 | $Na_v$1.4 | F1705I | $Na_v$1.4:F1705 | F2078I | HYPP*<br>PMC*<br>CAM | | Imax:none<br>τrec:none<br>Ip:none<br>V1/2act:none<br>V1/2inact:→ | (Biswas et al., 2013)<br>(Groome et al., 2008) |

| | Protein | Mutation | Isoform | Mutation (MSA) | Phenotype | GoF/LoF | Gating Properties | Reference |
|---|---|---|---|---|---|---|---|---|
| | | | | | | | Iω:none<br>ExTraFol:none | |
| 199 | $Na_v1.4$ | H1782Q | $Na_v1.4$:H1782 | H2157Q | CMS16 | GOF | Imax:↑<br>τrec:none<br>Ip:none<br>V1/2act:none<br>V1/2inact:none<br>Iω:none<br>ExTraFol:none | (Zaharieva et al., 2016) |
| 200 | $Na_v1.5$ | G145R | - | 0 | BRGDA1 | LOF | Imax:↓<br>τrec:none<br>Ip:↑<br>V1/2act:none<br>V1/2inact:none<br>Iω:none<br>ExTraFol:none | (Bersell et al., 2023) |
| 201 | $Na_v1.5$ | G9V | $Na_v1.5$:G9<br>$Na_v1.5$-2:G9<br>$Na_v1.5$-3:G9<br>$Na_v1.5$-4:G9<br>$Na_v1.5$-5:G9<br>$Na_v1.5$-6:G9 | G13V | LQT3*<br>RWS* | | Imax:none<br>τrec:none<br>Ip:none<br>V1/2act:none<br>V1/2inact:←<br>Iω:none<br>ExTraFol:none | (Glazer et al., 2020) |
| 202 | $Na_v1.5$ | R27H | $Na_v1.5$:R27<br>$Na_v1.5$-2:R27<br>$Na_v1.5$-3:R27<br>$Na_v1.5$-4:R27<br>$Na_v1.5$-5:R27<br>$Na_v1.5$-6:R27 | R31H | LQT3*<br>BRGDA1 | LOF | Imax:↓<br>τrec:↓<br>Ip:none<br>V1/2act:→<br>V1/2inact:none<br>Iω:none<br>ExTraFol:none | (Gütter et al., 2013) |
| 203 | $Na_v1.5$ | R43Q | $Na_v1.5$:R43<br>$Na_v1.5$-2:R43<br>$Na_v1.5$-3:R43<br>$Na_v1.5$-4:R43<br>$Na_v1.5$-5:R43<br>$Na_v1.5$-6:R43 | R47Q | LQT3 | GOF | Imax:none<br>τrec:none<br>Ip:none<br>V1/2act:←<br>V1/2inact:none<br>Iω:none<br>ExTraFol:none | (Lin et al., 2008) |
| 204 | $Na_v1.5$ | D84N | $Na_v1.5$:D84<br>$Na_v1.5$-2:D84<br>$Na_v1.5$-3:D84<br>$Na_v1.5$-4:D84 | D88N | BRGDA1 | LOF | Imax:↓<br>τrec:none<br>Ip:none<br>V1/2act:none<br>V1/2inact:none | (Glazer et al., 2020) |

| | Protein | Mutation | Isoform | Mutation (MSA) | Phenotype | GoF/LoF | Gating Properties | Reference |
|---|---|---|---|---|---|---|---|---|
| | | | Na$_v$1.5-5:D84<br>Na$_v$1.5-6:D84 | | | | Iω:none<br>ExTraFol:none | |
| 205 | Na$_v$1.5 | Y87C | Na$_v$1.5:Y87<br>Na$_v$1.5-2:Y87<br>Na$_v$1.5-3:Y87<br>Na$_v$1.5-4:Y87<br>Na$_v$1.5-5:Y87<br>Na$_v$1.5-6:Y87 | Y91C | BRGDA1 | LOF | Imax:↓<br>τrec:none<br>Ip:none<br>V1/2act:→<br>V1/2inact:none<br>Iω:none<br>ExTraFol:none | (Z. Wang et al., 2020) |
| 206 | Na$_v$1.5 | R104Q | Na$_v$1.5:R104<br>Na$_v$1.5-2:R104<br>Na$_v$1.5-3:R104<br>Na$_v$1.5-4:R104<br>Na$_v$1.5-5:R104<br>Na$_v$1.5-6:R104 | R108Q | BRGDA1 | LOF | Imax:↓<br>τrec:none<br>Ip:none<br>V1/2act:none<br>V1/2inact:←<br>Iω:none<br>ExTraFol:none | (Gütter et al., 2013) |
| 207 | Na$_v$1.5 | R104W | Na$_v$1.5:R104<br>Na$_v$1.5-2:R104<br>Na$_v$1.5-3:R104<br>Na$_v$1.5-4:R104<br>Na$_v$1.5-5:R104<br>Na$_v$1.5-6:R104 | R108W | BRGDA1 | LOF | Imax:↓<br>τrec:none<br>Ip:none<br>V1/2act:→<br>V1/2inact:none<br>Iω:none<br>ExTraFol:none | (Doisne et al., 2021) |
| 208 | Na$_v$1.5 | S106G | Na$_v$1.5:S106<br>Na$_v$1.5-2:S106<br>Na$_v$1.5-3:S106<br>Na$_v$1.5-4:S106<br>Na$_v$1.5-5:S106<br>Na$_v$1.5-6:S106 | S110G | LQT3 | GOF | Imax:↑<br>τrec:none<br>Ip:none<br>V1/2act:←<br>V1/2inact:→<br>Iω:none<br>ExTraFol:none | (Scheiper-Welling et al., 2020) |
| 209 | Na$_v$1.5 | V125L | Na$_v$1.5:V125<br>Na$_v$1.5-2:V125<br>Na$_v$1.5-3:V125<br>Na$_v$1.5-4:V125<br>Na$_v$1.5-5:V125<br>Na$_v$1.5-6:V125 | V129L | LQT3 | GOF | Imax:none<br>τrec:↑<br>Ip:none<br>V1/2act:none<br>V1/2inact:→<br>Iω:none<br>ExTraFol:none | (Gütter et al., 2013) |
| 210 | Na$_v$1.5 | K126E | Na$_v$1.5:K126<br>Na$_v$1.5-2:K126<br>Na$_v$1.5-3:K126<br>Na$_v$1.5-4:K126 | K130E | BRGDA1 | LOF | Imax:↓<br>τrec:none<br>Ip:none<br>V1/2act:→<br>V1/2inact:→ | (Gütter et al., 2013) |

| | Protein | Mutation | Isoform | Mutation (MSA) | Phenotype | GoF/LoF | Gating Properties | Reference |
|---|---|---|---|---|---|---|---|---|
| | | | $Na_v$1.5-5:K126<br>$Na_v$1.5-6:K126 | | | | Iω:none<br>ExTraFol:none | |
| 211 | $Na_v$1.5 | L136P | $Na_v$1.5:L136<br>$Na_v$1.5-2:L136<br>$Na_v$1.5-3:L136<br>$Na_v$1.5-4:L136<br>$Na_v$1.5-5:L136<br>$Na_v$1.5-6:L136 | L140P | BRGDA1 | LOF | Imax:↓<br>τrec:none<br>Ip:none<br>V1/2act:none<br>V1/2inact:none<br>Iω:none<br>ExTraFol:none | (Glazer et al., 2020) |
| 212 | $Na_v$1.5 | I141V | $Na_v$1.5:I141<br>$Na_v$1.5-2:I141<br>$Na_v$1.5-3:I141<br>$Na_v$1.5-4:I141<br>$Na_v$1.5-5:I141<br>$Na_v$1.5-6:I141 | I145V | Hyperexcitability | GOF | Imax:none<br>τrec:none<br>Ip:none<br>V1/2act:←<br>V1/2inact:none<br>Iω:none<br>ExTraFol:none | (Amarouch et al., 2014) |
| 213 | $Na_v$1.5 | E161K | $Na_v$1.5:E161<br>$Na_v$1.5-2:E161<br>$Na_v$1.5-3:E161<br>$Na_v$1.5-4:E161<br>$Na_v$1.5-5:E161<br>$Na_v$1.5-6:E161 | E169K | SSS1<br>PFHB1A<br>BRGDA1 | LOF | Imax:↓<br>τrec:none<br>Ip:none<br>V1/2act:→<br>V1/2inact:←<br>Iω:none<br>ExTraFol:none | (Gui et al., 2010)<br>(Smits, Koopmann, et al., 2005)<br>(Meregalli et al., 2009) |
| 214 | $Na_v$1.5 | Y168F | $Na_v$1.5:Y168<br>$Na_v$1.5-2:Y168<br>$Na_v$1.5-3:Y168<br>$Na_v$1.5-4:Y168<br>$Na_v$1.5-5:Y168<br>$Na_v$1.5-6:Y168 | Y176F | Hyperexcitability | LOF | Imax:none<br>τrec:none<br>Ip:none<br>V1/2act:→<br>V1/2inact:none<br>Iω:none<br>ExTraFol:none | (Amarouch et al., 2014) |
| 215 | $Na_v$1.5 | K175N | $Na_v$1.5:K175<br>$Na_v$1.5-2:K175<br>$Na_v$1.5-3:K175<br>$Na_v$1.5-4:K175<br>$Na_v$1.5-5:K175<br>$Na_v$1.5-6:K175 | K183N | BRGDA1 | | Imax:none<br>τrec:none<br>Ip:none<br>V1/2act:none<br>V1/2inact:←<br>Iω:none<br>ExTraFol:none | (Glazer et al., 2020) |
| 216 | $Na_v$1.5 | A204E | $Na_v$1.5:A204<br>$Na_v$1.5-2:A204<br>$Na_v$1.5-3:A204<br>$Na_v$1.5-4:A204 | A212E | multifocal ectopic Purkinje-related premature | GOF | Imax:none<br>τrec:none<br>Ip:none<br>V1/2act:←<br>V1/2inact:none | (Doisne et al., 2020a) |

| | Protein | Mutation | Isoform | Mutation (MSA) | Phenotype | GoF/LoF | Gating Properties | Reference |
|---|---|---|---|---|---|---|---|---|
| | | | $Na_v1.5$-5:A204<br>$Na_v1.5$-6:A204 | | contractions | | Iω:none<br>ExTraFol:none | |
| 217 | $Na_v1.5$ | L212P | $Na_v1.5$:L212<br>$Na_v1.5$-2:L212<br>$Na_v1.5$-3:L212<br>$Na_v1.5$-4:L212<br>$Na_v1.5$-5:L212<br>$Na_v1.5$-6:L212 | L220P | LQT3*<br>RWS*<br>CCD | LOF | Imax:none<br>τrec:none<br>Ip:none<br>V1/2act:←<br>V1/2inact:←<br>Iω:none<br>ExTraFol:none | (Makita et al., 2005) |
| 218 | $Na_v1.5$ | L212P | $Na_v1.5$:L212<br>$Na_v1.5$-2:L212<br>$Na_v1.5$-3:L212<br>$Na_v1.5$-4:L212<br>$Na_v1.5$-5:L212<br>$Na_v1.5$-6:L212 | L220P | LQT3*<br>RWS*<br>CCD*<br>SSS1 | LOF | Imax:none<br>τrec:↓<br>Ip:none<br>V1/2act:←<br>V1/2inact:→<br>Iω:none<br>ExTraFol:none | (Gui et al., 2010) |
| 219 | $Na_v1.5$ | G213D | $Na_v1.5$:G213<br>$Na_v1.5$-2:G213<br>$Na_v1.5$-3:G213<br>$Na_v1.5$-4:G213<br>$Na_v1.5$-5:G213<br>$Na_v1.5$-6:G213 | G221D | DCM | GOF | Imax:↓<br>τrec:none<br>Ip:none<br>V1/2act:←<br>V1/2inact:→<br>Iω:none<br>ExTraFol:none | (Calloe et al., 2018) |
| 220 | $Na_v1.5$ | S216L | $Na_v1.5$:S216<br>$Na_v1.5$-2:S216<br>$Na_v1.5$-3:S216<br>$Na_v1.5$-4:S216<br>$Na_v1.5$-5:S216<br>$Na_v1.5$-6:S216 | S225L | BRGDA1*<br>LQT3 | GOF | Imax:none<br>τrec:none<br>Ip:↑<br>V1/2act:none<br>V1/2inact:→<br>Iω:none<br>ExTraFol:none | (D. W. Wang et al., 2007) |
| 221 | $Na_v1.5$ | R219Q | $Na_v1.5$:R219<br>$Na_v1.5$-2:R219<br>$Na_v1.5$-3:R219<br>$Na_v1.5$-4:R219<br>$Na_v1.5$-5:R219<br>$Na_v1.5$-6:R219 | R228Q | | | Imax:none<br>τrec:none<br>Ip:none<br>V1/2act:→<br>V1/2inact:none<br>Iω:none<br>ExTraFol:none | (L. Q. Chen et al., 1996) |
| 222 | $Na_v1.5$ | R219H | $Na_v1.5$:R219<br>$Na_v1.5$-2:R219<br>$Na_v1.5$-3:R219<br>$Na_v1.5$-4:R219 | R228H | DCM | GOF | Imax:none<br>τrec:none<br>Ip:none<br>V1/2act:none<br>V1/2inact:none | (Gosselin-Badaroudine et al., 2012) |

| | Protein | Mutation | Isoform | Mutation (MSA) | Phenotype | GoF/LoF | Gating Properties | Reference |
|---|---|---|---|---|---|---|---|---|
| | | | $Na_v$1.5-5:R219<br>$Na_v$1.5-6:R219 | | | | Iω:↑<br>ExTraFol:none | |
| 223 | $Na_v$1.5 | R219H | $Na_v$1.5:R219<br>$Na_v$1.5-2:R219<br>$Na_v$1.5-3:R219<br>$Na_v$1.5-4:R219<br>$Na_v$1.5-5:R219<br>$Na_v$1.5-6:R219 | R228H | | | Imax:none<br>τrec:none<br>Ip:none<br>V1/2act:none<br>V1/2inact:←<br>Iω:none<br>ExTraFol:none | (L. Q. Chen et al., 1996) |
| 224 | $Na_v$1.5 | T220I | $Na_v$1.5:T220<br>$Na_v$1.5-2:T220<br>$Na_v$1.5-3:T220<br>$Na_v$1.5-4:T220<br>$Na_v$1.5-5:T220<br>$Na_v$1.5-6:T220 | T229I | BRGDA1*<br>SSS1 | LOF | Imax:↓<br>τrec:↓<br>Ip:none<br>V1/2act:none<br>V1/2inact:←<br>Iω:none<br>ExTraFol:none | (Gui et al., 2010)<br>(Glazer et al., 2020) |
| 225 | $Na_v$1.5 | R222Q | $Na_v$1.5:R222<br>$Na_v$1.5-2:R222<br>$Na_v$1.5-3:R222<br>$Na_v$1.5-4:R222<br>$Na_v$1.5-5:R222<br>$Na_v$1.5-6:R222 | R231Q | CMD1E*<br>RWS*<br>DCM<br>LQT3<br>BRGDA1 | MIX | Imax:↑<br>τrec:none<br>Ip:none<br>V1/2act:←<br>V1/2inact:←<br>Iω:↑<br>ExTraFol:none | (Mann et al., 2012)<br>(Laurent et al., 2012)<br>(Moreau, Gosselin-Badaroudine, Delemotte, et al., 2015)<br>(Daniel et al., 2019)<br>(Nair et al., 2012) |
| 226 | $Na_v$1.5 | V223L | $Na_v$1.5:V223<br>$Na_v$1.5-2:V223<br>$Na_v$1.5-3:V223<br>$Na_v$1.5-4:V223<br>$Na_v$1.5-5:V223<br>$Na_v$1.5-6:V223 | V232L | BRGDA1 | LOF | Imax:↓<br>τrec:none<br>Ip:none<br>V1/2act:none<br>V1/2inact:none<br>Iω:none<br>ExTraFol:none | (Glazer et al., 2020) |
| 227 | $Na_v$1.5 | R225P | $Na_v$1.5:R225<br>$Na_v$1.5-2:R225<br>$Na_v$1.5-3:R225<br>$Na_v$1.5-4:R225<br>$Na_v$1.5-5:R225<br>$Na_v$1.5-6:R225 | R234P | arrhythmia-associated cardiomyopathy | GOF | Imax:↑<br>τrec:↓<br>Ip:↑<br>V1/2act:←<br>V1/2inact:none<br>Iω:↑<br>ExTraFol:none | (Beckermann et al., 2014)<br>(Moreau, Gosselin-Badaroudine, Boutjdir, et al., 2015) |

| | Protein | Mutation | Isoform | Mutation (MSA) | Phenotype | GoF/LoF | Gating Properties | Reference |
|---|---|---|---|---|---|---|---|---|
| 228 | Na$_v$1.5 | R225Q | Na$_v$1.5:R225<br>Na$_v$1.5-2:R225<br>Na$_v$1.5-3:R225<br>Na$_v$1.5-4:R225<br>Na$_v$1.5-5:R225<br>Na$_v$1.5-6:R225 | R234Q | LQT3* | | Imax:none<br>τrec:none<br>Ip:none<br>V1/2act:→<br>V1/2inact:←<br>Iω:none<br>ExTraFol:none | (L. Q. Chen et al., 1996) |
| 229 | Na$_v$1.5 | R225W | Na$_v$1.5:R225<br>Na$_v$1.5-2:R225<br>Na$_v$1.5-3:R225<br>Na$_v$1.5-4:R225<br>Na$_v$1.5-5:R225<br>Na$_v$1.5-6:R225 | R234W | PFHB1A*<br>LQT3*<br>BRGDA1*<br>CCD*<br>IBS | MIX | Imax:↓<br>τrec:↑<br>Ip:none<br>V1/2act:→<br>V1/2inact:→<br>Iω:↑<br>ExTraFol:none | (Bezzina et al., 2003)<br>(Moreau, Gosselin-Badaroudine, Delemotte, et al., 2015)<br>(Strege et al., 2018) |
| 230 | Na$_v$1.5 | A226D | Na$_v$1.5:A226<br>Na$_v$1.5-2:A226<br>Na$_v$1.5-3:A226<br>Na$_v$1.5-4:A226<br>Na$_v$1.5-5:A226<br>Na$_v$1.5-6:A226 | A235D | IVF | LOF | Imax:↓<br>τrec:none<br>Ip:none<br>V1/2act:none<br>V1/2inact:none<br>Iω:none<br>ExTraFol:↓ | (Watanabe, Nogami, Ohkubo, Kawata, Hayashi, Ishikawa, Makiyama, Nagao, Yagihara, Takehara, Shimizu, & Makita, 2011) |
| 231 | Na$_v$1.5 | A226V | Na$_v$1.5:A226<br>Na$_v$1.5-2:A226<br>Na$_v$1.5-3:A226<br>Na$_v$1.5-4:A226<br>Na$_v$1.5-5:A226<br>Na$_v$1.5-6:A226 | A235V | BRGDA1 | LOF | Imax:↓<br>τrec:none<br>Ip:none<br>V1/2act:none<br>V1/2inact:none<br>Iω:none<br>ExTraFol:none | (B. Y. B. Y. Tan et al., 2016) |
| 232 | Na$_v$1.5 | Q270K | Na$_v$1.5:Q270<br>Na$_v$1.5-2:Q270<br>Na$_v$1.5-3:Q270<br>Na$_v$1.5-4:Q270<br>Na$_v$1.5-5:Q270<br>Na$_v$1.5-6:Q270 | Q279K | LQT3<br>BRGDA1 | GOF | Imax:↓<br>τrec:none<br>Ip:↑<br>V1/2act:→<br>V1/2inact:→<br>Iω:none<br>ExTraFol:none | (Calloe et al., 2011) |
| 233 | Na$_v$1.5 | R282C | Na$_v$1.5:R282<br>Na$_v$1.5-2:R282<br>Na$_v$1.5-3:R282<br>Na$_v$1.5-4:R282 | R291C | BRGDA1 | LOF | Imax:↓<br>τrec:none<br>Ip:none<br>V1/2act:none<br>V1/2inact:none | (Glazer et al., 2020) |

| | Protein | Mutation | Isoform | Mutation (MSA) | Phenotype | GoF/LoF | Gating Properties | Reference |
|---|---|---|---|---|---|---|---|---|
| | | | $Na_v$1.5-5:R282<br>$Na_v$1.5-6:R282 | | | | Iω:none<br>ExTraFol:none | |
| 234 | $Na_v$1.5 | R282H | $Na_v$1.5:R282<br>$Na_v$1.5-2:R282<br>$Na_v$1.5-3:R282<br>$Na_v$1.5-4:R282<br>$Na_v$1.5-5:R282<br>$Na_v$1.5-6:R282 | R291H | BRGDA1 | LOF | Imax:↓<br>τrec:none<br>Ip:none<br>V1/2act:→<br>V1/2inact:→<br>Iω:none<br>ExTraFol:none | (Itoh, Shimizu, Mabuchi, et al., 2005) |
| 235 | $Na_v$1.5 | G298S | $Na_v$1.5:G298<br>$Na_v$1.5-2:G298<br>$Na_v$1.5-3:G298<br>$Na_v$1.5-4:G298<br>$Na_v$1.5-5:G298<br>$Na_v$1.5-6:G298 | G361S | PFHB1A*<br>CCD | LOF | Imax:↓<br>τrec:none<br>Ip:none<br>V1/2act:→<br>V1/2inact:→<br>Iω:none<br>ExTraFol:none | (D. W. Wang et al., 2002) |
| 236 | $Na_v$1.5 | L325R | $Na_v$1.5:L325<br>$Na_v$1.5-2:L325<br>$Na_v$1.5-3:L325<br>$Na_v$1.5-4:L325<br>$Na_v$1.5-5:L325<br>$Na_v$1.5-6:L325 | L392R | BRGDA1 | LOF | Imax:↓<br>τrec:none<br>Ip:none<br>V1/2act:→<br>V1/2inact:none<br>Iω:none<br>ExTraFol:none | (Keller et al., 2005) |
| 237 | $Na_v$1.5 | C335R | $Na_v$1.5:C335<br>$Na_v$1.5-2:C335<br>$Na_v$1.5-3:C335<br>$Na_v$1.5-4:C335<br>$Na_v$1.5-5:C335<br>$Na_v$1.5-6:C335 | C402R | DCM | LOF | Imax:↓<br>τrec:none<br>Ip:none<br>V1/2act:none<br>V1/2inact:none<br>Iω:none<br>ExTraFol:none | (Sedaghat-hamedani et al., 2021) |
| 238 | $Na_v$1.5 | P336L | $Na_v$1.5:P336<br>$Na_v$1.5-2:P336<br>$Na_v$1.5-3:P336<br>$Na_v$1.5-4:P336<br>$Na_v$1.5-5:P336<br>$Na_v$1.5-6:P336 | P403L | BRGDA1 | LOF | Imax:↓<br>τrec:none<br>Ip:none<br>V1/2act:→<br>V1/2inact:←<br>Iω:none<br>ExTraFol:none | (Cordeiro et al., 2006) |
| 239 | $Na_v$1.5 | R340Q | $Na_v$1.5:R340<br>$Na_v$1.5-2:R340<br>$Na_v$1.5-3:R340<br>$Na_v$1.5-4:R340 | R407Q | LQT3 | GOF | Imax:↓<br>τrec:↑<br>Ip:none<br>V1/2act:←<br>V1/2inact:← | (Olesen et al., 2012) |

| | Protein | Mutation | Isoform | Mutation (MSA) | Phenotype | GoF/LoF | Gating Properties | Reference |
|---|---|---|---|---|---|---|---|---|
| | | | Na$_v$1.5-5:R340<br>Na$_v$1.5-6:R340 | | | | Iω:none<br>ExTraFol:none | |
| 240 | Na$_v$1.5 | K343Q | Na$_v$1.5:K343<br>Na$_v$1.5-2:K343<br>Na$_v$1.5-3:K343<br>Na$_v$1.5-4:K343<br>Na$_v$1.5-5:K343<br>Na$_v$1.5-6:K343 | K410Q | | GOF | Imax:none<br>τrec:none<br>Ip:↑<br>V1/2act:none<br>V1/2inact:none<br>Iω:none<br>ExTraFol:none | (Holland et al., 2008) |
| 241 | Na$_v$1.5 | E346D | Na$_v$1.5:E346<br>Na$_v$1.5-2:E346<br>Na$_v$1.5-3:E346<br>Na$_v$1.5-4:E346<br>Na$_v$1.5-5:E346<br>Na$_v$1.5-6:E346 | E413D | BRGDA1 | | Imax:none<br>τrec:↓<br>Ip:none<br>V1/2act:none<br>V1/2inact:none<br>Iω:none<br>ExTraFol:none | (Glazer et al., 2020) |
| 242 | Na$_v$1.5 | D349N | Na$_v$1.5:D349<br>Na$_v$1.5-2:D349<br>Na$_v$1.5-3:D349<br>Na$_v$1.5-4:D349<br>Na$_v$1.5-5:D349<br>Na$_v$1.5-6:D349 | D416N | BRGDA1 | LOF | Imax:↓<br>τrec:none<br>Ip:none<br>V1/2act:none<br>V1/2inact:none<br>Iω:none<br>ExTraFol:none | (Glazer et al., 2020) |
| 243 | Na$_v$1.5 | G351V | Na$_v$1.5:G351<br>Na$_v$1.5-2:G351<br>Na$_v$1.5-3:G351<br>Na$_v$1.5-4:G351<br>Na$_v$1.5-5:G351<br>Na$_v$1.5-6:G351 | G418V | BRGDA1 | LOF | Imax:↓<br>τrec:none<br>Ip:none<br>V1/2act:→<br>V1/2inact:none<br>Iω:none<br>ExTraFol:none | (Vatta, Dumaine, Antzelevitch, et al., 2002) |
| 244 | Na$_v$1.5 | T353I | Na$_v$1.5:T353<br>Na$_v$1.5-2:T353<br>Na$_v$1.5-3:T353<br>Na$_v$1.5-4:T353<br>Na$_v$1.5-5:T353<br>Na$_v$1.5-6:T353 | T420I | BRGDA1 | LOF | Imax:↓<br>τrec:none<br>Ip:none<br>V1/2act:none<br>V1/2inact:←<br>Iω:none<br>ExTraFol:none | (Pfahnl et al., 2007) |
| 245 | Na$_v$1.5 | R367H | Na$_v$1.5:R367<br>Na$_v$1.5-2:R367<br>Na$_v$1.5-3:R367<br>Na$_v$1.5-4:R367 | R434H | BRGDA1 | LOF | Imax:↓<br>τrec:↑<br>Ip:none<br>V1/2act:→<br>V1/2inact:← | (Vatta, Dumaine, Varghese, et al., 2002) (Watanabe, Nogami, |

| | Protein | Mutation | Isoform | Mutation (MSA) | Phenotype | GoF/LoF | Gating Properties | Reference |
|---|---|---|---|---|---|---|---|---|
| | | | Na$_v$1.5-5:R367<br>Na$_v$1.5-6:R367 | | | | Iω:none<br>ExTraFol:none | Ohkubo, Kawata, Hayashi, Ishikawa, Makiyama, Nagao, Yagihara, Takehara, Shimizu, & Makita, 2011)<br>(Selga et al., 2018) |
| 246 | Na$_v$1.5 | R367S | Na$_v$1.5:R367<br>Na$_v$1.5-2:R367<br>Na$_v$1.5-3:R367<br>Na$_v$1.5-4:R367<br>Na$_v$1.5-5:R367<br>Na$_v$1.5-6:R367 | R434S | BRGDA1 | LOF | Imax:↓<br>τrec:none<br>Ip:none<br>V1/2act:none<br>V1/2inact:none<br>Iω:none<br>ExTraFol:none | (Suzuki et al., 2022)v |
| 247 | Na$_v$1.5 | M369K | Na$_v$1.5:M369<br>Na$_v$1.5-2:M369<br>Na$_v$1.5-3:M369<br>Na$_v$1.5-4:M369<br>Na$_v$1.5-5:M369<br>Na$_v$1.5-6:M369 | M436K | BRGDA1 | LOF | Imax:↓<br>τrec:none<br>Ip:none<br>V1/2act:none<br>V1/2inact:none<br>Iω:none<br>ExTraFol:none | (Glazer et al., 2020) |
| 248 | Na$_v$1.5 | R376H | Na$_v$1.5:R376<br>Na$_v$1.5-2:R376<br>Na$_v$1.5-3:R376<br>Na$_v$1.5-4:R376<br>Na$_v$1.5-5:R376<br>Na$_v$1.5-6:R376 | R443H | BRGDA1 | LOF | Imax:↓<br>τrec:none<br>Ip:none<br>V1/2act:none<br>V1/2inact:none<br>Iω:none<br>ExTraFol:none | (Rossenbacker et al., 2004) |
| 249 | Na$_v$1.5 | V396L | Na$_v$1.5:V396<br>Na$_v$1.5-2:V396<br>Na$_v$1.5-3:V396<br>Na$_v$1.5-4:V396<br>Na$_v$1.5-5:V396<br>Na$_v$1.5-6:V396 | V463L | BRGDA1 | LOF | Imax:↓<br>τrec:none<br>Ip:none<br>V1/2act:none<br>V1/2inact:none<br>Iω:none<br>ExTraFol:none | (Glazer et al., 2020) |
| 250 | Na$_v$1.5 | N406K | Na$_v$1.5:N406<br>Na$_v$1.5-2:N406<br>Na$_v$1.5-3:N406<br>Na$_v$1.5-4:N406 | N473K | RWS*<br>LQT3 | | Imax:↓<br>τrec:↓<br>Ip:↑<br>V1/2act:→<br>V1/2inact:none | (Kato et al., 2014)<br>(R. M. Hu et al., 2018) |

| | Protein | Mutation | Isoform | Mutation (MSA) | Phenotype | GoF/LoF | Gating Properties | Reference |
|---|---|---|---|---|---|---|---|---|
| | | | $Na_v$1.5-5:N406<br>$Na_v$1.5-6:N406 | | | | Iω:none<br>ExTraFol:↓ | |
| 251 | $Na_v$1.5 | N406S | $Na_v$1.5:N406<br>$Na_v$1.5-2:N406<br>$Na_v$1.5-3:N406<br>$Na_v$1.5-4:N406<br>$Na_v$1.5-5:N406<br>$Na_v$1.5-6:N406 | N473S | BRGDA1 | LOF | Imax:none<br>τrec:none<br>Ip:none<br>V1/2act:→<br>V1/2inact:→<br>Iω:none<br>ExTraFol:none | (Itoh, Shimizu, Takata, et al., 2005) |
| 252 | $Na_v$1.5 | V411M | $Na_v$1.5:V411<br>$Na_v$1.5-2:V411<br>$Na_v$1.5-3:V411<br>$Na_v$1.5-4:V411<br>$Na_v$1.5-5:V411<br>$Na_v$1.5-6:V411 | V478M | RWS*<br>LQT3 | GOF | Imax:none<br>τrec:none<br>Ip:↑<br>V1/2act:←<br>V1/2inact:←<br>Iω:none<br>ExTraFol:none | (Horne et al., 2011) |
| 253 | $Na_v$1.5 | E428K | $Na_v$1.5:E428<br>$Na_v$1.5-2:E428<br>$Na_v$1.5-3:E428<br>$Na_v$1.5-4:E428<br>$Na_v$1.5-5:E428<br>$Na_v$1.5-6:E428 | E495K | ATFB10*<br>AF | GOF | Imax:none<br>τrec:none<br>Ip:↑<br>V1/2act:←<br>V1/2inact:none<br>Iω:none<br>ExTraFol:none | (Hong et al., 2021) |
| 254 | $Na_v$1.5 | R433C | $Na_v$1.5:R433<br>$Na_v$1.5-2:R433<br>$Na_v$1.5-3:R433<br>$Na_v$1.5-4:R433<br>$Na_v$1.5-5:R433<br>$Na_v$1.5-6:R433 | R500C | IBS | LOF | Imax:↓<br>τrec:none<br>Ip:none<br>V1/2act:none<br>V1/2inact:←<br>Iω:none<br>ExTraFol:none | (Strege et al., 2018) |
| 255 | $Na_v$1.5 | K480N | $Na_v$1.5:K480<br>$Na_v$1.5-2:K480<br>$Na_v$1.5-3:K480<br>$Na_v$1.5-4:K480<br>$Na_v$1.5-5:K480<br>$Na_v$1.5-6:K480 | K570N | syncopes | GOF | Imax:none<br>τrec:↓<br>Ip:none<br>V1/2act:none<br>V1/2inact:→<br>Iω:none<br>ExTraFol:none | (Ortiz-Bonnin et al., 2016) |
| 256 | $Na_v$1.5 | T512I | $Na_v$1.5:T512<br>$Na_v$1.5-2:T512<br>$Na_v$1.5-3:T512<br>$Na_v$1.5-4:T512 | T625I | PFHB1A*<br>CCD | LOF | Imax:none<br>τrec:none<br>Ip:none<br>V1/2act:←<br>V1/2inact:← | (Viswanathan et al., 2003) |

| | Protein | Mutation | Isoform | Mutation (MSA) | Phenotype | GoF/LoF | Gating Properties | Reference |
|---|---|---|---|---|---|---|---|---|
| | | | Na$_v$1.5-5:T512<br>Na$_v$1.5-6:T512 | | | | Iω:none<br>ExTraFol:none | |
| 257 | Na$_v$1.5 | G514C | Na$_v$1.5:G514<br>Na$_v$1.5-2:G514<br>Na$_v$1.5-3:G514<br>Na$_v$1.5-4:G514<br>Na$_v$1.5-5:G514<br>Na$_v$1.5-6:G514 | G627C | PFHB1A*<br>BRGDA1*<br>CCD | LOF | Imax:none<br>τrec:none<br>Ip:none<br>V1/2act:→<br>V1/2inact:→<br>Iω:none<br>ExTraFol:none | (H. L. Tan et al., 2001) |
| 258 | Na$_v$1.5 | A551T | Na$_v$1.5:A551<br>Na$_v$1.5-2:A551<br>Na$_v$1.5-3:A551<br>Na$_v$1.5-4:A551<br>Na$_v$1.5-5:A551<br>Na$_v$1.5-6:A551 | A668T | BRGDA1 | | Imax:↓<br>τrec:none<br>Ip:none<br>V1/2act:none<br>V1/2inact:←<br>Iω:none<br>ExTraFol:none | (Chiang et al., 2009) |
| 259 | Na$_v$1.5 | L567Q | Na$_v$1.5:L567<br>Na$_v$1.5-2:L567<br>Na$_v$1.5-3:L567<br>Na$_v$1.5-4:L567<br>Na$_v$1.5-5:L567<br>Na$_v$1.5-6:L567 | L684Q | BRGDA1 | LOF | Imax:none<br>τrec:none<br>Ip:none<br>V1/2act:→<br>V1/2inact:←<br>Iω:none<br>ExTraFol:none | (Wan et al., 2001) |
| 260 | Na$_v$1.5 | R568H | Na$_v$1.5:R568<br>Na$_v$1.5-2:R568<br>Na$_v$1.5-3:R568<br>Na$_v$1.5-4:R568<br>Na$_v$1.5-5:R568<br>Na$_v$1.5-6:R568 | R687H | LQT3 | GOF | Imax:↑<br>τrec:↑<br>Ip:none<br>V1/2act:none<br>V1/2inact:→<br>Iω:none<br>ExTraFol:none | (Ortiz-Bonnin et al., 2016) |
| 261 | Na$_v$1.5 | A572D | Na$_v$1.5:A572<br>Na$_v$1.5-2:A572<br>Na$_v$1.5-3:A572<br>Na$_v$1.5-4:A572<br>Na$_v$1.5-5:A572<br>Na$_v$1.5-6:A572 | A696D | ATFB10*<br>LQT3 | GOF | Imax:none<br>τrec:none<br>Ip:none<br>V1/2act:none<br>V1/2inact:→<br>Iω:none<br>ExTraFol:none | (Tester et al., 2005)<br>(Ortiz-Bonnin et al., 2016) |
| 262 | Na$_v$1.5 | G615E | Na$_v$1.5:G615<br>Na$_v$1.5-2:G615<br>Na$_v$1.5-3:G615<br>Na$_v$1.5-4:G615 | G740E | BRGDA1*<br>LQT3 | GOF | Imax:↓<br>τrec:none<br>Ip:none<br>V1/2act:none<br>V1/2inact:→ | (P. Yang et al., 2002) |

| | Protein | Mutation | Isoform | Mutation (MSA) | Phenotype | GoF/LoF | Gating Properties | Reference |
|---|---|---|---|---|---|---|---|---|
| | | | Na$_v$1.5-5:G615<br>Na$_v$1.5-6:G615 | | | | Iω:none<br>ExTraFol:none | |
| 263 | Na$_v$1.5 | L618F | Na$_v$1.5:L618<br>Na$_v$1.5-2:L618<br>Na$_v$1.5-3:L618<br>Na$_v$1.5-4:L618<br>Na$_v$1.5-5:L618<br>Na$_v$1.5-6:L618 | L743F | LQT3 | GOF | Imax:↑<br>τrec:none<br>Ip:none<br>V1/2act:←<br>V1/2inact:→<br>Iω:none<br>ExTraFol:none | (P. Yang et al., 2002) |
| 264 | Na$_v$1.5 | L619F | Na$_v$1.5:L619<br>Na$_v$1.5-2:L619<br>Na$_v$1.5-3:L619<br>Na$_v$1.5-4:L619<br>Na$_v$1.5-5:L619<br>Na$_v$1.5-6:L619 | L744F | BRGDA1*<br>LQT3 | GOF | Imax:none<br>τrec:none<br>Ip:↑<br>V1/2act:none<br>V1/2inact:→<br>Iω:none<br>ExTraFol:none | (Wehrens et al., 2003) |
| 265 | Na$_v$1.5 | R680H | Na$_v$1.5:R680<br>Na$_v$1.5-2:R680<br>Na$_v$1.5-3:R680<br>Na$_v$1.5-4:R680<br>Na$_v$1.5-5:R680<br>Na$_v$1.5-6:R680 | R811H | LQT3 | GOF | Imax:none<br>τrec:none<br>Ip:↑<br>V1/2act:←<br>V1/2inact:none<br>Iω:none<br>ExTraFol:none | (Remme, 2013) |
| 266 | Na$_v$1.5 | H681P | Na$_v$1.5:H681<br>Na$_v$1.5-2:H681<br>Na$_v$1.5-3:H681<br>Na$_v$1.5-4:H681<br>Na$_v$1.5-5:H681<br>Na$_v$1.5-6:H681 | H812P | BRGDA1 | LOF | Imax:none<br>τrec:none<br>Ip:none<br>V1/2act:←<br>V1/2inact:none<br>Iω:none<br>ExTraFol:none | (Mok et al., 2003) |
| 267 | Na$_v$1.5 | C683R | Na$_v$1.5:C683<br>Na$_v$1.5-2:C683<br>Na$_v$1.5-3:C683<br>Na$_v$1.5-4:C683<br>Na$_v$1.5-5:C683<br>Na$_v$1.5-6:C683 | C814R | adrenaline-triggered ventricular arrhythmia | GOF | Imax:none<br>τrec:↓<br>Ip:none<br>V1/2act:←<br>V1/2inact:none<br>Iω:none<br>ExTraFol:none | (Steinberg et al., 2021) |
| 268 | Na$_v$1.5 | M734V | Na$_v$1.5:M734<br>Na$_v$1.5-2:M734<br>Na$_v$1.5-3:M734<br>Na$_v$1.5-4:M734 | M865V | BRGDA1 | LOF | Imax:none<br>τrec:↓<br>Ip:none<br>V1/2act:none<br>V1/2inact:← | (Glazer et al., 2020) |

| | Protein | Mutation | Isoform | Mutation (MSA) | Phenotype | GoF/LoF | Gating Properties | Reference |
|---|---|---|---|---|---|---|---|---|
| | | | $Na_v$1.5-5:M734<br>$Na_v$1.5-6:M734 | | | | Iω:none<br>ExTraFol:none | |
| 269 | $Na_v$1.5 | A735T | $Na_v$1.5:A735<br>$Na_v$1.5-2:A735<br>$Na_v$1.5-3:A735<br>$Na_v$1.5-4:A735<br>$Na_v$1.5-5:A735<br>$Na_v$1.5-6:A735 | A866T | BRGDA1 | LOF | Imax:none<br>τrec:↑<br>Ip:none<br>V1/2act:none<br>V1/2inact:none<br>Iω:none<br>ExTraFol:none | (Glazer et al., 2020) |
| 270 | $Na_v$1.5 | A735V | $Na_v$1.5:A735<br>$Na_v$1.5-2:A735<br>$Na_v$1.5-3:A735<br>$Na_v$1.5-4:A735<br>$Na_v$1.5-5:A735<br>$Na_v$1.5-6:A735 | A866V | SSS1*<br>BRGDA1 | LOF | Imax:none<br>τrec:↓<br>Ip:none<br>V1/2act:→<br>V1/2inact:none<br>Iω:none<br>ExTraFol:none | (de la Roche et al., 2019) |
| 271 | $Na_v$1.5 | Y739D | $Na_v$1.5:Y739<br>$Na_v$1.5-2:Y739<br>$Na_v$1.5-3:Y739<br>$Na_v$1.5-4:Y739<br>$Na_v$1.5-5:Y739<br>$Na_v$1.5-6:Y739 | Y870D | BRGDA1 | LOF | Imax:↓<br>τrec:none<br>Ip:none<br>V1/2act:→<br>V1/2inact:←<br>Iω:none<br>ExTraFol:none | (Zaytseva et al., 2022) |
| 272 | $Na_v$1.5 | E746K | $Na_v$1.5:E746<br>$Na_v$1.5-2:E746<br>$Na_v$1.5-3:E746<br>$Na_v$1.5-4:E746<br>$Na_v$1.5-5:E746<br>$Na_v$1.5-6:E746 | E877K | BRGDA1 | LOF | Imax:↓<br>τrec:none<br>Ip:none<br>V1/2act:none<br>V1/2inact:none<br>Iω:none<br>ExTraFol:none | (Glazer et al., 2020) |
| 273 | $Na_v$1.5 | G752R | $Na_v$1.5:G752<br>$Na_v$1.5-2:G752<br>$Na_v$1.5-3:G752<br>$Na_v$1.5-4:G752<br>$Na_v$1.5-5:G752<br>$Na_v$1.5-6:G752 | G883R | PFHB1A*<br>BRGDA1 | LOF | Imax:↓<br>τrec:none<br>Ip:none<br>V1/2act:→<br>V1/2inact:→<br>Iω:none<br>ExTraFol:none | (Potet et al., 2003)<br>(Glazer et al., 2020) |
| 274 | $Na_v$1.5 | P773S | $Na_v$1.5:P773<br>$Na_v$1.5-2:P773<br>$Na_v$1.5-3:P773<br>$Na_v$1.5-4:P773 | P904S | BRGDA1 | | Imax:none<br>τrec:↓<br>Ip:none<br>V1/2act:none<br>V1/2inact:← | (Glazer et al., 2020) |

| | Protein | Mutation | Isoform | Mutation (MSA) | Phenotype | GoF/LoF | Gating Properties | Reference |
|---|---|---|---|---|---|---|---|---|
| | | | $Na_v$1.5-5:P773<br>$Na_v$1.5-6:P773 | | | | Iω:none<br>ExTraFol:none | |
| 275 | $Na_v$1.5 | D785N | $Na_v$1.5:D785<br>$Na_v$1.5-2:D785<br>$Na_v$1.5-3:D785<br>$Na_v$1.5-4:D785<br>$Na_v$1.5-5:D785<br>$Na_v$1.5-6:D785 | D916N | BRGDA1 | LOF | Imax:↓<br>τrec:↓<br>Ip:none<br>V1/2act:→<br>V1/2inact:←<br>Iω:none<br>ExTraFol:none | (Glazer et al., 2020) |
| 276 | $Na_v$1.5 | R800L | $Na_v$1.5:R800<br>$Na_v$1.5-2:R800<br>$Na_v$1.5-3:R800<br>$Na_v$1.5-4:R800<br>$Na_v$1.5-5:R800<br>$Na_v$1.5-6:R800 | R931L | LQT3 | GOF | Imax:none<br>τrec:none<br>Ip:↑<br>V1/2act:none<br>V1/2inact:none<br>Iω:none<br>ExTraFol:none | (R.-M. Hu et al., 2013) |
| 277 | $Na_v$1.5 | R808C | $Na_v$1.5:R808<br>$Na_v$1.5-2:R808<br>$Na_v$1.5-3:R808<br>$Na_v$1.5-4:R808<br>$Na_v$1.5-5:R808<br>$Na_v$1.5-6:R808 | R941C | BRGDA1 | LOF | Imax:↓<br>τrec:↓<br>Ip:none<br>V1/2act:none<br>V1/2inact:←<br>Iω:none<br>ExTraFol:none | (R.-M. Hu et al., 2013) |
| 278 | $Na_v$1.5 | R808H | $Na_v$1.5:R808<br>$Na_v$1.5-2:R808<br>$Na_v$1.5-3:R808<br>$Na_v$1.5-4:R808<br>$Na_v$1.5-5:R808<br>$Na_v$1.5-6:R808 | R941H | | | Imax:none<br>τrec:none<br>Ip:none<br>V1/2act:→<br>V1/2inact:none<br>Iω:none<br>ExTraFol:none | (L. Q. Chen et al., 1996) |
| 279 | $Na_v$1.5 | R808Q | $Na_v$1.5:R808<br>$Na_v$1.5-2:R808<br>$Na_v$1.5-3:R808<br>$Na_v$1.5-4:R808<br>$Na_v$1.5-5:R808<br>$Na_v$1.5-6:R808 | R941Q | | | Imax:none<br>τrec:none<br>Ip:none<br>V1/2act:→<br>V1/2inact:→<br>Iω:none<br>ExTraFol:none | (L. Q. Chen et al., 1996) |
| 280 | $Na_v$1.5 | R811H | $Na_v$1.5:R811<br>$Na_v$1.5-2:R811<br>$Na_v$1.5-3:R811<br>$Na_v$1.5-4:R811 | R944H | BRGDA1 | LOF | Imax:↓<br>τrec:↓<br>Ip:none<br>V1/2act:none<br>V1/2inact:← | (Calloe et al., 2013) |

| | Protein | Mutation | Isoform | Mutation (MSA) | Phenotype | GoF/LoF | Gating Properties | Reference |
|---|---|---|---|---|---|---|---|---|
| | | | Nav1.5-5:R811<br>Nav1.5-6:R811 | | | | Iω:none<br>ExTraFol:none | |
| 281 | Nav1.5 | R814E | Nav1.5:R814<br>Nav1.5-2:R814<br>Nav1.5-3:R814<br>Nav1.5-4:R814<br>Nav1.5-5:R814<br>Nav1.5-6:R814 | R947E | | | Imax:none<br>τrec:none<br>Ip:none<br>V1/2act:→<br>V1/2inact:←<br>Iω:none<br>ExTraFol:none | (L. Q. Chen et al., 1996) |
| 282 | Nav1.5 | R814Q | Nav1.5:R814<br>Nav1.5-2:R814<br>Nav1.5-3:R814<br>Nav1.5-4:R814<br>Nav1.5-5:R814<br>Nav1.5-6:R814 | R947Q | BRGDA1 | GOF | Imax:none<br>τrec:none<br>Ip:↑<br>V1/2act:none<br>V1/2inact:←<br>Iω:none<br>ExTraFol:none | (Glazer et al., 2020) |
| 283 | Nav1.5 | R814W | Nav1.5:R814<br>Nav1.5-2:R814<br>Nav1.5-3:R814<br>Nav1.5-4:R814<br>Nav1.5-5:R814<br>Nav1.5-6:R814 | R947W | CMD1E*<br>DCM | GOF | Imax:↓<br>τrec:↓<br>Ip:none<br>V1/2act:←<br>V1/2inact:←<br>Iω:↑<br>ExTraFol:none | (Moreau, Gosselin-Badaroudine, Boutjdir, et al., 2015) |
| 284 | Nav1.5 | R814W | Nav1.5:R814<br>Nav1.5-2:R814<br>Nav1.5-3:R814<br>Nav1.5-4:R814<br>Nav1.5-5:R814<br>Nav1.5-6:R814 | R947W | CMD1E*<br>DCM | GOF | Imax:↑<br>τrec:none<br>Ip:none<br>V1/2act:←<br>V1/2inact:none<br>Iω:none<br>ExTraFol:none | (Nguyen et al., 2008) |
| 285 | Nav1.5 | L839P | Nav1.5:L839<br>Nav1.5-2:L839<br>Nav1.5-3:L839<br>Nav1.5-4:L839<br>Nav1.5-5:L839<br>Nav1.5-6:L839 | L972P | BRGDA1 | LOF | Imax:↓<br>τrec:none<br>Ip:none<br>V1/2act:none<br>V1/2inact:none<br>Iω:none<br>ExTraFol:none | (Glazer et al., 2020) |
| 286 | Nav1.5 | L846R | Nav1.5:L846<br>Nav1.5-2:L846<br>Nav1.5-3:L846<br>Nav1.5-4:L846 | L979R | IVF | LOF | Imax:↓<br>τrec:none<br>Ip:none<br>V1/2act:none<br>V1/2inact:none | (Watanabe, Nogami, Ohkubo, Kawata, Hayashi, Ishikawa, |

| | Protein | Mutation | Isoform | Mutation (MSA) | Phenotype | GoF/LoF | Gating Properties | Reference |
|---|---|---|---|---|---|---|---|---|
| | | | Na$_v$1.5-5:L846<br>Na$_v$1.5-6:L846 | | | | Iω:none<br>ExTraFol:none | Makiyama, Nagao, Yagihara, Takehara, Shimizu, & Makita, 2011) |
| 287 | Na$_v$1.5 | F851L | Na$_v$1.5:F851<br>Na$_v$1.5-2:F851<br>Na$_v$1.5-3:F851<br>Na$_v$1.5-4:F851<br>Na$_v$1.5-5:F851<br>Na$_v$1.5-6:F851 | F984L | BRGDA1 | LOF | Imax:↓<br>τrec:none<br>Ip:none<br>V1/2act:none<br>V1/2inact:none<br>Iω:none<br>ExTraFol:none | (Glazer et al., 2020) |
| 288 | Na$_v$1.5 | R878C | Na$_v$1.5:R878<br>Na$_v$1.5-2:R878<br>Na$_v$1.5-3:R878<br>Na$_v$1.5-4:R878<br>Na$_v$1.5-5:R878<br>Na$_v$1.5-6:R878 | R1020C | BRGDA1 | LOF | Imax:↑<br>τrec:none<br>Ip:none<br>V1/2act:none<br>V1/2inact:none<br>Iω:none<br>ExTraFol:none | (Y. Zhang et al., 2008) (Clatot et al., 2020) |
| 289 | Na$_v$1.5 | R893C | Na$_v$1.5:R893<br>Na$_v$1.5-2:R893<br>Na$_v$1.5-3:R893<br>Na$_v$1.5-4:R893<br>Na$_v$1.5-5:R893<br>Na$_v$1.5-6:R893 | R1035C | BRGDA1 | LOF | Imax:↓<br>τrec:none<br>Ip:none<br>V1/2act:none<br>V1/2inact:none<br>Iω:none<br>ExTraFol:none | (Ishikawa et al., 2021) |
| 290 | Na$_v$1.5 | E901K | Na$_v$1.5:E901<br>Na$_v$1.5-2:E901<br>Na$_v$1.5-3:E901<br>Na$_v$1.5-4:E901<br>Na$_v$1.5-5:E901<br>Na$_v$1.5-6:E901 | E1043K | BRGDA1 | LOF | Imax:↓<br>τrec:none<br>Ip:none<br>V1/2act:none<br>V1/2inact:none<br>Iω:none<br>ExTraFol:none | (Glazer et al., 2020) |
| 291 | Na$_v$1.5 | N927S | Na$_v$1.5:N927<br>Na$_v$1.5-2:N927<br>Na$_v$1.5-3:N927<br>Na$_v$1.5-4:N927<br>Na$_v$1.5-5:N927<br>Na$_v$1.5-6:N927 | N1071S | BRGDA1 | LOF | Imax:↓<br>τrec:none<br>Ip:none<br>V1/2act:none<br>V1/2inact:none<br>Iω:none<br>ExTraFol:none | (Glazer et al., 2020) |
| 292 | Na$_v$1.5 | S941N | Na$_v$1.5:S941<br>Na$_v$1.5-2:S941<br>Na$_v$1.5-3:S941 | S1085N | LQT3 | GOF | Imax:none<br>τrec:none<br>Ip:↑ | (Ruan et al., 2007) |

| | Protein | Mutation | Isoform | Mutation (MSA) | Phenotype | GoF/LoF | Gating Properties | Reference |
|---|---|---|---|---|---|---|---|---|
| | | | Na$_v$1.5-4:S941<br>Na$_v$1.5-5:S941<br>Na$_v$1.5-6:S941 | | | | V1/2act:none<br>V1/2inact:none<br>Iω:none<br>ExTraFol:none | |
| 293 | Na$_v$1.5 | R965C | Na$_v$1.5:R965<br>Na$_v$1.5-2:R965<br>Na$_v$1.5-3:R965<br>Na$_v$1.5-4:R965<br>Na$_v$1.5-5:R965<br>Na$_v$1.5-6:R965 | R1110C | BRGDA1 | LOF | Imax:none<br>τrec:↓<br>Ip:none<br>V1/2act:none<br>V1/2inact:←<br>Iω:none<br>ExTraFol:none | (Hsueh et al., 2009) |
| 294 | Na$_v$1.5 | G983D | Na$_v$1.5:G983<br>Na$_v$1.5-2:G983<br>Na$_v$1.5-3:G983<br>Na$_v$1.5-4:G983<br>Na$_v$1.5-5:G983<br>Na$_v$1.5-6:G983 | G1128D | bizarre T-wave | GOF | Imax:↑<br>τrec:↑<br>Ip:none<br>V1/2act:←<br>V1/2inact:none<br>Iω:none<br>ExTraFol:none | (Ortiz-Bonnin et al., 2016) |
| 295 | Na$_v$1.5 | R986Q | Na$_v$1.5:R986<br>Na$_v$1.5-2:R986<br>Na$_v$1.5-3:R986<br>Na$_v$1.5-4:R986<br>Na$_v$1.5-5:R986<br>Na$_v$1.5-6:R986 | R1131Q | IBS | LOF | Imax:↓<br>τrec:↓<br>Ip:none<br>V1/2act:→<br>V1/2inact:→<br>Iω:none<br>ExTraFol:none | (Strege et al., 2018)<br>(Hayashi et al., 2015) |
| 296 | Na$_v$1.5 | A993T | Na$_v$1.5:A993<br>Na$_v$1.5-2:A993<br>Na$_v$1.5-3:A993<br>Na$_v$1.5-4:A993<br>Na$_v$1.5-5:A993<br>Na$_v$1.5-6:A993 | A1153T | LQT3 | GOF | Imax:↑<br>τrec:none<br>Ip:none<br>V1/2act:none<br>V1/2inact:none<br>Iω:none<br>ExTraFol:none | (Ortiz-Bonnin et al., 2016) |
| 297 | Na$_v$1.5 | A997S | Na$_v$1.5:A997<br>Na$_v$1.5-2:A997<br>Na$_v$1.5-3:A997<br>Na$_v$1.5-4:A997<br>Na$_v$1.5-5:A997<br>Na$_v$1.5-6:A997 | A1157S | RWS*<br>LQT3 | GOF | Imax:none<br>τrec:none<br>Ip:↑<br>V1/2act:none<br>V1/2inact:none<br>Iω:none<br>ExTraFol:none | (Ackerman et al., 2001) |
| 298 | Na$_v$1.5 | A997T | Na$_v$1.5:A997<br>Na$_v$1.5-2:A997<br>Na$_v$1.5-3:A997<br>Na$_v$1.5-4:A997 | A1157T | LQT3*<br>BRGDA1 | LOF | Imax:↓<br>τrec:none<br>Ip:↓<br>V1/2act:→ | (Beyder et al., 2014) |

| | Protein | Mutation | Isoform | Mutation (MSA) | Phenotype | GoF/LoF | Gating Properties | Reference |
|---|---|---|---|---|---|---|---|---|
| | | | $Na_v$1.5-5:A997<br>$Na_v$1.5-6:A997 | | | | V1/2inact:none<br>Iω:none<br>ExTraFol:none | |
| 299 | $Na_v$1.5 | P1008S | $Na_v$1.5:P1008<br>$Na_v$1.5-2:P1008<br>$Na_v$1.5-3:P1008<br>$Na_v$1.5-4:P1008<br>$Na_v$1.5-5:P1008<br>$Na_v$1.5-6:P1008 | P1178S | CCD | LOF | Imax:↓<br>τrec:↓<br>Ip:none<br>V1/2act:→<br>V1/2inact:none<br>Iω:none<br>ExTraFol:↓ | (D. Hu et al., 2010) |
| 300 | $Na_v$1.5 | P1014S | $Na_v$1.5:P1014<br>$Na_v$1.5-2:P1014<br>$Na_v$1.5-3:P1014<br>$Na_v$1.5-4:P1014<br>$Na_v$1.5-5:P1014<br>$Na_v$1.5-6:P1014 | P1184S | BRGDA1 | | Imax:none<br>τrec:none<br>Ip:none<br>V1/2act:←<br>V1/2inact:none<br>Iω:none<br>ExTraFol:none | (Glazer et al., 2020) |
| 301 | $Na_v$1.5 | R1023H | $Na_v$1.5:R1023<br>$Na_v$1.5-2:R1023<br>$Na_v$1.5-3:R1023<br>$Na_v$1.5-4:R1023<br>$Na_v$1.5-5:R1023<br>$Na_v$1.5-6:R1023 | R1193H | BRGDA1 | LOF | Imax:↓<br>τrec:↑<br>Ip:none<br>V1/2act:none<br>V1/2inact:none<br>Iω:none<br>ExTraFol:none | (Frustaci et al., 2005) |
| 302 | $Na_v$1.5 | R1192Q | $Na_v$1.5:R1193<br>$Na_v$1.5-2:R1192<br>$Na_v$1.5-3:R1192<br>$Na_v$1.5-4:R1193<br>$Na_v$1.5-5:R1139<br>$Na_v$1.5-6:R1193 | R1193Q | BRGDA1 | LOF | Imax:↓<br>τrec:none<br>Ip:none<br>V1/2act:none<br>V1/2inact:→<br>Iω:none<br>ExTraFol:none | (Vatta, Dumaine, Varghese, et al., 2002) |
| 303 | $Na_v$1.5 | E1053K | $Na_v$1.5:E1053<br>$Na_v$1.5-2:E1053<br>$Na_v$1.5-3:E1053<br>$Na_v$1.5-4:E1053<br>$Na_v$1.5-5:E1053<br>$Na_v$1.5-6:E1053 | E1241K | ATFB10*<br>LQT3*<br>BRGDA1 | LOF | Imax:↓<br>τrec:none<br>Ip:none<br>V1/2act:←<br>V1/2inact:none<br>Iω:none<br>ExTraFol:none | (Mohler et al., 2004) |
| 304 | $Na_v$1.5 | P1090L | $Na_v$1.5:P1090<br>$Na_v$1.5-2:P1089<br>$Na_v$1.5-3:P1089<br>$Na_v$1.5-4:P1090 | P1278L | SSS1*<br>LQT3 | GOF | Imax:none<br>τrec:none<br>Ip:none<br>V1/2act:←<br>V1/2inact:→ | (X. Wu et al., 2022)<br>(B.-H. Tan et al., 2005) |

| | Protein | Mutation | Isoform | Mutation (MSA) | Phenotype | GoF/LoF | Gating Properties | Reference |
|---|---|---|---|---|---|---|---|---|
| | | | Na$_v$1.5-5:--<br>Na$_v$1.5-6:P1090 | | | | Iω:none<br>ExTraFol:none | |
| 305 | Na$_v$1.5 | S1103Y | Na$_v$1.5:S1103<br>Na$_v$1.5-2:S1102<br>Na$_v$1.5-3:S1102<br>Na$_v$1.5-4:S1103<br>Na$_v$1.5-5:--<br>Na$_v$1.5-6:S1103 | S1291Y | BRGDA1*<br>LQT3 | GOF | Imax:none<br>τrec:↑<br>Ip:none<br>V1/2act:←<br>V1/2inact:→<br>Iω:none<br>ExTraFol:none | (J. Cheng et al., 2011)<br>(B.-H. Tan et al., 2005) |
| 306 | Na$_v$1.5 | P1177L | Na$_v$1.5:P1177<br>Na$_v$1.5-2:P1176<br>Na$_v$1.5-3:P1176<br>Na$_v$1.5-4:P1177<br>Na$_v$1.5-5:P1123<br>Na$_v$1.5-6:P1177 | P1368L | LQT3 | GOF | Imax:none<br>τrec:↓<br>Ip:↑<br>V1/2act:none<br>V1/2inact:none<br>Iω:none<br>ExTraFol:none | (Winkel et al., 2012) |
| 307 | Na$_v$1.5 | A1180V | Na$_v$1.5:A1180<br>Na$_v$1.5-2:A1179<br>Na$_v$1.5-3:A1179<br>Na$_v$1.5-4:A1180<br>Na$_v$1.5-5:A1126<br>Na$_v$1.5-6:A1180 | A1371V | DCM | LOF | Imax:none<br>τrec:↓<br>Ip:↑<br>V1/2act:none<br>V1/2inact:←<br>Iω:none<br>ExTraFol:none | (Ge et al., 2008) |
| 308 | Na$_v$1.5 | R1193Q | Na$_v$1.5:R1193<br>Na$_v$1.5-2:R1192<br>Na$_v$1.5-3:R1192<br>Na$_v$1.5-4:R1193<br>Na$_v$1.5-5:R1139<br>Na$_v$1.5-6:R1193 | R1384Q | BRGDA1<br>LQT3 | MIX | Imax:↑<br>τrec:none<br>Ip:↑<br>V1/2act:←<br>V1/2inact:←<br>Iω:none<br>ExTraFol:none | (Q. Wang et al., 2004) |
| 309 | Na$_v$1.5 | E1225K | Na$_v$1.5:E1225<br>Na$_v$1.5-2:E1224<br>Na$_v$1.5-3:E1224<br>Na$_v$1.5-4:E1225<br>Na$_v$1.5-5:E1171<br>Na$_v$1.5-6:E1225 | E1416K | LQT3*<br>BRGDA1 | LOF | Imax:↓<br>τrec:none<br>Ip:none<br>V1/2act:none<br>V1/2inact:→<br>Iω:none<br>ExTraFol:none | (Glazer et al., 2020) |
| 310 | Na$_v$1.5 | L1239P | Na$_v$1.5:L1239<br>Na$_v$1.5-2:L1238<br>Na$_v$1.5-3:L1238<br>Na$_v$1.5-4:L1239 | L1430P | BRGDA1 | LOF | Imax:↓<br>τrec:none<br>Ip:none<br>V1/2act:→<br>V1/2inact:none | (Y. Wang et al., 2020) |

| | Protein | Mutation | Isoform | Mutation (MSA) | Phenotype | GoF/LoF | Gating Properties | Reference |
|---|---|---|---|---|---|---|---|---|
| | | | Na$_v$1.5-5:L1185<br>Na$_v$1.5-6:L1239 | | | | Iω:none<br>ExTraFol:none | |
| 311 | Na$_v$1.5 | D1243N | Na$_v$1.5:D1243<br>Na$_v$1.5-2:D1242<br>Na$_v$1.5-3:D1242<br>Na$_v$1.5-4:D1243<br>Na$_v$1.5-5:D1189<br>Na$_v$1.5-6:D1243 | D1434N | BRGDA1 | | Imax:none<br>τrec:↑<br>Ip:none<br>V1/2act:→<br>V1/2inact:→<br>Iω:none<br>ExTraFol:none | (Glazer et al., 2020) |
| 312 | Na$_v$1.5 | F1250L | Na$_v$1.5:F1250<br>Na$_v$1.5-2:F1249<br>Na$_v$1.5-3:F1249<br>Na$_v$1.5-4:F1250<br>Na$_v$1.5-5:F1196<br>Na$_v$1.5-6:F1250 | F1441L | LQT3 | GOF | Imax:↓<br>τrec:none<br>Ip:none<br>V1/2act:→<br>V1/2inact:→<br>Iω:none<br>ExTraFol:none | (D. Han et al., 2018) |
| 313 | Na$_v$1.5 | D1275N | Na$_v$1.5:D1275<br>Na$_v$1.5-2:D1274<br>Na$_v$1.5-3:D1274<br>Na$_v$1.5-4:D1275<br>Na$_v$1.5-5:D1221<br>Na$_v$1.5-6:D1275 | D1466N | PFHB1A*<br>CMD1E*<br>ATFB10*<br>LQT3*<br>BRGDA1*<br>ATRST1*<br>CCD<br>SSS1 | LOF | Imax:↓<br>τrec:↓<br>Ip:none<br>V1/2act:→<br>V1/2inact:←<br>Iω:none<br>ExTraFol:none | (Gui et al., 2010) (Watanabe, Yang, et al., 2011) (Groenewegen et al., 2003) (Hayano et al., 2017) |
| 314 | Na$_v$1.5 | F1293S | Na$_v$1.5:F1293<br>Na$_v$1.5-2:F1292<br>Na$_v$1.5-3:F1292<br>Na$_v$1.5-4:F1293<br>Na$_v$1.5-5:F1239<br>Na$_v$1.5-6:F1293 | F1484S | BRGDA1*<br>IBS | LOF | Imax:↓<br>τrec:none<br>Ip:none<br>V1/2act:→<br>V1/2inact:none<br>Iω:none<br>ExTraFol:none | (Strege et al., 2018) |
| 315 | Na$_v$1.5 | E1295K | Na$_v$1.5:E1295<br>Na$_v$1.5-2:E1294<br>Na$_v$1.5-3:E1294<br>Na$_v$1.5-4:E1295<br>Na$_v$1.5-5:E1241<br>Na$_v$1.5-6:E1295 | E1486K | LQT3 | GOF | Imax:none<br>τrec:none<br>Ip:none<br>V1/2act:none<br>V1/2inact:→<br>Iω:none<br>ExTraFol:none | (Abriel et al., 2001) |
| 316 | Na$_v$1.5 | P1298L | Na$_v$1.5:P1298<br>Na$_v$1.5-2:P1297<br>Na$_v$1.5-3:P1297<br>Na$_v$1.5-4:P1298 | P1489L | SSS1 | LOF | Imax:↓<br>τrec:none<br>Ip:none<br>V1/2act:none<br>V1/2inact:← | (Gui et al., 2010) |

| | Protein | Mutation | Isoform | Mutation (MSA) | Phenotype | GoF/LoF | Gating Properties | Reference |
|---|---|---|---|---|---|---|---|---|
| | | | Na$_v$1.5-5:P1244<br>Na$_v$1.5-6:P1298 | | | | Iω:none<br>ExTraFol:none | |
| 317 | Na$_v$1.5 | K1300H | Na$_v$1.5:K1300<br>Na$_v$1.5-2:K1299<br>Na$_v$1.5-3:K1299<br>Na$_v$1.5-4:K1300<br>Na$_v$1.5-5:K1246<br>Na$_v$1.5-6:K1300 | K1491H | | | Imax:none<br>τrec:none<br>Ip:none<br>V1/2act:none<br>V1/2inact:←<br>Iω:none<br>ExTraFol:none | (L. Q. Chen et al., 1996) |
| 318 | Na$_v$1.5 | K1300Q | Na$_v$1.5:K1300<br>Na$_v$1.5-2:K1299<br>Na$_v$1.5-3:K1299<br>Na$_v$1.5-4:K1300<br>Na$_v$1.5-5:K1246<br>Na$_v$1.5-6:K1300 | K1491Q | | | Imax:none<br>τrec:none<br>Ip:none<br>V1/2act:←<br>V1/2inact:←<br>Iω:none<br>ExTraFol:none | (L. Q. Chen et al., 1996) |
| 319 | Na$_v$1.5 | T1304M | Na$_v$1.5:T1304<br>Na$_v$1.5-2:T1303<br>Na$_v$1.5-3:T1303<br>Na$_v$1.5-4:T1304<br>Na$_v$1.5-5:T1250<br>Na$_v$1.5-6:T1304 | T1495M | LQT3 | GOF | Imax:none<br>τrec:↑<br>Ip:↑<br>V1/2act:→<br>V1/2inact:→<br>Iω:none<br>ExTraFol:none | (Beyder et al., 2014)<br>(Olesen et al., 2012) |
| 320 | Na$_v$1.5 | R1306E | Na$_v$1.5:R1306<br>Na$_v$1.5-2:R1305<br>Na$_v$1.5-3:R1305<br>Na$_v$1.5-4:R1306<br>Na$_v$1.5-5:R1252<br>Na$_v$1.5-6:R1306 | R1497E | | | Imax:none<br>τrec:none<br>Ip:none<br>V1/2act:→<br>V1/2inact:←<br>Iω:none<br>ExTraFol:none | (L. Q. Chen et al., 1996) |
| 321 | Na$_v$1.5 | R1306Q | Na$_v$1.5:R1306<br>Na$_v$1.5-2:R1305<br>Na$_v$1.5-3:R1305<br>Na$_v$1.5-4:R1306<br>Na$_v$1.5-5:R1252<br>Na$_v$1.5-6:R1306 | R1497Q | | | Imax:none<br>τrec:none<br>Ip:none<br>V1/2act:none<br>V1/2inact:←<br>Iω:none<br>ExTraFol:none | (L. Q. Chen et al., 1996) |
| 322 | Na$_v$1.5 | P1310L | Na$_v$1.5:P1310<br>Na$_v$1.5-2:P1309<br>Na$_v$1.5-3:P1309<br>Na$_v$1.5-4:P1310 | P1501L | BRGDA1 | LOF | Imax:↓<br>τrec:↑<br>Ip:none<br>V1/2act:→<br>V1/2inact:none | (Balla et al., 2021) |

| | Protein | Mutation | Isoform | Mutation (MSA) | Phenotype | GoF/LoF | Gating Properties | Reference |
|---|---|---|---|---|---|---|---|---|
| | | | $Na_v1.5$-5:P1256<br>$Na_v1.5$-6:P1310 | | | | Iω:none<br>ExTraFol:none | |
| 323 | $Na_v1.5$ | G1319V | $Na_v1.5$:G1319<br>$Na_v1.5$-2:G1318<br>$Na_v1.5$-3:G1318<br>$Na_v1.5$-4:G1319<br>$Na_v1.5$-5:G1265<br>$Na_v1.5$-6:G1319 | G1510V | BRGDA1 | LOF | Imax:none<br>τrec:none<br>Ip:none<br>V1/2act:→<br>V1/2inact:←<br>Iω:none<br>ExTraFol:none | (Casini et al., 2007) |
| 324 | $Na_v1.5$ | N1325S | $Na_v1.5$:N1325<br>$Na_v1.5$-2:N1324<br>$Na_v1.5$-3:N1324<br>$Na_v1.5$-4:N1325<br>$Na_v1.5$-5:N1271<br>$Na_v1.5$-6:N1325 | N1516S | RWS*<br>LQT3 | GOF | Imax:none<br>τrec:none<br>Ip:↑<br>V1/2act:←<br>V1/2inact:→<br>Iω:none<br>ExTraFol:none | (D. W. Wang et al., 1996) (Glazer et al., 2020) |
| 325 | $Na_v1.5$ | A1330P | $Na_v1.5$:A1330<br>$Na_v1.5$-2:A1329<br>$Na_v1.5$-3:A1329<br>$Na_v1.5$-4:A1330<br>$Na_v1.5$-5:A1276<br>$Na_v1.5$-6:A1330 | A1521P | RWS*<br>LQT3 | GOF | Imax:none<br>τrec:none<br>Ip:none<br>V1/2act:none<br>V1/2inact:→<br>Iω:none<br>ExTraFol:none | (Wedekind et al., 2001) (Berecki et al., 2006) |
| 326 | $Na_v1.5$ | A1330T | $Na_v1.5$:A1330<br>$Na_v1.5$-2:A1329<br>$Na_v1.5$-3:A1329<br>$Na_v1.5$-4:A1330<br>$Na_v1.5$-5:A1276<br>$Na_v1.5$-6:A1330 | A1521T | RWS*<br>LQT3 | GOF | Imax:none<br>τrec:↑<br>Ip:none<br>V1/2act:none<br>V1/2inact:→<br>Iω:none<br>ExTraFol:none | (Smits, Veldkamp, et al., 2005) |
| 327 | $Na_v1.5$ | P1332L | $Na_v1.5$:P1332<br>$Na_v1.5$-2:P1331<br>$Na_v1.5$-3:P1331<br>$Na_v1.5$-4:P1332<br>$Na_v1.5$-5:P1278<br>$Na_v1.5$-6:P1332 | P1523L | BRGDA1*<br>LQT3 | GOF | Imax:none<br>τrec:↓<br>Ip:none<br>V1/2act:←<br>V1/2inact:←<br>Iω:none<br>ExTraFol:none | (Ruan et al., 2007) |
| 328 | $Na_v1.5$ | S1333Y | $Na_v1.5$:S1333<br>$Na_v1.5$-2:S1332<br>$Na_v1.5$-3:S1332<br>$Na_v1.5$-4:S1333 | S1524Y | RWS*<br>SIDS*<br>LQT3 | GOF | Imax:none<br>τrec:none<br>Ip:none<br>V1/2act:←<br>V1/2inact:→ | (H. Huang et al., 2009a) |

| | Protein | Mutation | Isoform | Mutation (MSA) | Phenotype | GoF/LoF | Gating Properties | Reference |
|---|---|---|---|---|---|---|---|---|
| | | | Na$_v$1.5-5:S1279<br>Na$_v$1.5-6:S1333 | | | | Iω:none<br>ExTraFol:none | |
| 329 | Na$_v$1.5 | V1340L | Na$_v$1.5:V1340<br>Na$_v$1.5-2:V1339<br>Na$_v$1.5-3:V1339<br>Na$_v$1.5-4:V1340<br>Na$_v$1.5-5:V1286<br>Na$_v$1.5-6:V1340 | V1531L | AS | LOF | Imax:↓<br>τrec:none<br>Ip:none<br>V1/2act:→<br>V1/2inact:none<br>Iω:none<br>ExTraFol:none | (R. B. Tan et al., 2018) |
| 330 | Na$_v$1.5 | I1343V | Na$_v$1.5:I1343<br>Na$_v$1.5-2:I1342<br>Na$_v$1.5-3:I1342<br>Na$_v$1.5-4:I1343<br>Na$_v$1.5-5:I1289<br>Na$_v$1.5-6:I1343 | I1534V | AF | LOF | Imax:↓<br>τrec:none<br>Ip:none<br>V1/2act:none<br>V1/2inact:none<br>Iω:none<br>ExTraFol:none | (Boddum et al., 2018) |
| 331 | Na$_v$1.5 | F1344S | Na$_v$1.5:F1344<br>Na$_v$1.5-2:F1343<br>Na$_v$1.5-3:F1343<br>Na$_v$1.5-4:F1344<br>Na$_v$1.5-5:F1290<br>Na$_v$1.5-6:F1344 | F1535S | BRGDA1 | LOF | Imax:↓<br>τrec:none<br>Ip:none<br>V1/2act:→<br>V1/2inact:none<br>Iω:none<br>ExTraFol:none | (Keller et al., 2006) |
| 332 | Na$_v$1.5 | W1345C | Na$_v$1.5:W1345<br>Na$_v$1.5-2:W1344<br>Na$_v$1.5-3:W1344<br>Na$_v$1.5-4:W1345<br>Na$_v$1.5-5:W1291<br>Na$_v$1.5-6:W1345 | W1536C | BRGDA1 | LOF | Imax:↓<br>τrec:none<br>Ip:none<br>V1/2act:none<br>V1/2inact:none<br>Iω:none<br>ExTraFol:none | (Glazer et al., 2020) |
| 333 | Na$_v$1.5 | L1346P | Na$_v$1.5:L1346<br>Na$_v$1.5-2:L1345<br>Na$_v$1.5-3:L1345<br>Na$_v$1.5-4:L1346<br>Na$_v$1.5-5:L1292<br>Na$_v$1.5-6:L1346 | L1537P | BRGDA1 | LOF | Imax:↓<br>τrec:none<br>Ip:none<br>V1/2act:none<br>V1/2inact:none<br>Iω:none<br>ExTraFol:none | (Glazer et al., 2020) |
| 334 | Na$_v$1.5 | S1382I | Na$_v$1.5:S1382<br>Na$_v$1.5-2:S1381<br>Na$_v$1.5-3:S1381<br>Na$_v$1.5-4:S1382 | S1574I | BRGDA1 | LOF | Imax:↓<br>τrec:none<br>Ip:none<br>V1/2act:none<br>V1/2inact:none | (Glazer et al., 2020) |

| | Protein | Mutation | Isoform | Mutation (MSA) | Phenotype | GoF/LoF | Gating Properties | Reference |
|---|---|---|---|---|---|---|---|---|
| | | | $Na_v$1.5-5:S1328<br>$Na_v$1.5-6:S1382 | | | | Iω:none<br>ExTraFol:none | |
| 335 | $Na_v$1.5 | K1397R | $Na_v$1.5:K1397<br>$Na_v$1.5-2:K1396<br>$Na_v$1.5-3:K1396<br>$Na_v$1.5-4:K1397<br>$Na_v$1.5-5:K1343<br>$Na_v$1.5-6:K1397 | K1591R | BRGDA1 | LOF | Imax:↓<br>τrec:none<br>Ip:none<br>V1/2act:none<br>V1/2inact:none<br>Iω:none<br>ExTraFol:none | (Q. Chen et al., 1998) |
| 336 | $Na_v$1.5 | V1405L | $Na_v$1.5:V1405<br>$Na_v$1.5-2:V1404<br>$Na_v$1.5-3:V1404<br>$Na_v$1.5-4:V1405<br>$Na_v$1.5-5:V1351<br>$Na_v$1.5-6:V1405 | V1599L | BRGDA1 | LOF | Imax:↓<br>τrec:none<br>Ip:none<br>V1/2act:none<br>V1/2inact:none<br>Iω:none<br>ExTraFol:none | (Glazer et al., 2020) |
| 337 | $Na_v$1.5 | V1405M | $Na_v$1.5:V1405<br>$Na_v$1.5-2:V1404<br>$Na_v$1.5-3:V1404<br>$Na_v$1.5-4:V1405<br>$Na_v$1.5-5:V1351<br>$Na_v$1.5-6:V1405 | V1599M | BRGDA1 | LOF | Imax:↓<br>τrec:none<br>Ip:none<br>V1/2act:none<br>V1/2inact:none<br>Iω:none<br>ExTraFol:none | (Zhu et al., 2021)<br>(Glazer et al., 2020) |
| 338 | $Na_v$1.5 | G1406E | $Na_v$1.5:G1406<br>$Na_v$1.5-2:G1405<br>$Na_v$1.5-3:G1405<br>$Na_v$1.5-4:G1406<br>$Na_v$1.5-5:G1352<br>$Na_v$1.5-6:G1406 | G1600E | BRGDA1 | LOF | Imax:↓<br>τrec:none<br>Ip:none<br>V1/2act:none<br>V1/2inact:none<br>Iω:none<br>ExTraFol:none | (Glazer et al., 2020) |
| 339 | $Na_v$1.5 | G1406R | $Na_v$1.5:G1406<br>$Na_v$1.5-2:G1405<br>$Na_v$1.5-3:G1405<br>$Na_v$1.5-4:G1406<br>$Na_v$1.5-5:G1352<br>$Na_v$1.5-6:G1406 | G1600R | BRGDA1 | LOF | Imax:↓<br>τrec:none<br>Ip:none<br>V1/2act:none<br>V1/2inact:none<br>Iω:none<br>ExTraFol:none | (Kyndt et al., 2001) |
| 340 | $Na_v$1.5 | G1408R | $Na_v$1.5:G1408<br>$Na_v$1.5-2:G1407<br>$Na_v$1.5-3:G1407<br>$Na_v$1.5-4:G1408 | G1602R | SSS1*<br>BRGDA1 | LOF | Imax:↓<br>τrec:none<br>Ip:none<br>V1/2act:none<br>V1/2inact:none | (Kyndt et al., 2001) |

| | Protein | Mutation | Isoform | Mutation (MSA) | Phenotype | GoF/LoF | Gating Properties | Reference |
|---|---|---|---|---|---|---|---|---|
| | | | $Na_v$1.5-5:G1354<br>$Na_v$1.5-6:G1408 | | | | Iω:none<br>ExTraFol:none | |
| 341 | $Na_v$1.5 | L1412F | $Na_v$1.5:L1412<br>$Na_v$1.5-2:L1411<br>$Na_v$1.5-3:L1411<br>$Na_v$1.5-4:L1412<br>$Na_v$1.5-5:L1358<br>$Na_v$1.5-6:L1412 | L1606F | BRGDA1*<br>ERS | LOF | Imax:↓<br>τrec:none<br>Ip:none<br>V1/2act:none<br>V1/2inact:←<br>Iω:none<br>ExTraFol:none | (Z.-H. Zhang et al., 2021) |
| 342 | $Na_v$1.5 | G1420R | $Na_v$1.5:G1420<br>$Na_v$1.5-2:G1419<br>$Na_v$1.5-3:G1419<br>$Na_v$1.5-4:G1420<br>$Na_v$1.5-5:G1366<br>$Na_v$1.5-6:-- | G1614R | BRGDA1 | LOF | Imax:↓<br>τrec:none<br>Ip:none<br>V1/2act:none<br>V1/2inact:none<br>Iω:none<br>ExTraFol:none | (Glazer et al., 2020) |
| 343 | $Na_v$1.5 | D1430N | $Na_v$1.5:D1430<br>$Na_v$1.5-2:D1429<br>$Na_v$1.5-3:D1429<br>$Na_v$1.5-4:D1430<br>$Na_v$1.5-5:D1376<br>$Na_v$1.5-6:-- | D1624N | BRGDA1 | LOF | Imax:↓<br>τrec:none<br>Ip:none<br>V1/2act:none<br>V1/2inact:none<br>Iω:none<br>ExTraFol:none | (Maury et al., 2013) |
| 344 | $Na_v$1.5 | R1432G | $Na_v$1.5:R1432<br>$Na_v$1.5-2:R1431<br>$Na_v$1.5-3:R1431<br>$Na_v$1.5-4:R1432<br>$Na_v$1.5-5:R1378<br>$Na_v$1.5-6:-- | R1626G | BRGDA1 | LOF | Imax:↓<br>τrec:none<br>Ip:none<br>V1/2act:none<br>V1/2inact:none<br>Iω:none<br>ExTraFol:none | (Deschênes et al., 2000)<br>(Baroudi et al., 2001)<br>(Mercier et al., 2012)<br>(Glazer et al., 2020) |
| 345 | $Na_v$1.5 | P1438L | $Na_v$1.5:P1438<br>$Na_v$1.5-2:P1437<br>$Na_v$1.5-3:P1437<br>$Na_v$1.5-4:P1438<br>$Na_v$1.5-5:P1384<br>$Na_v$1.5-6:P1420 | P1632L | BRGDA1 | LOF | Imax:↓<br>τrec:none<br>Ip:none<br>V1/2act:none<br>V1/2inact:none<br>Iω:none<br>ExTraFol:none | (Six et al., 2008) |
| 346 | $Na_v$1.5 | Y1449C | $Na_v$1.5:Y1449<br>$Na_v$1.5-2:Y1448<br>$Na_v$1.5-3:Y1448<br>$Na_v$1.5-4:Y1449 | Y1643C | BRGDA1 | LOF | Imax:↓<br>τrec:none<br>Ip:none<br>V1/2act:none<br>V1/2inact:none | (Glazer et al., 2020) |

| | Protein | Mutation | Isoform | Mutation (MSA) | Phenotype | GoF/LoF | Gating Properties | Reference |
|---|---|---|---|---|---|---|---|---|
| | | | Na$_v$1.5-5:Y1395<br>Na$_v$1.5-6:Y1431 | | | | Iω:none<br>ExTraFol:none | |
| 347 | Na$_v$1.5 | Y1449H | Na$_v$1.5:Y1449<br>Na$_v$1.5-2:Y1448<br>Na$_v$1.5-3:Y1448<br>Na$_v$1.5-4:Y1449<br>Na$_v$1.5-5:Y1395<br>Na$_v$1.5-6:Y1431 | Y1643H | BRGDA1 | LOF | Imax:↓<br>τrec:none<br>Ip:none<br>V1/2act:none<br>V1/2inact:none<br>Iω:none<br>ExTraFol:none | (Arana-Rueda et al., 2021) |
| 348 | Na$_v$1.5 | T1461S | Na$_v$1.5:T1461<br>Na$_v$1.5-2:T1460<br>Na$_v$1.5-3:T1460<br>Na$_v$1.5-4:T1461<br>Na$_v$1.5-5:T1407<br>Na$_v$1.5-6:T1443 | T1655S | BRGDA1 | LOF | Imax:↓<br>τrec:↑<br>Ip:none<br>V1/2act:none<br>V1/2inact:none<br>Iω:none<br>ExTraFol:none | (Glazer et al., 2020) |
| 349 | Na$_v$1.5 | F1465L | Na$_v$1.5:F1465<br>Na$_v$1.5-2:F1464<br>Na$_v$1.5-3:F1464<br>Na$_v$1.5-4:F1465<br>Na$_v$1.5-5:F1411<br>Na$_v$1.5-6:F1447 | F1659L | LQT3*<br>Tachycardia | GOF | Imax:none<br>τrec:none<br>Ip:none<br>V1/2act:←<br>V1/2inact:none<br>Iω:none<br>ExTraFol:none | (Goktas Sahoglu et al., 2022) |
| 350 | Na$_v$1.5 | F1473C | Na$_v$1.5:F1473<br>Na$_v$1.5-2:F1472<br>Na$_v$1.5-3:F1472<br>Na$_v$1.5-4:F1473<br>Na$_v$1.5-5:F1419<br>Na$_v$1.5-6:F1455 | F1667C | RWS*<br>LQT3 | GOF | Imax:none<br>τrec:none<br>Ip:↑<br>V1/2act:none<br>V1/2inact:→<br>Iω:none<br>ExTraFol:none | (Bankston et al., 2007) |
| 351 | Na$_v$1.5 | F1473S | Na$_v$1.5:F1473<br>Na$_v$1.5-2:F1472<br>Na$_v$1.5-3:F1472<br>Na$_v$1.5-4:F1473<br>Na$_v$1.5-5:F1419<br>Na$_v$1.5-6:F1455 | F1667S | RWS*<br>LQT3 | GOF | Imax:↑<br>τrec:none<br>Ip:↑<br>V1/2act:none<br>V1/2inact:→<br>Iω:none<br>ExTraFol:none | (Ruan et al., 2010) |
| 352 | Na$_v$1.5 | Q1476R | Na$_v$1.5:Q1476<br>Na$_v$1.5-2:Q1475<br>Na$_v$1.5-3:Q1475<br>Na$_v$1.5-4:Q1476 | Q1670R | LQT3 | | Imax:none<br>τrec:none<br>Ip:↑<br>V1/2act:none<br>V1/2inact:→ | (Moreau et al., 2013) |

| | Protein | Mutation | Isoform | Mutation (MSA) | Phenotype | GoF/LoF | Gating Properties | Reference |
|---|---|---|---|---|---|---|---|---|
| | | | $Na_v$1.5-5:Q1422<br>$Na_v$1.5-6:Q1458 | | | | Iω:none<br>ExTraFol:none | |
| 353 | $Na_v$1.5 | F1486L | $Na_v$1.5:F1486<br>$Na_v$1.5-2:F1485<br>$Na_v$1.5-3:F1485<br>$Na_v$1.5-4:F1486<br>$Na_v$1.5-5:F1432<br>$Na_v$1.5-6:F1468 | F1680L | SIDS*<br>LQT3 | GOF | Imax:none<br>τrec:none<br>Ip:↑<br>V1/2act:none<br>V1/2inact:→<br>Iω:none<br>ExTraFol:none | (D. W. Wang et al., 2007) |
| 354 | $Na_v$1.5 | K1493R | $Na_v$1.5:K1493<br>$Na_v$1.5-2:K1492<br>$Na_v$1.5-3:K1492<br>$Na_v$1.5-4:K1493<br>$Na_v$1.5-5:K1439<br>$Na_v$1.5-6:K1475 | K1687R | LQT3*<br>AF | GOF | Imax:none<br>τrec:none<br>Ip:none<br>V1/2act:none<br>V1/2inact:→<br>Iω:none<br>ExTraFol:none | (Q. Li et al., 2009) |
| 355 | $Na_v$1.5 | Y1495F | $Na_v$1.5:Y1495<br>$Na_v$1.5-2:Y1494<br>$Na_v$1.5-3:Y1494<br>$Na_v$1.5-4:Y1495<br>$Na_v$1.5-5:Y1441<br>$Na_v$1.5-6:Y1477 | Y1689F | | | Imax:none<br>τrec:none<br>Ip:none<br>V1/2act:none<br>V1/2inact:→<br>Iω:none<br>ExTraFol:none | (Ahern et al., 2005) |
| 356 | $Na_v$1.5 | R1512W | $Na_v$1.5:R1512<br>$Na_v$1.5-2:R1511<br>$Na_v$1.5-3:R1511<br>$Na_v$1.5-4:R1512<br>$Na_v$1.5-5:R1458<br>$Na_v$1.5-6:R1494 | R1706W | BRGDA1 | LOF | Imax:none<br>τrec:none<br>Ip:none<br>V1/2act:←<br>V1/2inact:none<br>Iω:none<br>ExTraFol:none | (Makiyama et al., 2005) |
| 357 | $Na_v$1.5 | K1527R | $Na_v$1.5:K1527<br>$Na_v$1.5-2:K1526<br>$Na_v$1.5-3:K1526<br>$Na_v$1.5-4:K1527<br>$Na_v$1.5-5:K1473<br>$Na_v$1.5-6:K1509 | K1721R | BRGDA1 | LOF | Imax:none<br>τrec:none<br>Ip:none<br>V1/2act:none<br>V1/2inact:←<br>Iω:none<br>ExTraFol:none | (Yokoi et al., 2005) |
| 358 | $Na_v$1.5 | V1762L | $Na_v$1.5:V1763<br>$Na_v$1.5-2:V1762<br>$Na_v$1.5-3:V1730<br>$Na_v$1.5-4:V1763 | V1763L | LQT3 | GOF | Imax:none<br>τrec:none<br>Ip:↑<br>V1/2act:none<br>V1/2inact:none | (Priest et al., 2016) |

| | Protein | Mutation | Isoform | Mutation (MSA) | Phenotype | GoF/LoF | Gating Properties | Reference |
|---|---|---|---|---|---|---|---|---|
| | | | Nav1.5-5:V1709<br>Nav1.5-6:V1745 | | | | Iω:none<br>ExTraFol:none | |
| 359 | Nav1.5 | E1574K | Nav1.5:E1574<br>Nav1.5-2:E1573<br>Nav1.5-3:--<br>Nav1.5-4:E1574<br>Nav1.5-5:E1520<br>Nav1.5-6:E1556 | E1768K | BRGDA1 | LOF | Imax:↓<br>τrec:none<br>Ip:none<br>V1/2act:→<br>V1/2inact:none<br>Iω:none<br>ExTraFol:none | (Glazer et al., 2020) |
| 360 | Nav1.5 | R1583C | Nav1.5:R1583<br>Nav1.5-2:R1582<br>Nav1.5-3:--<br>Nav1.5-4:R1583<br>Nav1.5-5:R1529<br>Nav1.5-6:R1565 | R1777C | BRGDA1 | | Imax:none<br>τrec:↑<br>Ip:none<br>V1/2act:none<br>V1/2inact:none<br>Iω:none<br>ExTraFol:none | (Glazer et al., 2020) |
| 361 | Nav1.5 | D1595H | Nav1.5:D1595<br>Nav1.5-2:D1594<br>Nav1.5-3:--<br>Nav1.5-4:D1595<br>Nav1.5-5:D1541<br>Nav1.5-6:D1577 | D1789H | CMD1E*<br>BRGDA1*<br>DCM | | Imax:↓<br>τrec:↓<br>Ip:none<br>V1/2act:none<br>V1/2inact:←<br>Iω:none<br>ExTraFol:none | (Nguyen et al., 2008) |
| 362 | Nav1.5 | D1595N | Nav1.5:D1595<br>Nav1.5-2:D1594<br>Nav1.5-3:--<br>Nav1.5-4:D1595<br>Nav1.5-5:D1541<br>Nav1.5-6:D1577 | D1789N | PFHB1A*<br>ATFB10*<br>CCD | LOF | Imax:↓<br>τrec:none<br>Ip:none<br>V1/2act:→<br>V1/2inact:→<br>Iω:none<br>ExTraFol:none | (D. W. Wang et al., 2002) |
| 363 | Nav1.5 | T1620M | Nav1.5:T1620<br>Nav1.5-2:T1619<br>Nav1.5-3:T1587<br>Nav1.5-4:T1620<br>Nav1.5-5:T1566<br>Nav1.5-6:T1602 | T1818M | BRGDA1 | LOF | Imax:↓<br>τrec:none<br>Ip:none<br>V1/2act:→<br>V1/2inact:←<br>Iω:none<br>ExTraFol:none | (Shirai et al., 2002)<br>(Dumaine et al., 1999) |
| 364 | Nav1.5 | T1620K | Nav1.5:T1620<br>Nav1.5-2:T1619<br>Nav1.5-3:T1587<br>Nav1.5-4:T1620 | T1818K | PFHB1A*<br>BRGDA1*<br>CCD<br>LQT3 | LOF | Imax:none<br>τrec:↑<br>Ip:↑<br>V1/2act:←<br>V1/2inact:← | (Surber et al., 2008)<br>(Walzik et al., 2011) |

| | Protein | Mutation | Isoform | Mutation (MSA) | Phenotype | GoF/LoF | Gating Properties | Reference |
|---|---|---|---|---|---|---|---|---|
| | | | $Na_v$1.5-5:T1566<br>$Na_v$1.5-6:T1602 | | | | Iω:none<br>ExTraFol:none | |
| 365 | $Na_v$1.5 | R1623Q | $Na_v$1.5:R1623<br>$Na_v$1.5-2:R1622<br>$Na_v$1.5-3:R1590<br>$Na_v$1.5-4:R1623<br>$Na_v$1.5-5:R1569<br>$Na_v$1.5-6:R1605 | R1821Q | BRGDA1*<br>RWS*<br>LQT3 | GOF | Imax:none<br>τrec:↓<br>Ip:↑<br>V1/2act:none<br>V1/2inact:→<br>Iω:none<br>ExTraFol:none | (Kambouris et al., 1998)<br>(Rajamani et al., 2009)<br>(Philippaert et al., 2021) |
| 366 | $Na_v$1.5 | R1626P | $Na_v$1.5:R1626<br>$Na_v$1.5-2:R1625<br>$Na_v$1.5-3:R1593<br>$Na_v$1.5-4:R1626<br>$Na_v$1.5-5:R1572<br>$Na_v$1.5-6:R1608 | R1824P | RWS*<br>LQT3 | GOF | Imax:none<br>τrec:none<br>Ip:↑<br>V1/2act:→<br>V1/2inact:←<br>Iω:none<br>ExTraFol:none | (Ruan et al., 2007)<br>(Olesen et al., 2012) |
| 367 | $Na_v$1.5 | R1629E | $Na_v$1.5:R1629<br>$Na_v$1.5-2:R1628<br>$Na_v$1.5-3:R1596<br>$Na_v$1.5-4:R1629<br>$Na_v$1.5-5:R1575<br>$Na_v$1.5-6:R1611 | R1827E | | | Imax:none<br>τrec:none<br>Ip:none<br>V1/2act:→<br>V1/2inact:←<br>Iω:none<br>ExTraFol:none | (L. Q. Chen et al., 1996) |
| 368 | $Na_v$1.5 | R1629Q | $Na_v$1.5:R1629<br>$Na_v$1.5-2:R1628<br>$Na_v$1.5-3:R1596<br>$Na_v$1.5-4:R1629<br>$Na_v$1.5-5:R1575<br>$Na_v$1.5-6:R1611 | R1827Q | LQT3*<br>BRGDA1 | LOF | Imax:none<br>τrec:↓<br>Ip:none<br>V1/2act:→<br>V1/2inact:←<br>Iω:none<br>ExTraFol:none | (Zeng et al., 2013)<br>(L. Q. Chen et al., 1996) |
| 369 | $Na_v$1.5 | R1632C | $Na_v$1.5:R1632<br>$Na_v$1.5-2:R1631<br>$Na_v$1.5-3:R1599<br>$Na_v$1.5-4:R1632<br>$Na_v$1.5-5:R1578<br>$Na_v$1.5-6:R1614 | R1830C | BRGDA1 | LOF | Imax:↓<br>τrec:↓<br>Ip:none<br>V1/2act:none<br>V1/2inact:←<br>Iω:none<br>ExTraFol:none | (Nakajima et al., 2015) |
| 370 | $Na_v$1.5 | R1632H | $Na_v$1.5:R1632<br>$Na_v$1.5-2:R1631<br>$Na_v$1.5-3:R1599<br>$Na_v$1.5-4:R1632 | R1830H | BRGDA1*<br>SIDS*<br>SSS1 | LOF | Imax:none<br>τrec:↓<br>Ip:none<br>V1/2act:none<br>V1/2inact:← | (Benson et al., 2003)<br>(Gui et al., 2010)<br>(Glazer et al., 2020) |

| | Protein | Mutation | Isoform | Mutation (MSA) | Phenotype | GoF/LoF | Gating Properties | Reference |
|---|---|---|---|---|---|---|---|---|
| | | | Na$_v$1.5-5:R1578<br>Na$_v$1.5-6:R1614 | | | | Iω:none<br>ExTraFol:none | |
| 371 | Na$_v$1.5 | G1642E | Na$_v$1.5:G1642<br>Na$_v$1.5-2:G1641<br>Na$_v$1.5-3:G1609<br>Na$_v$1.5-4:G1642<br>Na$_v$1.5-5:G1588<br>Na$_v$1.5-6:G1624 | G1840E | BRGDA1 | LOF | Imax:↓<br>τrec:↑<br>Ip:none<br>V1/2act:none<br>V1/2inact:→<br>Iω:none<br>ExTraFol:none | (Glazer et al., 2020) |
| 372 | Na$_v$1.5 | R1644C | Na$_v$1.5:R1644<br>Na$_v$1.5-2:R1643<br>Na$_v$1.5-3:R1611<br>Na$_v$1.5-4:R1644<br>Na$_v$1.5-5:R1590<br>Na$_v$1.5-6:R1626 | R1842C | LQT3*<br>RWS*<br>BRGDA1 | LOF | Imax:none<br>τrec:↑<br>Ip:none<br>V1/2act:→<br>V1/2inact:none<br>Iω:none<br>ExTraFol:none | (Kroncke et al., 2018) |
| 373 | Na$_v$1.5 | R1644H | Na$_v$1.5:R1644<br>Na$_v$1.5-2:R1643<br>Na$_v$1.5-3:R1611<br>Na$_v$1.5-4:R1644<br>Na$_v$1.5-5:R1590<br>Na$_v$1.5-6:R1626 | R1842H | RWS*<br>LQT3 | GOF | Imax:none<br>τrec:none<br>Ip:↑<br>V1/2act:none<br>V1/2inact:→<br>Iω:none<br>ExTraFol:none | (D. W. Wang et al., 1996) |
| 374 | Na$_v$1.5 | A1649V | Na$_v$1.5:A1649<br>Na$_v$1.5-2:A1648<br>Na$_v$1.5-3:A1616<br>Na$_v$1.5-4:A1649<br>Na$_v$1.5-5:A1595<br>Na$_v$1.5-6:A1631 | A1847V | BRGDA1 | LOF | Imax:none<br>τrec:none<br>Ip:none<br>V1/2act:←<br>V1/2inact:←<br>Iω:none<br>ExTraFol:none | (L. Tang et al., 1998) |
| 375 | Na$_v$1.5 | M1652R | Na$_v$1.5:M1652<br>Na$_v$1.5-2:M1651<br>Na$_v$1.5-3:M1619<br>Na$_v$1.5-4:M1652<br>Na$_v$1.5-5:M1598<br>Na$_v$1.5-6:M1634 | M1850R | RWS*<br>LQT3 | GOF | Imax:none<br>τrec:none<br>Ip:↑<br>V1/2act:→<br>V1/2inact:→<br>Iω:none<br>ExTraFol:none | (Ruan et al., 2007) |
| 376 | Na$_v$1.5 | I1660V | Na$_v$1.5:I1660<br>Na$_v$1.5-2:I1659<br>Na$_v$1.5-3:I1627<br>Na$_v$1.5-4:I1660 | I1858V | LQT3*<br>BRGDA1 | LOF | Imax:↓<br>τrec:none<br>Ip:none<br>V1/2act:→<br>V1/2inact:← | (Cordeiro et al., 2006) |

| | Protein | Mutation | Isoform | Mutation (MSA) | Phenotype | GoF/LoF | Gating Properties | Reference |
|---|---|---|---|---|---|---|---|---|
| | | | Na$_v$1.5-5:I1606<br>Na$_v$1.5-6:I1642 | | | | Iω:none<br>ExTraFol:none | |
| 377 | Na$_v$1.5 | G1661R | Na$_v$1.5:G1661<br>Na$_v$1.5-2:G1660<br>Na$_v$1.5-3:G1628<br>Na$_v$1.5-4:G1661<br>Na$_v$1.5-5:G1607<br>Na$_v$1.5-6:G1643 | G1859R | BRGDA1 | LOF | Imax:↓<br>τrec:none<br>Ip:none<br>V1/2act:none<br>V1/2inact:none<br>Iω:none<br>ExTraFol:none | (Glazer et al., 2020) |
| 378 | Na$_v$1.5 | V1667D | Na$_v$1.5:V1667<br>Na$_v$1.5-2:V1666<br>Na$_v$1.5-3:V1634<br>Na$_v$1.5-4:V1667<br>Na$_v$1.5-5:V1613<br>Na$_v$1.5-6:V1649 | V1865D | BRGDA1 | LOF | Imax:↓<br>τrec:↓<br>Ip:none<br>V1/2act:none<br>V1/2inact:none<br>Iω:none<br>ExTraFol:none | (Monasky et al., 2021) |
| 379 | Na$_v$1.5 | T1709M | Na$_v$1.5:T1709<br>Na$_v$1.5-2:T1708<br>Na$_v$1.5-3:T1676<br>Na$_v$1.5-4:T1709<br>Na$_v$1.5-5:T1655<br>Na$_v$1.5-6:T1691 | T1907M | BRGDA1 | LOF | Imax:↓<br>τrec:↑<br>Ip:none<br>V1/2act:none<br>V1/2inact:none<br>Iω:none<br>ExTraFol:none | (Glazer et al., 2020) |
| 380 | Na$_v$1.5 | S1710L | Na$_v$1.5:S1710<br>Na$_v$1.5-2:S1709<br>Na$_v$1.5-3:S1677<br>Na$_v$1.5-4:S1710<br>Na$_v$1.5-5:S1656<br>Na$_v$1.5-6:S1692 | S1908L | VF1*<br>IVF<br>BRGDA1 | LOF | Imax:↓<br>τrec:none<br>Ip:none<br>V1/2act:→<br>V1/2inact:←<br>Iω:none<br>ExTraFol:none | (Shirai et al., 2002)<br>(Akai et al., 2000) |
| 381 | Na$_v$1.5 | G1712S | Na$_v$1.5:G1712<br>Na$_v$1.5-2:G1711<br>Na$_v$1.5-3:G1679<br>Na$_v$1.5-4:G1712<br>Na$_v$1.5-5:G1658<br>Na$_v$1.5-6:G1694 | G1910S | BRGDA1 | LOF | Imax:↓<br>τrec:none<br>Ip:none<br>V1/2act:none<br>V1/2inact:none<br>Iω:none<br>ExTraFol:none | (Sanner et al., 2021) |
| 382 | Na$_v$1.5 | D1714G | Na$_v$1.5:D1714<br>Na$_v$1.5-2:D1713<br>Na$_v$1.5-3:D1681<br>Na$_v$1.5-4:D1714 | D1912G | BRGDA1 | LOF | Imax:↓<br>τrec:none<br>Ip:none<br>V1/2act:none<br>V1/2inact:none | (Amin et al., 2005) |

| | Protein | Mutation | Isoform | Mutation (MSA) | Phenotype | GoF/LoF | Gating Properties | Reference |
|---|---|---|---|---|---|---|---|---|
| | | | $Na_v$1.5-5:D1660<br>$Na_v$1.5-6:D1696 | | | | Iω:none<br>ExTraFol:none | |
| 383 | $Na_v$1.5 | N1722D | $Na_v$1.5:N1722<br>$Na_v$1.5-2:N1721<br>$Na_v$1.5-3:N1689<br>$Na_v$1.5-4:N1722<br>$Na_v$1.5-5:N1668<br>$Na_v$1.5-6:N1704 | N1920D | BRGDA1 | LOF | Imax:↓<br>τrec:↑<br>Ip:none<br>V1/2act:→<br>V1/2inact:→<br>Iω:none<br>ExTraFol:none | (Glazer et al., 2020) |
| 384 | $Na_v$1.5 | P1730H | $Na_v$1.5:P1730<br>$Na_v$1.5-2:P1729<br>$Na_v$1.5-3:P1697<br>$Na_v$1.5-4:P1730<br>$Na_v$1.5-5:P1676<br>$Na_v$1.5-6:P1712 | P1928H | BRGDA1 | LOF | Imax:↓<br>τrec:↑<br>Ip:none<br>V1/2act:→<br>V1/2inact:→<br>Iω:none<br>ExTraFol:none | (Glazer et al., 2020) |
| 385 | $Na_v$1.5 | G1740R | $Na_v$1.5:G1740<br>$Na_v$1.5-2:G1739<br>$Na_v$1.5-3:G1707<br>$Na_v$1.5-4:G1740<br>$Na_v$1.5-5:G1686<br>$Na_v$1.5-6:G1722 | G1939R | BRGDA1 | LOF | Imax:↓<br>τrec:none<br>Ip:none<br>V1/2act:none<br>V1/2inact:none<br>Iω:none<br>ExTraFol:none | (Baroudi et al., 2004) |
| 386 | $Na_v$1.5 | G1743R | $Na_v$1.5:G1743<br>$Na_v$1.5-2:G1742<br>$Na_v$1.5-3:G1710<br>$Na_v$1.5-4:G1743<br>$Na_v$1.5-5:G1689<br>$Na_v$1.5-6:G1725 | G1942R | BRGDA1 | LOF | Imax:↓<br>τrec:none<br>Ip:none<br>V1/2act:←<br>V1/2inact:none<br>Iω:none<br>ExTraFol:none | (Valdivia et al., 2004)<br>(Clatot et al., 2020) |
| 387 | $Na_v$1.5 | V1763M | $Na_v$1.5:V1763<br>$Na_v$1.5-2:V1762<br>$Na_v$1.5-3:V1730<br>$Na_v$1.5-4:V1763<br>$Na_v$1.5-5:V1709<br>$Na_v$1.5-6:V1745 | V1962M | RWS*<br>SIDS*<br>LQT3 | GOF | Imax:none<br>τrec:none<br>Ip:↑<br>V1/2act:←<br>V1/2inact:→<br>Iω:none<br>ExTraFol:none | (Chang et al., 2004) |
| 388 | $Na_v$1.5 | M1766L | $Na_v$1.5:M1766<br>$Na_v$1.5-2:M1765<br>$Na_v$1.5-3:M1733<br>$Na_v$1.5-4:M1766 | M1965L | RWS*<br>LQT3 | GOF | Imax:none<br>τrec:none<br>Ip:↑<br>V1/2act:none<br>V1/2inact:→ | (Valdivia et al., 2002) |

| | Protein | Mutation | Isoform | Mutation (MSA) | Phenotype | GoF/LoF | Gating Properties | Reference |
|---|---|---|---|---|---|---|---|---|
| | | | Nav1.5-5:M1712<br>Nav1.5-6:M1748 | | | | Iω:none<br>ExTraFol:none | |
| 389 | Nav1.5 | Y1767C | Nav1.5:Y1767<br>Nav1.5-2:Y1766<br>Nav1.5-3:Y1734<br>Nav1.5-4:Y1767<br>Nav1.5-5:Y1713<br>Nav1.5-6:Y1749 | Y1966C | RWS*<br>LQT3 | GOF | Imax:none<br>τrec:↑<br>Ip:↑<br>V1/2act:none<br>V1/2inact:none<br>Iω:none<br>ExTraFol:none | (H. Huang et al., 2011) |
| 390 | Nav1.5 | I1768V | Nav1.5:I1768<br>Nav1.5-2:I1767<br>Nav1.5-3:I1735<br>Nav1.5-4:I1768<br>Nav1.5-5:I1714<br>Nav1.5-6:I1750 | I1967V | RWS*<br>LQT3 | GOF | Imax:none<br>τrec:none<br>Ip:none<br>V1/2act:←<br>V1/2inact:→<br>Iω:none<br>ExTraFol:none | (Rivolta et al., 2002) |
| 391 | Nav1.5 | N1774D | Nav1.5:N1774<br>Nav1.5-2:N1773<br>Nav1.5-3:N1741<br>Nav1.5-4:N1774<br>Nav1.5-5:N1720<br>Nav1.5-6:N1756 | N1973D | RWS*<br>LQT3 | GOF | Imax:↑<br>τrec:none<br>Ip:↑<br>V1/2act:←<br>V1/2inact:none<br>Iω:none<br>ExTraFol:none | (Kato et al., 2014)<br>(Hirose et al., 2020) |
| 392 | Nav1.5 | V1777M | Nav1.5:V1777<br>Nav1.5-2:V1776<br>Nav1.5-3:V1744<br>Nav1.5-4:V1777<br>Nav1.5-5:V1723<br>Nav1.5-6:V1759 | V1976M | LQT3 | GOF | Imax:none<br>τrec:none<br>Ip:↑<br>V1/2act:←<br>V1/2inact:←<br>Iω:none<br>ExTraFol:none | (Lupoglazoff et al., 2001) |
| 393 | Nav1.5 | E1784K | Nav1.5:E1784<br>Nav1.5-2:E1783<br>Nav1.5-3:E1751<br>Nav1.5-4:E1784<br>Nav1.5-5:E1730<br>Nav1.5-6:E1766 | E1983K | RWS*<br>LQT3<br>BRGDA1 | MIX | Imax:↓<br>τrec:none<br>Ip:↑<br>V1/2act:→<br>V1/2inact:←<br>Iω:none<br>ExTraFol:none | (Makita et al., 2008)<br>(Veltmann et al., 2016)<br>(Comollo et al., 2022)<br>(Wei et al., 1999) |
| 394 | Nav1.5 | L1786Q | Nav1.5:L1786<br>Nav1.5-2:L1785<br>Nav1.5-3:L1753<br>Nav1.5-4:L1786 | L1985Q | RWS*<br>BRGDA1 | | Imax:↓<br>τrec:none<br>Ip:↑<br>V1/2act:→<br>V1/2inact:← | (Kanters et al., 2014) |

| | Protein | Mutation | Isoform | Mutation (MSA) | Phenotype | GoF/LoF | Gating Properties | Reference |
|---|---|---|---|---|---|---|---|---|
| | | | $Na_v$1.5-5:L1732<br>$Na_v$1.5-6:L1768 | | | | Iω:none<br>ExTraFol:none | |
| 395 | $Na_v$1.5 | D1790G | $Na_v$1.5:D1790<br>$Na_v$1.5-2:D1789<br>$Na_v$1.5-3:D1757<br>$Na_v$1.5-4:D1790<br>$Na_v$1.5-5:D1736<br>$Na_v$1.5-6:D1772 | D1989G | RWS*<br>LQT3<br>BRGDA1 | MIX | Imax:↑<br>τrec:none<br>Ip:↑<br>V1/2act:→<br>V1/2inact:←<br>Iω:none<br>ExTraFol:none | (Blich et al., 2015)<br>(Baroudi & Chahine, 2000) |
| 396 | $Na_v$1.5 | Y1795C | $Na_v$1.5:Y1795<br>$Na_v$1.5-2:Y1794<br>$Na_v$1.5-3:Y1762<br>$Na_v$1.5-4:Y1795<br>$Na_v$1.5-5:Y1741<br>$Na_v$1.5-6:Y1777 | Y1994C | RWS*<br>LQT3 | GOF | Imax:none<br>τrec:none<br>Ip:↑<br>V1/2act:none<br>V1/2inact:none<br>Iω:none<br>ExTraFol:none | (Rivolta et al., 2001)<br>(Clancy et al., 2002) |
| 397 | $Na_v$1.5 | Y1795H | $Na_v$1.5:Y1795<br>$Na_v$1.5-2:Y1794<br>$Na_v$1.5-3:Y1762<br>$Na_v$1.5-4:Y1795<br>$Na_v$1.5-5:Y1741<br>$Na_v$1.5-6:Y1777 | Y1994H | BRGDA1 | LOF | Imax:↓<br>τrec:none<br>Ip:none<br>V1/2act:none<br>V1/2inact:←<br>Iω:none<br>ExTraFol:none | (Rivolta et al., 2001) |
| 398 | $Na_v$1.5 | D1819N | $Na_v$1.5:D1819<br>$Na_v$1.5-2:D1818<br>$Na_v$1.5-3:D1786<br>$Na_v$1.5-4:D1819<br>$Na_v$1.5-5:D1765<br>$Na_v$1.5-6:D1801 | D2018N | LQT3 | GOF | Imax:none<br>τrec:none<br>Ip:↑<br>V1/2act:none<br>V1/2inact:←<br>Iω:none<br>ExTraFol:none | (Olesen et al., 2012) |
| 399 | $Na_v$1.5 | L1825P | $Na_v$1.5:L1825<br>$Na_v$1.5-2:L1824<br>$Na_v$1.5-3:L1792<br>$Na_v$1.5-4:L1825<br>$Na_v$1.5-5:L1771<br>$Na_v$1.5-6:L1807 | L2024P | LQT3 | GOF | Imax:↓<br>τrec:none<br>Ip:↑<br>V1/2act:none<br>V1/2inact:←<br>Iω:none<br>ExTraFol:none | (K. Liu et al., 2005) |
| 400 | $Na_v$1.5 | R1826H | $Na_v$1.5:R1826<br>$Na_v$1.5-2:R1825<br>$Na_v$1.5-3:R1793<br>$Na_v$1.5-4:R1826 | R2025H | LQT3 | GOF | Imax:none<br>τrec:none<br>Ip:↑<br>V1/2act:none<br>V1/2inact:none | (Ackerman et al., 2001) |

| | Protein | Mutation | Isoform | Mutation (MSA) | Phenotype | GoF/LoF | Gating Properties | Reference |
|---|---|---|---|---|---|---|---|---|
| | | | Na$_v$1.5-5:R1772<br>Na$_v$1.5-6:R1808 | | | | Iω:none<br>ExTraFol:none | |
| 401 | Na$_v$1.5 | C1850S | Na$_v$1.5:C1850<br>Na$_v$1.5-2:C1849<br>Na$_v$1.5-3:C1817<br>Na$_v$1.5-4:C1850<br>Na$_v$1.5-5:C1796<br>Na$_v$1.5-6:C1832 | C2049S | BRGDA1 | LOF | Imax:↓<br>τrec:none<br>Ip:none<br>V1/2act:none<br>V1/2inact:←<br>Iω:none<br>ExTraFol:none | (Petitprez, Jespersen, et al., 2008) |
| 402 | Na$_v$1.5 | M1851V | Na$_v$1.5:M1851<br>Na$_v$1.5-2:M1850<br>Na$_v$1.5-3:M1818<br>Na$_v$1.5-4:M1851<br>Na$_v$1.5-5:M1797<br>Na$_v$1.5-6:M1833 | M2050V | AF | GOF | Imax:none<br>τrec:↑<br>Ip:none<br>V1/2act:←<br>V1/2inact:→<br>Iω:none<br>ExTraFol:none | (Lieve et al., 2017) |
| 403 | Na$_v$1.5 | T1858I | Na$_v$1.5:T1858<br>Na$_v$1.5-2:T1857<br>Na$_v$1.5-3:T1825<br>Na$_v$1.5-4:T1858<br>Na$_v$1.5-5:T1804<br>Na$_v$1.5-6:T1840 | T2057I | SCD | GOF | Imax:↓<br>τrec:↓<br>Ip:none<br>V1/2act:→<br>V1/2inact:→<br>Iω:none<br>ExTraFol:none | (Ghovanloo et al., 2020) |
| 404 | Na$_v$1.5 | R1897W | Na$_v$1.5:R1897<br>Na$_v$1.5-2:R1896<br>Na$_v$1.5-3:R1864<br>Na$_v$1.5-4:R1897<br>Na$_v$1.5-5:R1843<br>Na$_v$1.5-6:R1879 | R2096W | LQT3*<br>AF | | Imax:↓<br>τrec:none<br>Ip:none<br>V1/2act:none<br>V1/2inact:←<br>Iω:none<br>ExTraFol:none | (Olesen et al., 2012) |
| 405 | Na$_v$1.5 | R1898C | Na$_v$1.5:R1898<br>Na$_v$1.5-2:R1897<br>Na$_v$1.5-3:R1865<br>Na$_v$1.5-4:R1898<br>Na$_v$1.5-5:R1844<br>Na$_v$1.5-6:R1880 | R2097C | BRGDA1 | LOF | Imax:↓<br>τrec:none<br>Ip:none<br>V1/2act:none<br>V1/2inact:none<br>Iω:none<br>ExTraFol:none | (Glazer et al., 2020) |
| 406 | Na$_v$1.5 | R1898H | Na$_v$1.5:R1898<br>Na$_v$1.5-2:R1897<br>Na$_v$1.5-3:R1865<br>Na$_v$1.5-4:R1898 | R2097H | LQT3*<br>ARVD | LOF | Imax:↓<br>τrec:none<br>Ip:none<br>V1/2act:none<br>V1/2inact:none | (Te Riele et al., 2017) |

| | Protein | Mutation | Isoform | Mutation (MSA) | Phenotype | GoF/LoF | Gating Properties | Reference |
|---|---|---|---|---|---|---|---|---|
| | | | $Na_v$1.5-5:R1844<br>$Na_v$1.5-6:R1880 | | | | Iω:none<br>ExTraFol:none | |
| 407 | $Na_v$1.5 | A1905G | $Na_v$1.5:A1905<br>$Na_v$1.5-2:A1904<br>$Na_v$1.5-3:A1872<br>$Na_v$1.5-4:A1905<br>$Na_v$1.5-5:A1851<br>$Na_v$1.5-6:A1887 | A2104G | Tachycardia | | Imax:↓<br>τrec:none<br>Ip:none<br>V1/2act:→<br>V1/2inact:←<br>Iω:none<br>ExTraFol:none | (Roberts et al., 2017) |
| 408 | $Na_v$1.5 | Q1909R | $Na_v$1.5:Q1909<br>$Na_v$1.5-2:Q1908<br>$Na_v$1.5-3:Q1876<br>$Na_v$1.5-4:Q1909<br>$Na_v$1.5-5:Q1855<br>$Na_v$1.5-6:Q1891 | Q2108R | LQT3*<br>SIDS | Mixed | Imax:none<br>τrec:↓<br>Ip:none<br>V1/2act:←<br>V1/2inact:none<br>Iω:none<br>ExTraFol:none | (Winkel et al., 2015)<br>(Abdelsayed et al., 2017) |
| 409 | $Na_v$1.5 | V1951M | $Na_v$1.5:V1951<br>$Na_v$1.5-2:V1950<br>$Na_v$1.5-3:V1918<br>$Na_v$1.5-4:V1951<br>$Na_v$1.5-5:V1897<br>$Na_v$1.5-6:V1933 | V2150M | ATFB10*<br>SSS1*<br>LQT3*<br>AF | | Imax:↑<br>τrec:none<br>Ip:none<br>V1/2act:none<br>V1/2inact:→<br>Iω:none<br>ExTraFol:none | (Olesen et al., 2012) |
| 410 | $Na_v$1.5 | V1951L | $Na_v$1.5:V1951<br>$Na_v$1.5-2:V1950<br>$Na_v$1.5-3:V1918<br>$Na_v$1.5-4:V1951<br>$Na_v$1.5-5:V1897<br>$Na_v$1.5-6:V1933 | V2150L | SSS1*<br>LQT3<br>BRGDA1 | MIX | Imax:↑<br>τrec:none<br>Ip:↑<br>V1/2act:←<br>V1/2inact:none<br>Iω:none<br>ExTraFol:none | (D. W. Wang et al., 2007) |
| 411 | $Na_v$1.5 | I1968S | $Na_v$1.5:I1968<br>$Na_v$1.5-2:I1967<br>$Na_v$1.5-3:I1935<br>$Na_v$1.5-4:I1968<br>$Na_v$1.5-5:I1914<br>$Na_v$1.5-6:I1950 | I2190S | BRGDA1 | LOF | Imax:↓<br>τrec:none<br>Ip:none<br>V1/2act:none<br>V1/2inact:none<br>Iω:none<br>ExTraFol:none | (Frustaci et al., 2005) |
| 412 | $Na_v$1.5 | F2004L | $Na_v$1.5:F2004<br>$Na_v$1.5-2:F2003<br>$Na_v$1.5-3:F1971<br>$Na_v$1.5-4:F2004 | F2229L | LQT3*<br>BRGDA1 | LOF | Imax:↓<br>τrec:none<br>Ip:none<br>V1/2act:→<br>V1/2inact:← | (Bébarová et al., 2008) |

| | Protein | Mutation | Isoform | Mutation (MSA) | Phenotype | GoF/LoF | Gating Properties | Reference |
|---|---|---|---|---|---|---|---|---|
| | | | $Na_v$1.5-5:F1950<br>$Na_v$1.5-6:F1986 | | | | Iω:none<br>ExTraFol:none | |
| 413 | $Na_v$1.5 | F2004L | $Na_v$1.5:F2004<br>$Na_v$1.5-2:F2003<br>$Na_v$1.5-3:F1971<br>$Na_v$1.5-4:F2004<br>$Na_v$1.5-5:F1950<br>$Na_v$1.5-6:F1986 | F2229L | BRGDA1*<br>LQT3 | | Imax:none<br>τrec:↑<br>Ip:↑<br>V1/2act:none<br>V1/2inact:→<br>Iω:none<br>ExTraFol:none | (Olesen et al., 2012)<br>(D. W. Wang et al., 2007) |
| 414 | $Na_v$1.5 | P2006A | $Na_v$1.5:P2006<br>$Na_v$1.5-2:P2005<br>$Na_v$1.5-3:P1973<br>$Na_v$1.5-4:P2006<br>$Na_v$1.5-5:P1952<br>$Na_v$1.5-6:P1988 | P2231A | BRGDA1*<br>LQT3 | GOF | Imax:none<br>τrec:none<br>Ip:↑<br>V1/2act:none<br>V1/2inact:none<br>Iω:none<br>ExTraFol:none | (D. W. Wang et al., 2007) |
| 415 | $Na_v$1.5 | R2012H | $Na_v$1.5:R2012<br>$Na_v$1.5-2:R2011<br>$Na_v$1.5-3:R1979<br>$Na_v$1.5-4:R2012<br>$Na_v$1.5-5:R1958<br>$Na_v$1.5-6:R1994 | R2237H | BRGDA1 | LOF | Imax:↓<br>τrec:none<br>Ip:none<br>V1/2act:none<br>V1/2inact:←<br>Iω:none<br>ExTraFol:none | (Ortiz-Bonnin et al., 2016) |
| 416 | $Na_v$1.5 | V2016M | $Na_v$1.5:V2016<br>$Na_v$1.5-2:V2015<br>$Na_v$1.5-3:V1983<br>$Na_v$1.5-4:V2016<br>$Na_v$1.5-5:V1962<br>$Na_v$1.5-6:V1998 | V2241M | LQT3 | GOF | Imax:↓<br>τrec:none<br>Ip:↑<br>V1/2act:none<br>V1/2inact:none<br>Iω:none<br>ExTraFol:none | (J. Chen et al., 2016) |
| 417 | $Na_v$1.6 | E1100A | - | 0 | DEE13 | | Imax:↓<br>τrec:none<br>Ip:none<br>V1/2act:none<br>V1/2inact:none<br>Iω:none<br>ExTraFol:none | (Gasser et al., 2012) |
| 418 | $Na_v$1.6 | M139I | $Na_v$1.6:M139<br>$Na_v$1.6-2:M139<br>$Na_v$1.6-3:M139<br>$Na_v$1.6-4:M139<br>$Na_v$1.6-5:M139 | M142I | EIEE*<br>ID | | Imax:none<br>τrec:none<br>Ip:↑<br>V1/2act:←<br>V1/2inact:none | (Zaman et al., 2019) |

| | Protein | Mutation | Isoform | Mutation (MSA) | Phenotype | GoF/LoF | Gating Properties | Reference |
|---|---|---|---|---|---|---|---|---|
| | | | | | | | Iω:none<br>ExTraFol:none | |
| 419 | Nav1.6 | R223G | Nav1.6:R223<br>Nav1.6-2:R223<br>Nav1.6-3:R223<br>Nav1.6-4:R223<br>Nav1.6-5:R223 | R231G | DEE13*<br>DD<br>DEE | GOF | Imax:↓<br>τrec:↑<br>Ip:none<br>V1/2act:←<br>V1/2inact:←<br>Iω:none<br>ExTraFol:none | (Y. Liu et al., 2021)<br>(de Kovel et al., 2014) |
| 420 | Nav1.6 | G269R | Nav1.6:G269<br>Nav1.6-2:G269<br>Nav1.6-3:G269<br>Nav1.6-4:G269<br>Nav1.6-5:G269 | G277R | DEE13 | | Imax:↓<br>τrec:none<br>Ip:none<br>V1/2act:none<br>V1/2inact:none<br>Iω:none<br>ExTraFol:none | (Wengert, Tronhjem, et al., 2019) |
| 421 | Nav1.6 | N374K | Nav1.6:N374<br>Nav1.6-2:N374<br>Nav1.6-3:N374<br>Nav1.6-4:N374<br>Nav1.6-5:N374 | N443K | EPI | | Imax:none<br>τrec:none<br>Ip:↑<br>V1/2act:←<br>V1/2inact:none<br>Iω:none<br>ExTraFol:none | (Zaman et al., 2019) |
| 422 | Nav1.6 | R639C | Nav1.6:R639<br>Nav1.6-2:R639<br>Nav1.6-3:R639<br>Nav1.6-4:R639<br>Nav1.6-5:R639 | R716C | EIEE | | Imax:↑<br>τrec:none<br>Ip:none<br>V1/2act:none<br>V1/2inact:none<br>Iω:none<br>ExTraFol:none | (Zybura et al., 2022) |
| 423 | Nav1.6 | T767I | Nav1.6:T767<br>Nav1.6-2:T767<br>Nav1.6-3:T778<br>Nav1.6-4:T767<br>Nav1.6-5:T767 | T862I | EIEE*<br>DEE13 | GOF | Imax:↓<br>τrec:none<br>Ip:↑<br>V1/2act:←<br>V1/2inact:none<br>Iω:none<br>ExTraFol:none | (Pan & Cummins, 2020) |
| 424 | Nav1.6 | G822R | Nav1.6:G822<br>Nav1.6-2:G822<br>Nav1.6-3:G833<br>Nav1.6-4:G822<br>Nav1.6-5:G822 | G917R | DEE13 | | Imax:↓<br>τrec:none<br>Ip:none<br>V1/2act:none<br>V1/2inact:none | (Wengert, Tronhjem, et al., 2019) |

| | Protein | Mutation | Isoform | Mutation (MSA) | Phenotype | GoF/LoF | Gating Properties | Reference |
|---|---|---|---|---|---|---|---|---|
| | | | | | | | Iω:none<br>ExTraFol:none | |
| 425 | Nav1.6 | R850Q | Nav1.6:R850<br>Nav1.6-2:R850<br>Nav1.6-3:R861<br>Nav1.6-4:R850<br>Nav1.6-5:R850 | R947Q | DEE13 | GOF | Imax:none<br>τrec:none<br>Ip:↑<br>V1/2act:←<br>V1/2inact:none<br>Iω:none<br>ExTraFol:none | (Pan & Cummins, 2020) |
| 426 | Nav1.6 | G964R | Nav1.6:G964<br>Nav1.6-2:G964<br>Nav1.6-3:G975<br>Nav1.6-4:G964<br>Nav1.6-5:G964 | G1070R | NDD | LOF | Imax:↓<br>τrec:none<br>Ip:none<br>V1/2act:none<br>V1/2inact:none<br>Iω:none<br>ExTraFol:none | (Wagnon et al., 2017) |
| 427 | Nav1.6 | N984K | Nav1.6:N984<br>Nav1.6-2:N984<br>Nav1.6-3:N995<br>Nav1.6-4:N984<br>Nav1.6-5:N984 | N1090K | DEE13*<br>EE<br>ID | GOF | Imax:none<br>τrec:none<br>Ip:none<br>V1/2act:←<br>V1/2inact:none<br>Iω:none<br>ExTraFol:none | (Blanchard et al., 2015) |
| 428 | Nav1.6 | E1218K | Nav1.6:E1218<br>Nav1.6-2:E1218<br>Nav1.6-3:E1229<br>Nav1.6-4:E1218<br>Nav1.6-5:E1218 | E1416K | ID | LOF | Imax:none<br>τrec:none<br>Ip:none<br>V1/2act:none<br>V1/2inact:none<br>Iω:none<br>ExTraFol:↓ | (Wagnon et al., 2017) |
| 429 | Nav1.6 | I1327V | Nav1.6:I1327<br>Nav1.6-2:I1327<br>Nav1.6-3:I1338<br>Nav1.6-4:--<br>Nav1.6-5:I1286 | I1525V | DEE13* | | Imax:none<br>τrec:none<br>Ip:none<br>V1/2act:←<br>V1/2inact:→<br>Iω:none<br>ExTraFol:none | (Barker et al., 2016) |
| 430 | Nav1.6 | T1360N | Nav1.6:T1360<br>Nav1.6-2:T1360<br>Nav1.6-3:T1371<br>Nav1.6-4:--<br>Nav1.6-5:T1319 | T1558N | DEE13 | | Imax:none<br>τrec:none<br>Ip:none<br>V1/2act:none<br>V1/2inact:← | (Wengert, Tronhjem, et al., 2019) |

| | Protein | Mutation | Isoform | Mutation (MSA) | Phenotype | GoF/LoF | Gating Properties | Reference |
|---|---|---|---|---|---|---|---|---|
| | | | | | | | Iω:none<br>ExTraFol:none | |
| 431 | Na$_v$1.6 | R1628C | Na$_v$1.6:R1617<br>Na$_v$1.6-2:R1617<br>Na$_v$1.6-3:R1628<br>Na$_v$1.6-4:--<br>Na$_v$1.6-5:R1576 | R1617C | DEE13 | | Imax:none<br>τrec:none<br>Ip:none<br>V1/2act:→<br>V1/2inact:none<br>Iω:none<br>ExTraFol:none | (Wengert, Tronhjem, et al., 2019) |
| 432 | Na$_v$1.6 | G1451S | Na$_v$1.6:G1451<br>Na$_v$1.6-2:G1451<br>Na$_v$1.6-3:G1462<br>Na$_v$1.6-4:--<br>Na$_v$1.6-5:G1410 | G1651S | DEE13*<br>EE<br>ID | LOF | Imax:↓<br>τrec:none<br>Ip:none<br>V1/2act:none<br>V1/2inact:none<br>Iω:none<br>ExTraFol:none | (Blanchard et al., 2015) |
| 433 | Na$_v$1.6 | G1475R | Na$_v$1.6:G1475<br>Na$_v$1.6-2:G1475<br>Na$_v$1.6-3:G1486<br>Na$_v$1.6-4:--<br>Na$_v$1.6-5:G1434 | G1675R | EIEE*<br>DEE13 | GOF | Imax:none<br>τrec:none<br>Ip:↑<br>V1/2act:←<br>V1/2inact:→<br>Iω:none<br>ExTraFol:none | (Y. Liu et al., 2019)<br>(Zaman et al., 2019) |
| 434 | Na$_v$1.6 | A1491V | Na$_v$1.6:A1491<br>Na$_v$1.6-2:A1491<br>Na$_v$1.6-3:A1502<br>Na$_v$1.6-4:--<br>Na$_v$1.6-5:A1450 | A1691V | EPI | | Imax:none<br>τrec:none<br>Ip:↑<br>V1/2act:←<br>V1/2inact:none<br>Iω:none<br>ExTraFol:none | (Zaman et al., 2019) |
| 435 | Na$_v$1.6 | R1617Q | Na$_v$1.6:R1617<br>Na$_v$1.6-2:R1617<br>Na$_v$1.6-3:R1628<br>Na$_v$1.6-4:--<br>Na$_v$1.6-5:R1576 | R1821Q | DEE13 | GOF | Imax:none<br>τrec:none<br>Ip:↑<br>V1/2act:←<br>V1/2inact:→<br>Iω:none<br>ExTraFol:none | (Wagnon et al., 2016) |
| 436 | Na$_v$1.6 | R1620L | Na$_v$1.6:R1620<br>Na$_v$1.6-2:R1620<br>Na$_v$1.6-3:R1631<br>Na$_v$1.6-4:--<br>Na$_v$1.6-5:R1579 | R1824L | ID | LOF | Imax:↓<br>τrec:↑<br>Ip:none<br>V1/2act:none<br>V1/2inact:← | (Y. Liu et al., 2019) |

| | Protein | Mutation | Isoform | Mutation (MSA) | Phenotype | GoF/LoF | Gating Properties | Reference |
|---|---|---|---|---|---|---|---|---|
| | | | | | | | Iω:none<br>ExTraFol:none | |
| 437 | $Na_v1.6$ | A1622D | $Na_v1.6$:A1622<br>$Na_v1.6$-2:A1622<br>$Na_v1.6$-3:A1633<br>$Na_v1.6$-4:--<br>$Na_v1.6$-5:A1581 | A1826D | ID | GOF | Imax:none<br>τrec:↑<br>Ip:↑<br>V1/2act:→<br>V1/2inact:→<br>Iω:none<br>ExTraFol:none | (Y. Liu et al., 2019) |
| 438 | $Na_v1.6$ | R1638C | $Na_v1.6$:R1638<br>$Na_v1.6$-2:R1638<br>$Na_v1.6$-3:R1649<br>$Na_v1.6$-4:--<br>$Na_v1.6$-5:R1597 | R1842C | NDD | LOF | Imax:none<br>τrec:none<br>Ip:none<br>V1/2act:→<br>V1/2inact:none<br>Iω:none<br>ExTraFol:none | (Wengert, Tronhjem, et al., 2019) |
| 439 | $Na_v1.6$ | M1760I | $Na_v1.6$:M1760<br>$Na_v1.6$-2:M1760<br>$Na_v1.6$-3:M1771<br>$Na_v1.6$-4:--<br>$Na_v1.6$-5:M1719 | M1965I | EIEE*<br>DEE13 | GOF | Imax:none<br>τrec:none<br>Ip:none<br>V1/2act:←<br>V1/2inact:none<br>Iω:none<br>ExTraFol:none | (Y. Liu et al., 2019) |
| 440 | $Na_v1.6$ | N1768D | $Na_v1.6$:N1768<br>$Na_v1.6$-2:N1768<br>$Na_v1.6$-3:N1779<br>$Na_v1.6$-4:--<br>$Na_v1.6$-5:N1727 | N1973D | DEE13 | GOF | Imax:↓<br>τrec:none<br>Ip:↑<br>V1/2act:none<br>V1/2inact:→<br>Iω:none<br>ExTraFol:none | (Veeramah et al., 2012)<br>(Wengert, Saga, et al., 2019)<br>(Patel et al., 2016)<br>(Baker et al., 2018) |
| 441 | $Na_v1.6$ | R1872L | $Na_v1.6$:R1872<br>$Na_v1.6$-2:R1872<br>$Na_v1.6$-3:R1883<br>$Na_v1.6$-4:--<br>$Na_v1.6$-5:R1831 | R2077L | DEE13*<br>EIEE | | Imax:none<br>τrec:none<br>Ip:↑<br>V1/2act:←<br>V1/2inact:none<br>Iω:none<br>ExTraFol:none | (Baker et al., 2018)<br>(Y. Liu et al., 2019) |
| 442 | $Na_v1.6$ | R1872Q | $Na_v1.6$:R1872<br>$Na_v1.6$-2:R1872<br>$Na_v1.6$-3:R1883 | R2077Q | DEE13 | GOF | Imax:↑<br>τrec:none<br>Ip:↑<br>V1/2act:←<br>V1/2inact:→ | (Pan & Cummins, 2020)<br>(Wagnon et al., 2016) |

| | Protein | Mutation | Isoform | Mutation (MSA) | Phenotype | GoF/LoF | Gating Properties | Reference |
|---|---|---|---|---|---|---|---|---|
| | | | Nav1.6-4:--<br>Nav1.6-5:R1831 | | | | Iω:none<br>ExTraFol:none | (Atkin et al., 2018) |
| 443 | Nav1.6 | R1872W | Nav1.6:R1872<br>Nav1.6-2:R1872<br>Nav1.6-3:R1883<br>Nav1.6-4:--<br>Nav1.6-5:R1831 | R2077W | DEE13 | GOF | Imax:none<br>τrec:none<br>Ip:↑<br>V1/2act:←<br>V1/2inact:none<br>Iω:none<br>ExTraFol:none | (Bunton-Stasyshyn et al., 2019)<br>(Y. Liu et al., 2019)<br>(Zaman et al., 2019) |
| 444 | Nav1.7 | R1150W | - | 0 | IEM | | Imax:none<br>τrec:none<br>Ip:none<br>V1/2act:→<br>V1/2inact:none<br>Iω:none<br>ExTraFol:none | (Estacion et al., 2009) |
| 445 | Nav1.7 | Q10R | Nav1.7:Q10<br>Nav1.7-2:Q10<br>Nav1.7-3:Q10<br>Nav1.7-4:Q10 | Q15R | PEXPD*<br>PERYTHM*<br>PEM | GOF | Imax:↑<br>τrec:none<br>Ip:none<br>V1/2act:←<br>V1/2inact:none<br>Iω:none<br>ExTraFol:none | (C. Han et al., 2009) |
| 446 | Nav1.7 | R99H | Nav1.7:R99<br>Nav1.7-2:R99<br>Nav1.7-3:R99<br>Nav1.7-4:R99 | R108H | CIP | LOF | Imax:↓<br>τrec:none<br>Ip:none<br>V1/2act:none<br>V1/2inact:→<br>Iω:none<br>ExTraFol:↓ | (J. Sun et al., 2020) |
| 447 | Nav1.7 | I136V | Nav1.7:I136<br>Nav1.7-2:I136<br>Nav1.7-3:I136<br>Nav1.7-4:I136 | I145V | PEM | GOF | Imax:none<br>τrec:↑<br>Ip:↑<br>V1/2act:←<br>V1/2inact:→<br>Iω:none<br>ExTraFol:none | (X. Cheng et al., 2008a)<br>(M. T. Wu et al., 2013) |
| 448 | Nav1.7 | S211P | Nav1.7:S211<br>Nav1.7-2:S211<br>Nav1.7-3:S211<br>Nav1.7-4:S211 | S225P | PEM | GOF | Imax:↑<br>τrec:none<br>Ip:none<br>V1/2act:←<br>V1/2inact:← | (Estacion, Choi, et al., 2010) |

| | Protein | Mutation | Isoform | Mutation (MSA) | Phenotype | GoF/LoF | Gating Properties | Reference |
|---|---|---|---|---|---|---|---|---|
| | | | | | | | Iω:none<br>ExTraFol:none | |
| 449 | Na$_v$1.7 | F216S | Na$_v$1.7:F216<br>Na$_v$1.7-2:F216<br>Na$_v$1.7-3:F216<br>Na$_v$1.7-4:F216 | F230S | PERYTHM*<br>PEM | GOF | Imax:↑<br>τrec:none<br>Ip:none<br>V1/2act:←<br>V1/2inact:none<br>Iω:none<br>ExTraFol:none | (Estacion, Choi, et al., 2010) |
| 450 | Na$_v$1.7 | I228M | Na$_v$1.7:I228<br>Na$_v$1.7-2:I228<br>Na$_v$1.7-3:I228<br>Na$_v$1.7-4:I228 | I242M | PEM/PEXPD/SFNP | GOF | Imax:none<br>τrec:none<br>Ip:none<br>V1/2act:none<br>V1/2inact:→<br>Iω:none<br>ExTraFol:none | (Estacion et al., 2011) |
| 451 | Na$_v$1.7 | I234T | Na$_v$1.7:I234<br>Na$_v$1.7-2:I234<br>Na$_v$1.7-3:I234<br>Na$_v$1.7-4:I234 | I248T | PEM | GOF | Imax:↑<br>τrec:none<br>Ip:none<br>V1/2act:←<br>V1/2inact:none<br>Iω:none<br>ExTraFol:none | (Ahn et al., 2010)<br>(M. T. Wu et al., 2013) |
| 452 | Na$_v$1.7 | S241T | Na$_v$1.7:S241<br>Na$_v$1.7-2:S241<br>Na$_v$1.7-3:S241<br>Na$_v$1.7-4:S241 | S255T | PERYTHM*<br>PEM | GOF | Imax:none<br>τrec:none<br>Ip:↑<br>V1/2act:←<br>V1/2inact:none<br>Iω:none<br>ExTraFol:none | (Lampert et al., 2006)<br>(Alsaloum et al., 2023) |
| 453 | Na$_v$1.7 | L245V | Na$_v$1.7:L245<br>Na$_v$1.7-2:L245<br>Na$_v$1.7-3:L245<br>Na$_v$1.7-4:L245 | L259V | IEM | | Imax:none<br>τrec:none<br>Ip:↑<br>V1/2act:none<br>V1/2inact:none<br>Iω:none<br>ExTraFol:none | (Emery et al., 2015) |
| 454 | Na$_v$1.7 | N395K | Na$_v$1.7:N395<br>Na$_v$1.7-2:N395<br>Na$_v$1.7-3:N395<br>Na$_v$1.7-4:N395 | N473K | PERYTHM*<br>PEM | GOF | Imax:none<br>τrec:none<br>Ip:none<br>V1/2act:←<br>V1/2inact:none | (Sheets et al., 2007) |

| | Protein | Mutation | Isoform | Mutation (MSA) | Phenotype | GoF/LoF | Gating Properties | Reference |
|---|---|---|---|---|---|---|---|---|
| | | | | | | | Iω:none<br>ExTraFol:none | |
| 455 | Na$_v$1.7 | V400M | Na$_v$1.7:V400<br>Na$_v$1.7-2:V400<br>Na$_v$1.7-3:V400<br>Na$_v$1.7-4:V400 | V478M | PEM | GOF | Imax:none<br>τrec:none<br>Ip:none<br>V1/2act:←<br>V1/2inact:none<br>Iω:none<br>ExTraFol:none | (Y. Yang et al., 2012)<br>(Fischer et al., 2009) |
| 456 | Na$_v$1.7 | K655R | Na$_v$1.7:K666<br>Na$_v$1.7-2:K666<br>Na$_v$1.7-3:K655<br>Na$_v$1.7-4:K655 | K666R | FS | | Imax:none<br>τrec:↑<br>Ip:↑<br>V1/2act:→<br>V1/2inact:none<br>Iω:none<br>ExTraFol:none | (S. Zhang et al., 2020) |
| 457 | Na$_v$1.7 | G616R | Na$_v$1.7:G616<br>Na$_v$1.7-2:G616<br>Na$_v$1.7-3:G616<br>Na$_v$1.7-4:G616 | G714R | PEM | GOF | Imax:↑<br>τrec:none<br>Ip:none<br>V1/2act:none<br>V1/2inact:→<br>Iω:none<br>ExTraFol:none | (Choi et al., 2010) |
| 458 | Na$_v$1.7 | D623N | Na$_v$1.7:D623<br>Na$_v$1.7-2:D623<br>Na$_v$1.7-3:D623<br>Na$_v$1.7-4:D623 | D721N | SFNP*<br>I-SFN | GOF | Imax:none<br>τrec:none<br>Ip:none<br>V1/2act:→<br>V1/2inact:→<br>Iω:none<br>ExTraFol:none | (Faber, Hoeijmakers, et al., 2012) |
| 459 | Na$_v$1.7 | I720K | Na$_v$1.7:I731<br>Na$_v$1.7-2:I731<br>Na$_v$1.7-3:I720<br>Na$_v$1.7-4:I720 | I731K | SFNP | GOF | Imax:none<br>τrec:none<br>Ip:none<br>V1/2act:none<br>V1/2inact:→<br>Iω:none<br>ExTraFol:none | (Faber, Hoeijmakers, et al., 2012) |
| 460 | Na$_v$1.7 | N641Y | Na$_v$1.7:N641<br>Na$_v$1.7-2:N641<br>Na$_v$1.7-3:N641<br>Na$_v$1.7-4:N641 | N740Y | GEFSP27*<br>FS | GOF | Imax:none<br>τrec:↑<br>Ip:↑<br>V1/2act:→<br>V1/2inact:none | (S. Zhang et al., 2020) |

| | Protein | Mutation | Isoform | Mutation (MSA) | Phenotype | GoF/LoF | Gating Properties | Reference |
|---|---|---|---|---|---|---|---|---|
| | | | | | | | Iω:none<br>ExTraFol:none | |
| 461 | Nav1.7 | I739V | Nav1.7:I750<br>Nav1.7-2:I750<br>Nav1.7-3:I739<br>Nav1.7-4:I739 | I750V | SFNP | GOF | Imax:none<br>τrec:none<br>Ip:none<br>V1/2act:←<br>V1/2inact:→<br>Iω:none<br>ExTraFol:none | (Faber, Hoeijmakers, et al., 2012) |
| 462 | Nav1.7 | F826Y | Nav1.7:F837<br>Nav1.7-2:F837<br>Nav1.7-3:F826<br>Nav1.7-4:F826 | F837Y | PEM | GOF | Imax:none<br>τrec:none<br>Ip:none<br>V1/2act:←<br>V1/2inact:→<br>Iω:none<br>ExTraFol:none | (B. Wu et al., 2017) |
| 463 | Nav1.7 | I848E | Nav1.7:I859<br>Nav1.7-2:I859<br>Nav1.7-3:I848<br>Nav1.7-4:I848 | I859E | IEM | | Imax:↓<br>τrec:none<br>Ip:none<br>V1/2act:←<br>V1/2inact:none<br>Iω:none<br>ExTraFol:none | (Kerth et al., 2021) |
| 464 | Nav1.7 | I848T | Nav1.7:I859<br>Nav1.7-2:I859<br>Nav1.7-3:I848<br>Nav1.7-4:I848 | I859T | PEM | GOF | Imax:↑<br>τrec:↑<br>Ip:↑<br>V1/2act:←<br>V1/2inact:→<br>Iω:none<br>ExTraFol:none | (M. T. Wu et al., 2013)<br>(Cummins et al., 2004)<br>(Drenth et al., 2005)<br>(Namer et al., 2015)<br>(Theile et al., 2011) |
| 465 | Nav1.7 | I848T | Nav1.7:I859<br>Nav1.7-2:I859<br>Nav1.7-3:I848<br>Nav1.7-4:I848 | I859T | IEM | GOF | Imax:↓<br>τrec:none<br>Ip:none<br>V1/2act:←<br>V1/2inact:none<br>Iω:none<br>ExTraFol:none | (Kerth et al., 2021) |
| 466 | Nav1.7 | G856D | Nav1.7:G867<br>Nav1.7-2:G867 | G867D | SFNP | GOF | Imax:none<br>τrec:none<br>Ip:↑<br>V1/2act:← | (Hoeijmakers et al., 2012) |

| | Protein | Mutation | Isoform | Mutation (MSA) | Phenotype | GoF/LoF | Gating Properties | Reference |
|---|---|---|---|---|---|---|---|---|
| | | | $Na_v1.7$-3:G856<br>$Na_v1.7$-4:G856 | | | | V1/2inact:→<br>Iω:none<br>ExTraFol:none | |
| 467 | $Na_v1.7$ | G856R | $Na_v1.7$:G867<br>$Na_v1.7$-2:G867<br>$Na_v1.7$-3:G856<br>$Na_v1.7$-4:G856 | G867R | peripheral pain | GOF | Imax:none<br>τrec:none<br>Ip:none<br>V1/2act:←<br>V1/2inact:none<br>Iω:none<br>ExTraFol:none | (Tanaka et al., 2017) |
| 468 | $Na_v1.7$ | L858F | $Na_v1.7$:L869<br>$Na_v1.7$-2:L869<br>$Na_v1.7$-3:L858<br>$Na_v1.7$-4:L858 | L869F | PEM | GOF | Imax:↑<br>τrec:none<br>Ip:none<br>V1/2act:←<br>V1/2inact:→<br>Iω:none<br>ExTraFol:none | (Drenth et al., 2005)<br>(C. Han et al., 2007) |
| 469 | $Na_v1.7$ | L858H | $Na_v1.7$:L869<br>$Na_v1.7$-2:L869<br>$Na_v1.7$-3:L858<br>$Na_v1.7$-4:L858 | L869H | PEM | GOF | Imax:↑<br>τrec:none<br>Ip:none<br>V1/2act:←<br>V1/2inact:none<br>Iω:none<br>ExTraFol:none | (Cummins et al., 2004)<br>(Y. Yang et al., 2004)<br>(Estacion & Waxman, 2017)<br>(Estacion, Waxman, et al., 2010) |
| 470 | $Na_v1.7$ | A863P | $Na_v1.7$:A874<br>$Na_v1.7$-2:A874<br>$Na_v1.7$-3:A863<br>$Na_v1.7$-4:A863 | A874P | PEM | GOF | Imax:↑<br>τrec:none<br>Ip:none<br>V1/2act:←<br>V1/2inact:→<br>Iω:none<br>ExTraFol:none | (Harty et al., 2006) |
| 471 | $Na_v1.7$ | Q875E | $Na_v1.7$:Q886<br>$Na_v1.7$-2:Q886<br>$Na_v1.7$-3:Q875<br>$Na_v1.7$-4:Q875 | Q886E | PEM | GOF | Imax:none<br>τrec:none<br>Ip:none<br>V1/2act:←<br>V1/2inact:none<br>Iω:none<br>ExTraFol:none | (Stadler et al., 2015) |
| 472 | $Na_v1.7$ | L823R | $Na_v1.7$:L823<br>$Na_v1.7$-2:L823 | L927R | PEM | GOF | Imax:none<br>τrec:none<br>Ip:none | (Lampert et al., 2009) |

| | Protein | Mutation | Isoform | Mutation (MSA) | Phenotype | GoF/LoF | Gating Properties | Reference |
|---|---|---|---|---|---|---|---|---|
| | | | $Na_v$1.7-3:L812 $Na_v$1.7-4:L812 | | | | V1/2act:← V1/2inact:← Iω:none ExTraFol:none | |
| 473 | $Na_v$1.7 | W917G | $Na_v$1.7:W928 $Na_v$1.7-2:W928 $Na_v$1.7-3:W917 $Na_v$1.7-4:W917 | W928G | CIP | LOF | Imax:↓ τrec:none Ip:none V1/2act:none V1/2inact:none Iω:none ExTraFol:none | (J. Sun et al., 2020) |
| 474 | $Na_v$1.7 | V872G | $Na_v$1.7:V872 $Na_v$1.7-2:V872 $Na_v$1.7-3:V861 $Na_v$1.7-4:V861 | V978G | PEM | GOF | Imax:↑ τrec:none Ip:none V1/2act:← V1/2inact:none Iω:none ExTraFol:none | (Choi et al., 2009) |
| 475 | $Na_v$1.7 | V991L | $Na_v$1.7:V1002 $Na_v$1.7-2:V1002 $Na_v$1.7-3:V991 $Na_v$1.7-4:V991 | V1002L | SFNP | GOF | Imax:none τrec:none Ip:none V1/2act:none V1/2inact:→ Iω:none ExTraFol:none | (Faber, Hoeijmakers, et al., 2012) |
| 476 | $Na_v$1.7 | R996C | $Na_v$1.7:R1007 $Na_v$1.7-2:R1007 $Na_v$1.7-3:R996 $Na_v$1.7-4:R996 | R1007C | PEXPD | GOF | Imax:none τrec:none Ip:↑ V1/2act:← V1/2inact:none Iω:none ExTraFol:none | (Fertleman et al., 2006) |
| 477 | $Na_v$1.7 | M932L | $Na_v$1.7:M932 $Na_v$1.7-2:M932 $Na_v$1.7-3:M921 $Na_v$1.7-4:M921 | M1045L | SFNP | GOF | Imax:none τrec:none Ip:none V1/2act:none V1/2inact:→ Iω:none ExTraFol:none | (Faber, Hoeijmakers, et al., 2012) |
| 478 | $Na_v$1.7 | C1143F | $Na_v$1.7:C1154 $Na_v$1.7-2:C1154 | C1154F | ASD | LOF | Imax:none τrec:↓ Ip:none V1/2act:none | (Rubinstein et al., 2018) |

| | Protein | Mutation | Isoform | Mutation (MSA) | Phenotype | GoF/LoF | Gating Properties | Reference |
|---|---|---|---|---|---|---|---|---|
| | | | $Na_v$1.7-3:C1143<br>$Na_v$1.7-4:C1143 | | | | V1/2inact:none<br>Iω:none<br>ExTraFol:none | |
| 479 | $Na_v$1.7 | W1150R | $Na_v$1.7:W1161<br>$Na_v$1.7-2:W1161<br>$Na_v$1.7-3:W1150<br>$Na_v$1.7-4:W1150 | W1161R | FS | | Imax:none<br>τrec:↓<br>Ip:↑<br>V1/2act:←<br>V1/2inact:none<br>Iω:none<br>ExTraFol:none | (S. Zhang et al., 2020) |
| 480 | $Na_v$1.7 | A1236E | $Na_v$1.7:A1247<br>$Na_v$1.7-2:A1247<br>$Na_v$1.7-3:A1236<br>$Na_v$1.7-4:A1236 | A1247E | CIP | | Imax:none<br>τrec:none<br>Ip:none<br>V1/2act:→<br>V1/2inact:←<br>Iω:none<br>ExTraFol:none | (Emery et al., 2015) |
| 481 | $Na_v$1.7 | R1279P | $Na_v$1.7:R1290<br>$Na_v$1.7-2:R1290<br>$Na_v$1.7-3:R1279<br>$Na_v$1.7-4:R1279 | R1290P | distal leg pain | GOF | Imax:none<br>τrec:none<br>Ip:↑<br>V1/2act:→<br>V1/2inact:→<br>Iω:none<br>ExTraFol:none | (J. Huang, Yang, et al., 2014) |
| 482 | $Na_v$1.7 | V1298D | $Na_v$1.7:V1309<br>$Na_v$1.7-2:V1309<br>$Na_v$1.7-3:V1298<br>$Na_v$1.7-4:V1298 | V1309D | PEXPD | GOF | Imax:none<br>τrec:none<br>Ip:↑<br>V1/2act:←<br>V1/2inact:none<br>Iω:none<br>ExTraFol:none | (Fertleman et al., 2006) |
| 483 | $Na_v$1.7 | V1298F | $Na_v$1.7:V1309<br>$Na_v$1.7-2:V1309<br>$Na_v$1.7-3:V1298<br>$Na_v$1.7-4:V1298 | V1309F | PEXPD | GOF | Imax:↓<br>τrec:↑<br>Ip:↑<br>V1/2act:none<br>V1/2inact:→<br>Iω:none<br>ExTraFol:none | (Fertleman et al., 2006)<br>(X. Cheng et al., 2010)<br>(Estacion, Waxman, et al., 2010) |
| 484 | $Na_v$1.7 | V1299F | $Na_v$1.7:V1310<br>$Na_v$1.7-2:V1310<br>$Na_v$1.7-3:V1299<br>$Na_v$1.7-4:V1299 | V1310F | PEXPD | | Imax:none<br>τrec:none<br>Ip:↑<br>V1/2act:none<br>V1/2inact:→ | (Jarecki et al., 2008) |

| | Protein | Mutation | Isoform | Mutation (MSA) | Phenotype | GoF/LoF | Gating Properties | Reference |
|---|---|---|---|---|---|---|---|---|
| | | | | | | | Iω:none ExTraFol:none | |
| 485 | $Na_v1.7$ | P1308L | $Na_v1.7$:P1319 $Na_v1.7$-2:P1319 $Na_v1.7$-3:P1308 $Na_v1.7$-4:P1308 | P1319L | PEM/IEM | | Imax:↓ τrec:none Ip:↑ V1/2act:← V1/2inact:none Iω:none ExTraFol:none | (X. Cheng et al., 2010) (M. T. Wu et al., 2013) |
| 486 | $Na_v1.7$ | V1316A | $Na_v1.7$:V1327 $Na_v1.7$-2:V1327 $Na_v1.7$-3:V1316 $Na_v1.7$-4:V1316 | V1327A | PEM | GOF | Imax:none τrec:none Ip:none V1/2act:← V1/2inact:→ Iω:none ExTraFol:none | (Cregg et al., 2013) (C. W. Huang et al., 2016) |
| 487 | $Na_v1.7$ | F1449W | $Na_v1.7$:F1460 $Na_v1.7$-2:F1460 $Na_v1.7$-3:F1449 $Na_v1.7$-4:F1449 | F1460W | erythromelalgia | | Imax:↓ τrec:none Ip:none V1/2act:none V1/2inact:→ Iω:none ExTraFol:none | (Lampert et al., 2008) |
| 488 | $Na_v1.7$ | F1449Y | $Na_v1.7$:F1460 $Na_v1.7$-2:F1460 $Na_v1.7$-3:F1449 $Na_v1.7$-4:F1449 | F1460Y | erythromelalgia | | Imax:none τrec:none Ip:none V1/2act:none V1/2inact:→ Iω:none ExTraFol:none | (Lampert et al., 2008) |
| 489 | $Na_v1.7$ | F1449V | $Na_v1.7$:F1460 $Na_v1.7$-2:F1460 $Na_v1.7$-3:F1449 $Na_v1.7$-4:F1449 | F1460V | IEM/PEM | GOF | Imax:none τrec:none Ip:none V1/2act:← V1/2inact:→ Iω:none ExTraFol:none | (Dib-Hajj et al., 2005) (Lampert et al., 2008) |
| 490 | $Na_v1.7$ | I1461T | $Na_v1.7$:I1472 $Na_v1.7$-2:I1472 $Na_v1.7$-3:I1461 $Na_v1.7$-4:I1461 | I1472T | PEXPD | GOF | Imax:none τrec:none Ip:↑ V1/2act:← V1/2inact:none | (Fertleman et al., 2006) |

| | Protein | Mutation | Isoform | Mutation (MSA) | Phenotype | GoF/LoF | Gating Properties | Reference |
|---|---|---|---|---|---|---|---|---|
| | | | | | | | Iω:none<br>ExTraFol:none | |
| 491 | $Na_v$1.7 | I1461T | $Na_v$1.7:I1472<br>$Na_v$1.7-2:I1472<br>$Na_v$1.7-3:I1461<br>$Na_v$1.7-4:I1461 | I1472T | PEXPD | GOF | Imax:none<br>τrec:none<br>Ip:↑<br>V1/2act:→<br>V1/2inact:→<br>Iω:none<br>ExTraFol:none | (Fertleman et al., 2006) |
| 492 | $Na_v$1.7 | F1462V | $Na_v$1.7:F1473<br>$Na_v$1.7-2:F1473<br>$Na_v$1.7-3:F1462<br>$Na_v$1.7-4:F1462 | F1473V | PEXPD | GOF | Imax:none<br>τrec:none<br>Ip:↑<br>V1/2act:←<br>V1/2inact:none<br>Iω:none<br>ExTraFol:none | (Fertleman et al., 2006) |
| 493 | $Na_v$1.7 | T1464I | $Na_v$1.7:T1475<br>$Na_v$1.7-2:T1475<br>$Na_v$1.7-3:T1464<br>$Na_v$1.7-4:T1464 | T1475I | PEXPD | GOF | Imax:none<br>τrec:none<br>Ip:↑<br>V1/2act:→<br>V1/2inact:→<br>Iω:none<br>ExTraFol:none | (Fertleman et al., 2006) (Theile et al., 2011) |
| 494 | $Na_v$1.7 | S1479D | $Na_v$1.7:S1490<br>$Na_v$1.7-2:S1490<br>$Na_v$1.7-3:S1479<br>$Na_v$1.7-4:S1479 | S1490D | | | Imax:none<br>τrec:none<br>Ip:none<br>V1/2act:none<br>V1/2inact:→<br>Iω:none<br>ExTraFol:none | (Z. Y. Tan et al., 2014) |
| 495 | $Na_v$1.7 | S1479E | $Na_v$1.7:S1490<br>$Na_v$1.7-2:S1490<br>$Na_v$1.7-3:S1479<br>$Na_v$1.7-4:S1479 | S1490E | | | Imax:none<br>τrec:none<br>Ip:none<br>V1/2act:→<br>V1/2inact:none<br>Iω:none<br>ExTraFol:none | (Z. Y. Tan et al., 2014) |
| 496 | $Na_v$1.7 | W1538R | $Na_v$1.7:W1549<br>$Na_v$1.7-2:W1549<br>$Na_v$1.7-3:W1538<br>$Na_v$1.7-4:W1538 | W1549R | PEM | GOF | Imax:none<br>τrec:none<br>Ip:none<br>V1/2act:←<br>V1/2inact:none | (Cregg et al., 2013) |

| | Protein | Mutation | Isoform | Mutation (MSA) | Phenotype | GoF/LoF | Gating Properties | Reference |
|---|---|---|---|---|---|---|---|---|
| | | | | | | | Iω:none ExTraFol:none | |
| 497 | $Na_v1.7$ | G1607R | $Na_v1.7$:G1618 $Na_v1.7$-2:G1618 $Na_v1.7$-3:G1607 $Na_v1.7$-4:G1607 | G1618R | PEXPD | GOF | Imax:none τrec:none Ip:↑ V1/2act:none V1/2inact:→ Iω:none ExTraFol:none | (Choi et al., 2011) |
| 498 | $Na_v1.7$ | L1612P | $Na_v1.7$:L1623 $Na_v1.7$-2:L1623 $Na_v1.7$-3:L1612 $Na_v1.7$-4:L1612 | L1623P | PEXPD | GOF | Imax:↑ τrec:none Ip:none V1/2act:← V1/2inact:→ Iω:none ExTraFol:none | (Suter et al., 2015) |
| 499 | $Na_v1.7$ | M1627K | $Na_v1.7$:M1638 $Na_v1.7$-2:M1638 $Na_v1.7$-3:M1627 $Na_v1.7$-4:M1627 | M1638K | PEXPD | GOF | Imax:none τrec:none Ip:↑ V1/2act:none V1/2inact:→ Iω:none ExTraFol:none | (Fertleman et al., 2006) (Dib-Hajj et al., 2008) (Theile & Cummins, 2011) |
| 500 | $Na_v1.7$ | A1632G | $Na_v1.7$:A1643 $Na_v1.7$-2:A1643 $Na_v1.7$-3:A1632 $Na_v1.7$-4:A1632 | A1643G | IEM | GOF | Imax:none τrec:none Ip:none V1/2act:← V1/2inact:→ Iω:none ExTraFol:none | (Y. Yang et al., 2016) |
| 501 | $Na_v1.7$ | A1632T | $Na_v1.7$:A1643 $Na_v1.7$-2:A1643 $Na_v1.7$-3:A1632 $Na_v1.7$-4:A1632 | A1643T | | | Imax:none τrec:none Ip:↑ V1/2act:none V1/2inact:→ Iω:none ExTraFol:none | (Eberhardt et al., 2014) |
| 502 | $Na_v1.7$ | A1632E | $Na_v1.7$:A1643 $Na_v1.7$-2:A1643 $Na_v1.7$-3:A1632 $Na_v1.7$-4:A1632 | A1643E | PEXPD/PEM | GOF | Imax:↑ τrec:none Ip:↑ V1/2act:← V1/2inact:→ | (Estacion et al., 2008) (Rühlmann et al., 2020) (Estacion et al., 2008) |

| | Protein | Mutation | Isoform | Mutation (MSA) | Phenotype | GoF/LoF | Gating Properties | Reference |
|---|---|---|---|---|---|---|---|---|
| | | | | | | | Iω:none<br>ExTraFol:none | |
| 503 | $Na_v1.7$ | M1532I | $Na_v1.7$:M1532<br>$Na_v1.7$-2:M1532<br>$Na_v1.7$-3:M1521<br>$Na_v1.7$-4:M1521 | M1739I | SFNP | GOF | Imax:none<br>τrec:none<br>Ip:none<br>V1/2act:←<br>V1/2inact:→<br>Iω:none<br>ExTraFol:none | (Faber, Hoeijmakers, et al., 2012) |
| 504 | $Na_v1.7$ | A1746G | $Na_v1.7$:A1757<br>$Na_v1.7$-2:A1757<br>$Na_v1.7$-3:A1746<br>$Na_v1.7$-4:A1746 | A1757G | PEM | GOF | Imax:none<br>τrec:none<br>Ip:none<br>V1/2act:←<br>V1/2inact:none<br>Iω:none<br>ExTraFol:none | (Cregg et al., 2013) |
| 505 | $Na_v1.7$ | W1775R | $Na_v1.7$:W1786<br>$Na_v1.7$-2:W1786<br>$Na_v1.7$-3:W1775<br>$Na_v1.7$-4:W1775 | W1786R | CIP | | Imax:none<br>τrec:none<br>Ip:none<br>V1/2act:→<br>V1/2inact:none<br>Iω:none<br>ExTraFol:none | (Emery et al., 2015) |
| 506 | $Na_v1.7$ | F1748A | $Na_v1.7$:F1748<br>$Na_v1.7$-2:F1748<br>$Na_v1.7$-3:F1737<br>$Na_v1.7$-4:F1737 | F1959A | | | Imax:none<br>τrec:none<br>Ip:none<br>V1/2act:none<br>V1/2inact:→<br>Iω:none<br>ExTraFol:none | (Klasfauseweh et al., 2022) |
| 507 | $Na_v1.8$ | V94G | $Na_v1.8$:V94 | V99G | BRGDA1*<br>AF | LOF | Imax:↓<br>τrec:none<br>Ip:none<br>V1/2act:none<br>V1/2inact:none<br>Iω:none<br>ExTraFol:none | (Jabbari et al., 2015) |
| 508 | $Na_v1.8$ | Y158D | $Na_v1.8$:Y158 | Y170D | FEPS2*<br>AF | | Imax:↑<br>τrec:↑<br>Ip:↑<br>V1/2act:→<br>V1/2inact:← | (Savio-Galimberti et al., 2014) |

| | Protein | Mutation | Isoform | Mutation (MSA) | Phenotype | GoF/LoF | Gating Properties | Reference |
|---|---|---|---|---|---|---|---|---|
| | | | | | | | Iω:none ExTraFol:none | |
| 509 | Na$_v$1.8 | S242T | Na$_v$1.8:S242 | S255T | FEPS2* PDN PPN | GOF | Imax:none τrec:none Ip:none V1/2act:← V1/2inact:← Iω:none ExTraFol:none | (C. Han et al., 2018) |
| 510 | Na$_v$1.8 | L388M | Na$_v$1.8:L388 | L471M | SUD | | Imax:↓ τrec:none Ip:none V1/2act:none V1/2inact:none Iω:none ExTraFol:none | (Gando et al., 2019) |
| 511 | Na$_v$1.8 | L554P | Na$_v$1.8:L554 | L728P | FEPS2* SFNP | GOF | Imax:none τrec:↑ Ip:↑ V1/2act:none V1/2inact:none Iω:none ExTraFol:none | (Faber, Lauria, Merkies, Cheng, Han, Ahn, Persson, Hoeijmakers, Gerrits, Pierro, Lombardi, et al., 2012) |
| 512 | Na$_v$1.8 | M650K | Na$_v$1.8:M650 | M833K | IEM | | Imax:none τrec:none Ip:none V1/2act:none V1/2inact:← Iω:none ExTraFol:none | (Kist et al., 2016) |
| 513 | Na$_v$1.8 | R756W | Na$_v$1.8:R756 | R941W | FEPS2* SUD | | Imax:none τrec:none Ip:none V1/2act:→ V1/2inact:none Iω:none ExTraFol:none | (Gando et al., 2019) |
| 514 | Na$_v$1.8 | R814H | Na$_v$1.8:R814 | R1005H | FEPS2* AF | | Imax:↑ τrec:↓ Ip:↑ V1/2act:→ V1/2inact:← | (Savio-Galimberti et al., 2014) |

| | Protein | Mutation | Isoform | Mutation (MSA) | Phenotype | GoF/LoF | Gating Properties | Reference |
|---|---|---|---|---|---|---|---|---|
| | | | | | | | Iω:none<br>ExTraFol:none | |
| 515 | $Na_v1.8$ | V1073A | $Na_v1.8$:V1073 | V1316A | FEPS2*<br>AF | GOF | Imax:↑<br>τrec:none<br>Ip:↑<br>V1/2act:→<br>V1/2inact:none<br>Iω:none<br>ExTraFol:none | (Jabbari et al., 2015)<br>(Coates et al., 2019) |
| 516 | $Na_v1.8$ | L1092P | $Na_v1.8$:L1092 | L1336P | FEPS2*<br>AF | GOF | Imax:↑<br>τrec:↑<br>Ip:↑<br>V1/2act:→<br>V1/2inact:none<br>Iω:none<br>ExTraFol:none | (Jabbari et al., 2015) |
| 517 | $Na_v1.8$ | A1304T | $Na_v1.8$:A1304 | A1548T | FEPS2*<br>SFNP | GOF | Imax:none<br>τrec:none<br>Ip:↑<br>V1/2act:←<br>V1/2inact:none<br>Iω:none<br>ExTraFol:none | (Faber, Lauria, Merkies, Cheng, Han, Ahn, Persson, Hoeijmakers, Gerrits, Pierro, Lombardi, et al., 2012) |
| 518 | $Na_v1.8$ | V1518I | $Na_v1.8$:V1518 | V1764I | SUD | | Imax:none<br>τrec:none<br>Ip:none<br>V1/2act:→<br>V1/2inact:none<br>Iω:none<br>ExTraFol:none | (Gando et al., 2019) |
| 519 | $Na_v1.8$ | R1588Q | $Na_v1.8$:R1588 | R1836Q | AF | LOF | Imax:none<br>τrec:none<br>Ip:none<br>V1/2act:none<br>V1/2inact:←<br>Iω:none<br>ExTraFol:none | (Jabbari et al., 2015) |
| 520 | $Na_v1.8$ | D1639N | $Na_v1.8$:D1639 | D1887N | SFNP | LOF | Imax:↓<br>τrec:none<br>Ip:none<br>V1/2act:none<br>V1/2inact:none | (Kaluza et al., 2018) |

| | Protein | Mutation | Isoform | Mutation (MSA) | Phenotype | GoF/LoF | Gating Properties | Reference |
|---|---|---|---|---|---|---|---|---|
| | | | | | | | Iω:none<br>ExTraFol:↓ | |
| 521 | $Na_v$1.8 | G1662S | $Na_v$1.8:G1662 | G1910S | BRGDA1*<br>SFNP | | Imax:none<br>τrec:↑<br>Ip:↑<br>V1/2act:none<br>V1/2inact:→<br>Iω:none<br>ExTraFol:none | (C. Han et al., 2014) |
| 522 | $Na_v$1.8 | I1706V | $Na_v$1.8:I1706 | I1955V | small-fiber neuropathy-associated | | Imax:none<br>τrec:none<br>Ip:none<br>V1/2act:←<br>V1/2inact:none<br>Iω:none<br>ExTraFol:none | (Alsaloum et al., 2021) |
| 523 | $Na_v$1.8 | A1886V | $Na_v$1.8:A1886 | A2137V | BRGDA1*<br>AF | | Imax:↑<br>τrec:↓<br>Ip:↑<br>V1/2act:none<br>V1/2inact:none<br>Iω:none<br>ExTraFol:none | (Savio-Galimberti et al., 2014) |
| 524 | $Na_v$1.9 | R222H | $Na_v$1.9:R222<br>$Na_v$1.9-2:R222<br>$Na_v$1.9-3:R222 | R228H | FEPS3*<br>early-onset pain symptoms | GOF | Imax:none<br>τrec:none<br>Ip:↓<br>V1/2act:←<br>V1/2inact:none<br>Iω:none<br>ExTraFol:none | (C. Han et al., 2017) |
| 525 | $Na_v$1.9 | I381T | $Na_v$1.9:I381<br>$Na_v$1.9-2:I381<br>$Na_v$1.9-3:I381 | I461T | FEPS3*<br>Painful Neuropathy | | Imax:none<br>τrec:none<br>Ip:none<br>V1/2act:←<br>V1/2inact:→<br>Iω:none<br>ExTraFol:none | (J. Huang, Han, et al., 2014) |
| 526 | $Na_v$1.9 | K419N | $Na_v$1.9:K419<br>$Na_v$1.9-2:K419<br>$Na_v$1.9-3:K419 | K499N | Painful Neuropathy | | Imax:none<br>τrec:none<br>Ip:none<br>V1/2act:none<br>V1/2inact:→ | (Zhou et al., 2017) |

| | Protein | Mutation | Isoform | Mutation (MSA) | Phenotype | GoF/LoF | Gating Properties | Reference |
|---|---|---|---|---|---|---|---|---|
| | | | | | | | Iω:none<br>ExTraFol:none | |
| 527 | Na$_v$1.9 | A582T | Na$_v$1.9:A582<br>Na$_v$1.9-2:A582<br>Na$_v$1.9-3:A582 | A853T | HSAN7*<br>Painful Neuropathy | | Imax:none<br>τrec:none<br>Ip:none<br>V1/2act:none<br>V1/2inact:→<br>Iω:none<br>ExTraFol:none | (Zhou et al., 2017) |
| 528 | Na$_v$1.9 | G699R | Na$_v$1.9:G699<br>Na$_v$1.9-2:G699<br>Na$_v$1.9-3:G699 | G970R | FEPS3*<br>HSAN7*<br>Painful Neuropathy | | Imax:none<br>τrec:none<br>Ip:↑<br>V1/2act:←<br>V1/2inact:→<br>Iω:none<br>ExTraFol:none | (C. Han et al., 2015) |
| 529 | Na$_v$1.9 | L811P | Na$_v$1.9:L811<br>Na$_v$1.9-2:L811<br>Na$_v$1.9-3:L811 | L1083P | HSAN7*<br>cold-aggravated peripheral pain | | Imax:none<br>τrec:none<br>Ip:none<br>V1/2act:←<br>V1/2inact:none<br>Iω:none<br>ExTraFol:none | (Leipold et al., 2015) |
| 530 | Na$_v$1.9 | N816K | Na$_v$1.9:N816<br>Na$_v$1.9-2:N816<br>Na$_v$1.9-3:N816 | N1088K | | GOF | Imax:↑<br>τrec:none<br>Ip:none<br>V1/2act:←<br>V1/2inact:none<br>Iω:none<br>ExTraFol:none | (J. Huang et al., 2019) |
| 531 | Na$_v$1.9 | A842P | Na$_v$1.9:A842<br>Na$_v$1.9-2:A842<br>Na$_v$1.9-3:A842 | A1114P | Painful Neuropathy | | Imax:none<br>τrec:none<br>Ip:none<br>V1/2act:none<br>V1/2inact:→<br>Iω:none<br>ExTraFol:none | (Zhou et al., 2017) |
| 532 | Na$_v$1.9 | L1158P | Na$_v$1.9:L1158<br>Na$_v$1.9-2:L1158<br>Na$_v$1.9-3:L1120 | L1505P | FEPS3*<br>Painful Neuropathy | | Imax:none<br>τrec:none<br>Ip:none<br>V1/2act:←<br>V1/2inact:none | (J. Huang, Han, et al., 2014) |

| | Protein | Mutation | Isoform | Mutation (MSA) | Phenotype | GoF/LoF | Gating Properties | Reference |
|---|---|---|---|---|---|---|---|---|
| | | | | | | | Iω:none<br>ExTraFol:none | |
| 533 | $Na_v$1.9 | V1184A | $Na_v$1.9:V1184<br>$Na_v$1.9-2:V1184<br>$Na_v$1.9-3:V1146 | V1531A | HSAN7*<br>cold-aggravated peripheral pain | GOF | Imax:↑<br>τrec:none<br>Ip:↑<br>V1/2act:←<br>V1/2inact:none<br>Iω:none<br>ExTraFol:none | (Leipold et al., 2015) |
| 534 | $Na_v$1.9 | L1302F | $Na_v$1.9:L1302<br>$Na_v$1.9-2:L1302<br>$Na_v$1.9-3:L1264 | L1658F | HSAN7*<br>insensitivity to pain | | Imax:none<br>τrec:none<br>Ip:none<br>V1/2act:←<br>V1/2inact:none<br>Iω:none<br>ExTraFol:none | (J. Huang et al., 2017) |
| 535 | $Na_v$1.9 | F1689L | $Na_v$1.9:F1689<br>$Na_v$1.9-2:--<br>$Na_v$1.9-3:F1651 | F2056L | HSAN7*<br>Painful Neuropathy | | Imax:none<br>τrec:none<br>Ip:none<br>V1/2act:none<br>V1/2inact:→<br>Iω:none<br>ExTraFol:none | (Zhou et al., 2017) |

* Phenotype Entries from UniProt

†Imax: Maximum current amplitude. τrec: Recovery form fast inactivation. Ip: Persistent current. V1/2act: Voltage half activation. V1/2inact: Voltage half inactivation. Iω: Omega current. ExTraFol: Expression, trafficking, folding.

A downward arrow (↓) indicates a decrease and an upward arrow (↑) will indicate an increase in that property.

A left pointing arrow (←) indicates a hyperpolarizing shift while a right pointing one (→) means a depolarizing shift in that property.

Table S4: Common mutations across $Na_v$ channels with gating properties.

| MSA | Protein | Phenotype | Mutation | Mutation (MSA ID) | Isoform | GoF/ LoF | Gating Properties | References |
|---|---|---|---|---|---|---|---|---|
| 108 | $Na_v1.7$ | CIP | R99H | R108H | $Na_v1.7$:R99<br>$Na_v1.7$-2:R99<br>$Na_v1.7$-3:R99<br>$Na_v1.7$-4:R99 | LOF | Imax:↓<br>τrec:none<br>Ip:none<br>V1/2act:none<br>V1/2inact:→<br>Iω:none<br>ExTraFol:↓ | (J. Sun et al., 2020) |
| 108 | $Na_v1.4$ | CMS16 | R104H | R108H | $Na_v1.4$:R104 | LOF | Imax:↓<br>τrec:none<br>Ip:none<br>V1/2act:→<br>V1/2inact:none<br>Iω:none<br>ExTraFol:none | (Zaharieva et al., 2016) |
| 145 | $Na_v1.4$ | SCM<br>PAM<br>PMC | I141V | I145V | $Na_v1.4$:I141 | GOF | Imax:↑<br>τrec:none<br>Ip:↑<br>V1/2act:←<br>V1/2inact:none<br>Iω:none<br>ExTraFol:none | (Petitprez, Tiab, et al., 2008) |
| 145 | $Na_v1.7$ | PEM | I136V | I145V | $Na_v1.7$:I136<br>$Na_v1.7$-2:I136<br>$Na_v1.7$-3:I136<br>$Na_v1.7$-4:I136 | GOF | Imax:none<br>τrec:↑<br>Ip:↑<br>V1/2act:←<br>V1/2inact:→<br>Iω:none<br>ExTraFol:none | (X. Cheng et al., 2008b) |
| 145 | $Na_v1.5$ | Hyperexcitability | I141V | I145V | $Na_v1.5$:I141<br>$Na_v1.5$-2:I141<br>$Na_v1.5$-3:I141<br>$Na_v1.5$-4:I141<br>$Na_v1.5$-5:I141<br>$Na_v1.5$-6:I141 | GOF | Imax:none<br>τrec:none<br>Ip:none<br>V1/2act:←<br>V1/2inact:none<br>Iω:none<br>ExTraFol:none | (Amarouch et al., 2014) |
| 212 | $Na_v1.4$ | HyperPP | A204E | A212E | $Na_v1.4$:A204 | | Imax:none<br>τrec:none<br>Ip:none<br>V1/2act:←<br>V1/2inact:← | (Kokunai et al., 2018) |

| MSA | Protein | Phenotype | Mutation | Mutation (MSA ID) | Isoform | GoF/ LoF | Gating Properties | References |
|---|---|---|---|---|---|---|---|---|
| | | | | | | | Iω:none<br>ExTraFol:none | |
| 212 | $Na_v1.5$ | multifocal ectopic purkinje-related premature contractions | A204E | A212E | $Na_v1.5$:A204<br>$Na_v1.5$-2:A204<br>$Na_v1.5$-3:A204<br>$Na_v1.5$-4:A204<br>$Na_v1.5$-5:A204<br>$Na_v1.5$-6:A204 | GOF | Imax:none<br>τrec:none<br>Ip:none<br>V1/2act:←<br>V1/2inact:none<br>Iω:none<br>ExTraFol:none | (Doisne et al., 2020b) |
| 234 | $Na_v1.2$ | BFIS3 | R223Q | R234Q | $Na_v1.2$:R223<br>$Na_v1.2$-2:R223 | GOF | Imax:none<br>τrec:none<br>Ip:none<br>V1/2act:→<br>V1/2inact:→<br>Iω:none<br>ExTraFol:none | (Scalmani et al., 2006) |
| 234 | $Na_v1.5$ | LQT3 | R225Q | R234Q | $Na_v1.5$:R225<br>$Na_v1.5$-2:R225<br>$Na_v1.5$-3:R225<br>$Na_v1.5$-4:R225<br>$Na_v1.5$-5:R225<br>$Na_v1.5$-6:R225 | | Imax:none<br>τrec:none<br>Ip:none<br>V1/2act:→<br>V1/2inact:←<br>Iω:none<br>ExTraFol:none | (L. Q. Chen et al., 1996) |
| 234 | $Na_v1.5$ | LQT3<br>CCD<br>PFHB1A<br>BRGDA1<br>IBS | R225W | R234W | $Na_v1.5$:R225<br>$Na_v1.5$-2:R225<br>$Na_v1.5$-3:R225<br>$Na_v1.5$-4:R225<br>$Na_v1.5$-5:R225<br>$Na_v1.5$-6:R225 | MIX | Imax:↓<br>τrec:↑<br>Ip:none<br>V1/2act:→<br>V1/2inact:→<br>Iω:↑<br>ExTraFol:none | (Bezzina et al., 2003)<br>(Moreau, Gosselin-Badaroudine, Delemotte, et al., 2015)<br>(Strege et al., 2018) |
| 234 | $Na_v1.4$ | SCM<br>PAM | R225W | R234W | $Na_v1.4$:R225 | LOF | Imax:↓<br>τrec:none<br>Ip:none<br>V1/2act:→<br>V1/2inact:none<br>Iω:none<br>ExTraFol:none | (Zaharieva et al., 2016) |
| 255 | $Na_v1.7$ | PERYTHM | S241T | S255T | $Na_v1.7$:S241<br>$Na_v1.7$-2:S241<br>$Na_v1.7$-3:S241<br>$Na_v1.7$-4:S241 | GOF | Imax:none<br>τrec:none<br>Ip:↑<br>V1/2act:← | (Lampert et al., 2006)<br>(Alsaloum et al., 2023) |

| MSA | Protein | Phenotype | Mutation | Mutation (MSA ID) | Isoform | GoF/ LoF | Gating Properties | References |
|---|---|---|---|---|---|---|---|---|
| | | | | | | | V1/2inact:none<br>Iω:none<br>ExTraFol:none | |
| 255 | $Na_v1.8$ | FEPS2 | S242T | S255T | $Na_v1.8$:S242 | GOF | Imax:none<br>τrec:none<br>Ip:none<br>V1/2act:←<br>V1/2inact:←<br>Iω:none<br>ExTraFol:none | (C. Han et al., 2018) |
| 275 | $Na_v1.1$ | FHM3 | L263V | L275V | $Na_v1.1$:L263<br>$Na_v1.1$-2:L263<br>$Na_v1.1$-3:L263 | GOF | Imax:none<br>τrec:none<br>Ip:↑<br>V1/2act:←<br>V1/2inact:→<br>Iω:none<br>ExTraFol:none | (Kahlig et al., 2008) |
| 275 | $Na_v1.4$ | PMC CAM | L266V | L275V | $Na_v1.4$:L266 | GOF | Imax:none<br>τrec:none<br>Ip:none<br>V1/2act:none<br>V1/2inact:→<br>Iω:none<br>ExTraFol:none | (F.-F. F. Wu et al., 2001) |
| 279 | $Na_v1.5$ | BRGDA1 | Q270K | Q279K | $Na_v1.5$:Q270<br>$Na_v1.5$-2:Q270<br>$Na_v1.5$-3:Q270<br>$Na_v1.5$-4:Q270<br>$Na_v1.5$-5:Q270<br>$Na_v1.5$-6:Q270 | GOF | Imax:↓<br>τrec:none<br>Ip:↑<br>V1/2act:→<br>V1/2inact:→<br>Iω:none<br>ExTraFol:none | (Calloe et al., 2011) |
| 279 | $Na_v1.4$ | PMC | Q270K | Q279K | $Na_v1.4$:Q270 | GOF | Imax:none<br>τrec:↑<br>Ip:none<br>V1/2act:→<br>V1/2inact:→<br>Iω:none<br>ExTraFol:none | (Carle et al., 2009) |
| 434 | $Na_v1.2$ | ASD | R379H | R434H | $Na_v1.2$:R379<br>$Na_v1.2$-2:R379 | LOF | Imax:↓<br>τrec:none<br>Ip:none<br>V1/2act:none<br>V1/2inact:none | (Ben-Shalom et al., 2017) |

| MSA | Protein | Phenotype | Mutation | Mutation (MSA ID) | Isoform | GoF/ LoF | Gating Properties | References |
|---|---|---|---|---|---|---|---|---|
| | | | | | | | Iω:none<br>ExTraFol:none | |
| 434 | $Na_v1.5$ | BRGDA1 | R367H | R434H | $Na_v1.5$:R367<br>$Na_v1.5$-2:R367<br>$Na_v1.5$-3:R367<br>$Na_v1.5$-4:R367<br>$Na_v1.5$-5:R367<br>$Na_v1.5$-6:R367 | LOF | Imax:↓<br>τrec:none<br>Ip:none<br>V1/2act:none<br>V1/2inact:none<br>Iω:none<br>ExTraFol:none | (Vatta, Dumaine, Varghese, et al., 2002)<br>(Watanabe, Nogami, Ohkubo, Kawata, Hayashi, Ishikawa, Makiyama, Nagao, Yagihara, Takehara, Shimizu, Makita, et al., 2011)<br>(Selga et al., 2018) |
| 473 | $Na_v1.4$ | PMC HYPP | N440K | N473K | $Na_v1.4$:N440 | GOF | Imax:none<br>τrec:none<br>Ip:↑<br>V1/2act:none<br>V1/2inact:→<br>Iω:none<br>ExTraFol:none | (Lossin, Nam, et al., 2012) |
| 473 | $Na_v1.7$ | PERYTHM | N395K | N473K | $Na_v1.7$:N395<br>$Na_v1.7$-2:N395<br>$Na_v1.7$-3:N395<br>$Na_v1.7$-4:N395 | GOF | Imax:none<br>τrec:none<br>Ip:none<br>V1/2act:←<br>V1/2inact:none<br>Iω:none<br>ExTraFol:none | (Sheets et al., 2007) |
| 473 | $Na_v1.5$ | RWS LQT3 | N406K | N473K | $Na_v1.5$:N406<br>$Na_v1.5$-2:N406<br>$Na_v1.5$-3:N406<br>$Na_v1.5$-4:N406<br>$Na_v1.5$-5:N406<br>$Na_v1.5$-6:N406 | GOF | Imax:↓<br>τrec:↓<br>Ip:↑<br>V1/2act:→<br>V1/2inact:none<br>Iω:none<br>ExTraFol:↓ | (R. M. Hu et al., 2018)<br>(Kato et al., 2014) |
| 478 | $Na_v1.5$ | RWS LQT3 | V411M | V478M | $Na_v1.5$:V411<br>$Na_v1.5$-2:V411<br>$Na_v1.5$-3:V411<br>$Na_v1.5$-4:V411<br>$Na_v1.5$-5:V411<br>$Na_v1.5$-6:V411 | GOF | Imax:none<br>τrec:none<br>Ip:↑<br>V1/2act:←<br>V1/2inact:←<br>Iω:none<br>ExTraFol:none | (Horne et al., 2011) |

| MSA | Protein | Phenotype | Mutation | Mutation (MSA ID) | Isoform | GoF/ LoF | Gating Properties | References |
|---|---|---|---|---|---|---|---|---|
| 478 | $Na_v1.4$ | PAM<br>HYPP | V445M | V478M | $Na_v1.4$:V445 | GOF | Imax:none<br>τrec:none<br>Ip:↑<br>V1/2act:←<br>V1/2inact:none<br>Iω:none<br>ExTraFol:none | (D. W. Wang et al., 1999) |
| 478 | $Na_v1.7$ | PEM | V400M | V478M | $Na_v1.7$:V400<br>$Na_v1.7$-2:V400<br>$Na_v1.7$-3:V400<br>$Na_v1.7$-4:V400 | GOF | Imax:none<br>τrec:none<br>Ip:none<br>V1/2act:←<br>V1/2inact:none<br>Iω:none<br>ExTraFol:none | (Y. Yang et al., 2012) |
| 862 | $Na_v1.2$ | DEE11 | T773I | T862I | $Na_v1.2$:T773<br>$Na_v1.2$-2:T773 | GoF | Imax:none<br>τrec:none<br>Ip:↑<br>V1/2act:←<br>V1/2inact:none<br>Iω:none<br>ExTraFol:none | (Lauxmann et al., 2018) |
| 862 | $Na_v1.6$ | DEE13<br>EIEE | T767I | T862I | $Na_v1.6$:T767<br>$Na_v1.6$-2:T767<br>$Na_v1.6$-3:T778<br>$Na_v1.6$-4:T767<br>$Na_v1.6$-5:T767 | GOF | Imax:↓<br>τrec:none<br>Ip:↑<br>V1/2act:←<br>V1/2inact:none<br>Iω:none<br>ExTraFol:none | (Pan & Cummins, 2020) |
| 941 | $Na_v1.4$ | HOKPP2 | R669H | R941H | $Na_v1.4$:R669 | LOF | Imax:none<br>τrec:↓<br>Ip:none<br>V1/2act:none<br>V1/2inact:←<br>Iω:none<br>ExTraFol:none | (Kuzmenkin et al., 2002) |
| 941 | $Na_v1.5$ | | R808H | R941H | $Na_v1.5$:R808<br>$Na_v1.5$-2:R808<br>$Na_v1.5$-3:R808<br>$Na_v1.5$-4:R808<br>$Na_v1.5$-5:R808<br>$Na_v1.5$-6:R808 | | Imax:none<br>τrec:none<br>Ip:none<br>V1/2act:→<br>V1/2inact:none<br>Iω:none<br>ExTraFol:none | (L. Q. Chen et al., 1996) |

| MSA | Protein | Phenotype | Mutation | Mutation (MSA ID) | Isoform | GoF/ LoF | Gating Properties | References |
|---|---|---|---|---|---|---|---|---|
| 941 | $Na_v1.1$ | GEFSP2 | R859H | R941H | $Na_v1.1$:R859<br>$Na_v1.1$-2:R848<br>$Na_v1.1$-3:R831 | LOF | Imax:none<br>τrec:↓<br>Ip:↑<br>V1/2act:←<br>V1/2inact:←<br>Iω:none<br>ExTraFol:none | (Brunklaus et al., 2020)<br>(Volkers et al., 2011) |
| 941 | $Na_v1.5$ | BRGDA1 | R808C | R941C | $Na_v1.5$:R808<br>$Na_v1.5$-2:R808<br>$Na_v1.5$-3:R808<br>$Na_v1.5$-4:R808<br>$Na_v1.5$-5:R808<br>$Na_v1.5$-6:R808 | LOF | Imax:↓<br>τrec:↓<br>Ip:none<br>V1/2act:none<br>V1/2inact:←<br>Iω:none<br>ExTraFol:none | (Glazer et al., 2020) |
| 941 | $Na_v1.1$ | DRVT<br>GEFSP2 | R859C | R941C | $Na_v1.1$:R859<br>$Na_v1.1$-2:R848<br>$Na_v1.1$-3:R831 | LOF | Imax:↓<br>τrec:none<br>Ip:none<br>V1/2act:none<br>V1/2inact:none<br>Iω:none<br>ExTraFol:none | (Bechi et al., 2015) |
| 944 | $Na_v1.5$ | BRGDA1 | R811H | R944H | $Na_v1.5$:R811<br>$Na_v1.5$-2:R811<br>$Na_v1.5$-3:R811<br>$Na_v1.5$-4:R811<br>$Na_v1.5$-5:R811<br>$Na_v1.5$-6:R811 | LOF | Imax:↓<br>τrec:↓<br>Ip:none<br>V1/2act:none<br>V1/2inact:←<br>Iω:none<br>ExTraFol:none | (Calloe et al., 2013) |
| 944 | $Na_v1.4$ | HOKPP2 | R672H | R944H | $Na_v1.4$:R672 | LOF | Imax:none<br>τrec:↓<br>Ip:none<br>V1/2act:→<br>V1/2inact:←<br>Iω:none<br>ExTraFol:↓ | (Jurkat-Rott et al., 2000)<br>(Kuzmenkin et al., 2002) |
| 947 | $Na_v1.5$ | BRGDA1 | R814Q | R947Q | $Na_v1.5$:R814<br>$Na_v1.5$-2:R814<br>$Na_v1.5$-3:R814<br>$Na_v1.5$-4:R814<br>$Na_v1.5$-5:R814<br>$Na_v1.5$-6:R814 | GOF | Imax:none<br>τrec:none<br>Ip:↑<br>V1/2act:none<br>V1/2inact:←<br>Iω:none<br>ExTraFol:none | (Glazer et al., 2020) |

| MSA | Protein | Phenotype | Mutation | Mutation (MSA ID) | Isoform | GoF/ LoF | Gating Properties | References |
|---|---|---|---|---|---|---|---|---|
| 947 | $Na_v1.6$ | DEE13 | R850Q | R947Q | $Na_v1.6$:R850<br>$Na_v1.6$-2:R850<br>$Na_v1.6$-3:R861<br>$Na_v1.6$-4:R850<br>$Na_v1.6$-5:R850 | GOF | Imax:none<br>τrec:none<br>Ip:↑<br>V1/2act:←<br>V1/2inact:none<br>Iω:none<br>ExTraFol:none | (Pan & Cummins, 2020) |
| 965 | $Na_v1.3$ | DEE62 | I875T | I965T | $Na_v1.3$:I875<br>$Na_v1.3$-2:I826<br>$Na_v1.3$-3:I875<br>$Na_v1.3$-4:I826 | GOF | Imax:↑<br>τrec:none<br>Ip:↑<br>V1/2act:←<br>V1/2inact:→<br>Iω:none<br>ExTraFol:none | (Zaman et al., 2018) |
| 965 | $Na_v1.4$ | PMC HYPP | I693T | I965T | $Na_v1.4$:I693 | GOF | Imax:none<br>τrec:none<br>Ip:none<br>V1/2act:←<br>V1/2inact:none<br>Iω:none<br>ExTraFol:none | (Bendahhou et al., 2002) |
| 970 | $Na_v1.2$ | EIMFS | G879R | G970R | $Na_v1.2$:G879<br>$Na_v1.2$-2:G879 | LOF | Imax:↓<br>τrec:none<br>Ip:none<br>V1/2act:←<br>V1/2inact:→<br>Iω:none<br>ExTraFol:↓ | (X. R. Yang et al., 2022) |
| 970 | $Na_v1.9$ | FEPS3 HSAN7 | G699R | G970R | $Na_v1.9$:G699<br>$Na_v1.9$-2:G699<br>$Na_v1.9$-3:G699 | | Imax:none<br>τrec:none<br>Ip:↑<br>V1/2act:←<br>V1/2inact:→<br>Iω:none<br>ExTraFol:none | (C. Han et al., 2015) |
| 1035 | $Na_v1.1$ | DRVT GEFSP2 | R946H | R1035H | $Na_v1.1$:R946<br>$Na_v1.1$-2:R935<br>$Na_v1.1$-3:R918 | LOF | Imax:↓<br>τrec:none<br>Ip:none<br>V1/2act:none<br>V1/2inact:none<br>Iω:none<br>ExTraFol:none | (Brunklaus et al., 2020) |

| MSA | Protein | Phenotype | Mutation | Mutation (MSA ID) | Isoform | GoF/ LoF | Gating Properties | References |
|---|---|---|---|---|---|---|---|---|
| 1035 | $Na_v1.2$ | ASD | R937H | R1035H | $Na_v1.2$:R937<br>$Na_v1.2$-2:R937 | LOF | Imax:↓<br>τrec:none<br>Ip:none<br>V1/2act:none<br>V1/2inact:none<br>Iω:none<br>ExTraFol:none | (Ben-Shalom et al., 2017) |
| 1035 | $Na_v1.5$ | BRGDA1 | R893C | R1035C | $Na_v1.5$:R893<br>$Na_v1.5$-2:R893<br>$Na_v1.5$-3:R893<br>$Na_v1.5$-4:R893<br>$Na_v1.5$-5:R893<br>$Na_v1.5$-6:R893 | LOF | Imax:↓<br>τrec:none<br>Ip:none<br>V1/2act:none<br>V1/2inact:none<br>Iω:none<br>ExTraFol:none | (Ishikawa et al., 2021) |
| 1035 | $Na_v1.2$ | ASD<br>BFIS3<br>ID | R937C | R1035C | $Na_v1.2$:R937<br>$Na_v1.2$-2:R937 | LOF | Imax:↓<br>τrec:none<br>Ip:none<br>V1/2act:none<br>V1/2inact:none<br>Iω:none<br>ExTraFol:none | (Ben-Shalom et al., 2017) |
| 1035 | $Na_v1.1$ | DRVT<br>GEFSP2<br>EIEE | R946C | R1035C | $Na_v1.1$:R946<br>$Na_v1.1$-2:R935<br>$Na_v1.1$-3:R918 | LOF | Imax:↓<br>τrec:none<br>Ip:none<br>V1/2act:none<br>V1/2inact:none<br>Iω:none<br>ExTraFol:none | (Brunklaus et al., 2020) |
| 1070 | $Na_v1.1$ | DRVT<br>ICEGTC | G979R | G1070R | $Na_v1.1$:G979<br>$Na_v1.1$-2:G968<br>$Na_v1.1$-3:G951 | LOF | Imax:↓<br>τrec:none<br>Ip:none<br>V1/2act:none<br>V1/2inact:none<br>Iω:none<br>ExTraFol:none | (Rhodes et al., 2005) |
| 1070 | $Na_v1.6$ | NDD | G964R | G1070R | $Na_v1.6$:G964<br>$Na_v1.6$-2:G964<br>$Na_v1.6$-3:G975<br>$Na_v1.6$-4:G964<br>$Na_v1.6$-5:G964 | LOF | Imax:↓<br>τrec:none<br>Ip:none<br>V1/2act:none<br>V1/2inact:none<br>Iω:none<br>ExTraFol:none | (Wagnon et al., 2017) |

| MSA | Protein | Phenotype | Mutation | Mutation (MSA ID) | Isoform | GoF/ LoF | Gating Properties | References |
|---|---|---|---|---|---|---|---|---|
| 1434 | Na$_v$1.4 | HOKPP2<br>CMS16 | D1069N | D1434N | Na$_v$1.4:D1069 | LOF | Imax:↓<br>τrec:none<br>Ip:none<br>V1/2act:→<br>V1/2inact:→<br>Iω:none<br>ExTraFol:none | (Zaharieva et al., 2016) |
| 1434 | Na$_v$1.5 | BRGDA1 | D1243N | D1434N | Na$_v$1.5:D1243<br>Na$_v$1.5-2:D1242<br>Na$_v$1.5-3:D1242<br>Na$_v$1.5-4:D1243<br>Na$_v$1.5-5:D1189<br>Na$_v$1.5-6:D1243 | | Imax:none<br>τrec:↑<br>Ip:none<br>V1/2act:→<br>V1/2inact:→<br>Iω:none<br>ExTraFol:none | (Glazer et al., 2020) |
| 1497 | Na$_v$1.5 | | R1306Q | R1497Q | Na$_v$1.5:R1306<br>Na$_v$1.5-2:R1305<br>Na$_v$1.5-3:R1305<br>Na$_v$1.5-4:R1306<br>Na$_v$1.5-5:R1252<br>Na$_v$1.5-6:R1306 | | Imax:none<br>τrec:none<br>Ip:none<br>V1/2act:none<br>V1/2inact:←<br>Iω:none<br>ExTraFol:none | (L. Q. Chen et al., 1996) |
| 1497 | Na$_v$1.4 | HOKPP2<br>HYPP | R1132Q | R1497Q | Na$_v$1.4:R1132 | LOF | Imax:none<br>τrec:none<br>Ip:none<br>V1/2act:→<br>V1/2inact:←<br>Iω:none<br>ExTraFol:none | (Carle et al., 2006) |
| 1521 | Na$_v$1.5 | RWS<br>LQT3 | A1330T | A1521T | Na$_v$1.5:A1330<br>Na$_v$1.5-2:A1329<br>Na$_v$1.5-3:A1329<br>Na$_v$1.5-4:A1330<br>Na$_v$1.5-5:A1276<br>Na$_v$1.5-6:A1330 | GOF | Imax:none<br>τrec:↑<br>Ip:none<br>V1/2act:none<br>V1/2inact:→<br>Iω:none<br>ExTraFol:none | (Smits, Veldkamp, et al., 2005) |
| 1521 | Na$_v$1.4 | PMC<br>PAM<br>HYPP | A1156T | A1521T | Na$_v$1.4:A1156 | GOF | Imax:none<br>τrec:↑<br>Ip:none<br>V1/2act:none<br>V1/2inact:→<br>Iω:none<br>ExTraFol:none | (Palmio et al., 2017) |

| MSA | Protein | Phenotype | Mutation | Mutation (MSA ID) | Isoform | GoF/ LoF | Gating Properties | References |
|---|---|---|---|---|---|---|---|---|
| 1523 | Na$_v$1.5 | LQT3<br>BRGDA1 | P1332L | P1523L | Na$_v$1.5:P1332<br>Na$_v$1.5-2:P1331<br>Na$_v$1.5-3:P1331<br>Na$_v$1.5-4:P1332<br>Na$_v$1.5-5:P1278<br>Na$_v$1.5-6:P1332 | GOF | Imax:none<br>τrec:↓<br>Ip:none<br>V1/2act:←<br>V1/2inact:←<br>Iω:none<br>ExTraFol:none | (Ruan et al., 2007) |
| 1523 | Na$_v$1.4 | HYPP | P1158L | P1523L | Na$_v$1.4:P1158 | GOF | Imax:none<br>τrec:none<br>Ip:none<br>V1/2act:none<br>V1/2inact:→<br>Iω:none<br>ExTraFol:none | (Desaphy et al., 2016) |
| 1523 | Na$_v$1.3 | DEE62 | P1333L | P1523L | Na$_v$1.3:P1333<br>Na$_v$1.3-2:P1284<br>Na$_v$1.3-3:P1333<br>Na$_v$1.3-4:P1284 | GOF | Imax:none<br>τrec:none<br>Ip:↑<br>V1/2act:←<br>V1/2inact:none<br>Iω:none<br>ExTraFol:none | (Zaman et al., 2018) |
| 1524 | Na$_v$1.2 | DEE11 | S1336Y | S1524Y | Na$_v$1.2:S1336<br>Na$_v$1.2-2:S1336 | MIX | Imax:↓<br>τrec:none<br>Ip:none<br>V1/2act:←<br>V1/2inact:→<br>Iω:none<br>ExTraFol:none | (Thompson et al., 2020) |
| 1524 | Na$_v$1.5 | RWS<br>LQT3<br>SIDS | S1333Y | S1524Y | Na$_v$1.5:S1333<br>Na$_v$1.5-2:S1332<br>Na$_v$1.5-3:S1332<br>Na$_v$1.5-4:S1333<br>Na$_v$1.5-5:S1279<br>Na$_v$1.5-6:S1333 | GOF | Imax:none<br>τrec:none<br>Ip:none<br>V1/2act:←<br>V1/2inact:→<br>Iω:none<br>ExTraFol:none | (H. Huang et al., 2009b) |
| 1525 | Na$_v$1.4 | PAM | I1160V | I1525V | Na$_v$1.4:I1160 | GOF | Imax:none<br>τrec:none<br>Ip:none<br>V1/2act:←<br>V1/2inact:none<br>Iω:none<br>ExTraFol:none | (Richmond et al., 1997) |

| MSA | Protein | Phenotype | Mutation | Mutation (MSA ID) | Isoform | GoF/ LoF | Gating Properties | References |
|---|---|---|---|---|---|---|---|---|
| 1525 | $Na_v1.6$ | DEE13 | I1327V | I1525V | $Na_v1.6$:I1327<br>$Na_v1.6$-2:I1327<br>$Na_v1.6$-3:I1338<br>$Na_v1.6$-4:--<br>$Na_v1.6$-5:I1286 | | Imax:none<br>τrec:none<br>Ip:none<br>V1/2act:←<br>V1/2inact:→<br>Iω:none<br>ExTraFol:none | (Barker et al., 2016) |
| 1531 | $Na_v1.1$ | GEFSP2 | V1353L | V1531L | $Na_v1.1$:V1353<br>$Na_v1.1$-2:V1342<br>$Na_v1.1$-3:V1325 | Mixed | Imax:↓<br>τrec:↑<br>Ip:none<br>V1/2act:none<br>V1/2inact:→<br>Iω:none<br>ExTraFol:none | (Berecki et al., 2019) |
| 1531 | $Na_v1.5$ | AS | V1340L | V1531L | $Na_v1.5$:V1340<br>$Na_v1.5$-2:V1339<br>$Na_v1.5$-3:V1339<br>$Na_v1.5$-4:V1340<br>$Na_v1.5$-5:V1286<br>$Na_v1.5$-6:V1340 | LOF | Imax:↓<br>τrec:none<br>Ip:none<br>V1/2act:→<br>V1/2inact:none<br>Iω:none<br>ExTraFol:none | (R. B. Tan et al., 2018) |
| 1667 | $Na_v1.4$ | SCM | F1298C | F1667C | $Na_v1.4$:F1298 | GOF | Imax:none<br>τrec:none<br>Ip:none<br>V1/2act:→<br>V1/2inact:→<br>Iω:none<br>ExTraFol:none | (Farinato et al., 2019) |
| 1667 | $Na_v1.5$ | RWS<br>LQT3 | F1473C | F1667C | $Na_v1.5$:F1473<br>$Na_v1.5$-2:F1472<br>$Na_v1.5$-3:F1472<br>$Na_v1.5$-4:F1473<br>$Na_v1.5$-5:F1419<br>$Na_v1.5$-6:F1455 | GOF | Imax:none<br>τrec:none<br>Ip:↑<br>V1/2act:none<br>V1/2inact:→<br>Iω:none<br>ExTraFol:none | (Bankston et al., 2007) |
| 1680 | $Na_v1.5$ | SIDS<br>LQT3 | F1486L | F1680L | $Na_v1.5$:F1486<br>$Na_v1.5$-2:F1485<br>$Na_v1.5$-3:F1485<br>$Na_v1.5$-4:F1486<br>$Na_v1.5$-5:F1432<br>$Na_v1.5$-6:F1468 | GOF | Imax:none<br>τrec:none<br>Ip:↑<br>V1/2act:none<br>V1/2inact:→<br>Iω:none<br>ExTraFol:none | (D. W. Wang et al., 2007) |

| MSA | Protein | Phenotype | Mutation | Mutation (MSA ID) | Isoform | GoF/ LoF | Gating Properties | References |
|---|---|---|---|---|---|---|---|---|
| 1680 | $Na_v$1.1 | FHM3 | F1499L | F1680L | $Na_v$1.1:F1499<br>$Na_v$1.1-2:F1488<br>$Na_v$1.1-3:F1471 | GOF | Imax:none<br>τrec:none<br>Ip:↑<br>V1/2act:none<br>V1/2inact:→<br>Iω:none<br>ExTraFol:none | (Barbieri et al., 2019) |
| 1768 | $Na_v$1.1 | EPI | E1587K | E1768K | $Na_v$1.1:E1587<br>$Na_v$1.1-2:E1576<br>$Na_v$1.1-3:E1559 | LOF | Imax:↓<br>τrec:none<br>Ip:none<br>V1/2act:none<br>V1/2inact:none<br>Iω:none<br>ExTraFol:none | (Kluckova et al., 2020) |
| 1768 | $Na_v$1.5 | BRGDA1 | E1574K | E1768K | $Na_v$1.5:E1574<br>$Na_v$1.5-2:E1573<br>$Na_v$1.5-3:--<br>$Na_v$1.5-4:E1574<br>$Na_v$1.5-5:E1520<br>$Na_v$1.5-6:E1556 | LOF | Imax:↓<br>τrec:none<br>Ip:none<br>V1/2act:→<br>V1/2inact:none<br>Iω:none<br>ExTraFol:none | (Glazer et al., 2020) |
| 1817 | $Na_v$1.2 | ICDEE | P1622S | P1817S | $Na_v$1.2:P1622<br>$Na_v$1.2-2:P1622 | LOF | Imax:none<br>τrec:↓<br>Ip:none<br>V1/2act:none<br>V1/2inact:←<br>Iω:none<br>ExTraFol:none | (Wolff et al., 2017) |
| 1817 | $Na_v$1.1 | DRVT ICEGTC | P1632S | P1817S | $Na_v$1.1:P1632<br>$Na_v$1.1-2:P1621<br>$Na_v$1.1-3:P1604 | LOF | Imax:none<br>τrec:none<br>Ip:↑<br>V1/2act:←<br>V1/2inact:←<br>Iω:none<br>ExTraFol:none | (Rhodes et al., 2005) |
| 1821 | $Na_v$1.1 | EIEE | R1636Q | R1821Q | $Na_v$1.1:R1636<br>$Na_v$1.1-2:R1625<br>$Na_v$1.1-3:R1608 | | Imax:none<br>τrec:↑<br>Ip:none<br>V1/2act:none<br>V1/2inact:none<br>Iω:none<br>ExTraFol:none | (Brunklaus et al., 2022) |

| MSA | Protein | Phenotype | Mutation | Mutation (MSA ID) | Isoform | GoF/ LoF | Gating Properties | References |
|---|---|---|---|---|---|---|---|---|
| 1821 | $Na_v1.5$ | RWS<br>LQT3<br>BRGDA1 | R1623Q | R1821Q | $Na_v1.5$:R1623<br>$Na_v1.5$-2:R1622<br>$Na_v1.5$-3:R1590<br>$Na_v1.5$-4:R1623<br>$Na_v1.5$-5:R1569<br>$Na_v1.5$-6:R1605 | GOF | Imax:none<br>τrec::none<br>τrec:↓<br>Ip:↑<br>V1/2act:none<br>V1/2inact:→<br>Iω:none<br>ExTraFol:none | (Rajamani et al., 2009)<br>(Philippaert et al., 2021)<br>(L. Q. Chen et al., 1996) |
| 1821 | $Na_v1.3$ | EPI<br>PMG | R1621Q | R1821Q | $Na_v1.3$:R1621<br>$Na_v1.3$-2:R1572<br>$Na_v1.3$-3:R1621<br>$Na_v1.3$-4:R1572 | GOF | Imax:none<br>τrec:none<br>Ip:↑<br>V1/2act:←<br>V1/2inact:none<br>Iω:none<br>ExTraFol:none | (Zaman et al., 2020) |
| 1821 | $Na_v1.6$ | DEE13 | R1617Q | R1821Q | $Na_v1.6$:R1617<br>$Na_v1.6$-2:R1617<br>$Na_v1.6$-3:R1628<br>$Na_v1.6$-4:--<br>$Na_v1.6$-5:R1576 | GOF | Imax:none<br>τrec:none<br>Ip:↑<br>V1/2act:←<br>V1/2inact:→<br>Iω:none<br>ExTraFol:none | (Wagnon et al., 2016) |
| 1824 | $Na_v1.4$ | SCM<br>HOKPP2 | R1451L | R1824L | $Na_v1.4$:R1451 | LOF | Imax:↓<br>τrec:↓<br>Ip:none<br>V1/2act:none<br>V1/2inact:←<br>Iω:none<br>ExTraFol:none | (Poulin et al., 2018)<br>(Luo et al., 2018) |
| 1824 | $Na_v1.6$ | ID | R1620L | R1824L | $Na_v1.6$:R1620<br>$Na_v1.6$-2:R1620<br>$Na_v1.6$-3:R1631<br>$Na_v1.6$-4:--<br>$Na_v1.6$-5:R1579 | LOF | Imax:↓<br>τrec:↑<br>Ip:none<br>V1/2act:none<br>V1/2inact:←<br>Iω:none<br>ExTraFol:none | (Y. Liu et al., 2019) |
| 1830 | $Na_v1.4$ | CMS16 | R1457H | R1830H | $Na_v1.4$:R1457 | LOF | Imax:none<br>τrec:↓<br>Ip:none<br>V1/2act:none<br>V1/2inact:←<br>Iω:none<br>ExTraFol:none | (Arnold et al., 2015) |

| MSA | Protein | Phenotype | Mutation | Mutation (MSA ID) | Isoform | GoF/ LoF | Gating Properties | References |
|---|---|---|---|---|---|---|---|---|
| 1830 | Na$_v$1.5 | SIDS<br>BRGDA1 | R1632H | R1830H | Na$_v$1.5:R1632<br>Na$_v$1.5-2:R1631<br>Na$_v$1.5-3:R1599<br>Na$_v$1.5-4:R1632<br>Na$_v$1.5-5:R1578<br>Na$_v$1.5-6:R1614 | LOF | Imax:none<br>τrec:↓<br>Ip:none<br>V1/2act:none<br>V1/2inact:←<br>Iω:none<br>ExTraFol:none | (Benson et al., 2003)<br>(Gui et al., 2010)<br>(Glazer et al., 2020) |
| 1842 | Na$_v$1.1 | GEFSP2 | R1657C | R1842C | Na$_v$1.1:R1657<br>Na$_v$1.1-2:R1646<br>Na$_v$1.1-3:R1629 | LOF | Imax:↓<br>τrec:↑<br>Ip:none<br>V1/2act:→<br>V1/2inact:none<br>Iω:none<br>ExTraFol:none | (Lossin et al., 2003) |
| 1842 | Na$_v$1.3 | DEE62 | R1642C | R1842C | Na$_v$1.3:R1642<br>Na$_v$1.3-2:R1593<br>Na$_v$1.3-3:R1642<br>Na$_v$1.3-4:R1593 | | Imax:↓<br>τrec:none<br>Ip:none<br>V1/2act:none<br>V1/2inact:none<br>Iω:none<br>ExTraFol:none | (Zaman et al., 2018) |
| 1842 | Na$_v$1.5 | RWS<br>LQT3<br>BRGDA1 | R1644C | R1842C | Na$_v$1.5:R1644<br>Na$_v$1.5-2:R1643<br>Na$_v$1.5-3:R1611<br>Na$_v$1.5-4:R1644<br>Na$_v$1.5-5:R1590<br>Na$_v$1.5-6:R1626 | LOF | Imax:none<br>τrec:↑<br>Ip:none<br>V1/2act:→<br>V1/2inact:none<br>Iω:none<br>ExTraFol:none | (Kroncke et al., 2018) |
| 1842 | Na$_v$1.6 | NDD | R1638C | R1842C | Na$_v$1.6:R1638<br>Na$_v$1.6-2:R1638<br>Na$_v$1.6-3:R1649<br>Na$_v$1.6-4:--<br>Na$_v$1.6-5:R1597 | LOF | Imax:none<br>τrec:none<br>Ip:none<br>V1/2act:→<br>V1/2inact:none<br>Iω:none<br>ExTraFol:none | (Wengert, Tronhjem, et al., 2019) |
| 1846 | Na$_v$1.1 | DRVT | F1661S | F1846S | Na$_v$1.1:F1661<br>Na$_v$1.1-2:F1650<br>Na$_v$1.1-3:F1633 | | Imax:↓<br>τrec:↑<br>Ip:↑<br>V1/2act:none<br>V1/2inact:→<br>Iω:none<br>ExTraFol:↓ | (Rhodes et al., 2004)<br>(Thompson et al., 2012) |

| MSA | Protein | Phenotype | Mutation | Mutation (MSA ID) | Isoform | GoF/ LoF | Gating Properties | References |
|---|---|---|---|---|---|---|---|---|
| 1846 | $Na_v$1.4 | PAM<br>PMC | F1473S | F1846S | $Na_v$1.4:F1473 | GOF | Imax:none<br>τrec:↑<br>Ip:↑<br>V1/2act:none<br>V1/2inact:→<br>Iω:none<br>ExTraFol:none | (Fleischhauer et al., 1998)<br>(Groome et al., 2008) |
| 1859 | $Na_v$1.1 | DRVT | G1674R | G1859R | $Na_v$1.1:G1674<br>$Na_v$1.1-2:G1663<br>$Na_v$1.1-3:G1646 | LOF | Imax:↓<br>τrec:none<br>Ip:none<br>V1/2act:none<br>V1/2inact:none<br>Iω:none<br>ExTraFol:none | (Rhodes et al., 2004) |
| 1859 | $Na_v$1.5 | BRGDA1 | G1661R | G1859R | $Na_v$1.5:G1661<br>$Na_v$1.5-2:G1660<br>$Na_v$1.5-3:G1628<br>$Na_v$1.5-4:G1661<br>$Na_v$1.5-5:G1607<br>$Na_v$1.5-6:G1643 | LOF | Imax:↓<br>τrec:none<br>Ip:none<br>V1/2act:none<br>V1/2inact:none<br>Iω:none<br>ExTraFol:none | (Glazer et al., 2020) |
| 1962 | $Na_v$1.5 | RWS<br>LQT3<br>SIDS | V1763M | V1962M | $Na_v$1.5:V1763<br>$Na_v$1.5-2:V1762<br>$Na_v$1.5-3:V1730<br>$Na_v$1.5-4:V1763<br>$Na_v$1.5-5:V1709<br>$Na_v$1.5-6:V1745 | GOF | Imax:none<br>τrec:none<br>Ip:↑<br>V1/2act:←<br>V1/2inact:→<br>Iω:none<br>ExTraFol:none | (Chang et al., 2004) |
| 1962 | $Na_v$1.4 | PAM<br>PMC | V1589M | V1962M | $Na_v$1.4:V1589 | GOF | Imax:none<br>τrec:none<br>Ip:none<br>V1/2act:none<br>V1/2inact:→<br>Iω:none<br>ExTraFol:none | (Mitrovié et al., 1994) |
| 1965 | $Na_v$1.6 | DEE13<br>EIEE | M1760I | M1965I | $Na_v$1.6:M1760<br>$Na_v$1.6-2:M1760<br>$Na_v$1.6-3:M1771<br>$Na_v$1.6-4:--<br>$Na_v$1.6-5:M1719 | GOF | Imax:none<br>τrec:none<br>Ip:none<br>V1/2act:←<br>V1/2inact:none<br>Iω:none<br>ExTraFol:none | (Y. Liu et al., 2019) |

| MSA | Protein | Phenotype | Mutation | Mutation (MSA ID) | Isoform | GoF/ LoF | Gating Properties | References |
|---|---|---|---|---|---|---|---|---|
| 1965 | $Na_v$1.3 | EPI | M1765I | M1965I | $Na_v$1.3:M1765<br>$Na_v$1.3-2:M1716<br>$Na_v$1.3-3:M1765<br>$Na_v$1.3-4:M1716 | GOF | Imax:↑<br>τrec:none<br>Ip:↑<br>V1/2act:←<br>V1/2inact:none<br>Iω:none<br>ExTraFol:none | (Zaman et al., 2020) |
| 1973 | $Na_v$1.5 | RWS | N1774D | N1973D | $Na_v$1.5:N1774<br>$Na_v$1.5-2:N1773<br>$Na_v$1.5-3:N1741<br>$Na_v$1.5-4:N1774<br>$Na_v$1.5-5:N1720<br>$Na_v$1.5-6:N1756 | GOF | Imax:↑<br>τrec:none<br>Ip:↑<br>V1/2act:←<br>V1/2inact:none<br>Iω:none<br>ExTraFol:none | (Kato et al., 2014)<br>(Hirose et al., 2020) |
| 1973 | $Na_v$1.6 | DEE13 | N1768D | N1973D | $Na_v$1.6:N1768<br>$Na_v$1.6-2:N1768<br>$Na_v$1.6-3:N1779<br>$Na_v$1.6-4:--<br>$Na_v$1.6-5:N1727 | GOF | Imax:↓<br>τrec:none<br>Ip:↑<br>V1/2act:none<br>V1/2inact:→<br>Iω:none<br>ExTraFol:none | (Veeramah et al., 2012)<br>(Wengert, Saga, et al., 2019) |
| 2077 | $Na_v$1.6 | DEE13 | R1872Q | R2077Q | $Na_v$1.6:R1872<br>$Na_v$1.6-2:R1872<br>$Na_v$1.6-3:R1883<br>$Na_v$1.6-4:--<br>$Na_v$1.6-5:R1831 | GOF | Imax:↑<br>τrec:none<br>Ip:↑<br>V1/2act:←<br>V1/2inact:→<br>Iω:none<br>ExTraFol:none | (Pan & Cummins, 2020) |
| 2077 | $Na_v$1.2 | NEIDEE<br>DEE11 | R1882Q | R2077Q | $Na_v$1.2:R1882<br>$Na_v$1.2-2:R1882 | GOF | Imax:↑<br>τrec:none<br>Ip:↑<br>V1/2act:←<br>V1/2inact:→<br>Iω:none<br>ExTraFol:none | (Schwarz et al., 2015)<br>(Mason et al., 2019)<br>(Berecki et al., 2018)<br>(Mason et al., 2019)<br>(Wolff et al., 2017)<br>(Thompson et al., 2020) |

Table S5: Common mutations with phenotypical GoF/LoF

| # | Mutation (MSA ID) | Na$_v$1.1 | Na$_v$1.2 | Na$_v$1.3 | Na$_v$1.4 | Na$_v$1.5 | Na$_v$1.6 | Na$_v$1.7 | Na$_v$1.8 | Na$_v$1.9 |
|---|---|---|---|---|---|---|---|---|---|---|
| 1 | R22K | GoF | GoF | | | | | | | |
| 2 | I71V | | | | GoF | | GoF | | | |
| 3 | Y91C | MIX | | | | LoF | | | | |
| 4 | F97S | LoF | | | | LoF | | | | |
| 5 | R108W | MIX | | | | LoF | | | | |
| 6 | R108Q | LoF | | | | LoF | | | | |
| 7 | M142T | LoF | | | | LoF | | | | |
| 8 | M142I | | | | | MIX | GoF | | | |
| 9 | E169Q | GoF | | | | LoF | | | | |
| 10 | C190R | LoF | | | | LoF | | | | |
| 11 | D205Y | GoF | | | | LoF | | | | |
| 12 | A212V | | | | GoF | LoF | | | LoF | |
| 13 | V224A | LoF | GoF | | | | GoF | | | |
| 14 | R231G | | GoF | | | | GoF | | | |
| 15 | R234Q | | GoF | | GoF | | | | | |
| 16 | R234W | | | | GoF | LoF | | | | |
| 17 | S240P | LoF | | | | | GoF | | | |
| 18 | A251T | LoF | GoF | | | | | | | |
| 19 | L275V | GoF | | | GoF | | | | | |
| 20 | Q279K | | | | GoF | LoF | | | | |
| 21 | R291H | | | | GoF | LoF | | | | |
| 22 | P415T | LoF | | | LoF | | | | | |
| 23 | R434H | | LoF | | | LoF | | | | |
| 24 | M436K | LoF | | | | LoF | | | | |
| 25 | R450C | MIX | | | | | | LoF | | |
| 26 | N473S | | GoF | | | LoF | | | | |
| 27 | L476P | | | | GoF | | | | | GoF |
| 28 | V478M | LoF | | | GoF | | | | | |
| 29 | V478L | | GoF | | GoF | | | | | |
| 30 | V479A | LoF | GoF | | | | | | | |
| 31 | R688C | GoF | GoF | | | | | | | |

| | | | | | | | | | | |
|---|---|---|---|---|---|---|---|---|---|---|
| 32 | D721N | GoF | | | | | | GoF | | |
| 33 | N723S | | | | GoF | | GoF | | | |
| 34 | P848L | LoF | | | | LoF | | | | GoF |
| 35 | R941W | | | | GoF | | | | GoF | |
| 36 | F943S | | | | GoF | | GoF | | | |
| 37 | R944G | LoF | | | MIX | | | | | |
| 38 | R944Q | LoF | GoF | | | | | | | |
| 39 | R947G | LoF | | | GoF | | | | | |
| 40 | R947Q | | GoF | | GoF | LoF | GoF | | | |
| 41 | R947W | | | | GoF | MIX | | | | |
| 42 | I965T | | | GoF | GoF | | | | | |
| 43 | L972P | MIX | | | | LoF | | | | |
| 44 | V978L | LoF | GoF | | | | | | | |
| 45 | A987T | | | | GoF | | GoF | | | |
| 46 | R1020H | MIX | | | | LoF | | | | |
| 47 | R1020C | LoF | | | | LoF | | | | |
| 48 | H1028P | LoF | | | | LoF | | | | |
| 49 | R1035C | MIX | MIX | | | LoF | | | | |
| 50 | R1035H | LoF | LoF | | | LoF | | | | GoF |
| 51 | C1038S | LoF | | | | LoF | | | | |
| 52 | E1043K | LoF | | | GoF | LoF | | | | |
| 53 | F1078L | LoF | | | GoF | | | | | |
| 54 | L1083P | LoF | | | | | | | | GoF |
| 55 | N1090K | GoF | | | | | GoF | | | |
| 56 | R1110H | | | | MIX | LoF | GoF | | | |
| 57 | R1123H | | GoF | | | | | GoF | | |
| 58 | E1399K | LoF | GoF | | | | | | | |
| 59 | Y1419H | | GoF | | | LoF | | | | |
| 60 | R1423Q | MIX | | | | LoF | | | | |
| 61 | D1434N | | | | MIX | LoF | | | | |
| 62 | D1466N | LoF | | | | MIX | | | | |
| 63 | L1505F | | GoF | | | | GoF | | | |
| 64 | R1507Q | | GoF | | GoF | | | | | |

| | | | | | | | | |
|---|---|---|---|---|---|---|---|---|
| 65 | V1514G | LoF | | | | LoF | | |
| 66 | A1521S | GoF | | | | | GoF | |
| 67 | P1523S | GoF | | | MIX | | | |
| 68 | P1523L | LoF | | GoF | GoF | LoF | | |
| 69 | I1525V | | | | GoF | | GoF | |
| 70 | V1599M | | | | GoF | LoF | | |
| 71 | Y1603C | LoF | | | | LoF | | |
| 72 | K1613E | | | | | LoF | GoF | |
| 73 | G1614R | LoF | | | | LoF | | |
| 74 | R1626G | | GoF | | | LoF | | |
| 75 | P1632L | LoF | | | | LoF | | |
| 76 | P1632S | LoF | | | | LoF | | |
| 77 | E1635K | MIX | | | GoF | | | |
| 78 | Y1643C | LoF | | | | LoF | | |
| 79 | N1666K | | | | GoF | | GoF | |
| 80 | I1679N | | | | GoF | | | GoF |
| 81 | M1681V | GoF | GoF | | | | GoF | |
| 82 | E1742K | LoF | | | | LoF | | |
| 83 | F1765L | | GoF | | | LoF | | |
| 84 | R1777H | GoF | | | | LoF | | |
| 85 | R1777C | LoF | | | GoF | LoF | | |
| 86 | D1789N | | | | MIX | MIX | | |
| 87 | V1792L | | GoF | | | | GoF | |
| 88 | V1793I | LoF | | | GoF | | | |
| 89 | L1805P | GoF | | | GoF | | | |
| 90 | V1815M | LoF | | | MIX | | | |
| 91 | T1818M | | | | GoF | LoF | | |
| 92 | R1821Q | GoF | GoF | | | LoF | GoF | |
| 93 | R1821L | | | | GoF | | GoF | |
| 94 | I1823T | LoF | GoF | | | | | |
| 95 | R1824H | | GoF | | GoF | | | |
| 96 | R1824L | | GoF | | MIX | | | |
| 97 | R1830H | | | | LoF | LoF | | |

| | | | | | | | | | | |
|---|---|---|---|---|---|---|---|---|---|---|
| 98 | R1833C | LoF | GoF | | | | GoF | | | |
| 99 | R1842C | | | GoF | | LoF | | | | |
| 100 | L1844P | GoF | GoF | | | | | | | |
| 101 | F1846S | LoF | | | GoF | | | | | |
| 102 | A1847V | LoF | | | | LoF | | | | |
| 103 | M1849K | LoF | | | | | | GoF | | |
| 104 | A1854E | LoF | | | | | | GoF | | |
| 105 | G1859R | LoF | | | | LoF | | | | |
| 106 | D1887N | GoF | | | | MIX | | | | |
| 107 | G1910S | LoF | | | GoF | LoF | | | LoF | |
| 108 | D1912G | LoF | | | | LoF | | | | |
| 109 | C1926R | LoF | | | | LoF | | | | |
| 110 | G1942R | GoF | | | | LoF | | | | |
| 111 | M1965I | | | | GoF | | GoF | | | |
| 112 | A1968V | LoF | GoF | | | | | | | |
| 113 | A1968T | LoF | GoF | | | | | | | |
| 114 | V1969A | LoF | | GoF | | | | | | |
| 115 | T1978M | | | | | LoF | | | GoF | |
| 116 | Y1994H | | | | | LoF | | | | GoF |
| 117 | Q2006E | | | | GoF | | GoF | | | |
| 118 | D2051G | GoF | | | GoF | | | | | |
| 119 | R2077Q | | GoF | | | | GoF | | | |
| 120 | N2082S | | | | GoF | | GoF | | | |
| 121 | R2097H | | GoF | | | | GoF | | | |
| 122 | R2097C | | GoF | | | LoF | | | | |
| 123 | R2113H | LoF | GoF | | | | | | | |

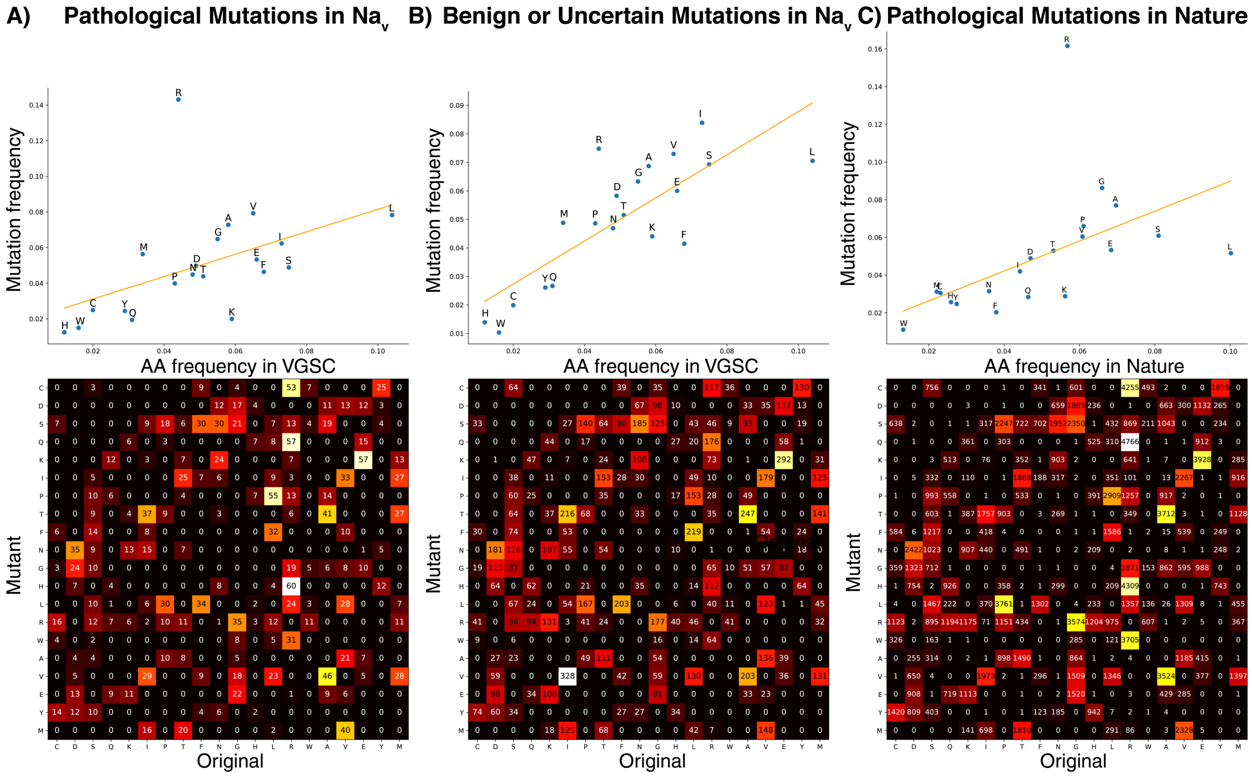

Figure S1: Amino acid change in pathological (A) and benign/uncertain (B) mutations in Na$_v$ and nature (C).
